# Using pose estimation and 3D rendered models to study leg-mediated self-righting by lanternflies

**DOI:** 10.1101/2023.02.06.527347

**Authors:** Theodore Bien, Benjamin H. Alexander, Chengpei Li, Natalie Goeler-Slough, S. Tonia Hsieh, Suzanne Amador Kane

**Author notes:** Contributed equally (Theodore Bien, Benjamin H. Alexander).

## Abstract

The ability to upright quickly and efficiently when overturned on the ground (terrestrial self-righting) is crucial for living organisms and robots. Previous studies have mapped the diverse behaviors used by various animals to self-right on different substrates, and proposed physical models to explain how body morphology can favor specific self-righting methods. However, to our knowledge no studies have quantified and modeled all of an animal’s limb motions during these complicated behaviors. Here, we studied terrestrial self-righting for nymphs of the invasive spotted lanternfly (*Lycorma delicatula*), an insect that must frequently recover from being overturned after jumping and falling in its native habitat. These nymphs self-righted successfully in 92-100% of trials on three substrates with different friction and roughness, with no significant difference in the time or number of attempts required. They accomplished this using three stereotypic sequences of movements. To understand these motions, we combined 3D poses tracked on multi-view high-speed video with articulated 3D models created using photogrammetry and Blender rendering software. The results were used to calculate the mechanical properties (e.g., potential and kinetic energy, angular speed, stability margin, torque, force, etc.) of these insects during righting trials. We used an inverted physical pendulum model (a “template”) to estimate the kinetic energy available in comparison to the increase in potential energy required to flip over. While these insects began righting using primarily quasistatic motions, they also used dynamic leg motions to achieve final tip-over. However, this template did not describe important features of the insect’s center of mass trajectory and rotational dynamics, necessitating the use of an “anchor” model comprising the 3D rendered body model and six articulated two-segment legs to model the body’s internal degrees of freedom and capture the role of the legs’ contribution to inertial reorientation. This anchor elucidated the sequence of highly coordinated leg movements these insects used for propulsion, adhesion, and inertial reorientation during righting, and how they frequently pivot about a body contact point on the ground to flip upright. In the most frequently used method, diagonal rotation, these motions allowed nymphs to spin their bodies to upright with lower force with a greater stability margin compared to the other less frequently-used methods. We provide a concise overview of necessary background on 3D orientation and rotational dynamics, and the resources required to apply these low-cost modeling methods to other problems in biomechanics.

## Introduction

Overturned insects benefit from being able to reassume an upright orientation quickly and with low metabolic energy investment, thereby mitigating the risk of predation, hunger, and desiccation. This behavior, which is called terrestrial self-righting when performed while on the ground, has been studied in insects (Full et al. 1995; Faisal and Matheson 2001; Li et al. 2019; Pace and Harris 2021; Zhang et al. 2021), spiders (O’Donnell 2018), frogs (Malashichev 2006), turtles (Domokos and Várkonyi 2007) and rats (Pellis et al. 1991). Animals use a rich, diverse range of body and appendage motions to rotate from supine to upright on the ground (Li et al. 2019). For example, (Frantsevich 2004) identified 20 different modes of self-righting in a study of 116 species of beetles. Ongoing research aims at understanding how the methods used by a specific organism to self-right depend on its morphology and locomotor abilities, as well as the substrate’s properties (Davis et al. 2011). Animal righting behaviors also have inspired designs for robots that can recover easily from upsets—a particular concern for operating in complex terrains (Saranli et al. 2004; Wang et al. 2022)—using coordinated motions of artificial legs (Saranli et al. 2004), wings (Li et al. 2016), and tails (Casarez and Fearing 2017).

Here we present the first study to our knowledge that measures and models the detailed motions of an animal’s body and all of its legs during terrestrial self-righting. We studied this behavior for nymphs of the spotted lanternfly (*Lycorma delicatula*), an invasive insect pest rapidly expanding its range in the U.S. (Urban and Leach 2022). Spotted lanternflies frequently jump and fall in their native habitats to disperse, evade predators, and respond to environmental disturbances (Kim et al. 2011). They therefore need to be able to self-right quickly and effectively under a variety of circumstances. In an earlier study, we found that falling spotted lanternfly nymphs are highly successful at self-righting both in the air and during landing (Kane et al. 2021). Here we turn our attention to how spotted lanternfly nymphs reorient to upright when starting from rest on their backs.

High-speed multi-angle video was used to record these insects as they attempted to self-right on several different substrates. These results were used to construct an ethogram of observed righting behaviors for this species and to determine how they depended on substrate and life stage. We also performed 3D tracking to quantify the detailed body and leg motions during righting (pose estimation). To interpret these data, we built on previous studies of terrestrial self-righting that considered how the animal’s gravitational potential energy and kinetic energy vary during righting (e.g., (Domokos and Várkonyi 2007; Li et al. 2019)). If an organism performs the motions required to self-right slowly enough that it remains approximately statically stable at every moment, it can be said to reorient quasistatically. Alternatively, it can use dynamic motions (inertial reorientation), e.g., by impulsively swinging an appendage (e.g., arm, leg or tail) so as to generate a reaction counter-torque on the body (Yeadon and King 2017). For a body in contact with the ground, such maneuvers generate ground reaction torques that can both enhance the ability to right and stabilize the final orientation after overturning (Frohlich 1979; Davis et al. 2011). Living organisms and robots can take advantage of combinations of quasistatic and dynamic righting strategies, so we considered both possibilities in our analysis. Earlier studies have proposed physical models with varying levels of detail (“templates and anchors” (Full and Koditschek 1999)) to determine whether animals meet the mechanical requirements for righting either quasi-statically or dynamically (Kessens et al. 2014; Li et al. 2016; Kessens and Dotterweich 2017; Othayoth and Li 2021). To this purpose, we created a detailed 3D rendered model of spotted lanternflies’ bodies using photogrammetry and Blender computer graphics software. We then used this model to calculate the mechanical properties required to test these predictions of righting strategies based on the tracked 3D kinematics of the insects’ body and legs measured from high-speed video.

This figure shows a scenario when the body and spatial frames do not coincide. F) Example of a 3D rendered model of a spotted lanternfly nymph used to estimate mechanical properties for modeling. G) Illustration of a 3D model of the spotted lanternfly nymph’s body in the orientation used to define the body (*xyz*) and spatial (*XYZ*) frames in the reference orientation, and the relationship between coordinate axes, roll, pitch and yaw rotations and Euler angles. (G) (H) Frame from the animation showing tracked landmarks (Fig. 1A) with the 3D body model and six jointed rod legs. (I-J) Animation frames showing examples of the overturned, flipping, and righted resting poses defined in the text.

**Fig. 1.**
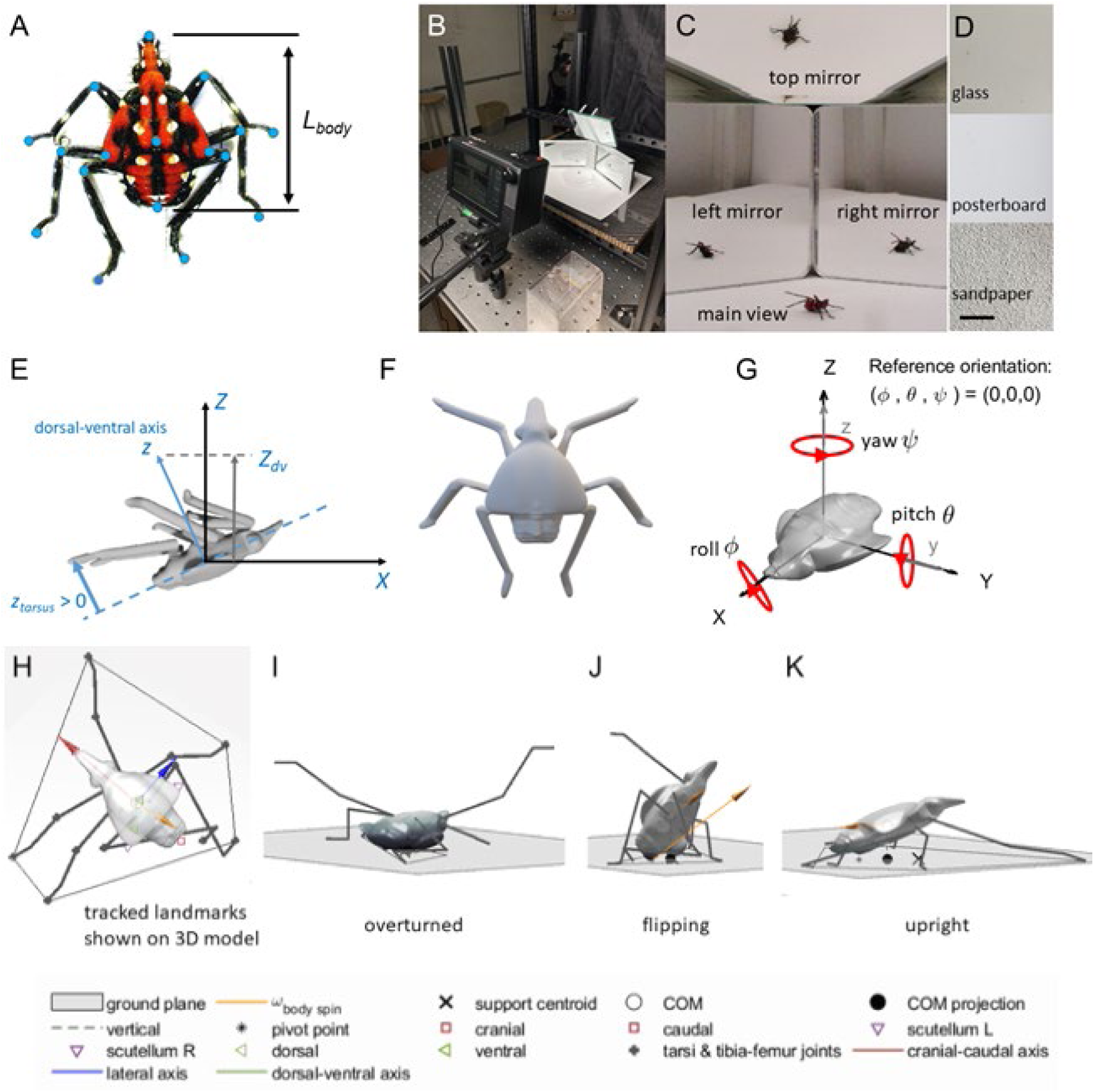
A) Fourth instar spotted lanternfly nymphs showing the definition of body length, *L_body_*, and the anatomical landmarks tracked on video (blue circles) B) Arena and video camera used to film terrestrial self-righting. C) Video frame showing the four camera views recorded of spotted lanternfly nymphs as they attempted to self-right. D) Substrates used in experiments. (Scale bar = 10 mm). E) Illustration of the geometry used to define the projection of the dorsal-ventral axis on the spatial Z direction used to quantify overturned vs righted orientation, −1 ≤ *Z_dv_* ≤ +1 and the tarsus (foot) *z*-position, *z_tarsus_*, in the body frame used to quantify foot motions.

## Materials and methods

### Data analysis and statistical methods

All image processing and data analysis were performed using MATLAB vR2023b (Mathworks, Natick MA, USA); specific MATLAB functions are given in italics. Results are reported as mean ± error bars based on instrumental uncertainties, unless noted otherwise.

### Insect collection, handling & morphometrics

Healthy, intact spotted lanternfly nymphs (Fig. 1A) were collected from tree of heaven (*Ailanthus altissima*) and wild grape vines (*Vitis vinifera*) in southeastern Pennsylvania (40°00’ 30.2” N; 75°18’ 22.0” W) in June 22-July 9, 2021. All specimens were collected mid-morning through mid-afternoon when they are most active (Kim et al. 2011). Experiments were performed in the laboratory at ambient conditions (23.2 [22.5, 24.1] °C, and relative humidity 62 [42, 70] % (mean, full range) within 5 minutes of collection to avoid specimen fatigue, hunger, or desiccation. To determine if self-righting behavior depended on life stage, we studied the two largest nymphal stages (3rd and 4th instars); earlier stages were too small for tracking on video. A combination of coloration and body morphometrics were used to determine the life stage of each specimen studied here, as previously described in (Bien et al. 2023). The body (head-caudal) length, *L_body_*, was measured to ± 0.05 mm using calipers and whole insect mass, *M*, was measured to ± 0.4 mg using an analytical balance (Fig. 1A), giving the following values: 3^rd^ instar: *L_body_* = 9.2 ± 0.7 mm, *M* = 31 ± 11 mg; 4^th^ instar: *L_body_* = 11.7 ± 0.8 mm, *M* = 59 ± 13 mg (mean ± SD).

### Self-righting experiments

Spotted lanternfly nymphs were recorded while attempting to self-right on flat, horizontal (± 0.05 deg) substrates made of posterboard, sandpaper and polished glass. (Fig. 1 B-D) Experiments were performed for 5 trials for each of 15 unique individuals for each of four different conditions (combination of instar and substrate considered); the limited number of trials per individual was chosen to minimize the effects of either learning or fatigue. This gave 299 total trials overall (75 trials/condition for all but the 4^th^ instars on sandpaper, for which one video was corrupted, so only 74 trials were analyzed).

At the start of each trial, we gently held a spotted lanternfly nymph for a few seconds to induce tonic immobility (Faisal and Matheson 2001). The nymph was then placed on its back on the substrate and allowed to attempt to self-right for ≤ 30 s. If the specimen failed to self-right in this time, the trial was terminated and recorded as a failure. An “attempt” at self-righting was defined as the time from when a specimen on its back began to lift its dorsum off the ground until it either turned upright successfully or else fell on its back after failing to right. Successful self-righting was defined as taking place at the moment when a specimen first came to rest stably on all of its feet with its dorsum uppermost. The mean attempt time was also measured for each specimen as the ratio of the time spent attempting to right divided by the number of attempts, including any time during which it was temporarily quiescent.

### Video analysis

High-speed videos of each trial were recorded using a Chronos 1.4 video camera (color, 1280 × 1024 pixels, 1000 fps; shutter speed 500 microsec, Krontech, Burnaby, BC, Canada) with a 12 mm focal length lens (f/# = 7, model M1214-MP2, Computar, Torrance, CA, USA). A total of four views (main camera and three mirror views) were filmed to allow 3D tracking of all body parts and legs with minimal occlusion. Data for 3D camera calibrations were obtained using wand calibration with easyWand software (wand score 0.64, reconstruction error 2.7-4.5 pixels, 7 pixel/mm) (Theriault et al. 2014) and the vertical direction was determined by tracking a falling ball. To approximate a natural light environment, the arena was illuminated with an LED light source (SL-200W, Godox, Shenzhen, China) covered by a diffusing softbox so as to provide 7,000 lux at the specimen, similar to the intensity measured in the understory of trees where the nymphs were collected.

Repeated viewings of all recorded videos were used to determine the stereotypic sequences of poses assumed while attempting to self-right, along with their frequency, success rate and timescales, the number of self-righting attempts, how the legs, feet, and body moved during different behaviors, and the behaviors associated with failed righting attempts. Note that we use “pose” to describe the body’s position and orientation in space combined with its posture (the detailed configuration of its leg joint angles and body parts.)

The insect’s pose was determined quantitatively using the program Direct Linear Transformation data viewer (DLTdv) (Hedrick 2008) to measure the image plane coordinates of multiple body anatomical landmarks. (Fig. 1A) These results were then used to compute the 3D *XYZ* coordinates of each landmark in the laboratory-fixed spatial frame. Manual tracking was necessary in most cases to measure the coordinates at sufficiently high resolution and accuracy to support 3D reconstruction of 16-18 body landmarks in four camera views for hundreds of frames in each video. Attempts to use automated tracking using DeepLabCut (Mathis et al. 2018) for this study were complicated by frequent occlusions, the small size of the features tracked, and the similar appearance of the legs and head; this resulted in points often failing to be tracked or to be tracked too inaccurately for 3D reconstruction, even after repeated rounds of supervised training. Custom MATLAB code was used to analyze the coordinates for velocity, angular velocity, kinetic energy, and potential energy as a function of time and body pose. Tracked coordinates and body orientation axes were smoothed by local regression using robust quadratic polynomial regression over a moving window of 25 ms (for slower-moving body parts) and 5 ms (for faster-moving legs) (MATLAB’s *smooth* with the “rloess” option, which ignores data that lie over 6 mean absolute deviations from the regression). Numerical derivatives were computed using quadratic polynomial regression using the same 25 ms time window (Li et al. 2023). We classified the rotational motions during self-righting using the orientation of the rotational axis in spatial and body fixed coordinates (defined in “Measuring and representing 3D orientation” below), the location of the pivot point about which rotations occur, and the distance, *d*, of the insect’s center of mass (COM) from the axis of rotation. The location of the *XY* ground plane on video was measured to 0.25 mm SD along *Z* by fitting a plane to a cloud of foot points visually determined to be in contact with the ground at a wide variety of locations throughout a trial.

### Substrate characterization

To test how surface properties affect terrestrial self-righting, we characterized the roughness of the three substrate materials used: posterboard for 3^rd^ and 4^th^ instars; and 80 grit sandpaper painted white and polished glass for 4^th^ instars only. (Fig. 1D) The feet of spotted lanternflies adhere to surfaces using a combination of their tarsal claws and adhesive pads (arolia) (Frantsevich et al. 2008). To characterize potential grasping by tarsal claws, we compared the feature sizes for each substrate with the claw dimensions of 3rd and 4th instar spotted lanternfly nymphs (> 1 µm radius tips and tip-tip spacing 660-849 µm) reported in (Avanesyan et al. 2019). The sizes of surface features of the sandpaper and posterboard were measured from micrographs using ImageJ (Schneider et al. 2012), and found to be large enough to allow nymphs to grasp the fibers of posterboard and sandpaper asperities using their tarsal claws. (Table 1) Polished glass, however, is reported to have sub-nm roughness values too small to afford a grip by these specimens (Yates and Duffy 2008). Combined with the fact that insect arolia adhere best to smooth substrates (Al Bitar et al. 2010), this suggests that spotted lanternfly nymphs should adhere to polished glass using their arolia, but not their claws.

**Table 1.** Substrate feature size and coefficients of static friction between the insect’s dorsum and substrate. ^§^ Roughness of polished glass from (Yates and Duffy 2008). (The frictional coefficient was so high during the sandpaper trials that only 2 trials out of 83 total resulted in sliding instead of rolling.)

| Substrate | Feature size | Coefficient of static friction relative to insect exoskeleton cuticle (mean ± SD) | N specimens (3rd, 4th instars); n trials |
| --- | --- | --- | --- |
| Posterboard | 40 ± 8 μm diameter fibers (median ± SD) | 0.36 ± 0.09 | 10 (5, 5); 63 |
| Glass | < 1 nm<sup>§</sup> | 0.55 ± 0.13 | 9 (5, 4); 73 |
| Sandpaper | 570 ± 100 μm diameter pores (mean ± SD) | $\geq 1.2 \pm 0.4$ | 2 (1, 1); 2 |

In addition to needing to hold onto and exert forces on the substrate, insects can rotate upright about a pivot point formed by a body part resting on the ground if frictional forces are sufficient at this contact region to prevent sliding. To characterize the friction between the exoskeleton and the substrate, freshly euthanized specimens were placed on their dorsum on an inclined plane tilted to find the minimum angle, *ψ*, at which sliding first occurs. The coefficient of static friction between the exoskeleton cuticle and substrate was then computed as: *µ_s_* = tan *ψ* (Halliday et al. 2013). While we were able to measure multiple instances of sliding on the inclined plane for glass and posterboard, the frictional force between the exoskeleton and sandpaper was so great that rolling occurred before sliding in only 2 of 83 inclined plane trials, so we were only able to determine a lower bound on the corresponding frictional coefficient. (Table 1)

Based on the combined effects of foot and back interactions, we expected that spotted lanternfly nymphs should right more effectively on sandpaper (which had the greatest frictional coefficient tested and substrate feature size amenable to tarsal claw grasping) than on lower-friction posterboard or glass.

### 3D rendered model and calculations of mechanical properties

We created 3D rendered models of spotted lanternfly nymphs to use in analyzing the tracked 3D coordinates. Photogrammetry and microscopy were used to determine the 3D geometry of the entire body and legs. Six 4^th^ instar nymphs also were dissected for use in separately measuring the dimensions and mass of the body and each leg. These data were used with Blender rendering software (www.blender.org, January 16, 2023) to create realistic 3D models of the spotted lanternfly nymph body; for details see (Li et al. 2023). (Fig. 1F) Three different 3D models were created to match the slightly different morphologies of the specimens analyzed in detail below to provide measures of uncertainty for the mechanical properties. (See Supplementary Dataset S1 for Blender files.) The 3D body models were used to create .stl files that contained the triangular mesh describing its surface geometry; these were used with the MATLAB package RigidBodyParams (Semechko 2023) to compute the body’s COM coordinates.

Mechanical properties during self-righting were computed using an “anchor” model in which the body (head, abdomen and thorax) was treated as a single rigid object represented by the 3D body model, and each leg was treated as an articulated assembly of two rigid rod-like segments (one segment each for the combined tibia and tarsus, and one for the combined femur and shorter coxa) based on the measured leg segment masses and lengths. The pose of the anchor’s body and legs at each timestep was determined using the results from 3D video tracking. (Fig. 1H) Because an earlier growth allometry study has shown that the body morphology of spotted lanternfly nymphs scales isometrically with body length (Bien et al. 2023), we analyzed the mechanical properties of tracked data for different specimens using 3D models scaled to a standardized body length and mass as follows. First, we multiplied the tracked coordinates for each trial analyzed by a scaling factor so the specimen’s body length in rescaled coordinates corresponded to a standardized value of *L_body_*= 8.9 mm. The rescaled coordinates were then analyzed in combination with a 3D model with this body length assuming a whole insect mass of 28.4 mg typical of this body length.

### Measuring & representing 3D orientation

We now consider how to represent orientation in 3D (attitude) during self-righting. In this study, attitude was defined as the 3D orientation of *xyz* axes defined by the insect’s anatomical axes (cranial-caudal, lateral, and dorsal-ventral, respectively), in a body frame of reference with respect to the *XYZ* axes of the laboratory-fixed spatial frame. (Fig. 1G) Attitude coordinates describe an ordered sequence of rotations that transform the body frame from a reference orientation to the orientation of interest with respect to the spatial frame. We use Euler angle attitude coordinates *ψ*, *θ*, *φ* (roll, pitch, yaw), which describe rotations about each of the *xyz* body axes, respectively. In the reference orientation used here, (*ψ*, *θ*, *φ*) *=* (0,0,0) corresponded to the body frame *xyz* axes aligned with the spatial frame *XYZ* axes. (Fig. 1G) The rotations used to relate the Euler angles to attitude were performed in as the order yaw, pitch, roll (Tait-Bryan convention, *zy’x’’* order), where, e.g., y’ refers to the new orientation of the pitch (lateral) axis after the first yaw rotation by *φ* about z, and x’’ refers to the direction of the cranial-caudal axis for roll rotations after rotation by *θ* about the y’-axis. (Fig. S1A) Note that if the rotations are performed in a different order, this results in a different 3D orientation (Schaub and Junkins 2003). (Fig. S1B) This also means that, in general, the Euler angles (*ψ*, *θ*, *φ*) do not correspond to the angles between the final *x’’’ y’’’ z’’’* body axis orientation and the *XYZ* spatial axes. (Fig. S1C) We therefore reported the *Z*-component of the body dorsal-ventral axis (−1 ≤ *Z_dv_* ≤ +1) in the spatial frame (Fig. 1E) as a measure of the body’s orientation during righting. When the insect is overturned and flat on its back, then *Z_dv_* = +1, and when it is upright with its dorsal-ventral axis pointed downward along the vertical, *Z_dv_* = −1. (Fig. 1I, K) We refer to the pose when *Z_dv_* = 0 as the flipping point. (Fig. 1J)

To avoid mathematical ambiguities that can arise when using Euler angles for calculating rotations (e.g., gimbal lock), we first performed calculations on orientations computed from the tracked coordinates using quaternion-based functions in MATLAB, and then converted the result to Euler angles (Wang et al. 2023). Given the measured 3D orientations of the body and appendages in the spatial frame at times *t* and *t* + *Δt*, we used MATLAB functions *absor* (Matt 2024) and *angvel* to compute the corresponding angular velocity vector, 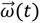, of each object. The rotation axis, 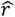, rotation angle, Δ*θ*, and angular speed, *ω*, in rad s^−1^ then were computed using 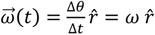, and the angular acceleration, *α*, was calculated as 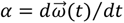.

### Stability metrics

The stability margin of poses during locomotion can be computed as a test for static mechanical stability (McGhee and Frank 1968; Ting et al. 1994). To this purpose, we used custom MATLAB code and the tracked insect body coordinates to compute the support polygon (the convex hull formed by all body parts in contact with the ground). (Fig. 2A) A tracked leg segment or point on the body was defined as contacting the ground if it lay within a distance comparable to the tracking uncertainty of the ground plane. The stability margin (*SM*) then was computed as the shortest distance from the boundary of the support polygon to the projection of the insect’s COM on the ground. If only pushing forces are exerted by the organism on the ground, then *SM > 0* for stable equilibrium, *SM* = 0 for metastable equilibrium, and *SM* < 0 for unstable poses. The value of *SM* is at its maximum (the ideal stability margin, *ISM*) when the projected COM agrees with the centroid of the support polygon. The ratio *SM*:*ISM* ≤ 100% then measures how close the insect’s pose is to optimal static stability. However, for insects that can exert pulling forces on the ground, either through adhesion with their arolia or by grasping the ground with tarsal claws, static stability can be achieved for *any* value of *SM* given a sufficiently high pulling force.

**Fig. 2.**
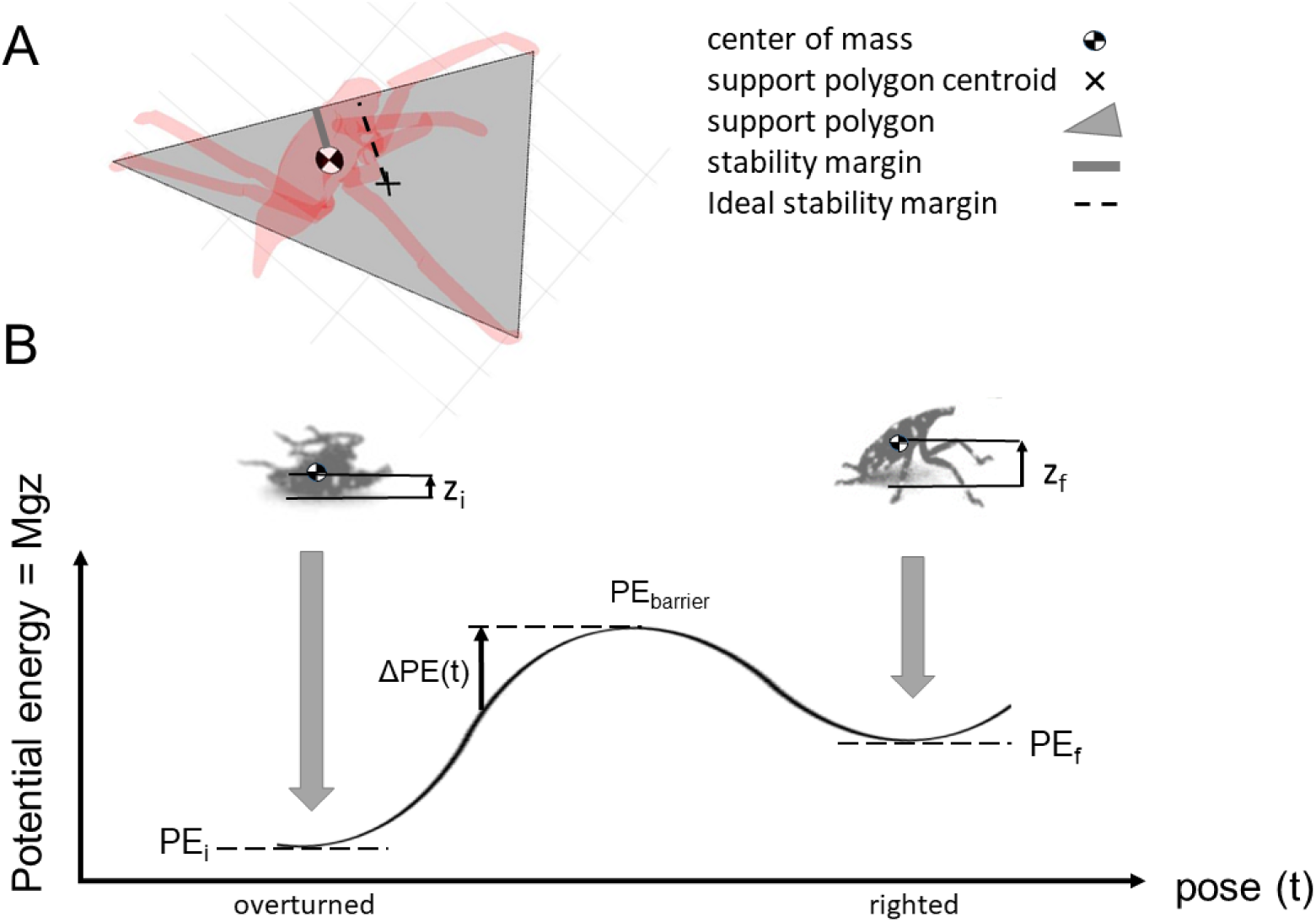
A) Top view of a 3D model of a spotted lanternfly nymph (red) showing the definition of the stability margin. The support polygon (gray) is the convex hull formed by all body parts in contact with the ground support. The stability margin (dashed line) is the shortest distance from the boundary of the support polygon to the projection of the COM on the ground, while the ideal stability margin (gray line) is the shortest distance from the support polygon’s in-plane centroid (×) to the boundary. (B) Schematic gravitational potential energy vs pose diagram showing the definition of the increase, *ΔPE*(*t*), in potential energy required to surmount the potential energy barrier starting at time *t*. Insets show sample images of a spotted lanternfly nymph in the overturned and righted poses. (Color online)

### Analyzing energy during righting

To determine to what extent an organism’s motions during righting contribute to inertial reorientation, we next consider how to compute its potential and kinetic energy. Many animals, including spotted lanternflies, must increase their gravitational potential energy (hereafter, potential energy or *PE*) between the initial overturned and final righted poses, as well as possibly overcoming one or more intervening *PE* maxima or “barriers” (*PE_barrier_*). (Fig. 2B) Following (Domokos and Várkonyi 2007; Othayoth et al. 2021), we computed *PE* = *MgZ* for each pose observed during self-righting, where *g* = 9.81 m s^−2^ is the acceleration of gravity and *Z* is the insect’s COM height relative to the ground. (Fig. 2B) The potential energy landscape for terrestrial self-righting is then the value of *PE* calculated for all possible poses consistent with the organism’s morphology and range of motion about each of its locomotor degrees of freedom that can result in a transition from overturned to upright (Othayoth et al. 2021). By analogy with free energy landscape models in physical chemistry, it has been proposed that self-righting by animals and robots can be modeled as a transition between the initial and final configurations, with the optimal trajectory in the *PE* landscape defined by following local PE basins so as to minimize any intervening energy barriers (Kessens et al. 2012; Li et al. 2019). However, in this study, the spotted lanternfly nymphs were observed to sometimes lift their bodies off the substrate and use all their legs during righting, so it was not feasible to compute *PE* for the full multidimensional configuration space during self-righting. Instead, we computed a simplified *PE* landscape for the 3D body model with one point of the body in contact with the ground for all possible values of pitch and roll to allow a comparison with measured values of body pitch and roll and with earlier studies of cockroach self-righting (Li et al. 2019).

Purely dynamic self-righting requires the organism to accumulate enough kinetic energy *KE*(*t*), to enable it to rotate upright while increasing its potential energy from the current value, *PE(t)*, to the maximum during righting, *PE_max_* (Robins et al. 1998; Kessens and Dotterweich 2017; Xuan and Li 2020). This requirement can be stated as *KE*(*t*) ≥ *ΔPE(t) = PE_max_ – PE(t)*. This time dependence indicates that the timing of a dynamic motion (e.g., a push that increases the body’s rotational *KE* or a rapid leg swing) determines the effectiveness of its contribution to inertial reorientation. For example, if an inverted pendulum is near the metastable apex of its trajectory, at which *ΔPE ≈ 0,* then *any* nonzero *KE* for rotations with the correct direction result in tip-over; farther from the apex, the inverted pendulum requires a greater kinetic energy to overcome the *ΔPE* required to reach the apex and tip it over. In addition, not all contributions to an object’s kinetic energy are available for conversion to work against gravity so as to promote righting. For example, the kinetic energy associated with rotations about a vertical axis or with horizontal translational motions is irrelevant for righting. For this reason, we define an available kinetic energy, *KE_avail_*(*t*), that can contribute to righting rotations and discuss below how to calculate it after developing the physical models used to interpret our tracked data.

To describe the contribution of inertial reorientation to righting at time *t*, we define a dimensionless righting number, *RNtt*) = *KE_avail_*(*t*)/Δ*PE*(*t*). If *RN*(*t*) ≥ 1, then the object has enough available kinetic energy to right at time *t* without exerting any additional torques (purely dynamic righting). At the other limit, *RN* << 1, righting is purely quasistatic. If *RN* < 1 but not << 1, *RN* quantifies how much the available kinetic energy contributes to inertial reorientation by reducing the ground reaction torques required to right.

### Rotational kinematics and dynamics

Using tracked coordinates and the 3D anchor model, we also computed a variety of measures for interpreting the rotational kinematics and dynamics during righting. We first briefly review some necessary background; for a full treatment of the relevant topics in advanced classical mechanics see (Goldstein 1980). An object’s angular acceleration, α, depends on net torque (moment of force), *τ_net_*, as (Fig. 3A):

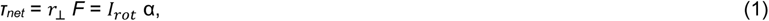

where *F* is force, *r*_⟂_ is the magnitude of the moment arm (the vector from the point at which the force acts to the axis of rotation) perpendicular to the force, and the moment of inertia, *I_rot_*, defines resistance to changes in angular velocity (Goldstein 1980). An object’s moment of inertia, *I_rot_*, depends on both the axis of rotation and its relation to the distribution of its mass, *M*, via (Fig. 3B):

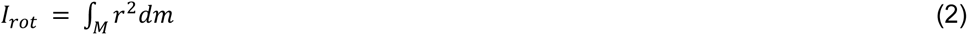

**Fig. 3.**
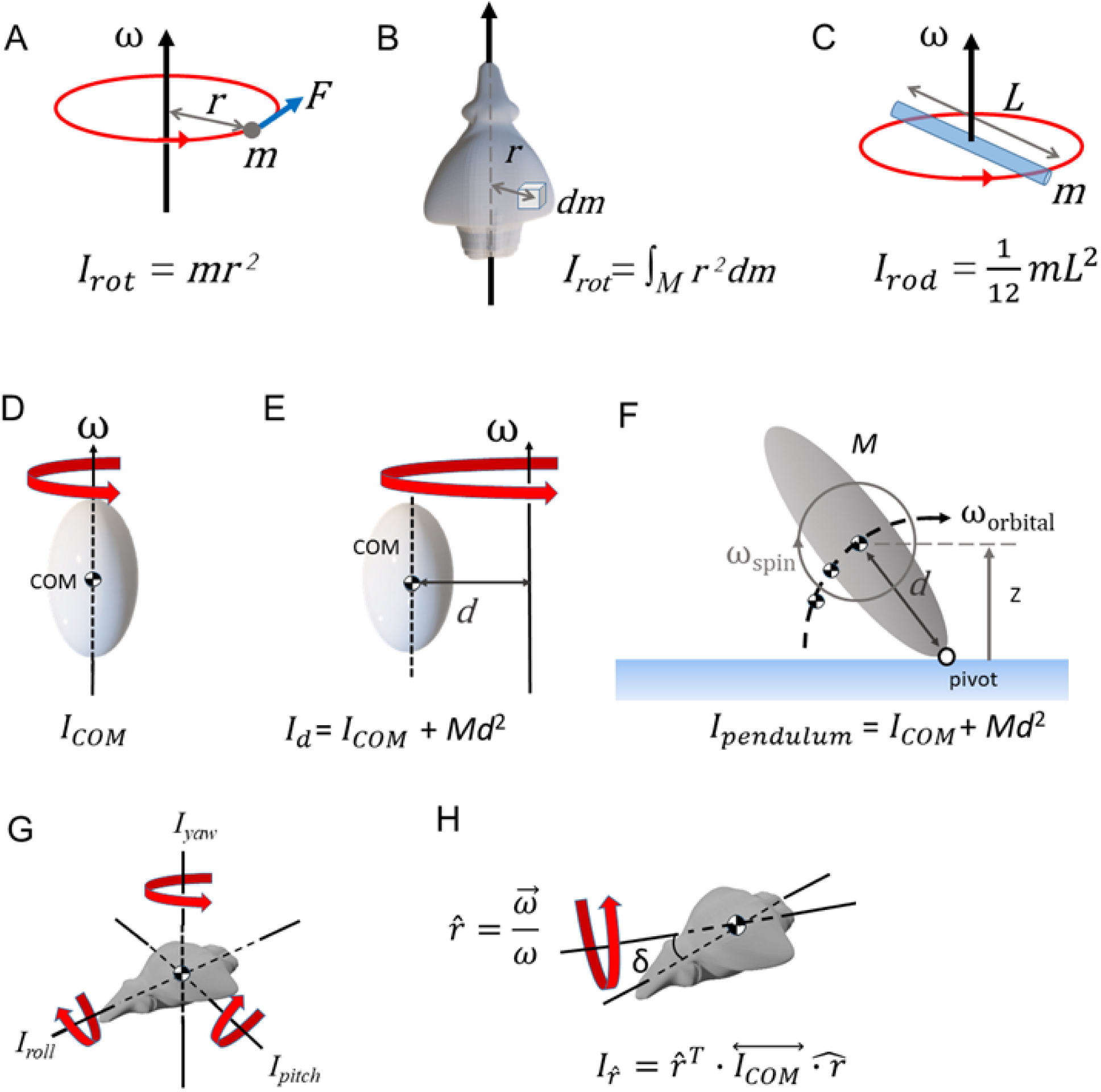
A) Geometry used in defining angular velocity, ω, force, *F*, moment arm, *r*, and the moment of inertia, *I_rot_*, for a point mass, *m*. Geometry for defining moment of inertia for the general case of (B) a distributed mass and C) a rod rotating about its center of mass (COM). The moment of inertia depends on the axis of rotation about the COM (D) and the distance, *d*, from the COM to the axis of rotation (E), as in the case of an inverted physical pendulum (F). The geometry for defining spin rotations about the COM and orbital motion of the COM is also shown in F. (G) Moments of inertia about each principal axis for the case where they correspond to the body’s roll, pitch and yaw axes. (H) Calculation of the moment of inertia about an arbitrary axis of rotation, 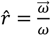. (Color online)

Note that *I_rot_* is computed in the body frame of the object, independent of its orientation in the spatial frame. For a rod with mass *m* and length *L* rotating about an axis passing through its COM, Equation (2) gives 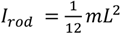. (Fig. 3C) We used this relationship to compute the moment of inertia of each of the two segments used to model each leg in the 3D model. We also needed to account for the distance between the COM of each segment and the whole insect COM using the parallel axis theorem. This states that, given the moment of inertia, *I_COM_*, for rotations about an axis that passes through an object’s COM, the value for rotations about the same axis displaced a distance *d* from the COM, *I_d_*, is (Fig. 3E, F):

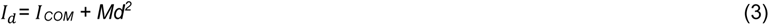

The tracked body coordinates were used with Equation 3 to determine the relevant geometry for computing each leg segment’s moment of inertia at a given time.

Insect bodies have complex shapes that influence their 3D rotational dynamics. In the most general case, the moment of inertia for rotations about the COM of a 3D object is a 3×3 matrix, the inertia tensor 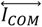. There always exists a body coordinate frame in which the inertia tensor is a diagonal matrix, 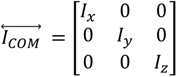, in which the *xyz* axes are called the principal axes and the object can undergo stable rotations for zero net torque about the *x, y*, or *z* principal axis with moment of inertia, *I_x_*, *I_y_*, or *I_z_*, respectively. For example, Fig. 3G shows the principal axes for the 3D model of the spotted lanternfly body, which happen to agree with the body frame’s roll, pitch, and yaw axes. In the most general case, the moment of inertia can be computed for rotation about any axis, 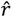, that passes through the object’s COM (Fig. 3H) as:

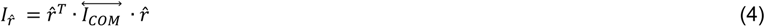

Values of the moment of inertia and principal axes for the spotted lanternfly’s body were computed using the MATLAB package RigidBodyParams (Semechko 2023) with the 3D body model. We then used Equations (3) and (4) with the body axes, calculated rotation axes, and pivot point to determine the moment of inertia of the body, which was added to the contributions of each leg segment to give the whole insect moment of inertia at each time step.

Following (Li et al. 2016), we also created a template that modeled the motion of the insect during righting as that of an inverted physical pendulum rotating about a fixed pivot point on the ground. (Fig. 3F) Nymphs indeed were observed to always maintained several points of no-slip contact with the ground during each successful righting attempt. This template also assumes that the relative motion of body and legs was slow enough to allow them to be approximated as a rigid object over times comparable to a few video frames. Using this model, we estimated the *KE_avail_*(*t*) for rotating the insect upright as the kinetic energy due to translation of its COM (orbital motion) and rotation about the COM (spin), assuming the same angular velocity for the orbital, *ω_orbital_*, and body roll and pitch spin, *ω_spin_*, motions. (Fig. 3F) For the inverted physical pendulum, this value of *KE_avail_* comprises the kinetic energy of all motions that could contribute to increasing the height of the COM, and hence increasing *PE*, while excluding the kinetic energy of motions that leave COM height unchanged (e.g., purely horizontal translations, rotations about a vertical axis, counter-rotations of two or more body components about the same axis with angular momenta that sum to zero, etc.) We computed for each time step the whole insect’s COM, its distance *d* from the pivot point, its COM speed, *ν_orbital_*, and the spin moment of inertia of the whole insect about its COM (Fig. 3D) This allowed us to compute the orbital angular velocity, *ω_orbital_* = *ν_orbital_*/*d*, and total moment of inertia about a pivot point on the ground a distance *d* away from the COM, *I_pendulum_* using Equation 3 (Halliday et al. 2013). The pendulum’s kinetic energy was then computed as 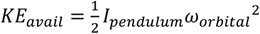.

The insect’s moment of inertia is necessary for computing its kinetic energy during righting, but it has other potential influences on righting success based on fundamental principles of rotational dynamics (Halliday et al. 2013). To see why, we note that an inverted physical pendulum with mass *M* has natural period, 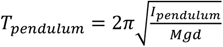 (Fig. 3F) This period sets the timescale for failure if the animal falls due to losing traction or to being unable to sustain sufficient force to stay raised, and the timescale for rotating upright from a given starting position. For example, gymnasts hold their arms outstretched when walking along a beam to increase the time allowed for making corrective motions to avoid falling. Furthermore, the inertial torque exerted on the body by corrective relative rotations of their arms is proportional to the arms’ moment of inertia, so increasing arm extension, and hence the moment of inertia, allows the gymnast to compensate for body tilting using smaller angular acceleration. In righting, the opposite is true: for an overturned animal, lower values of *I_pitch_* and *I_roll_* facilitate rotating upright more rapidly, while a larger value of *I_yaw_* allows for corrective motions to prevent yaw rotations, which are unhelpful in the overturned pose. Second, the time, *t_rot_*, required to rotate *any* object with moment of inertia *I_rot_* acted on by a constant torque, *τ*, through an angle *θ* is *θ* = ½ α *t_rot_* ^2^ = *½ τ/ II_rrrrtt_ t_rot_* ^2^, so 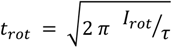. To compensate for an increase in moment of inertia, the animal therefore either must exert a greater torque to turn over in the same time, or exert the same torque for a longer duration. It can exert a greater torque by either exerting a greater force or by using a larger moment arm. However, the magnitude and duration of force production are limited because righting is a strenuous activity for insects (Full et al. 1995), and the maximum moment arm is limited by the length of the legs and their limited ranges of motion.

Using the template, we also estimated the net ground reaction torque acting on the insect with respect to the pivot point, *τ_net_* = *I_pendulum_α_orbital_*, where *α_orbital_* = *dω_orbital_*/*d*t is orbital angular acceleration. During self-righting, the forces acting on the insect include gravity, which acts only on the insect’s COM, and ground reaction forces at each contact between the body and ground. The net ground reaction torque for rotations about the pivot point is therefore:

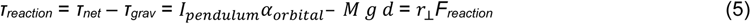

where *F_reaction_* is the net ground reaction force, and *r_COM_* and *r*_⟂_ are the moment arms for the force of gravity and the ground reaction force, respectively. We estimated *r_COM_* as the distance between the pivot point and the projection of the insect’s COM on the ground, and *r*_⟂_ using the mean distance from all tarsi in contact with the ground and the effective pivot point. We also computed the mean reaction force exerted by each leg (Full et al. 1995) by dividing the net force by the number of tarsi in contact with the ground (*n_tarsi_*) at each timestep: *F_leg_* = *F_reaction_* / *n_tarsi_* = *τ_reaction_* / (*n_tarsi_ r*_⟂_).

### Leg coordination

We also analyzed how leg motions might influence inertial reorientation. Because the relevant leg motions require changing the height of the tarsus above the body dorsal plane, we computed the value of *z_tarsus_* for each leg vs time for comparison with other measures during and after righting (Fig. 1E). Following, (Othayoth and Li 2021) the coordination between the z-positions of the *j*th and *k*th tarsi, z_j_ and z_k_, was analyzed using the discrete-time normalized cross-correlation, *z_j_* * *z_k_*(*t*_lag_), a type of inner product between two functions, one of which is offset in time from the other by *t_lag_* (the lag time):

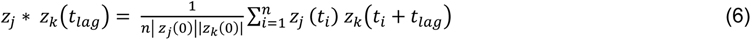

where the sum is performed over all n time samples (Press 2007). The magnitude (similarity) of the normalized cross-correlation of two signals is a measure of how correlated one signal remains with another when shifted by a positive or negative time offset called the lag time, where +1 is the maximum similarity possible for in-phase coordination, and −1 the maximum for out-of-phase motion. For example, the cross-correlation of any function with itself (its autocorrelation) is equal to 1 at zero lag time, while the autocorrelation of a random signal with zero mean decays rapidly to zero as lag time increases. (Fig. S6) The autocorrelation of a periodic signal has periodic peaks at nonzero lag times at integer multiples of the signal period, *T*_j_, with peak similarities that indicate the timescale over which the signal maintains correlation with itself. Fig. S6 illustrates this behavior using several examples: a sine wave has peaks with similarity = 1 at all values of *T*_j_, while a sine wave with added noise in its period, phase or amplitude has periodic peaks with similarities that decrease with increasing absolute lag time. If two sinusoidal signals have the same period but one is shifted by a constant phase offset, then their cross-correlation function has its first peak at a lag time equal to that corresponding to the phase offset.

We performed correlation analysis from the first frame analyzed to when the trajectory reached its apex using *xcorr* in MATLAB, after standardizing *z_tarsus_* to zero mean to remove trivial correlations between constant values. First, the normalized autocorrelation of *z_tarsus_* for each tarsus was computed as a function of lag time to measure the timescale over which each leg’s motion remained correlated, as well as its maximum peak similarity at nonzero lag time. Second, to determine the phase offset and maximum similarity of each pair’s motions, the cross-correlation function was measured between each pair of *z_tarsus_* values for the forelegs and midlegs (the hindlegs were not considered because they were primarily used for relatively static stabilizing, propelling or pulling motions). (Fig. S7-9)

## Results

### Terrestrial self-righting behaviors

Spotted lanternfly nymphs were found to self-right using a sequence of stereotypic motions. (Fig. 4A, Movie 1) Initially, the nymph lay flat on its back (overturned). Upon recovering from tonic immobility, the nymph moved its legs in probing motions in all possible directions (searching) in apparent attempts to locate the surface, similar to those described for beetles and spiders prior to righting (Frantsevich 2004; O’Donnell 2018). Once a foot made contact with the ground dorsally, the other legs were quickly elevated and extended so as to grasp the ground (planting). Next, the nymph primarily used its forelegs, with some assistance from the midlegs, to push against the ground so as to pitch and raise its cranial end off the ground (lifting). It then attempted to flip upright (righting) using a combination of propelling and swinging leg motions (Fig. 4A, Movie 1); we discuss more specifics of these movements below. During the post-tip-over recovery phase in successful trials, the nymph used its legs to achieve the pose typical of resting and walking (righted), in which its cranial-caudal axis was pitched upward from the horizontal. If self-righting was unsuccessful, the nymph toppled onto its side or dorsum (failed), after which it either rested or resumed searching.

**Fig. 4.**
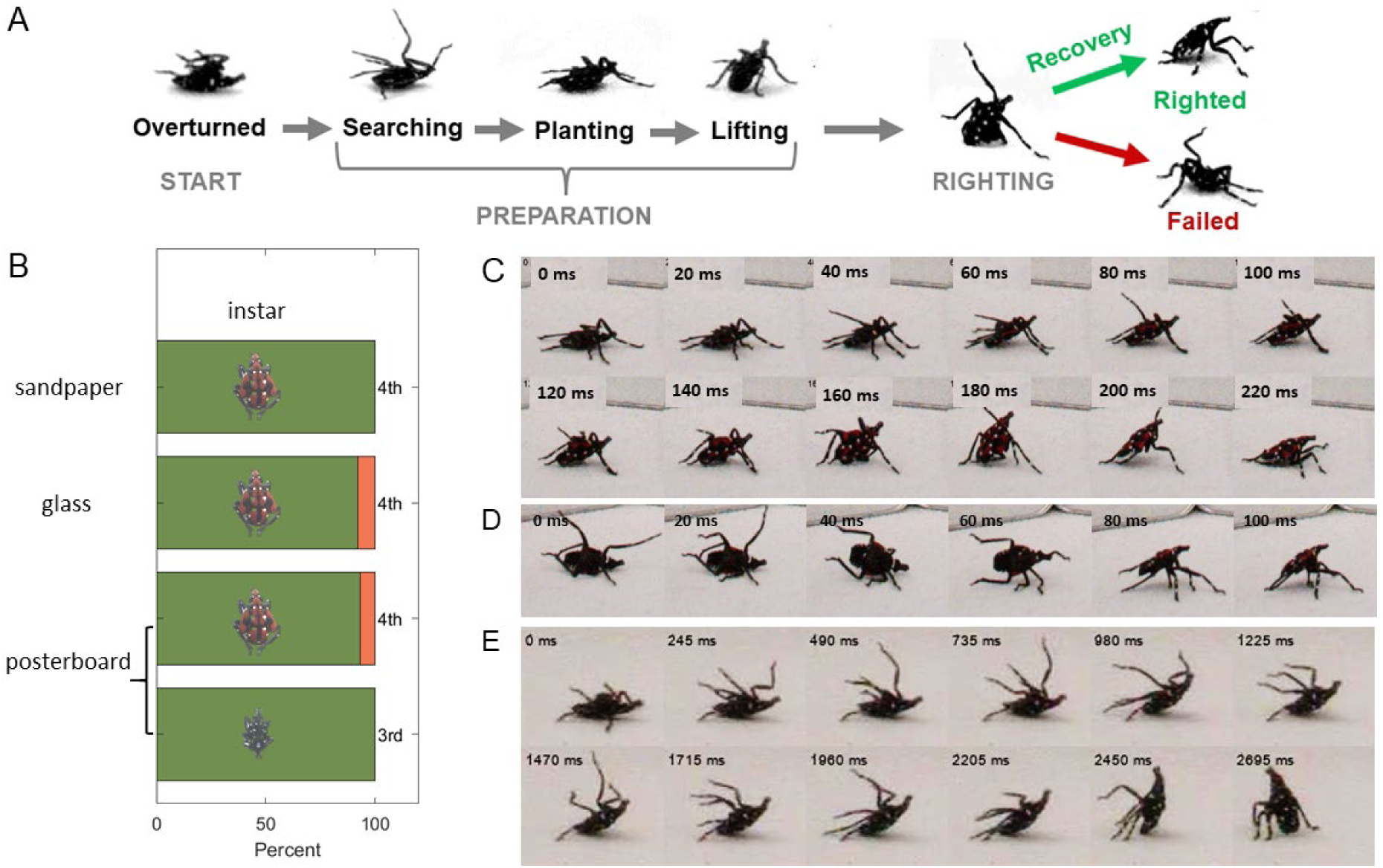
A) Ethogram showing stereotypic poses assumed by spotted lanternfly nymphs during terrestrial self-righting. B) Percent successful trials for each combination of substrate and life stage (instar) studied. (75 total trials per condition: N = 15, 5 trials/specimen). Sequences of closeup still images for (C) diagonal rotating about a pivot point on the caudal end (4^th^ instar on sandpaper), (D) lifted rotating with the legs but not the body contacting the substrate (4^th^ instar on sandpaper); and (E) pitching about a pivot point on the caudal end (3^rd^ instar on posterboard). (C-E) Times relative to the first image shown at top left in each frame. (Color online)

Spotted lanternfly nymphs self-righted successfully (92-100% all trials) on all three substrates for both life stages studied. (Fig. 4B; Table S1) We performed linear regression to test whether success rate, time/attempt and number of attempts were correlated with trial number to test the hypotheses that: 1) fatigue might result in poorer righting performance upon repeated attempts, or, conversely, that 2) learning from repeated attempts could improve righting outcomes. (Fig. S2A-C, Table S2) Only the number of attempts for the 3^rd^ instar nymphs on posterboard increased significantly with trial number. Given this result and the balanced study design, we consequently pooled data across trials and individuals before performing further tests. The percent of self-righting trials resulting in success did not depend significantly on the combination of instar and substrate (Fig. 4B; Table S3). All self-righting trials were successful for all 3^rd^ instar specimens for all three substrates and for all 4^th^ instar nymphs on sandpaper. All 4^th^ instars succeeded in the first trial and in at least 1 additional trial on posterboard and glass. The only trials in which a specimen failed to eventually self-right involved five failures by two 4^th^ instars on posterboard and five failures by five 4^th^ instars on posterboard or glass. Significance testing using Kruskal-Wallis revealed that the total time required to self-right, the time/attempt, and the number of attempts did not depend significantly on the combination of substrate and life stage. (Fig. S2D-F; Table S4)

Fig. 4C-E and Movie 1 show examples of the rotational body and leg motions used by spotted lanternfly nymphs for self-righting. In 96.7% (289:299) of all trials, this involved the nymph rotating its body upright about a rotational axis intermediate between pitching and rotating while pivoting about a contact point between its body and the ground (“diagonal rotating”). (Fig. 4C) During diagonal rotating, the hindlegs remained planted while the fore- and midlegs swung from the dorsal to ventral sides, consistent with swinging motions expected to promote inertial reorientation. Once the forelegs contacted the ground, the specimen was close to the resting pose for this righting mode because its cranial-caudal axis was already elevated relative to the ground.

In 3.0% of trials (9:299; four 4^th^ instars on sandpaper and 1 on posterboard, three 3^rd^ instars on posterboard), the nymph rotated upright with only its legs, but not its body, contacting the ground. (Fig. 4D, Movie 1) This lifted rotating behavior required the nymph to position itself on all three of its legs on the side nearer the substrate while swinging its raised contralateral legs dorsal-ventrally so as to flip rapidly upright in a single motion.

Only one specimen in one trial succeeded in righting by pitching about a pivot point formed by its caudal end resting on the ground (0.3% = 1:299; one 3^rd^ instar on posterboard), although several other specimens unsuccessfully attempted to right using this mode. (Fig. 4E, Movie 1) During pitching, the nymph first used its hindlegs to pull on the ground so as to rotate and lift its body onto a support tripod formed by its hindlegs and caudal end. It then swung its fore- and midlegs from the dorsal to ventral side consistent with attempting to grasp the ground and to provide inertial torques for righting. Once the forelegs touched the ground, the nymph used all its legs to lever its body forward about the caudal end as a pivot point to complete righting.

Pooling data across all trials, life stages and substrates, we found no significant difference among the time to self-right, time/attempt or number of attempts for diagonal rotating and lifted rotating. (Table 2, Table S5, Fig. S2G-I) While the mean time/attempt for the single pitching trial agreed with the mean of the other two righting methods, the total time required to self-right by pitching and the number of attempts before success lay near their outer ranges.

**Table 2.** Summary statistics for different methods used to self-right by spotted lanternfly nymphs; also see Table S5 for results of Kruskal-Wallis tests.

|  | Diagonal rotating<br>(289 trials, 60<br>individuals) | Lifted rotating<br>(9 trials, 7<br>individuals) | Pitching<br>1 trial |
| --- | --- | --- | --- |
| Time to self-right (s) mean ±<br>SEM (SD) | 0.98 ± 0.10 (1.73) | 1.72 ± 1.10 (3.28) | 7.65 |
| Time/attempt (s) mean ±<br>SEM (SD) | 0.31 ± 0.01 (0.17) | 0.31 ± 0.05 (0.14) | 0.32 |
| Number of attempts mean ±<br>SEM (SD) | 3.3 ± 0.2 (3.7) | 4.0 ± 1.8 (5.4) | 24 |

Failure to self-right was associated with several factors. Some specimens became fatigued and stopped trying to right after repeated attempts, consistent with earlier reports that self-righting is a strenuous activity (Full et al. 1995). (Fig. S3, Movie 1) Slipping at the contact point with either glass or posterboard sometimes compromised the nymph’s ability to maintain a secure foothold or stationary pivot point. (Fig. S3A) In other cases, the specimen appeared unable to exert sufficient torque to flip upright; e.g., for some failed attempts on posterboard, the nymph lifted and rested its body on its caudal end, then tried to rotate upright by a combination of swinging its legs from the dorsal to ventral side and pulling on the ground while pivoting on its caudal end, but finally fell back onto its dorsum instead of righting. (Fig. S3B)

### Mechanical properties during terrestrial self-righting

We tracked body and leg landmarks for the one successful pitching trial and for two trials each for diagonal rotating and lifted rotating (5 total trials, each for a different individual). Movie S1 illustrates the kinematic uniformity among trials that used each righting strategy. Results were scaled to standard body mass and dimensions to allow direct comparisons between data for different specimens. The results of the analyses are plotted in Fig. 5 and S4. We refer to the approximate interval of time in which the dorsal-ventral axis changes orientation most rapidly as “active overturning” (see gray shaded regions in Fig. 5).

**Fig. 5.**
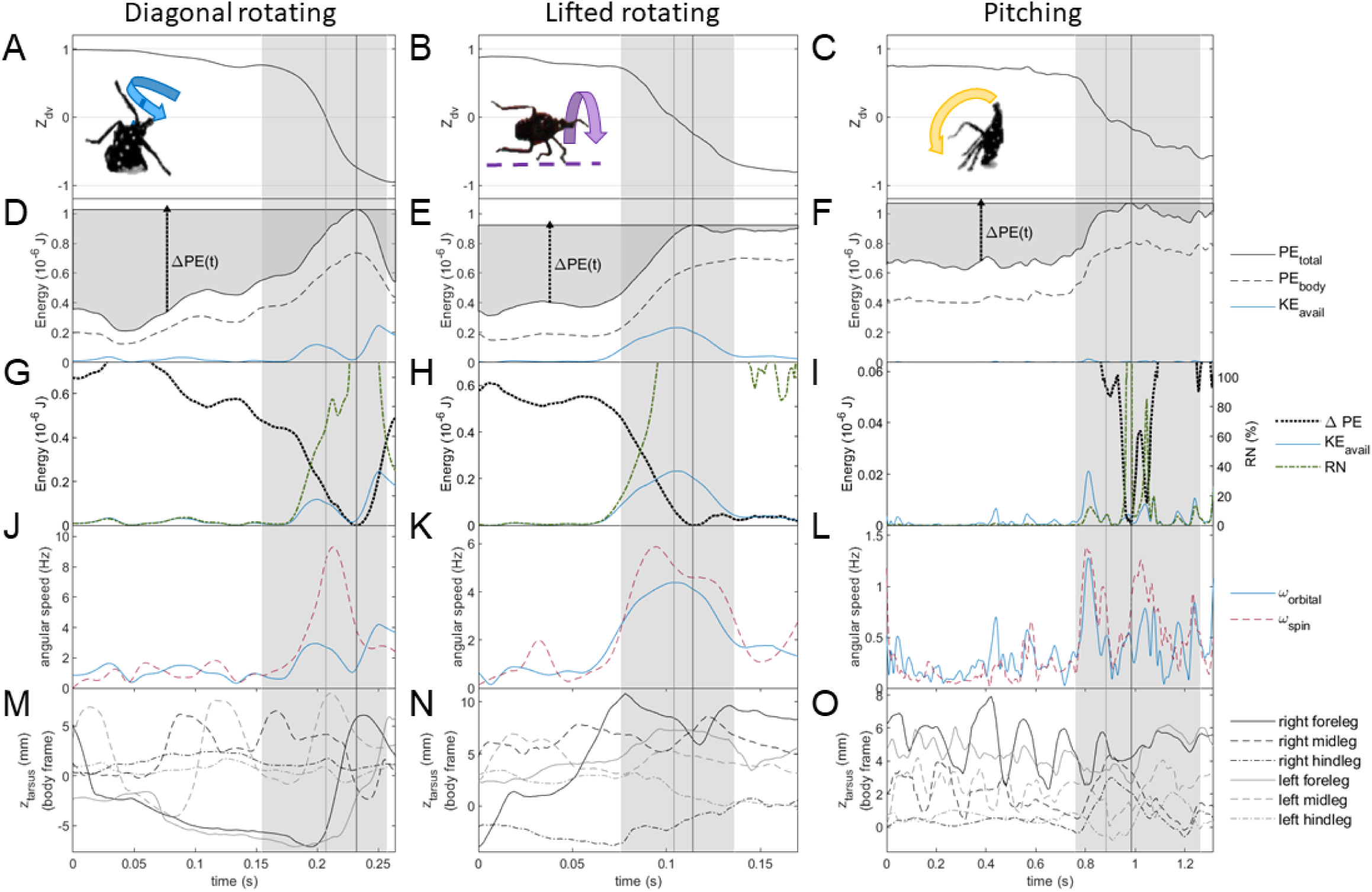
Results of analysis of the tracked coordinates and 3D model plotted for diagonal rotating, lifted rotating, and pitching. Note that different time intervals were plotted for each righting mode to display data for approximately equal relative time intervals before and during active overturning (gray shaded region) Fig. S4A,B show plots for the other diagonal rotating and lifted rotating trials, and Fig S4C shows full results for the pitching trial. (A-C) show the z-component of the dorsal-ventral axis, which transitions from ≈ +1 when overturned to 0 at the flipping point to a final value ≥ −1 with the body pitched upward in the resting pose. (D-F) show the time behavior of gravitational potential energy of the whole insect and its body, and available kinetic energy, *K_avail_*, computed from the inverted physical pendulum model (see text for details). These plots also illustrate the increase in potential energy, *ΔPE(t)*, required to right at any given time. The plots in (G-I) compare the time dependence of *ΔPE(t), K_avail_* and the righting number, *RN*, used to characterize the contribution of inertial reorientation to righting. Note especially that the energy scale for pitching in (I) is approximately an order of magnitude lower than that for the other two methods. The plots in (J-L) compare the time behavior of the angular velocity of the body’s spin motion along its pitch and roll axes with the orbital angular velocity computed from the inverted physical pendulum model; see Fig. 3F for the definitions of ω_orbital_ and ω_spin_; note that ω_spin_ only includes contributions from roll and pitch rotations. (M-O) show the motion of the tarsi (feet) along in the z-axis in the body frame, as defined in Fig. 1E. The light gray vertical line denotes the flipping point (*Z_dv_* = 0) when the specimen transitions from overturned, and the dark gray vertical line the apex of its trajectory when total PE is at its maximum. (Fig. 1I-K)

The tracking results were used to test the following predictions of the inverted physical pendulum template. First, the angular speeds for body spin, ω_spin_, and orbital motion, ω_orbital_, should be equal, so the body should flip its dorsal-ventral axis orientation and reach its maximum potential energy at the same time. (Fig. 5A-I) While ω_spin_ and ω_orbital_ were similar initially during all three methods, during active overturning the body’s angular speed was significantly greater than that for orbital motion, resulting in the body rotating to upright before the insect’s COM reached its apex. (Table 3, Fig. 5J-L) Second, the template predicts that during overturning the insect’s COM trajectory should be a circular arc curving downward and with radius comparable to the maximum height. (Fig. 3F) The measured trajectories were approximately circular during active overturning until the apex for all but one diagonal rotation trial, although only those for lifted rotation moved in an approximately vertical plane as predicted. (Fig. S5) These discrepancies indicate that the rigid object approximation used in the template did not capture fully all of the rotational dynamics during righting, highlighting the importance of measuring the dynamic leg motions in detail and using the more realistic anchor model.

**Table 3.** Summary statistics from analysis of the kinematics and dynamics during self-righting. Here, apex refers to the point at which the insect’s potential energy (*PE*) was at a maximum. Data are quoted as mean ± error bars unless otherwise stated. Error bars for *PE* were based on tracking errors and those for *KE* based on uncertainty in computing moment of inertia and angular speed. For other quantities, error bars were computed from 95% bootstrapped confidence intervals (MATLAB *bootci*, 1000 samples) added in quadrature with the variation between trials, when relevant.

|  | Diagonal rotating | Lifted rotating | Pitching |
| --- | --- | --- | --- |
| Number trials analyzed (specimens) | 2 (2) | 2 (2) | 1 (1) |
| Active overturning duration (s) | $0.078 \pm 0.006$ | $0.052 \pm 0.019$ | $0.51 \pm 0.01$ |
| $PE_{\max} - PE_i$ (J) | $7.7 \pm 1.1 \times 10^{-7}$ | $5.8 \pm 0.5 \times 10^{-7}$ | $7.0 \pm 1.0 \times 10^{-7}$ |
| Time from $Z_{dv} = 0$ to apex (ms) | $20 \pm 5$ | $8 \pm 5$ | $102 \pm 5$ |
| $\omega_{\text{spin}} : \omega_{\text{orbital}}$ | | | |
| before active overturning | $121 \pm 30\%$ | $97 \pm 14\%$ | $75 \pm 1\%$ |
| active overturning to apex | $242 \pm 31\%$ | $118 \pm 9\%$ | $131 \pm 1\%$ |
| $I_{\text{spin legs}} : I_{\text{total spin}}$ (all data) | $68 \pm 2\%$ | $72\% \pm 2\%$ | $65 \pm 1\%$ |
| $I_{\text{body}}$ ($\text{kg m}^2$) active overturning to apex | $6.0 \pm 0.2 \times 10^{-118}$ | $6.8 \pm 1.2 \times 10^{-11}$ | $9.2 \pm 0.2 \times 10^{-11}$ |
| Total torque (Nm) start to apex | $7.3 \pm 0.9 \times 10^{-7}$ | $9.1 \pm 1.0 \times 10^{-7}$ | $6.6 \pm 0.2 \times 10^{-7}$ |
| Force/leg (BW) start to apex | $0.19 \pm 0.04$ | $0.92 \pm 0.55$ | $0.33 \pm 0.02$ |
| Moment arm (mm) start to apex | $3.7 \pm 0.5$ | $2.8 \pm 0.8$ | $3.2 \pm 0.1$ |
| SM/ISM start to apex | $76 \pm 4\%$ | $-58 \pm 10\%$ | $-361 \pm 19\%$ |
| number leg ground contacts (mean ± SD)<br>start to apex | $4.8 \pm 0.1$ | $3.5 \pm 0.5$ | $3.4 \pm 0.1$ |

We now consider how energy varied throughout the righting process. (Fig. 5D-I) All specimens that successfully righted started out overturned with legs raised and ended in an upright resting pose. During planting, because the body rested on its dorsum and the legs extended laterally and dorsally to the ground, potential energy decreased to a global minimum from its initial value when overturned, *PE_i_*. The total *PE* then increased monotonically when the nymph lifted its body in preparation for attempting to right and during active overturning. The increase in total *PE* between its initial and maximum value was the same within measurement uncertainty for diagonal rotating and pitching, and somewhat lower for lifted rotating. (Table 3)

The comparison of potential and kinetic energy during righting revealed that spotted lanternfly nymphs relied primarily on quasistatic righting until shortly before flipping. The available kinetic energy estimated using the template was negligible compared to Δ*PE(t)* until the specimen began active overturning: specifically, *RN(t)* = *KE_avail_(t)*/Δ*PE(t)* < 25% until 36, 31 and 26ms before the apex for diagonal rotating, lifted rotating and pitching, respectively. During active overturning, inertial reorientation contributed substantially (*RN* > 50%) to righting within approximately 23-28 ms of the apex. (Fig. 5G-I) However, as mentioned above, the template’s estimate for *KE_avail_* did not include the body’s faster spin rotation that resulted in flipping before the apex, which indicated dynamical righting played a greater role than computed using the template.

Fig. 6 shows the measured self-righting trajectories in 2D body pitch-roll configuration space. The total *PE* computed for the legs and the body at its actual height relative to the substrate is indicated by the colored trajectory markers. The heatmap shows *PE* computed for the 3D body model, with no contributions from legs or the body’s vertical displacement from the ground. The measured trajectories and PE landscapes for diagonal rotating, lifted rotating, and pitching by spotted lanternfly nymphs were similar to those shown in (Li et al. 2019) for diagonal rotating, rolling on the ground (as opposed to lifted rotating), and pitching for a simple geometrical model of self-righting by cockroaches. For the PE landscape model used in the latter study, the predicted frequency of righting methods (rolling, diagonal rotating, and pitching, from highest to lowest frequency) based on PE barrier height agreed with the empirical data for Madagascar hissing and American cockroaches. However, the frequency of righting methods used by spotted lanternflies did not agree with this order, presumably due to the importance of their legs and body height from the ground in determining total PE, and the important contribution of their internal motions to inertial reorientation.

**Fig. 6.**
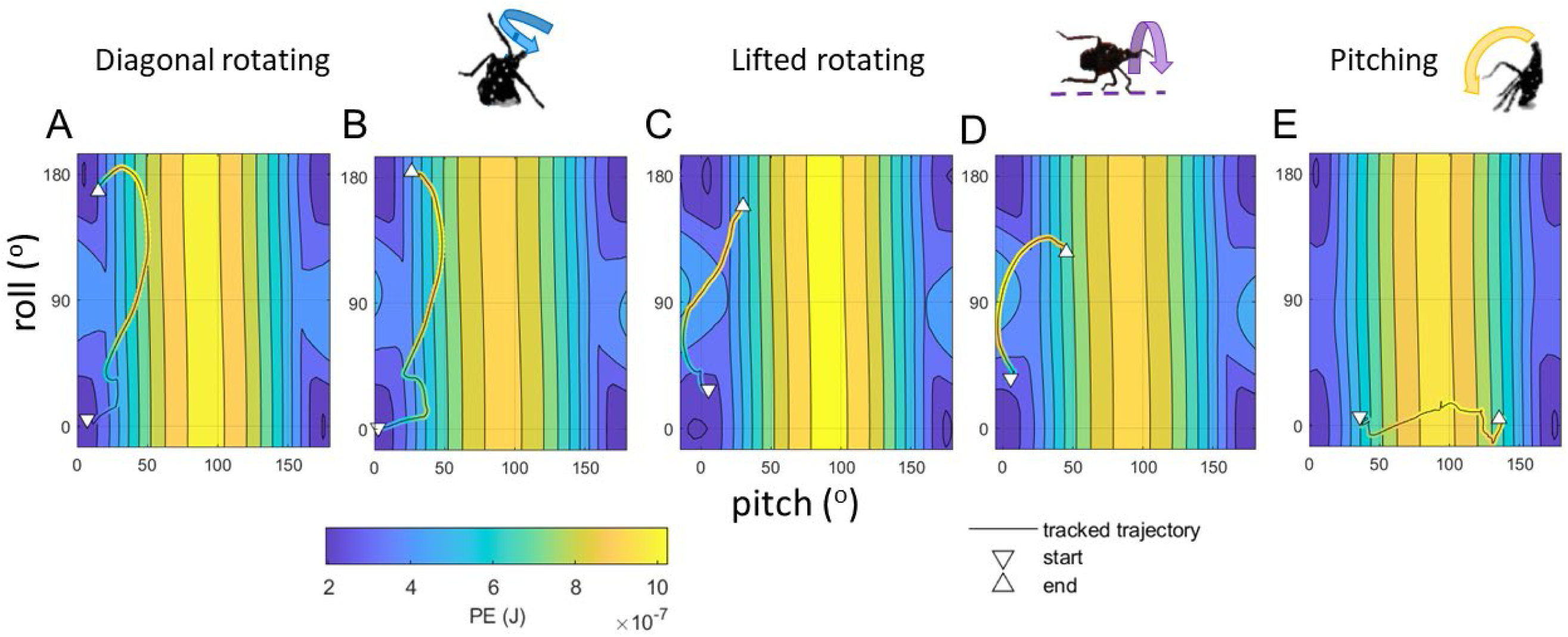
The measured pitch and roll angles vs time (black lines) for righting by A, B) diagonal rotating, C,D) lifted rotating, C) pitching shown superimposed on contour plots and heatmaps showing the gravitational potential energy (PE) landscape for the 3D rendered model of the spotted lanternfly body computed for each combination of elevation and bank angles with the lowest point on the body resting on the substrate. The actual measured values of the trajectory are also plotted using markers with colors corresponding to the computed whole insect potential energy at each time, using the colormap shown. Note that the heatmaps in A-D) were computed using a 4^th^ instar 3D model and E) for a 3^rd^ instar 3D model.

The stability margin had very different values during the three righting methods. During diagonal rotating, the nymph always maintained a positive stability margin near its ideal value by keeping one body point and 3-6 leg parts in contact with the ground. (Fig. S6D-F) By contrast, the stability margin was always negative during lifted rotating and pitching until the nymph was upright because the support polygon formed by the legs (for both methods) and caudal end (for pitching only) in contact with the ground did not enclose the COM’s projection on the ground until after flipping. (Fig. S6A-C) During pitching the nymph sometimes remained stationary when the stability margin was negative, indicating that it used its claws or arolia to adhere to the substrate.

While the mean ground reaction torques were similar within measurement uncertainty for all three righting methods, the mean and maximum force per leg, *F_leg_*, were lower during diagonal rotating than lifted rotating and pitching. This was because the most commonly-used method had more ground contacts and a somewhat greater moment arm, *r*_⟂_. (Table 3, Fig. S6) ground reaction torque and *F_leg_*, were positive until flipping, consistent with the legs providing an overturning torque during righting for all three methods. (Fig. S6G-I) During recovery, the direction of *F_leg_* varied as expected as the legs executed corrective motions required to reduce the body’s spin so that it came to rest in the righted pose.

Moment of inertia calculations were consistent with the frequencies of observed righting strategies. (See Table 3 for summary statistics, and Fig. S7 for plots of time dependence.) First, spotted lanternfly nymphs could significantly change their moment of inertia by flexing or extending their legs, which accounted for 65-72% of their total spin moment of inertia; this enhanced their ability to generate inertial torques using leg swings. Second, the body’s mean spin moment of inertia during diagonal rotating and lifted rotating, was 64% and 74%, respectively, of that for pitching. This means that either ground reaction or inertial torques performed for a given time resulted in a lower increase in ω_spin_ during pitching than the other two methods; this agreed with the measured values of ω_spin_ shown in Fig. 5J-L. Third, yaw rotations of the 3D body model had a greater moment of inertia than for roll or pitch (*I_roll_*, *I_pitch_*, *I_yaw_* = 0.46 ± 0.05, 1.25 ± 0.01, 1.50 ± 0.04 × 10^−10^ kg m^2^, respectively), which indicates that a greater torque is required to achieve the a given yaw angular acceleration than for pitch or roll. Pure yaw rotations not only do not contribute to righting when the nymph is on its back, they sometimes even lead to righting failure (Movie 1); therefore, a greater resistance to increases in yaw rate is helpful for righting.

The analysis of the tarsal motion vs time was consistent with the nymphs swinging their forelegs and midlegs to facilitate inertial reorientation. For example, in Fig. 4C and 5M, we see that during diagonal rotating, the uppermost foreleg and midleg were successively swung from the dorsal to ventral sides just before flipping. These legs also accelerated more rapidly during the dorsal-to-ventral stroke than when retracted in the reverse direction, as required for generating a reaction torque to promote righting (Frohlich 1979). The cross-correlation analysis revealed strong correlation between the forelegs and midlegs for diagonal rotating and lifted rotating (maximum similarity mean, full range 0.65 [0.46, 0.94] and 0.64 [0.34, 0.95], respectively), but lower correlation between the motions of these legs for pitching (maximum similarity 0.45 [0.37, 0.55]). During pitching, the forelegs were swung periodically, consistent with either providing inertial torque or attempting to grasp the substrate. (See Fig. S9-13 and Table S6 for full results.)

## Discussion

This study showed that determining the motions of all legs as well as the body’s orientation during righting yielded new insights into how insects perform terrestrial self-righting. Specifically, the complex, agile leg motions observed during diagonal rotating and lifted rotating by spotted lanternfly nymphs were different from those reported for other insects and proposed for legged robots (a topic we return to below). The nymphs constantly moved their legs to facilitate overturning by first raising and tilting their bodies, then pushing and pulling on the substrate (quasistatic righting), and finally swinging their legs in a coordinated fashion to generate inertial torques (dynamic self-righting). Each of the three observed righting methods involved a stereotypic sequence of motions involving all legs, with no one pair of legs playing a predominant role. These dynamic motions precluded assuming the low-dimensional potential energy landscape used effectively in earlier studies (Domokos and Várkonyi 2007; Li et al. 2019) because the insect’s COM height, and hence the potential energy, depends significantly on the leg joint angles as well as body orientation and height from the ground (Kessens et al. 2012). While a geometrical model comprising an ellipsoidal body and two legs was sufficient to describe terrestrial self-righting by discoid cockroaches (Othayoth and Li 2021), a 3D rendered body model and six articulated legs was necessary to represent the antero-posteriorly asymmetrical body shape of spotted lanternfly nymphs and the important contributions of their long, massive legs. Using this anchor model revealed that, for these insects, the legs constitute such a large fraction of the total mass that their configuration is important for understanding their dynamic mechanical properties. The relative geometry of the body and legs (i.e., body width to leg length ratio and the legs’ point of insertion relative to the body midline) also impacts an insect’s ability to self-right. For example, a wide, flattened body and relatively short legs can limit an overturned insect’s ability to grasp and push dorsally against the substrate (Faisal and Matheson 2001). Spotted lanternflies instead have legs long compared to their body width and the coxae of the fore- and midlegs are located closer to the body outer edges than the midline. Taken together, this allows them to reach their legs both laterally and dorsally, allowing them to use their legs effectively, even when overturned, as long levers to facilitate lifting and rotating.

Terrestrial self-righting methods used by spotted lanternfly nymphs had no significant dependence among success rate (92-100%), time required, number of attempts, substrate, and life stage. By contrast, the righting strategies used by beetles and stink bugs (Sasaki and Nonaka 2016; Pace and Harris 2021; Zhang et al. 2021) and the times required for righting (Zhang et al. 2021) depended significantly on substrate properties. The forces spotted lanternfly nymphs exerted on the ground by were consistent with those measured for terrestrial self-righting by discoid cockroaches (≤ 5.1 body weight/leg) (Full et al. 1995), wedging and pulling by beetles (Evans 1977; Evans and Forsythe 1984), and adhesion on glass by adult spotted lanternflies (1-2 body weight/leg) (Frantsevich et al. 2008). Spotted lanternflies are known to cling tenaciously to a wide variety of natural surfaces (Kim et al. 2011; Kane et al. 2021) using a combination of their tarsal claws and arolia (Avanesyan et al. 2019). Their use of multiple legs for grasping and adhesion likely provides them with adaptations for righting on the varied surfaces encountered in their native habitats, which range from smooth, waxy leaves to rugged ground (Sasaki and Nonaka 2016; Kane et al. 2021).

Spotted lanternfly nymphs favored self-righting by diagonal rotating, in which they used their legs to pivot about a body contact point, similar to motions used during righting by some species of beetles and cockroaches (Frantsevich 2004; Li et al. 2019). This method was found to ensure a greater stability margin and to require a lower force per leg during righting than the other two methods observed. All three methods were found to be primarily quasistatic until shortly before the flipping point, when rapid body spin and dynamic swinging leg motions became important. Spotted lanternfly nymphs also sometimes righted by using their legs to entirely lift their bodies off the substrate while overturning (lifted rotating), but rarely by pitching--a method commonly used by some species of beetles, cockroaches and stinkbugs (Frantsevich 2004; Li et al. 2019; Pace and Harris 2021). Pitching was found to correspond to a large, negative stability margin requiring stabilization by adhesion forces applied by a small number of legs until immediately before overturning; this means that even a momentary loss of contact with the ground would destabilize this method before tip-over, as observed during failed pitching attempts. (Movie 1)

It is also illuminating to compare the righting motions observed for spotted lanternfly nymphs to those proposed for righting by legged robots (Peng et al. 2017; Wang et al. 2022). The planting and lifting motions used by these insects correspond to steps required for robots to assume a starting posture suitable for using the legs to support, balance, and rotate the body. Because the nymphs cannot passively rotate upright given their flattened body geometry, they must generate the necessary overturning torque either by using pushing or pulling motions with the supporting feet or by employing dynamic swinging motions by the legs (Saranli et al. 2004). Interestingly, the motions used during diagonal rotating resemble those used in “pivoting” methods for object manipulation by robots, similar to how humans move a heavy object by lifting and balancing it on an edge (Yoshida et al. 2006). During the recovery phase, the legs supported and lowered the nymph’s body, avoiding jarring and tumbling on impact. After righting was completed, the legs were repositioned to serve as supports and to assume the typical resting pose.

We present tracking results for a small number of trials that is consistent with the number analyzed in earlier studies (Burrows et al. 2015; Othayoth and Li 2021), due to the difficulty of manually tracking dozens of points over thousands of video frames. Ongoing improvements in unsupervised tracking methods based on deep learning promise to facilitate gathering larger tracking datasets for inter- and intraspecies comparisons. This would also allow expanding the study of terrestrial self-righting to consider a wider range of insect species with diverse body and foot morphologies (e.g., tarsal claws vs adhesive pads). In future work, it would be interesting to see if these results for self-righting generalize to a wider range of substrates, including granular, compliant, and inhomogeneous materials. The analysis methods used in this study can be applied to a wide range of animals for which legs, tails and long necks are employed during righting, and therefore promise a new source of inspiration for robotic models of righting. On a practical note, this study demonstrates the feasibility of creating 3D models of external morphology using free software packages and low-cost photogrammetry for specimens too small for 3D scanning, and for studies that do not require the resolution and full 3D imaging capabilities of microCT. Given that this combination of 3D tracking and modeling has proved productive for understanding rotational dynamics during insect jumping (Li et al. 2023) and terrestrial self-righting (this study), these methods could also be extended to explore a wider range of motions (e.g., modes of locomotion) in combination with 3D prints based on realistic 3D models (Behm et al. 2018).

## Acknowledgements

The authors thank Robert Beyer and Kent Watson for instrumental design and fabrication assistance.

## Data availability

The data underlying this article are available in the article and in its online supplementary material.

