## Supplemental figures and tables for "Using pose estimation and 3D rendered models to study leg-mediated self-righting by lanternflies"

[§] Contributed equally

[*] Corresponding author

**Supplemental Dataset S1**

Dataset S1.zip: Datasets and computer code used for analysis and creating models and figures

**Movie 1.mp4**

Videos showing examples of the behaviors used by spotted lanternfly nymphs during successful and failed terrestrial self-righting attempts, and animations of the 3D models.

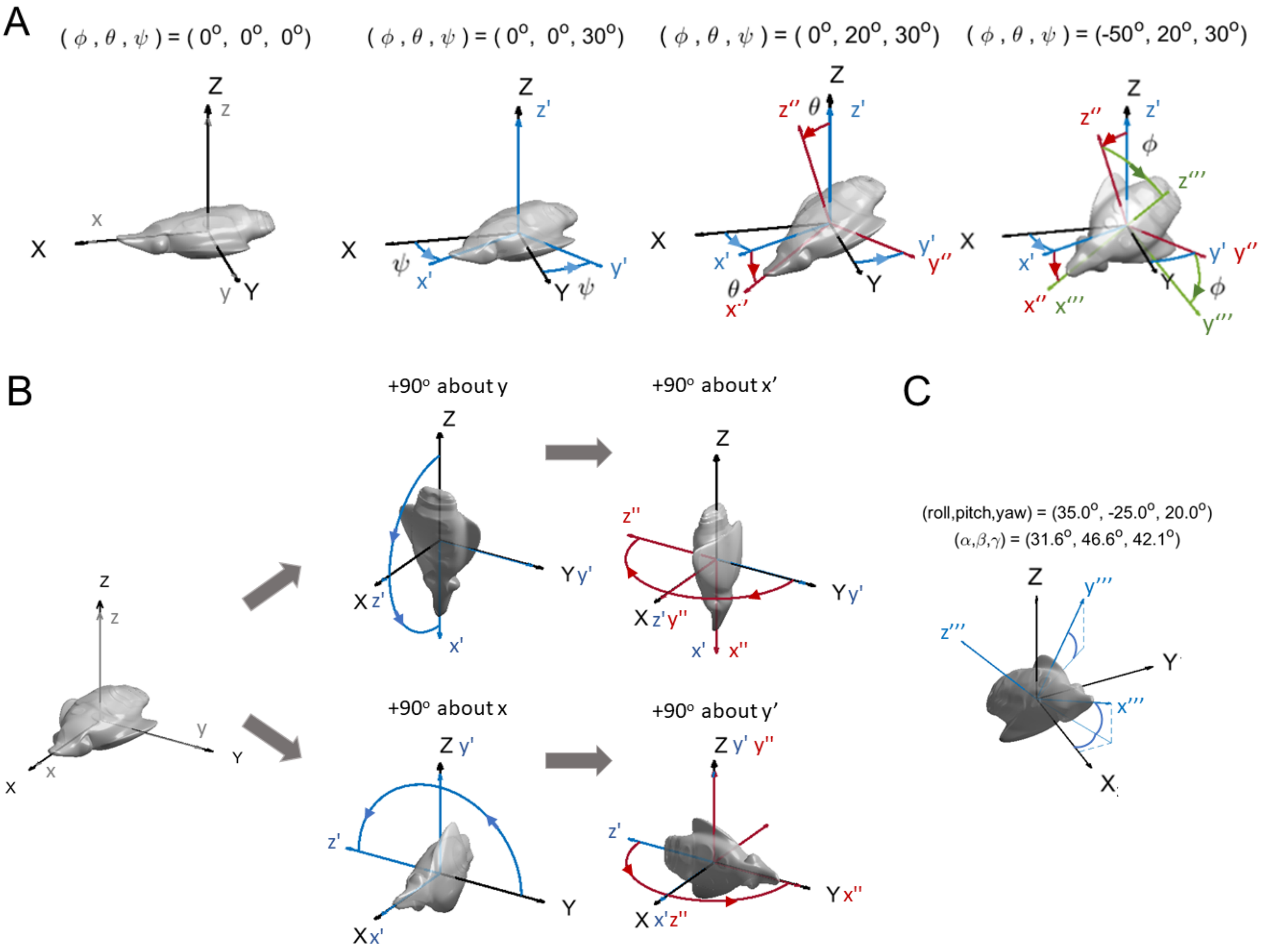
A
( φ , θ , ψ ) = ( 0°, 0°, 0°)
( φ , θ , ψ ) = ( 0°, 0°, 30°)
( φ , θ , ψ ) = ( 0°, 20°, 30°)
( φ , θ , ψ ) = (-50°, 20°, 30°)
B
+90° about y
+90° about x'
+90° about x
+90° about y'
C
(roll,pitch,yaw) = (35.0°, -25.0°, 20.0°)
(α,β,γ) = (31.6°, 46.6°, 42.1°)

**Fig. S1 (preceding page)**

A) Illustration of the three successive yaw-pitch-roll rotations, where the axes' orientations are shown as *x'y'z', x"y"z" and x"'y"'z"'* after the first, second and third rotations, respectively. B) Example image showing that performing the 3D rotations denoted by the Euler angles in different order result sin different final orientations. More generally, rotations in 3D do not commute: i.e., rotations about different axes that occur in a different order result in different final 3D orientations. C) Illustration of how the angles that describe the final orientation of the body *xyz* axes with respect to the *XYZ* axes of the spatial frame (i.e., the angle α between *x'''* and *X*, β between *y'"* and *Y*, and γ between *z*''' and *Z*) do not in general correspond to the roll, pitch and yaw Euler angles. (Figures created with code adapted from MATLAB example code (e.g. *dr.draw3DOrientation* at https://www.mathworks.com/help/nav/ug/rotations-orientation-and-quaternions.html, accessed January 16, 2024).

**Fig. S2**

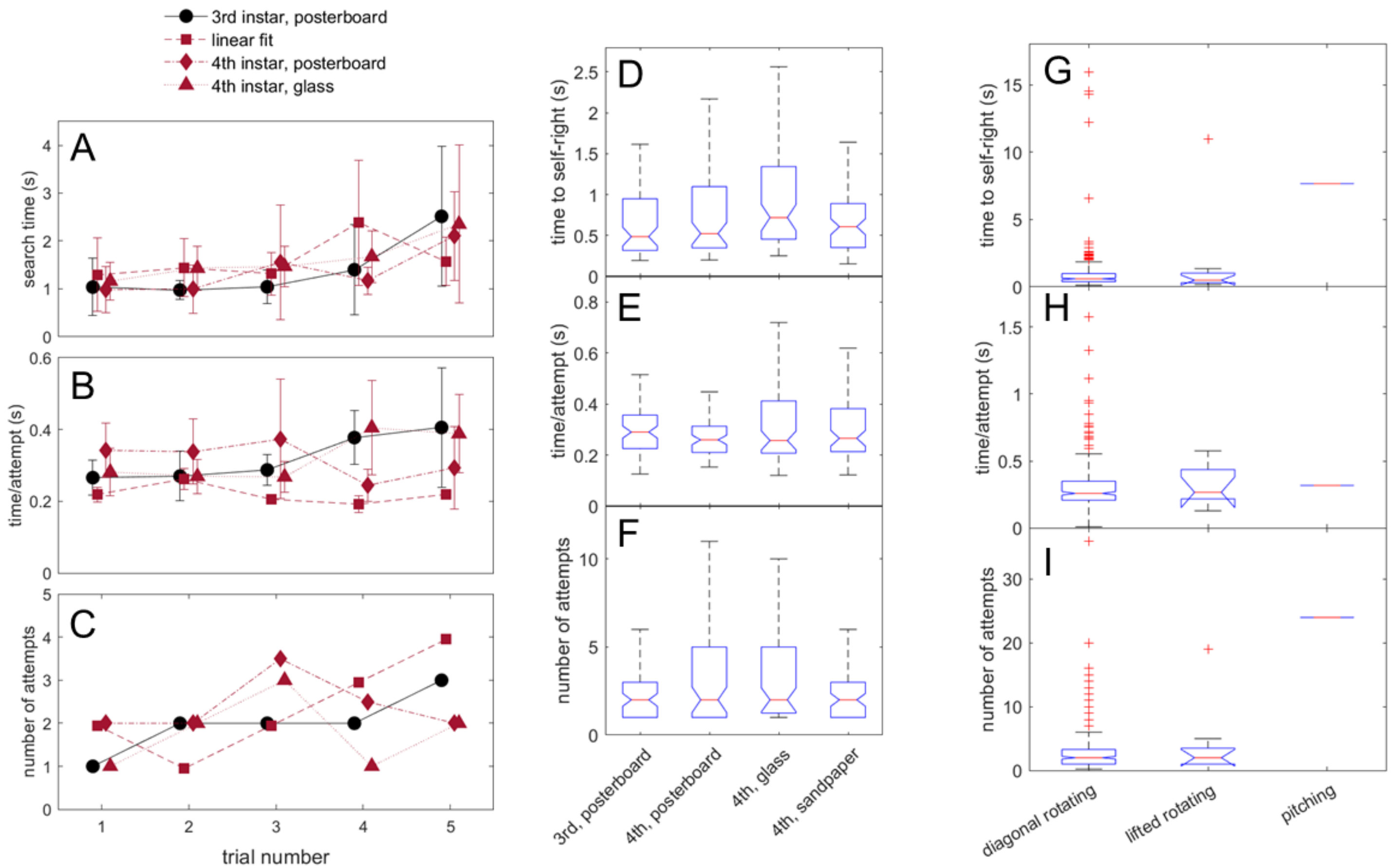

Dependence on trial number of the time spent searching before attempting righting (A), time/attempt (B), and (C) number of attempts during terrestrial self-righting, broken down by life stage and substrate. Error bars in A and B indicate 95% CI. Box-and-whisker plots of (D) time to self-right, (E) time/attempt and (F) number of attempts vs substrate and life stage combinations. (G-I) show the same quantities vs righting methods. In D-I, the red lines are medians, blue boxes the 25th to 75% percentiles, outliers are red + markers, and whiskers denote maximum and minimum values that are not outliers. If the notches around the medians of each pair of data fail to overlap, then the medians disagree at the 95% CI.

**Fig. S3**

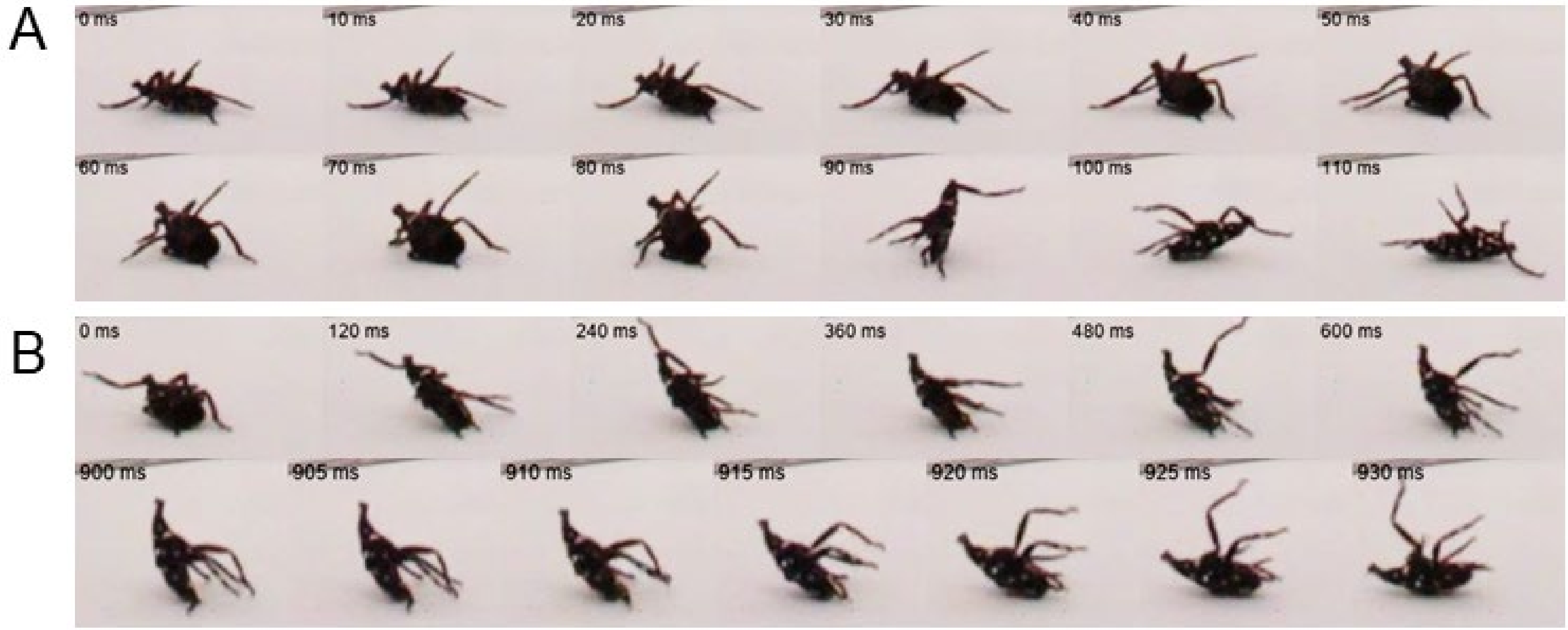

Sequences of closeup still images (times relative to the first image shown at top left; spacings between frames also indicated in caption) for attempts by 3rd instar spotted lanternfly nymphs to right on posterboard that failed because of A) slipping (every 20 ms) and B) inadequate torque (top row: every 120 ms; bottom row: every 5 ms).

**Fig. S4**

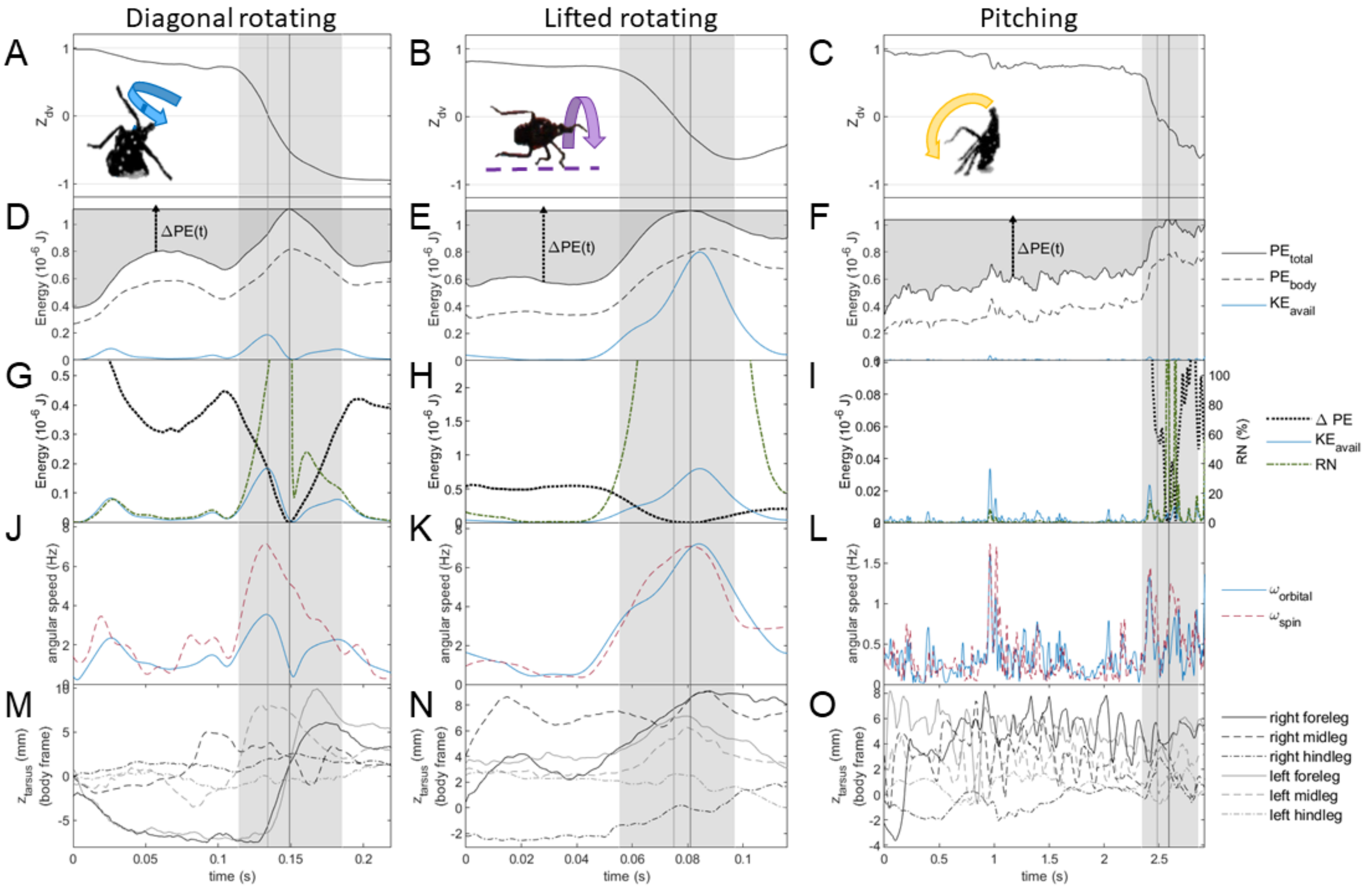

Additional results of analysis of the tracked coordinates and 3D model plotted for diagonal rotating and lifted rotating not shown in Fig. 5, and full results for pitching. Note that different time intervals were plotted for each righting mode for diagonal rotating and lifted rotating to display data for approximately equal relative time intervals before and during active overturning (gray shaded region) (A-C) show the z-component of the dorsal-ventral axis, which transitions from ≈ +1 when overturned to 0 at the flipping point to a final value ≥ -1 with the body pitched upward in the resting pose. (D-F) show the time behavior of gravitational potential energy of the whole insect and its body, and available kinetic energy, $K_{avail}$, computed from the inverted physical pendulum model (see text for details). These plots also illustrate the increase in potential energy, *ΔPE(t)*, required to right at any given time. The plots in (G-I) compare the time dependence of *ΔPE(t),* $K_{avail}$ and the righting number, *RN*, used to characterize the contribution of inertial reorientation to righting. Note especially that the energy scale for pitching in (I) is approximately an order of magnitude lower than that for the other two methods. The plots in (J-L) compare the time behavior of the angular velocity of the body's spin motion along its pitch and roll axes with the orbital angular velocity computed from the inverted physical pendulum model; see Fig. 3F for the definitions of $\omega_{orbital}$ and $\omega_{spin}$; note that $\omega_{spin}$ only includes contributions from roll and pitch rotations. (M-O) show the motion of the tarsi (feet) along in the z-axis in the body frame, as defined in Fig. 1E. The light gray vertical line denotes the flipping point ($Z_{dv}$ = 0) when the specimen transitions from overturned, and the dark gray vertical line the apex of its trajectory when total PE is at its maximum. (Fig. 1I-K)

**Fig. S5**

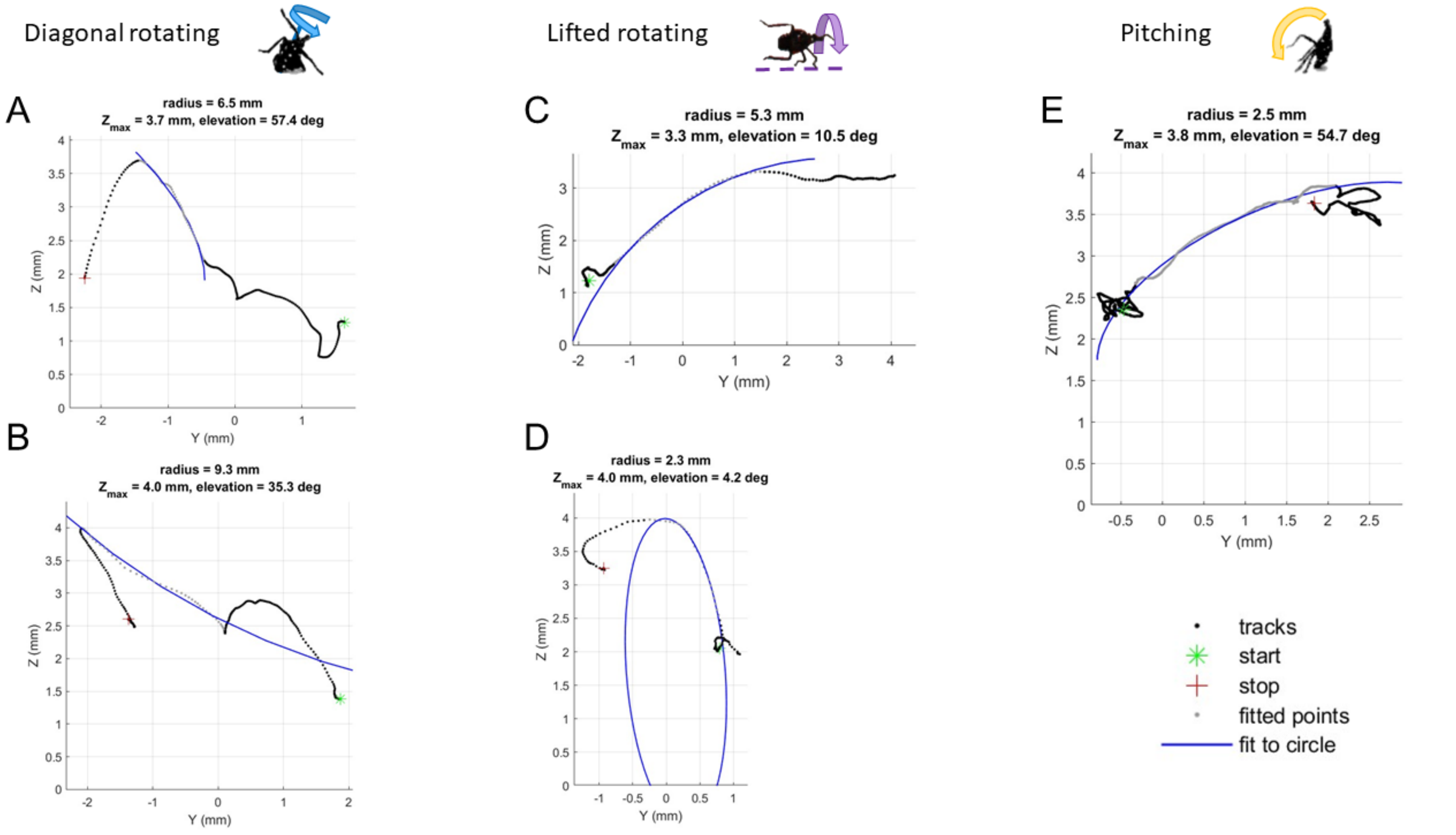

Insect center of mass (COM) trajectory during self-righting compared to a fit to a circle from the start of active overturning the apex (maximum height) of the trajectory. In each case, the maximum fit residual was less than the tracking uncertainty. Note that the fit in (B) does not correspond to the correct curvature to agree with geometry predicted by the inverted pendulum model

**Fig. S6**

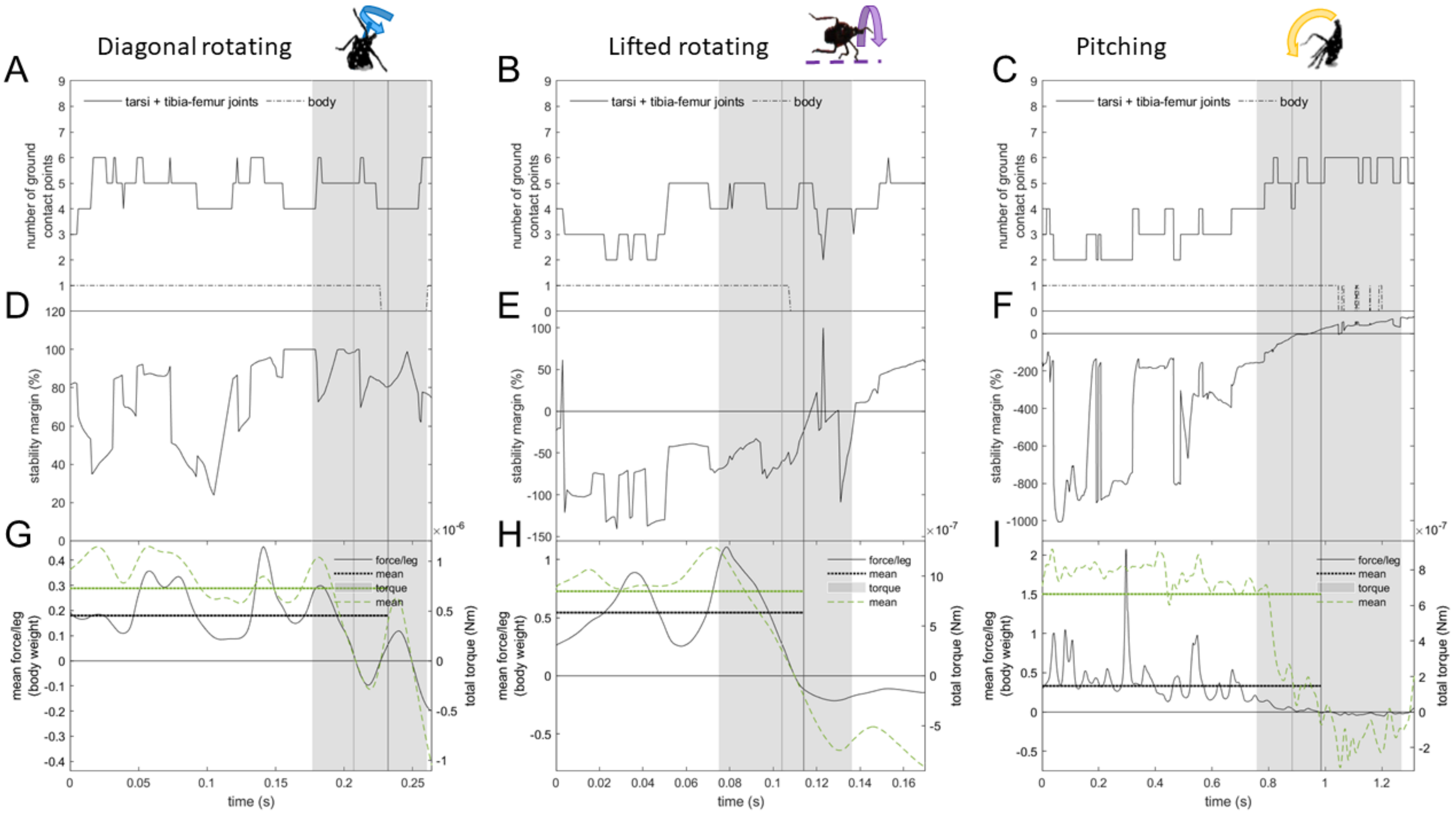

Diagonal rotating
Lifted rotating
Pitching
A
B
C
D
E
F
G
H
I
tarsi + tibia-femur joints
body
number of ground contact points
stability margin (%)
mean force/leg (body weight)
total torque (Nm)
force/leg
mean
torque
mean
time (s)

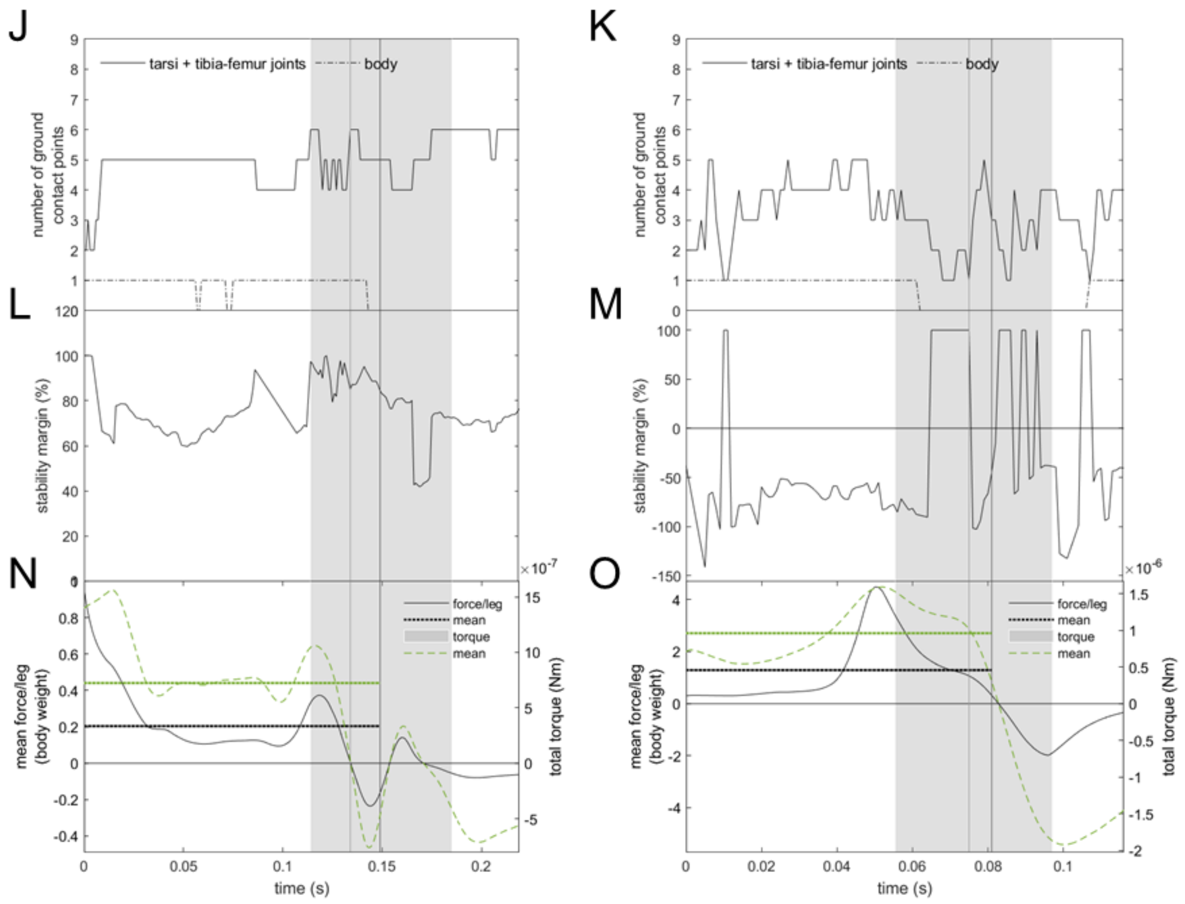

(A, B, C, J, K) number of leg and body points in contact with the ground, (D, E, F, L, M) stability margin/ideal stability margin (SM %) computed as percent ideal stability margin, (G, H, I, N, O) ground reaction force per leg and total torque vs time. Note that the abrupt changes in the stability margin and force per leg are due to changes in the number of leg parts contacting the ground. The light gray vertical line denotes the flipping point when the specimen transitions from overturned, and the dark gray vertical line the apex of its trajectory, while the shaded gray region denotes active overturning.

**Fig. S7**

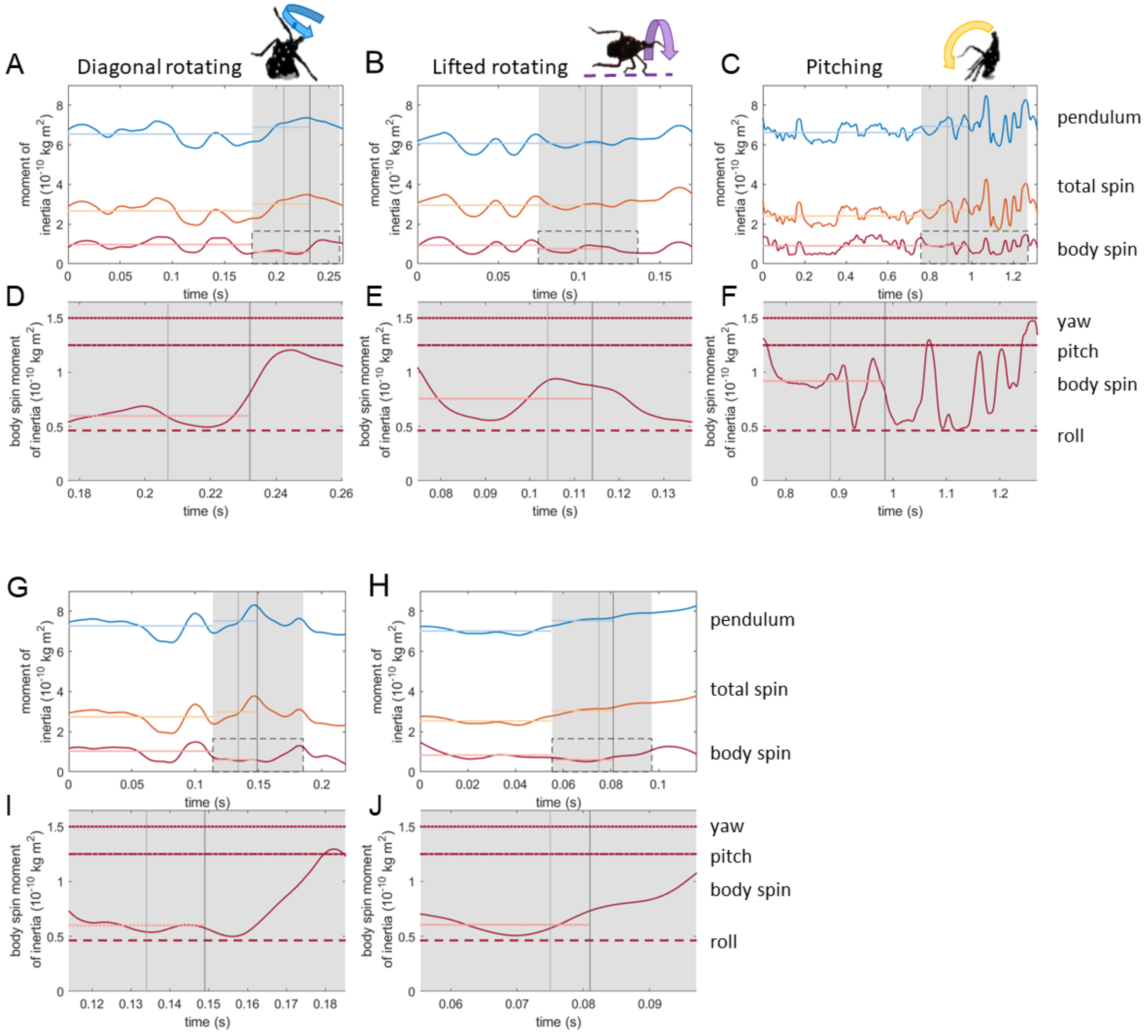

Moment of inertia values for the three rotational modes used by spotted lanternfly nymphs for self-righting. (A, B, C, G, H) Moment of inertia vs time immediately before and during righting for the physical pendulum model, $I_{pendulum}$, the whole insect spin moment of inertia including body and legs, $I_{spin,total}$, the spin moment of inertia of only the body, $I_{body}$, and the mean value of $I_{body}$. (D,E,F,I,J) Zoomed in plot of the region shown in the gray dashed boxes in A-C that shows the relationship of the body's spin moment of inertia, $I_{body}$, to its mean and to the values for pure roll, pitch and yaw rotations. These values correspond to a body spin rotation axis [roll, pitch, yaw] of [0.94, -0.17, 0.29] ± 0.02 for diagonal rotating, [-0.89,0.43,-0.13] ± 0.02 for lifted rotating, and [0.61, -0.77, 0.17] ± 0.02 for pitching, equivalent to the angle, δ, between the body's spin rotation axis & cranial-caudal axis (Fig. 3H) being equal to $18 \pm 3^{o}$, 28 ±

11 °, and 50 ± 8 °, respectively. The light gray vertical line denotes the flipping point when the specimen transitions from overturned, and the dark gray vertical line the apex of its trajectory, while the shaded gray region denotes active overturning.

**Fig. S8**

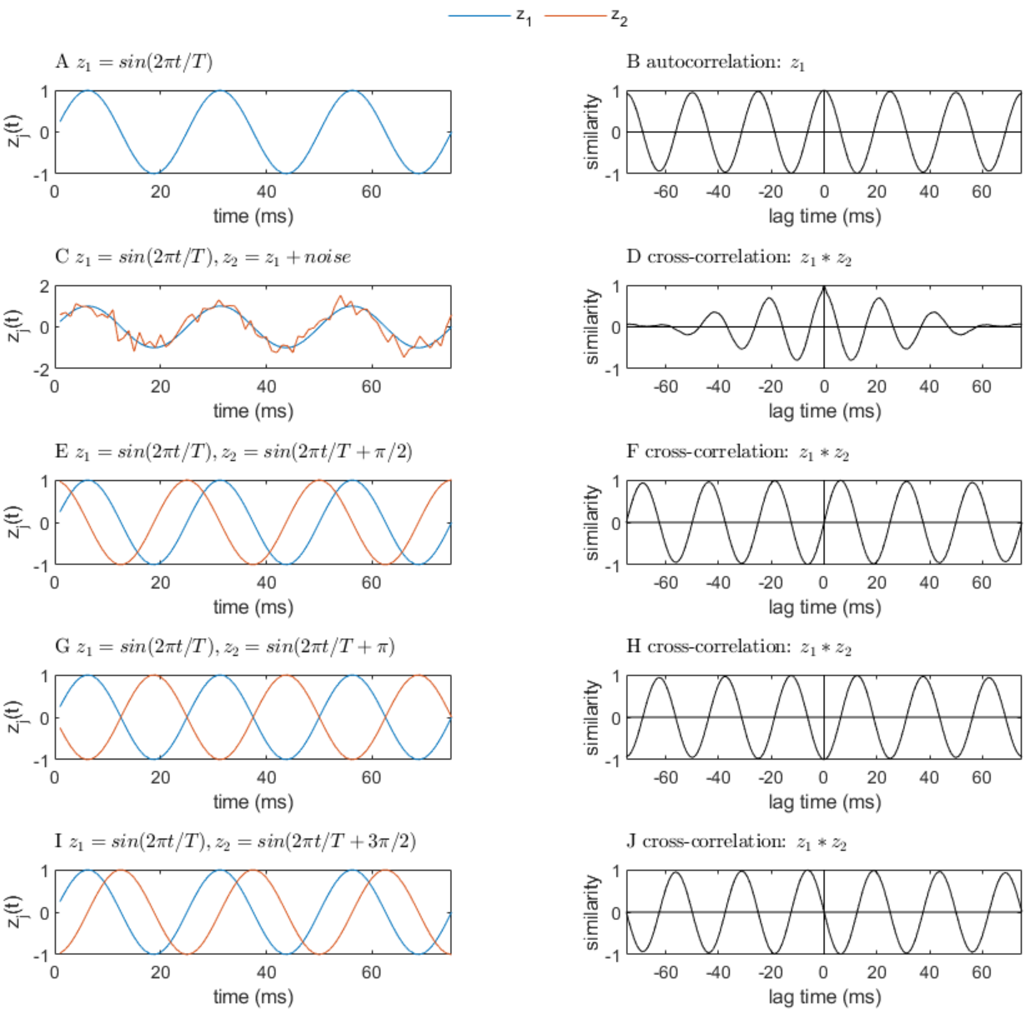

Illustrative examples of the cross-correlation vs lag time (time offset) useful in interpreting the results in Fig. S7-9. A, B) The normalized autocorrelation (cross-correlation of a function with itself) for a sine function, $z_1 = \sin(2\pi t / T)$, where the period T = 25 ms, has value 1 at zero lag time by definition, and peaks at integer multiples of the period. C, D) The cross-correlation of the sine function from A) with the sine function with noisy period and amplitude, still has periodic peaks, but their amplitude decreases to zero as absolute lag time increases. E-I) illustrate the cross-correlation of the sine function from A) with another sine with the same period, T, but a nonzero phase offset, φ: $z_2 = \sin(2\pi t / T + \varphi)$. The following plots show the cross-correlation between a sine and a sine with the same period with a phase offset of: B) $\varphi = 0$ (in phase), F) $\varphi = \pi /2$, H) $\varphi = \pi$ (out of phase), and J) $\varphi = 3\pi/2$.

**Fig. S9**

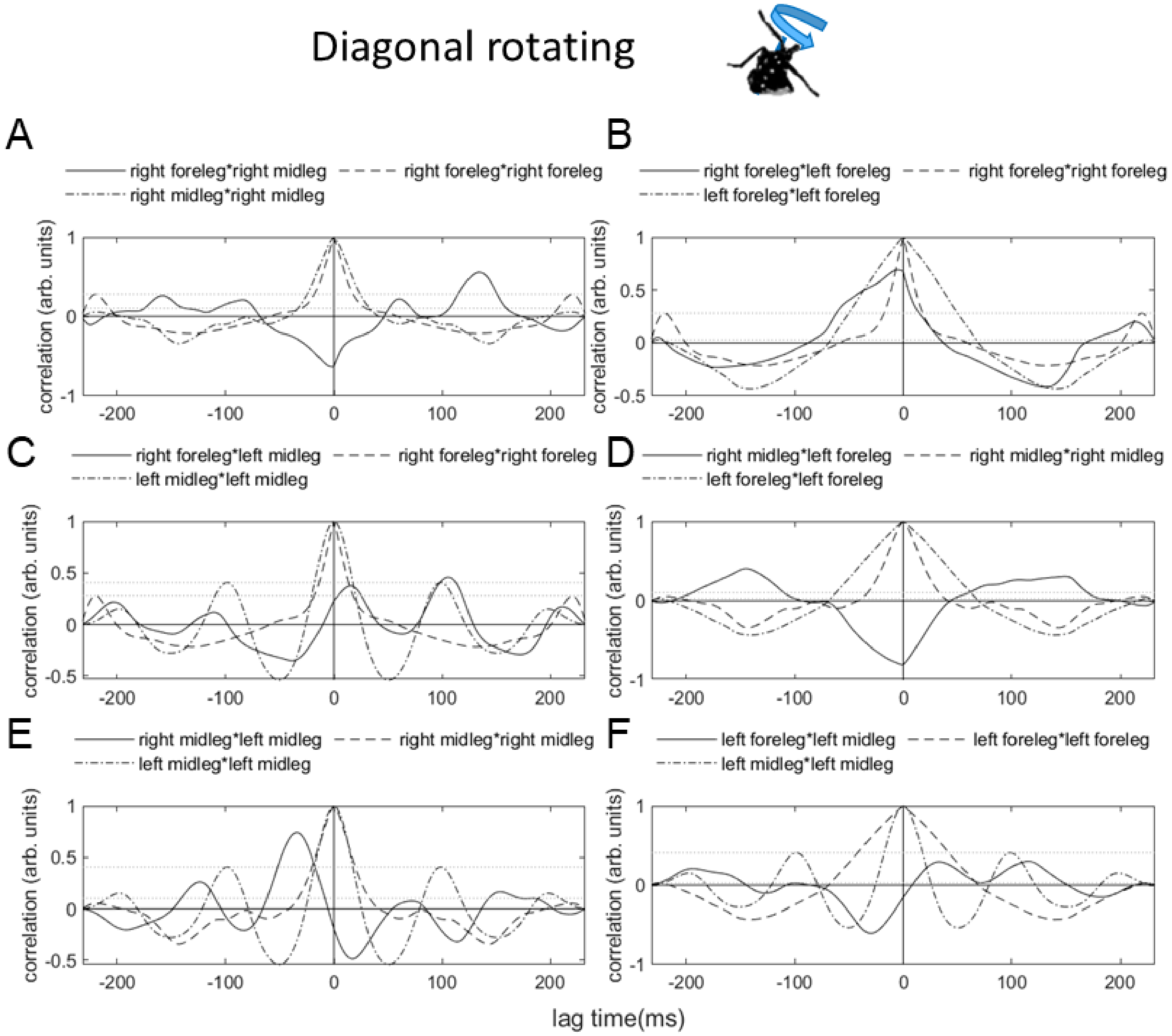

Cross-correlation of each pair of legs (solid lines) compared with the corresponding autocorrelation for each leg in the pair (dashed and dot-dashed line) during righting by diagonal rotating. Note that the right legs are uppermost during righting in this trial, while the legs on the left side primarily support the body. (Gray horizontal lines indicate the maximum values of the autocorrelation for each leg in the pair at nonzero lag time.)

**Fig. S10**

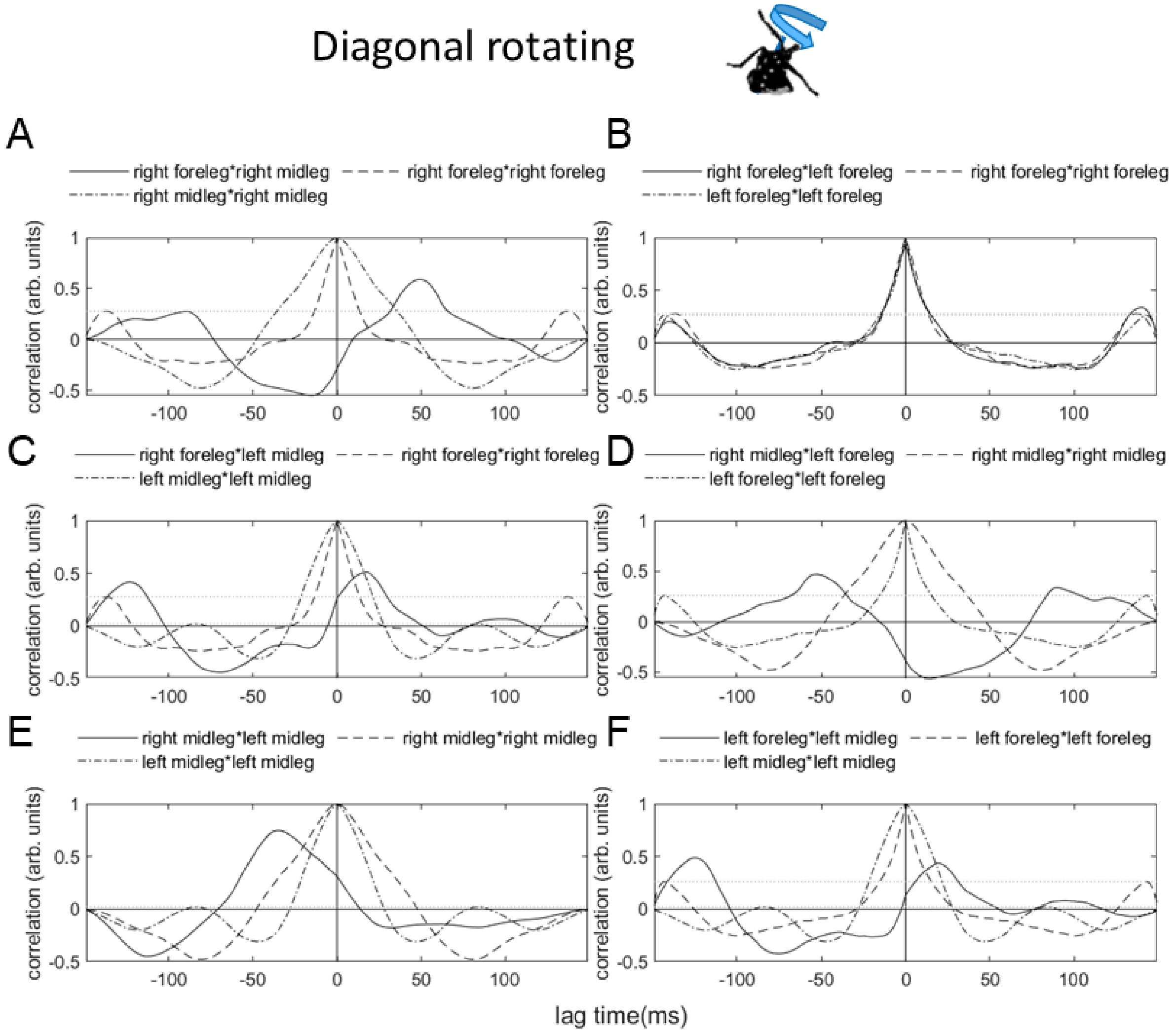

See caption for Fig. S9

**Fig. S11**

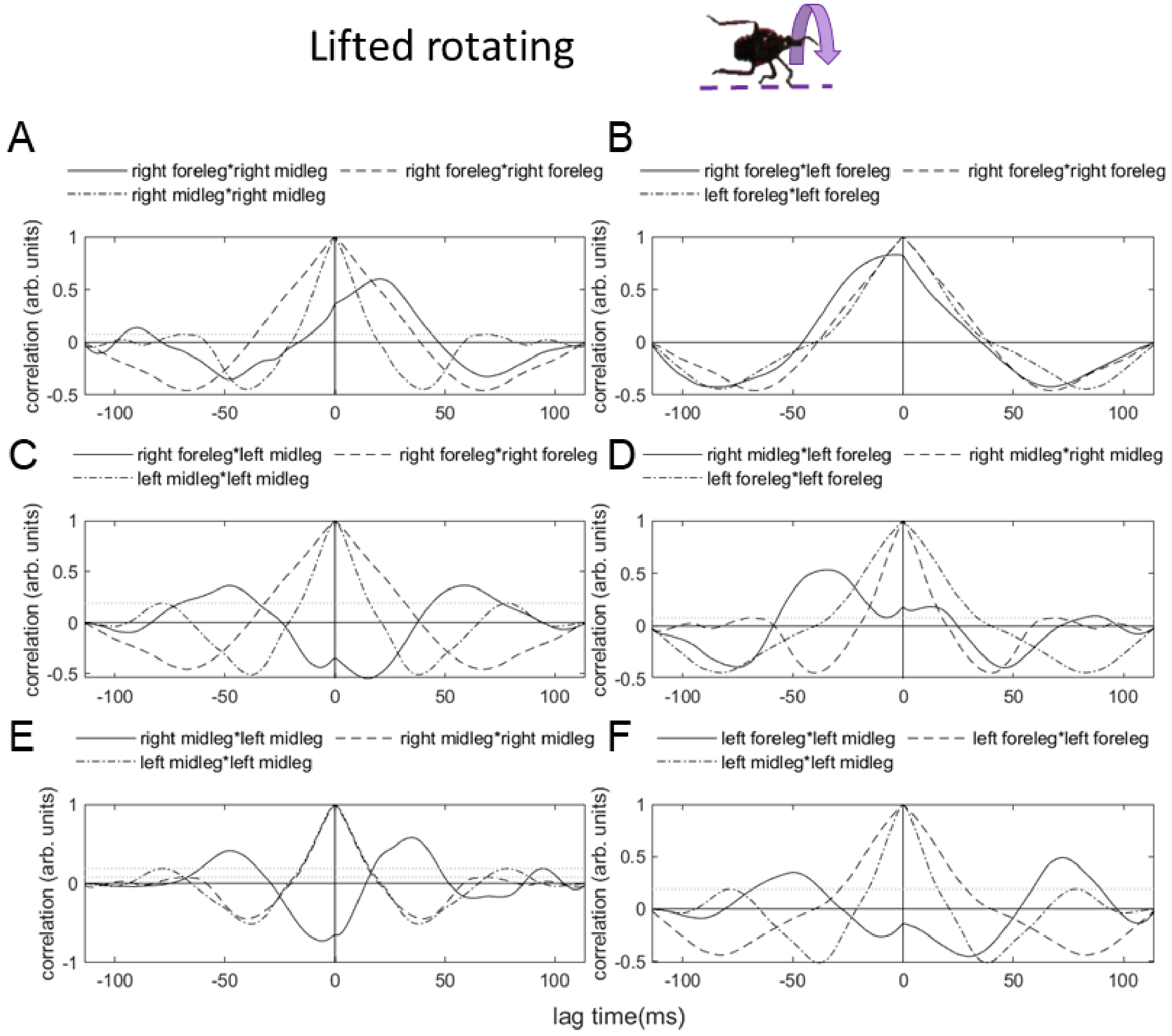

See caption for Fig. S9

**Fig. S12**

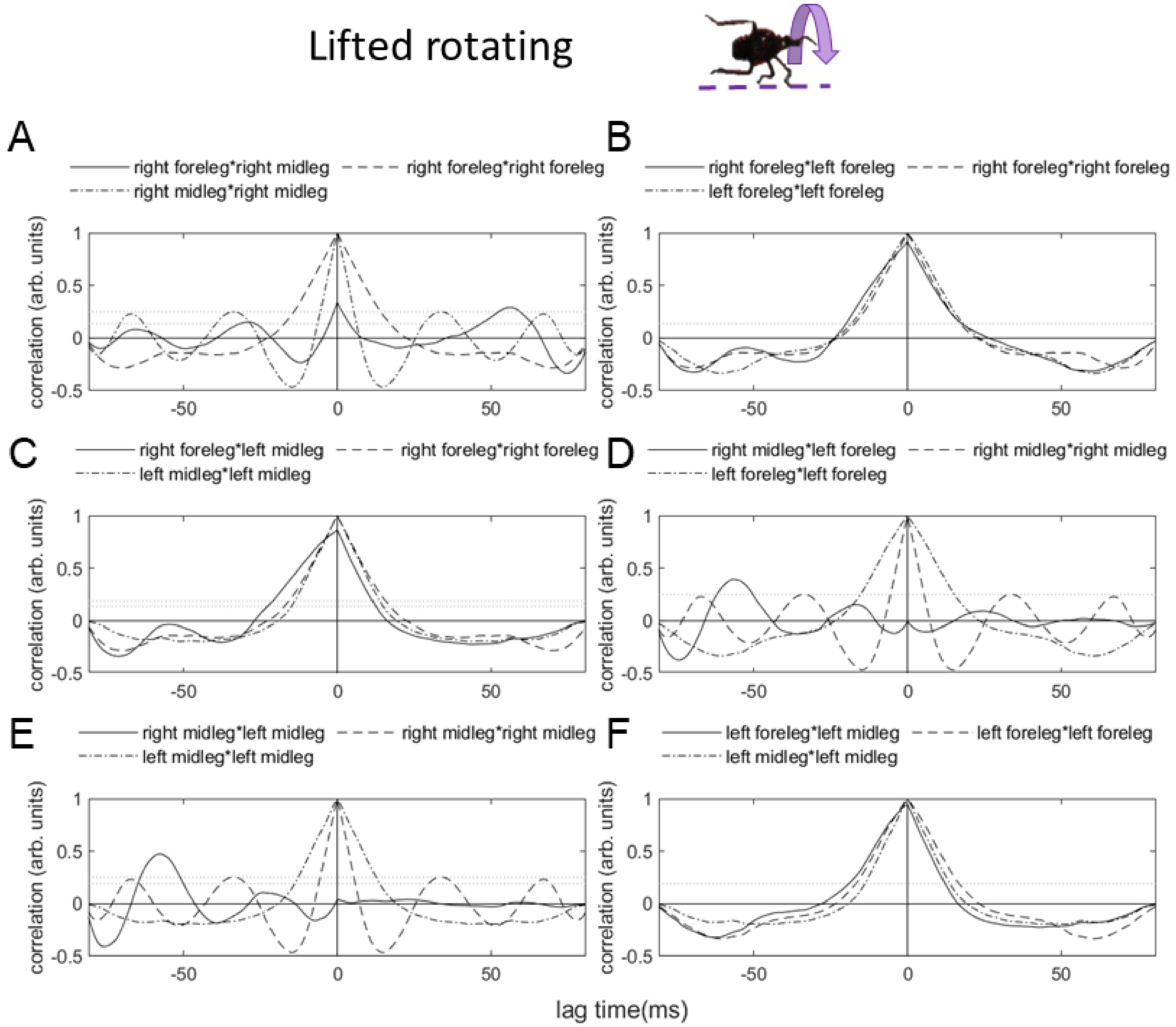

See caption for Fig. S9

**Fig. S13**

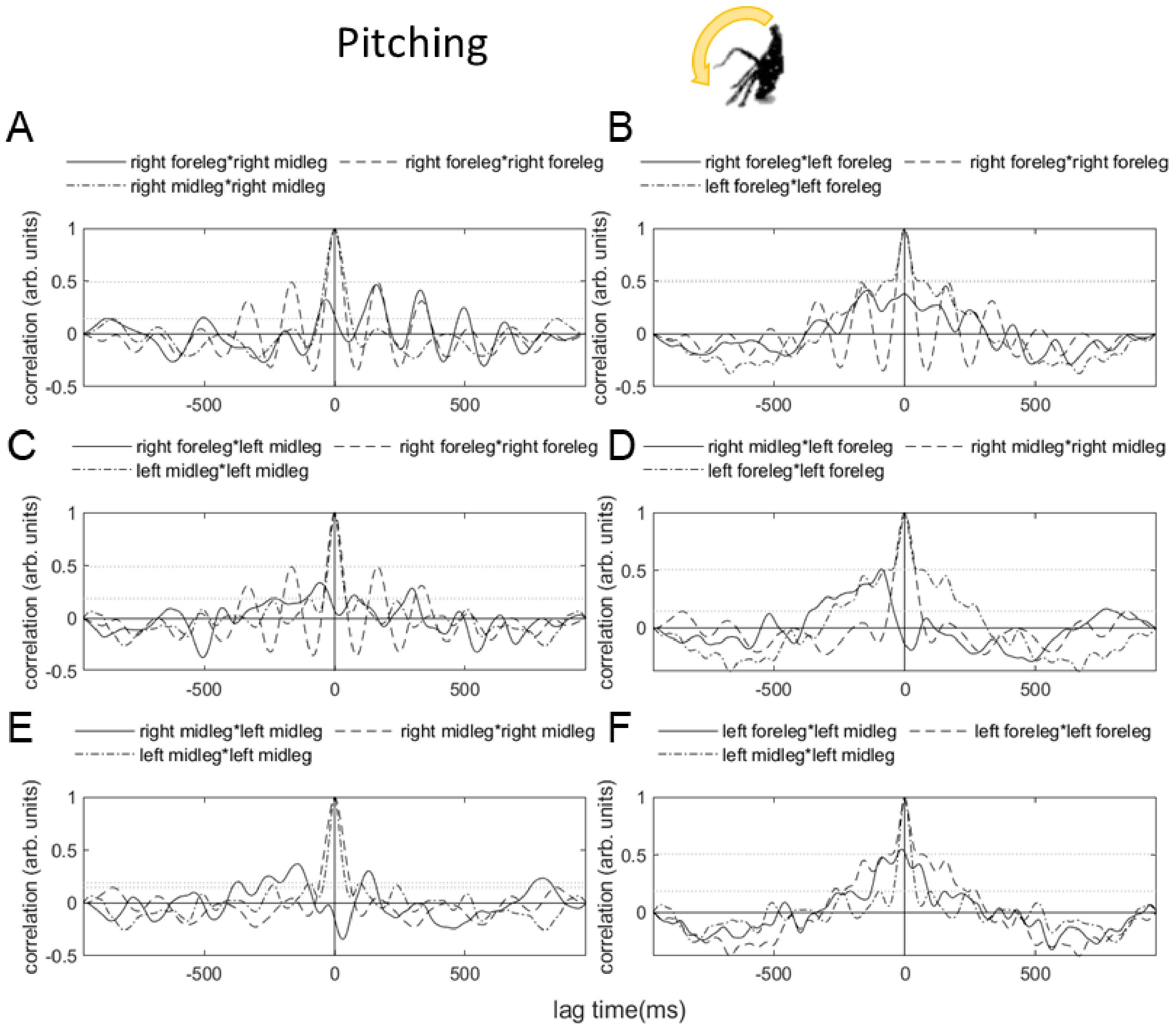

See caption for Fig. S9

Table S1. Summary statistics for terrestrial self-righting successful trials, times, attempt times and number of attempts per trial

Data for times and number of attempts are given as median ± MAD, where MAD = median absolute deviation.

| Substrate | Instar | Number of trials | Number (%) successful trials | time to self-right (ms) | attempt time (ms) | number of attempts |
|---|---|---|---|---|---|---|
| posterboard | 3rd | 75 | 75 (100%) | 486 ± 228 | 290 ± 65 | 2 ± 1 |
| posterboard | 4th | 75 | 70 (93%) | 517 ± 206 | 255 ± 45 | 2 ± 1 |
| glass | 4th | 75 | 69 (92%) | 687 ± 278 | 253 ± 69 | 2 ± 1 |
| sandpaper | 4th | 74 | 74 (100%) | 607 ± 255 | 266 ± 79 | 2 ± 1 |

**Table S2. Results of linear regression on success rate, percent diagonal somersaulting, time/attempt and attempt number vs trial number**

Sample size n = 15. Degrees of freedom = 3. * Results for the 3rd instar self-righting success rate on posterboard are not reported because they were 100% for all trials.

Self-righting success rate

| Life stage, substrate | R-squared | F-statistic | P-value |
|---|---|---|---|
| 3rd instar, posterboard * | --- | --- | --- |
| 4th instar, posterboard | 0.63 | 5.0 | .11 |
| 4th instar, glass | 0.59 | 4.4 | .13 |
| 4th instar, sandpaper | 0 | 0 | 1 |

Time/attempt

| Life stage, substrate | R-squared | F-statistic | P-value |
|---|---|---|---|
| 3rd instar, posterboard | 0.68 | 6.4 | 0.09 |
| 4th instar, posterboard | 0.30 | 1.3 | 0.34 |
| 4th instar, glass | 0.64 | 5.3 | 0.11 |
| 4th instar, sandpaper | 0.36 | 1.7 | 0.28 |

Attempt number

| Life stage, substrate | R-squared | F-statistic | P-value |
|---|---|---|---|
| 3rd instar, posterboard | 0.80 | 12 | 0.04 |
| 4th instar, posterboard | 0.69 | 6.8 | 0.08 |
| 4th instar, glass | 0.01 | 0.045 | 0.85 |
| 4th instar, sandpaper | 0.04 | 0.11 | 0.76 |

**Table S3. Results of chi-squared proportion tests on the percent of successful self-righting attempts.**

Group A, B = instar, substrate. Sample size: n = 75 for all conditions except 4$^{th}$ instars on sandpaper for which n = 74. Degrees of freedom = 1.

| Group A | Group B | p-value | chi-squared |
|---|---|---|---|
| 3$^{rd}$/posterboard | 4$^{th}$/posterboard | 0.07 | 3.3 |
| 4th, posterboard | 4th, glass | 0.73 | 0.11 |
| 4th, posterboard | 4$^{th,}$ sandpaper | 0.07 | 3.3 |
| 4th, glass | 4$^{th,}$ sandpaper | 0.07 | 3.3 |

**Table S4. Statistics from Kruskal-Wallis tests for self-righting times and attempt numbers for different life stages and substrates.**

Group A, B = instar, substrate. Sample size: n = 75 for all conditions except 4$^{th}$ instars on sandpaper for which n = 74. Limits are used for testing but are not simultaneous confidence intervals.

Time to self-right

| Group A | Group B | Lower Limit | A-B | Upper Limit | P-value |
|---|---|---|---|---|---|
| 3rd, posterboard | 4th, posterboard | -43.2 | -6.9 | 29. 4 | 0.96 |
| 4th, posterboard | 4th, glass | -65.3 | -29.0 | 7.3 | 0.17 |
| 4th, posterboard | 4th, sandpaper | -30.4 | 6.0 | 42.4 | 0.97 |
| 4th, glass | 4th, sandpaper | -1.4 | 35.0 | 71.4 | 0.065 |

Time/attempt

| Group A | Group B | Lower Limit | A-B | Upper Limit | P-value |
|---|---|---|---|---|---|
| 3rd, posterboard | 4th, posterboard | -20.3 | 16.0 | 52.2 | 0.67 |
| 4th, posterboard | 4th, glass | -48.7 | -12.5 | 23.8 | 0.81 |
| 4th, posterboard | 4th, sandpaper | -48.9 | -12.5 | 23.9 | 0.82 |
| 4th, glass | 4th, sandpaper | -36.4 | 0.023 | 36.4 | 1 |

Number of attempts

| Group A | Group B | Lower Limit | A-B | Upper Limit | P-value |
|---|---|---|---|---|---|
| 3rd, posterboard | 4th, posterboard | -54.4 | -19.1 | 16.1 | 0.50 |
| 4th, posterboard | 4th, glass | -42.3 | -7.0 | 28.2 | 0.96 |
| 4th, posterboard | 4th, sandpaper | -16.1 | 19.2 | 54.6 | 0.50 |
| 4th, glass | 4th, sandpaper | -9.1 | 26.3 | 61.6 | 0.22 |

**Table S5. Statistics from Kruskal-Wallis tests for self-righting times and attempt numbers for different righting methods.**

Group A, B = two different righting method. Sample size: n = 289 for diagonal rotating, 9 for lifted rotating and 1 for pitching. Limits are used for testing but are not simultaneous confidence intervals.

Time to self-right

| Group A | Group B | Lower Limit | A-B | Upper Limit | P-value |
|---|---|---|---|---|---|
| Diagonal righting | Lifted righting | -55.583 | 11.426 | 78.435 | 0.91574 |

Time/attempt

| Group A | Group B | Lower Limit | A-B | Upper Limit | P-value |
|---|---|---|---|---|---|
| Diagonal righting | Lifted righting | -74.376 | -7.3664 | 59.643 | 0.96407 |

Number of attempts

| Group A | Group B | Lower Limit | A-B | Upper Limit | P-value |
|---|---|---|---|---|---|
| Diagonal righting | Lifted righting | -62.248 | 2.7553 | 67.759 | 0.99457 |

**Table S6. Similarity values from cross-correlation analysis of tarsal z motion in the body frame**

Here these values correspond to the maximum similarity for each pair of legs in Fig. S7-9; note that 1 = perfect correlation, 0 = no correlation.

| **Righting mode** | **Diagonal rotating trial 1, trial 2** | **Lifted rotating trial 1, trial 2** | **Pitching** |
|---|---|---|---|
| **Leg pair** | **Maximum similarity** | | |
| right foreleg*right midleg (ipsilateral pair) | 0.62, 0.59 | 0.60, 0.34 | 0.47 |
| left foreleg*left midleg (ipsilateral pair) | 0.61, 0.94 | 0.49, 0.92 | 0.55 |
| right foreleg*left foreleg (contralateral pair) | 0.69, 0.51 | 0.83, 0.87 | 0.42 |
| right midleg*left midleg (contralateral pair) | 0.74, 0.56 | 0.73, 0.40 | 0.37 |
| right foreleg*left midleg | 0.46, 0.75 | 0.55, 0.47 | 0.37 |
| right midleg*left foreleg | 0.81, 0.49 | 0.53, 0.95 | 0.51 |
