## Supplementary material for "Using pose estimation and 3D rendered models to study leg-mediated self-righting by lanternflies": Dataset S1: analyze_leg_motion.m

```
%% analyze coordination among legs using tarsal motion along body z-axis

close all; clear;

%% load the .mat file with analysis data
[file, pathstr] = uigetfile('*.mat','Select .mat file with analysis data');
matdatafilename = [pathstr file];
load(matdatafilename);

%% initialization

% same linestyle pair of contralateral legs, same color each side
tarsuscolors = [0 0 0;0 0 0;0 0 0;darkgray;darkgray;darkgray];
tarsuslinestyles = {'-' '--' '-.' '-' '--' '-.'};

% the analyzed tarsi coordinates are in this order:
% j =        1            2            3           4           5           6
% leg = 'rightfore'	'rightmid'	'righthind'	'lefthind'	'leftmid'	'leftfore'

% 'right foreleg','right midleg','right hindleg' = 1 2 3
% 'left hindleg','left midleg''left foreleg' = 4 5 6

% here we will analyze and plot in this order:  1 2 3 6 5 4

newlegorder = [1 2 3 6 5 4];

leglabels = {'right foreleg' 'right midleg'	'right hindleg'...
    'left foreleg' 'left midleg' 'left hindleg'};

foremidlegs = [1 2 4 5];                % only consider their correlations
hindlegs = [3 6];

% get tarsi z in body frame
% tarsus_zbody = [];
% for j = 1:6
%     tarsus_zbody(:,j) = dot((tarsuscoords(:,(1:3)+(j-1)*3)-COM_body),Zbody,2);
% end
% tarsus_zbody(idxmax+1:end,:) =  [];
tarsus_zbody = tarsus_zbody(firstframe:idxmax,newlegorder);
nsignal = size(tarsus_zbody,1);         % number of samples analyzed

% put time in ms and start at firstframe analyzed
time = 1e3*time(firstframe:idxmax); time = time';

% correct other times so first frame is t = 0
idxdvz0 = idxdvz0 - firstframe + 1;
idxmax = idxmax  - firstframe + 1;
time_active = 1e3*time_active - firstframe + 1;

% compute frequency for Fourier analysis
fs = 1/dt;                              % sampling frequency (Hz)

% only keep positive frequencies (real signals)
freq = (fs/nsignal)*(0:round(nsignal/2));        % Fs/L*(0:L-1)

freq_err = 0.5/(1e-3*time(end));        % 1/2 smallest frequency increment
                                        % = 1/2 (1/duration of signal)

phase_diff = [];

% initialize data arrays
pklocs = nan(6,1+round(nframe/25));
pkprom = nan(6,1+round(nframe/25));
pkht = nan(6,1+round(nframe/25));

%% compute z coordinates with slow time variation substracted off
% and zero mean

tarsi_z = [];

for j = 1:6
    tarsi_z(:,j) = tarsus_zbody(:,j);
    % detrend if desired to remove slow time variation
    % tarsi_z(:,j) = detrend(tarsus_zbody(:,j),2);
    % subtract off mean for correlation analysis
    tarsi_z(:,j) = tarsi_z(:,j) - mean(tarsi_z(:,j));
end

%%-------------------------------------------------------------------------
% plot the legs' zero mean tarsus z body frame positions vs time
figure('Color','white','Position',[20 60 400 700]);
t1 = tiledlayout(6,1,"TileSpacing","tight",'Padding','tight');

for j = 1:6
    nexttile;
    plot(time,tarsi_z(:,j),'k-'); hold on;
    plot(time,zeros(size(time)),'k-')
    title(leglabels{j});
    if j < 6; set(gca,'XTick',[]); end
    % plot vertical gray line when dv axis becomes horizontal
    Yminmax = ylim;
    plot([time(idxdvz0),time(idxdvz0)],[Yminmax(1),Yminmax(2)],'-','Color',htmlgray);

    % plot vertical dark gray line when PE is at its maximum
    plot([time(idxmax),time(idxmax)],Yminmax,'-','Color',0.5*htmlgray,'LineWidth',0.5);

    % shade righting region in gray
    xregion(time_active(1),time_active(2),'FaceColor',htmlgray,'EdgeColor','white');
    xlim([0 time(end)]);
end

ylabel(t1,'z_{tarsus} (mm) (zero mean)');

xlabel(t1,'time (ms)');

 
%% use xcorr of the tarsi z-body coords to find the mean phase difference
%  between pairs of legs 

figure('Color','white','Position',[850 60 800 600]);

t2 = tiledlayout(3,2,"TileSpacing","tight",'Padding','compact');
alllegpairs = nchoosek(foremidlegs,2);        % pairs of mid & forelegs

phasediffarray = cell(size(alllegpairs,1),5);

for k = 1:size(alllegpairs,1)
    nexttile;

    k1 = alllegpairs(k,1); k2 = alllegpairs(k,2);
    legpair_label{k} = [leglabels{k1} '*' leglabels{k2}];

    % compute cross-correlation for each pair
    [ysim,lags] = xcorr(tarsi_z(:,k1),tarsi_z(:,k2),'normalized');
    lag0 = find(lags == 0);                 % index of zero lag
    ysimmax(k) = max(abs(ysim));                 % get maximum similarity

    [~,idxlagpklocs] = findpeaks(ysim);         % find peaks
    [~,idxlagtroughlocs] = findpeaks(-ysim);   % find troughs
    % use the peak or trough closest to zero lag time to estimate phase
    % offset
    % find first peak and first trough closest to zero lag time
    [~,idxpk] = min(abs(idxlagpklocs - lag0));
    [~,idxtrough] = min(abs(idxlagtroughlocs - lag0));
    % estimate periodicity in cross-correlation
    crosscorrperiod = median(diff(idxlagpklocs));
    % convert to fraction of a cycle
    if idxpk > idxtrough    % trough is closer to 0 lag
        phase_diff(k) = lags(idxlagtroughlocs(idxtrough))/crosscorrperiod;
    else                    % peak is closer to 0 lag
        phase_diff(k) = lags(idxlagpklocs(idxpk))/crosscorrperiod;
    end

    % plot cross-correlation 

    plot(lags,ysim,'k-'); hold on;

    % plot autocorrelation of each leg
    [ysimk1,lags] = xcorr(tarsi_z(:,k1),tarsi_z(:,k1),'normalized');
    plot(lags,ysimk1,'k--'); 
    % find largest nonzero peak that's not at zero lag time
    [pksk1,pklocs] = findpeaks(ysimk1(lag0:end));

    [ysimk2,lags] = xcorr(tarsi_z(:,k2),tarsi_z(:,k2),'normalized');
    plot(lags,ysimk2,'k-.');     
    % find largest nonzero peak that's not at zero lag time
    [pksk2,pklocs] = findpeaks(ysimk2(lag0:end));
    
    Ylim = ylim; xlim([min(lags) max(lags)]);

    if j < 6; set(gca,'XTick',[]); end

    % plot axes centered at the origini
    plot(lags,zeros(numel(lags)),'k-'); 
    plot([0,0],[Ylim(1),Ylim(2)],'k-');

    % plot maximum values of each leg's autocorrelation function at the
    % largest nonzero peak
    if numel(pksk1) > 0
        plot(lags,max(abs(pksk1))*ones(size(lags)),':','Color',darkgray,...
            'LineWidth',0.5);
    else
        pksk1 = NaN;
    end
    if numel(pksk2) > 0
        plot(lags,max(abs(pksk2))*ones(size(lags)),':','Color',darkgray,...
            'LineWidth',0.5);
    else
        pksk1 = NaN;
    end

    ylabel('correlation (arb. units)');
    legend({legpair_label{k},[leglabels{k1} '*' leglabels{k1}],...
        [leglabels{k2} '*' leglabels{k2}]},...
        'Location','northoutside','Orientation','horizontal',...
        'NumColumns',2,'FontSize',9);
    legend boxoff;

    fprintf('leg pair %s max similarity = %2.2f\n',legpair_label{k},ysimmax(k));

    phasediffarray(k,:) = {legpair_label{k} phase_diff(k) ysimmax(k) ...
        max(abs(pksk1)) max(abs(pksk2))};
end

xlabel(t2,'lag time(ms)'); 

legend boxoff;

% save in spreadsheet
% add header
phasediffarray = [{'leg' 'phase offset (cycle)' 'max cross-corr sim' 'max auto 1' 'max auto 2'};...
    phasediffarray];

spreadsheetfilename = fullfile(pathstr,[filename(1:23),'_legcoord.xlsx']);
xlswrite(spreadsheetfilename,phasediffarray,1);

xcorrfilename = fullfile(pathstr,[filename(1:23),'_xcorr.png']);
saveas(gcf,xcorrfilename)

%% done

%%%%%%%%%%%%%%%%%%%%%%%%%%%%%%%%%%%%%%%%%%%%%%%%%%%%%%%%%%%%%%%%%%%%%%%%%%%
% functions
%%%%%%%%%%%%%%%%%%%%%%%%%%%%%%%%%%%%%%%%%%%%%%%%%%%%%%%%%%%%%%%%%%%%%%%%%%%
```
