## Supplementary material for "Using pose estimation and 3D rendered models to study leg-mediated self-righting by lanternflies": Dataset S1: Dataset S1 Metadata.docx

Metadata for Supplemental Dataset S1

### Data and code used in analysis

Description: Spreadsheet with summary statistics about success/failure rates, strategies used, etc.

- SLF self-righting video summary statistics.xlsx

Description: MATLAB code used to analyze the 3D tracked xyz body keypoints and use to create Fig. 5, S4, S6, S7 plots and Table 2. To run, accept all default settings in the first window and follow instructions to select the points at the start and end of overturning, as shown in Fig. 6’s plots of the dorsal-ventral axis z-component vs time.

- Code in folder: SLFrightingposeanalyzer
- MATLAB version vR2023b; toolboxes: Signal Processing, Mapping, Image Processing, Statistics and Machine Learning, Curve Fitting, Phased Array System, Automated Driving, Lidar, Navigation, Robotics System, ROS, UAV
- Data in folders: lifted rotating, diagonal rotating, pitching

Description: MATLAB code used to create Fig. 5B, SA-F, Table S4

- Code: SLF_terrestrial_self_righting_Fig5BSAFTableS4.m
- Data: SLF self-righting video summary statistics.xlsx

Description: MATLAB code used to create Tables S2 and S3.

- Code: SLF_terrestrial_self_righting_TableS2S3
- Data: SLF self-righting video summary statistics.xlsx

Description: MATLAB code used to create Fig. S2G-I

- Code: FigS2GHIRightingMethodStatistics.m
- Data: SLF self-righting video summary statistics.xlsx

Description: MATLAB code used to create Figs S9-13 and Table S6.

- Code: analyze_leg_motion.m
- Data: *.mat files created with SLFrightingposeanalyzer.

Description: Code to create Fig. S5 plots

- Code: Fit3DpointsToCircle_FigS5.m
- Data: *.mat files created with SLFrightingposeanalyzer.

Description: MATLAB code used to create Fig 6.

- Code: PElandscape.m
- Data: *.mat files created with SLFrightingposeanalyzer.

### Blender 3D models

Description: 3D Blender model .stl files used to create triangular meshes.

SLFBody3rdinstar.stl	3^rd^ instar body only

SLFBody4thinstar.stl	4^th^ instar body only

SLFSWholeInsect_Righted.stl	whole insect in righted pose

### Code used to make Euler angle figures

EulerAngleFigs.m
