## Supplementary material for "Using pose estimation and 3D rendered models to study leg-mediated self-righting by lanternflies": Dataset S1: EulerAngleFigs.m

```
% creates Fig. S1 showing how Euler angle rotations work, and how these
% rotations do not commute in 3D

clear; close all;

linecolors = lines(7);
BlueLines = linecolors(1,:); RedLines = linecolors(7,:);
GreenLines = linecolors(5,:);

body_length = 8.9192;  % measured on plot; typical for SLF third instar

axis_length = 0.95*body_length;

az = 122; el = 26;                 % for view camera angle default -37.5, 30

% set up parameters for plotting
Xaxis_label_spacing_factor = 1.1; Yaxis_label_spacing_factor = 1; 
Zaxis_label_spacing_factor = 1;

title_text = '';

% axis directions & lengths to start

Xspatial = [1,0,0]; Yspatial = [0,1,0];
Zspatial = [0,0,1]; 

% used for plotting axes
Xspatial_plot = [axis_length,0,0]; Yspatial_plot = [0,axis_length,0];
Zspatial_plot = [0,0,axis_length]; 

%% Load STL mesh
bodymesh = stlread('SLFBody_HighPoly_rot.stl');

bodyPoints = bodymesh.Points;
bodyConnect = bodymesh.ConnectivityList;

% orient the body principal axes using standard reference orientation

rotmz = rotz(180); rotmx = rotx(180);rotm = rotmz*rotmx;
bodyPointsrot = rotm*bodyPoints';      % now perform 3D rotation on mesh
bodyPointsrot = bodyPointsrot';        % restore correct format
% make an updated mesh
bodymesh = triangulation(bodyConnect,bodyPointsrot);

% rotate so the snout-caudal end (cranial-caudal axis) is along +X
rotm = roty(4-180);

bodyPointsrot = rotm*bodyPoints';      % now perform 3D rotation on mesh
bodyPointsrot = bodyPointsrot';        % restore correct format

% make an updated mesh
bodymesh = triangulation(bodyConnect,bodyPointsrot);

%--------------------------------------------------------------------------
%% Fig. S1A shows three successive rotations in ZY'X'' order
figure('Color','white','Position',[10,10,1100,500]);
tiledlayout(1,4,'TileSpacing','loose')
az=167; el = 22;    % use viewing angles so we can compare with plots in 
                    % the MATLAB QuaternionExample.m which uses the same
                    % angles

%-------------------------
% plot 1 = reference orientation
nexttile;
psi = 0; theta = 0; phi = 0;

% initialize the body frame axis orientation to the reference orientation
% aligned with the spatial frame
xbody = Xspatial; ybody = Yspatial; zbody = Zspatial;

% original mesh
bodymesh = triangulation(bodyConnect,bodyPointsrot);

plot_3D_rotation_figure(bodymesh,Xspatial_plot,...
    Yspatial_plot,Zspatial_plot,title_text,Xaxis_label_spacing_factor,...
    Yaxis_label_spacing_factor,Zaxis_label_spacing_factor,0.75)

% draw the body axes
xbody_plot = axis_length*xbody; ybody_plot = axis_length*ybody;
zbody_plot = axis_length*zbody;
draw_body_axes(xbody_plot,ybody_plot,zbody_plot,0);

title_text = sprintf(' ( \\phi , \\theta , \\psi ) = (%2d^o, %2d^o, %2d^o)\n',...
    phi,theta,psi);
title(title_text);

view(az,el);                        % camera anglepsi = 15;

%------------
% plot 2 -- rotate by psi (yaw)
nexttile;
psi = 30;
Euler_angle = [psi theta phi];

[xbody1,ybody1,zbody1,bodyPointsrotz] = ...
    EulerRotate(Euler_angle,xbody,ybody,zbody,bodyPointsrot);

% make an updated mesh
bodymesh = triangulation(bodyConnect,bodyPointsrotz);

% plot

plot_3D_rotation_figure(bodymesh,Xspatial_plot,...
    Yspatial_plot,Zspatial_plot,title_text,Xaxis_label_spacing_factor,...
    Yaxis_label_spacing_factor,Zaxis_label_spacing_factor,0.75)

% draw the new body axes
xbody_plot = axis_length*xbody; ybody_plot = axis_length*ybody;
zbody_plot = axis_length*zbody;
xbody1_plot = axis_length*xbody1; ybody1_plot = axis_length*ybody1;
zbody1_plot = axis_length*zbody1;
draw_body_axes(xbody1_plot,ybody1_plot,zbody1_plot,1);

CreateCurvedArrow3(0.7*xbody_plot,0.7*xbody1_plot,[0 0 0],15,BlueLines,'b');
CreateCurvedArrow3(0.7*ybody_plot,0.7*ybody1_plot,[0 0 0],15,BlueLines,'b');
CreateCurvedArrow3(0.7*zbody_plot,0.7*zbody1_plot,[0 0 0],15,BlueLines,'b');

title_text = sprintf(' ( \\phi , \\theta , \\psi ) = (%2d^o, %2d^o, %2d^o)\n',...
    phi,theta,psi);
title(title_text);

view(az,el);                        % camera angle

%------------
% rotate by beta (elevation)
nexttile;
theta = 20;
Euler_angle = [psi theta phi];

[xbody2,ybody2,zbody2,bodyPointsrotzy] = ...
    EulerRotate(Euler_angle,xbody,ybody,zbody,bodyPointsrot);

% make an updated mesh
bodymesh = triangulation(bodyConnect,bodyPointsrotzy);

% plot

plot_3D_rotation_figure(bodymesh,Xspatial_plot,...
    Yspatial_plot,Zspatial_plot,title_text,Xaxis_label_spacing_factor,...
    Yaxis_label_spacing_factor,Zaxis_label_spacing_factor,0.75)

% draw all body axes
xbody_plot = axis_length*xbody; ybody_plot = axis_length*ybody;
zbody_plot = axis_length*zbody;
xbody1_plot = axis_length*xbody1; ybody1_plot = axis_length*ybody1;
zbody1_plot = axis_length*zbody1;
xbody2_plot = axis_length*xbody2; ybody2_plot = axis_length*ybody2;
zbody2_plot = axis_length*zbody2;

draw_body_axes(xbody1_plot,ybody1_plot,zbody1_plot,1);
draw_body_axes(xbody2_plot,ybody2_plot,zbody2_plot,2);

CreateCurvedArrow3(0.7*xbody_plot,0.7*xbody1_plot,[0 0 0],15,BlueLines,'b');
CreateCurvedArrow3(0.7*ybody_plot,0.7*ybody1_plot,[0 0 0],15,BlueLines,'b');
CreateCurvedArrow3(0.7*zbody_plot,0.7*zbody1_plot,[0 0 0],15,BlueLines,'b');

CreateCurvedArrow3(0.7*xbody1,0.7*xbody2,[0 0 0],15,RedLines,'r');
CreateCurvedArrow3(0.7*ybody1,0.7*ybody2,[0 0 0],15,RedLines,'r');
CreateCurvedArrow3(0.7*zbody1,0.7*zbody2,[0 0 0],15,RedLines,'r');

title_text = sprintf(' ( \\phi , \\theta , \\psi ) = (%2d^o, %2d^o, %2d^o)\n',...
    phi,theta,psi);
title(title_text);

view(az,el);                        % camera angle

%------------
% rotate by phi (roll)
nexttile;
phi = -50;
Euler_angle = [psi theta phi];

[xbody3,ybody3,zbody3,bodyPointsrotzyx] = ...
    EulerRotate(Euler_angle,xbody,ybody,zbody,bodyPointsrot);

% make an updated mesh
bodymesh = triangulation(bodyConnect,bodyPointsrotzyx);

% plot

plot_3D_rotation_figure(bodymesh,Xspatial_plot,...
    Yspatial_plot,Zspatial_plot,title_text,Xaxis_label_spacing_factor,...
    Yaxis_label_spacing_factor,Zaxis_label_spacing_factor,0.75)

% draw the body axes
xbody_plot = axis_length*xbody; ybody_plot = axis_length*ybody;
zbody_plot = axis_length*zbody;
xbody1_plot = axis_length*xbody1; ybody1_plot = axis_length*ybody1;
zbody1_plot = axis_length*zbody1;
xbody2_plot = axis_length*xbody2; ybody2_plot = axis_length*ybody2;
zbody2_plot = axis_length*zbody2;
xbody3_plot = axis_length*xbody3; ybody3_plot = axis_length*ybody3;
zbody3_plot = axis_length*zbody3;
draw_body_axes(xbody1_plot,ybody1_plot,zbody1_plot,1);
draw_body_axes(xbody2_plot,ybody2_plot,zbody2_plot,2);
draw_body_axes(xbody3_plot,ybody3_plot,zbody3_plot,3);

CreateCurvedArrow3(0.7*xbody_plot,0.7*xbody1_plot,[0 0 0],15,BlueLines,'b');
CreateCurvedArrow3(0.7*ybody_plot,0.7*ybody1_plot,[0 0 0],15,BlueLines,'b');
CreateCurvedArrow3(0.7*zbody_plot,0.7*zbody1_plot,[0 0 0],15,BlueLines,'b');

CreateCurvedArrow3(0.7*xbody1_plot,0.7*xbody2_plot,[0 0 0],15,RedLines,'r');
CreateCurvedArrow3(0.7*ybody1_plot,0.7*ybody2_plot,[0 0 0],15,RedLines,'r');
CreateCurvedArrow3(0.7*zbody1_plot,0.7*zbody2_plot,[0 0 0],15,RedLines,'r');

CreateCurvedArrow3(0.7*xbody3_plot,0.7*xbody2_plot,[0 0 0],15,GreenLines,'g');
CreateCurvedArrow3(0.7*ybody3_plot,0.7*ybody2_plot,[0 0 0],15,GreenLines,'g');
CreateCurvedArrow3(0.7*zbody3_plot,0.7*zbody2_plot,[0 0 0],15,GreenLines,'g');

title_text = sprintf(' ( \\phi , \\theta , \\psi ) = (%2d^o, %2d^o, %2d^o)\n',...
    phi,theta,psi);
title(title_text);

view(az,el);                        % camera angle

saveas(gcf,'FigS1A.png')
% Done with Fig. S1A Euler angle rotations

%--------------------------------------------------------------------------
%% Fig. S1B show two successive rotations made in different orders
figure('Color','white','Position',[10,10,500,500]);
tiledlayout(2,2,'TileSpacing','none','Padding','none')

az = 122; el = 26;                  % for view camera angle default -37.5, 30

phi = 90; theta = 90; psi = 0;      % Euler angles to rotate by in correct and incorrect order

showtitles = 0; title_text = '';     % don't display the titles

%%%%%%%%%%%%%%%%%%%%%%%%%%%%%%%%%%%%%%%%%%%%%%%%%%%%%%%%%%%%%%%%%%%%%%%%%%%
% FIG 1SB top plots: theta (pitch) then phi (roll) -- correct order
%-------------
% make first top plot -- rotate by theta (pitch) about +y axis
nexttile;

% initialize the body frame axis orientation to the reference orientation
% aligned with the spatial frame
xbody = Xspatial; ybody = Yspatial; zbody = Zspatial;

Euler_angle = [0 theta 0];

[xbody1,ybody1,zbody1,bodyPointsroty] = ...
    EulerRotate(Euler_angle,xbody,ybody,zbody,bodyPointsrot);

% make an updated mesh
bodymesh = triangulation(bodyConnect,bodyPointsroty);

% plot result of first rotation
plot_3D_rotation_figure(bodymesh,Xspatial_plot,...
    Yspatial_plot,Zspatial_plot,title_text,Xaxis_label_spacing_factor,...
    Yaxis_label_spacing_factor,Zaxis_label_spacing_factor,0.75)
hold on;

% draw the body axes
xbody_plot = axis_length*xbody; ybody_plot = axis_length*ybody;
zbody_plot = axis_length*zbody;
xbody1_plot = axis_length*xbody1; ybody1_plot = axis_length*ybody1;
zbody1_plot = axis_length*zbody1;

draw_body_axes(xbody1_plot,ybody1_plot,zbody1_plot,1);

CreateCurvedArrow3(0.7*Xspatial_plot,0.7*xbody1_plot,[0 0 0],15,BlueLines,'b');
CreateCurvedArrow3(0.7*Zspatial_plot,0.7*zbody1_plot,[0 0 0],15,BlueLines,'b');

if showtitles 
    title_text = sprintf('1st rotation: \\theta = %2d^o about y\n',theta);
    title(title_text); 
end
view(az,el);                        % camera angle
%-------------
% make second top plot -- roll by phi about +x axis

nexttile;

Euler_angle = [0 theta phi];            % next rotate by both theta & phi (roll)
[xbody2,ybody2,zbody2,bodyPointsrotyx] = ...
    EulerRotate(Euler_angle,xbody,ybody,zbody,bodyPointsrot);

% make an updated mesh
bodymesh = triangulation(bodyConnect,bodyPointsrotyx);

% plot result of second rotation
plot_3D_rotation_figure(bodymesh,Xspatial_plot,...
    Yspatial_plot,Zspatial_plot,title_text,Xaxis_label_spacing_factor,...
    Yaxis_label_spacing_factor,Zaxis_label_spacing_factor,0.75)
hold on;

% draw the body axes
xbody2_plot = axis_length*xbody2; ybody2_plot = axis_length*ybody2;
zbody2_plot = axis_length*zbody2;

draw_body_axes(xbody1_plot,ybody1_plot,zbody1_plot,1);
draw_body_axes(xbody2_plot,ybody2_plot,zbody2_plot,2);

CreateCurvedArrow3(-0.7*Yspatial_plot,0.7*Xspatial_plot,[0 0 0],15,RedLines,'r');
CreateCurvedArrow3(0.7*Xspatial_plot,0.7*Yspatial_plot,[0 0 0],15,RedLines,'r');

if showtitles 
    title_text = sprintf('2nd rotation: \\phi = %2d^o about x''\n',phi);
    title(title_text); 
end
view(az,el);                        % camera angle
%%%%%%%%%%%%%%%%%%%%%%%%%%%%%%%%%%%%%%%%%%%%%%%%%%%%%%%%%%%%%%%%%%%%%%%%%%%
% FIG 1SB bottom plots: roll (phi) then pitch (theta) -- reversed order

% initialize the body frame axis orientation to the reference orientation
% aligned with the spatial frame
xbody = Xspatial; ybody = Yspatial; zbody = Zspatial;
%------------------------
% first bottom plot -- roll by phi about x-axis
nexttile;               
Euler_angle = [0 0 phi];
[xbody1,ybody1,zbody1,bodyPointsrotx] = ...
    EulerRotate(Euler_angle,xbody,ybody,zbody,bodyPointsrot);

% make an updated mesh
bodymesh = triangulation(bodyConnect,bodyPointsrotx);

plot_3D_rotation_figure(bodymesh,Xspatial_plot,...
    Yspatial_plot,Zspatial_plot,title_text,Xaxis_label_spacing_factor,...
    Yaxis_label_spacing_factor,Zaxis_label_spacing_factor,0.75)

% draw the body axes
xbody_plot = axis_length*xbody; ybody_plot = axis_length*ybody;
zbody_plot = axis_length*zbody;
xbody1_plot = axis_length*xbody1; ybody1_plot = axis_length*ybody1;
zbody1_plot = axis_length*zbody1;

draw_body_axes(xbody1_plot,ybody1_plot,zbody1_plot,1);

CreateCurvedArrow3(0.7*Xspatial_plot,0.7*xbody1_plot,[0 0 0],15,BlueLines,'b');
CreateCurvedArrow3(0.7*Yspatial_plot,0.7*ybody1_plot,[0 0 0],15,BlueLines,'b');
CreateCurvedArrow3(0.7*Zspatial_plot,0.7*zbody1_plot,[0 0 0],15,BlueLines,'b');

if showtitles 
    title_text = sprintf('1st rotation: \\phi = %2d^o about x',phi);
    title(title_text); 
end
view(az,el);                        % camera angle
%------------------------
% second plot -- rotate by theta about the y' axis, which is the world z
% axis so...
nexttile;               
Euler_angle = [theta 0 0];

[xbody2,ybody2,zbody2,bodyPointsrotxy] = ...
    EulerRotate(Euler_angle,xbody,ybody,zbody,bodyPointsrotx);

% make an updated mesh
bodymesh = triangulation(bodyConnect,bodyPointsrotxy);

% plot result of second rotation
plot_3D_rotation_figure(bodymesh,Xspatial_plot,...
    Yspatial_plot,Zspatial_plot,title_text,Xaxis_label_spacing_factor,...
    Yaxis_label_spacing_factor,Zaxis_label_spacing_factor,0.75)

% draw the body axes
xbody_plot = axis_length*xbody; ybody_plot = axis_length*ybody;
zbody_plot = axis_length*zbody;
xbody1_plot = axis_length*xbody1; ybody1_plot = axis_length*ybody1;
zbody1_plot = axis_length*zbody1;
xbody2_plot = axis_length*xbody2; ybody2_plot = axis_length*ybody2;
zbody2_plot = axis_length*zbody2;
 
draw_body_axes(xbody1_plot,ybody1_plot,zbody1_plot,1);
draw_body_axes(xbody2_plot,ybody2_plot,zbody2_plot,2);

CreateCurvedArrow3(-0.7*Yspatial_plot,0.7*Xspatial_plot,[0 0 0],15,RedLines,'r');
CreateCurvedArrow3(0.7*Xspatial_plot,0.7*Yspatial_plot,[0 0 0],15,RedLines,'r');

if showtitles 
    title_text = sprintf('2nd rotation: \\theta = %2d^o about y''\n',theta);
    title(title_text); 
end
view(az,el);                        % camera angle

%-----------------
saveas(gcf,'FigS1B.png')
% Done with Fig. S1B showing non commutativity of 3D rotations


%% Fig. S1C shows how the Euler angles can disagree with the final 
% orientation angles in the spatial frame
figure('Color','white','Position',[10,10,500,500]);
tiledlayout(2,2,'TileSpacing','none','Padding','none')

az = 122; el = 26;                  % for view camera angle

% initialize the body frame axis orientation to the reference orientation
% aligned with the spatial frame
xbody = Xspatial; ybody = Yspatial; zbody = Zspatial;

phi = 35; theta = -25; psi = 20;      % Euler angles to rotate by
Euler_angle = [psi theta phi];          % yaw,pitch,roll angles

[xbody1,ybody1,zbody1,bodyPointsrotyx] = ...
    EulerRotate(Euler_angle,xbody,ybody,zbody,bodyPointsrot);

% make an updated mesh
bodymesh = triangulation(bodyConnect,bodyPointsrotyx);

% plot result of first rotation
plot_3D_rotation_figure(bodymesh,Xspatial_plot,...
    Yspatial_plot,Zspatial_plot,title_text,Xaxis_label_spacing_factor,...
    Yaxis_label_spacing_factor,Zaxis_label_spacing_factor,0.75)
hold on;

% draw the body axes
xbody1_plot = axis_length*xbody1; ybody1_plot = axis_length*ybody1;
zbody1_plot = axis_length*zbody1;

draw_body_axes(xbody1_plot,ybody1_plot,zbody1_plot,1);

% compute α, β, γ (bank, elevation, heading) defined with respect to the XYZ axes
alpha = acos(dot(xbody1,Xspatial))*180/pi; 
beta = acos(dot(ybody1,Yspatial))*180/pi;
gamma = acos(dot(zbody1,Zspatial))*180/pi;

% compute the angle of the lateral (ybody''') pitch axis with the horizontal
% XY plane
lataxis_w_z = 180/pi*acos(ybody1(3));  % angle between ybody and XY ground plane
if lataxis_w_z <= 90
    latXY = 90 - abs(lataxis_w_z);
else
    latXY = abs(lataxis_w_z)-90;
end

% compute the angle of the cranial-caudal (xbody''') axis with the horizontal
% XY plane (elevation)
dvaxis_w_z = 180/pi*acos(xbody1(3));  % angle between xbody and XY ground plane
if dvaxis_w_z <= 90
    elev = 90 - abs(dvaxis_w_z);
else
    elev = abs(dvaxis_w_z)-90;
end

% bearing angle between X and projected dv axis on XY plane (deg)
bearing = acos(xbody1(1))*180/pi; 

% draw dashed lines showing their projections onto the horizontal plane

% get length of projection of each axis on the plane

% the factor of *0.81 takes into account how draw_body_axes rescales the
% axes after each distinct rotation to avoid overlapping different axis
% sets plotted successively in different colors (e.g., xyz, x'y'z', etc.)

% draw dashed projection and thin projected axis on XY plane
plot3(xbody1_plot(1)*0.81*[1,1],xbody1_plot(2)*0.81*[1,1],...
    [0,xbody1_plot(3)*0.81],'--','Color',BlueLines);

plot3(xbody1_plot(1)*0.81*[1,1],xbody1_plot(2)*0.81*[0,1],...
    [0,0],'--','Color',BlueLines);

plot3([0,xbody1_plot(1)*0.81],[0,xbody1_plot(2)*0.81],[0,0],'-',...
    'Color',BlueLines);

% draw dashed projection and thin projected axis on XY plane
plot3(ybody1_plot(1)*0.81*[1,1],ybody1_plot(2)*0.81*[1,1],...
    [ybody1_plot(3)*0.81,0],'--','Color',BlueLines);
plot3([0,ybody1_plot(1)*0.81],[0,ybody1_plot(2)*0.81],[0,0],'-',...
    'Color',BlueLines);

text1 = ['(roll,pitch,yaw) = (%2.1f^o, %2.1f^o, %2.1f^o)\n' ...
    '(\\alpha,\\beta,\\gamma) = (%2.1f^o, %2.1f^o, %2.1f^o)'];

title1_text = sprintf(text1,phi,theta,psi,alpha,beta,gamma);

title({title1_text}); 

view(az,el);                        % camera angle

saveas(gcf,'FigS1C.png')
% Done with Fig. showing how Euler angles of pitch and roll do not agree
% with the angle of the specimens wrt XYZ in spatial frame after rotations


%%*************************************************************************
% supporting functions
%%*************************************************************************
%% --- Creates a curved arrow
% https://stackoverflow.com/questions/25895072/curved-arrows-in-matlab
% usage: hold on; CreateCurvedArrow3(0.3*[1 0 0], 0.3*[0 1 0]);
% from: Starting position - (x,y,z) upplet
% to: Final position - (x,y,z) upplet
% center: Center of arc - (x,y,z) upplet => by default the origin
% count: The number of segment to draw the arrow => by default 15
function [h] = CreateCurvedArrow3(from,to,center,count,arrowcolor,arrowcolorname)
%[
% Inputs
if from == to
    display('cannot draw arrow');
else
    display('trying to plot arrows')
    % fill in missing parameters
    if (nargin < 4), count = 15; end
    if (nargin < 3), center = [0 0 0]; end

    center = center(:); from = from(:); to = to(:);

    % Start, stop and normal vectors
    start = from - center; rstart = norm(start);
    stop = to - center; rstop = norm(stop);
    angle = atan2(norm(cross(start,stop)), dot(start,stop));
    normal = cross(start, stop); normal = normal / norm(normal);

    % Compute intermediate points by rotating 'start' vector
    % toward 'end' vector around 'normal' axis
    % See: http://inside.mines.edu/fs_home/gmurray/ArbitraryAxisRotation/
    phiAngles = linspace(0, angle, count);
    r = linspace(rstart, rstop, count) / rstart;
    intermediates = zeros(3, count);
    a = center(1); b = center(2); c = center(3);
    u = normal(1); v = normal(2); w = normal(3);
    x = from(1); y = from(2); z = from(3);
    for ki = 1:count
        phi = phiAngles(ki);
        cosp = cos(phi); sinp = sin(phi);
        T = [(u^2+(v^2+w^2)*cosp)  (u*v*(1-cosp)-w*sinp)  (u*w*(1-cosp)+v*sinp) ((a*(v^2+w^2)-u*(b*v+c*w))*(1-cosp)+(b*w-c*v)*sinp); ...
            (u*v*(1-cosp)+w*sinp) (v^2+(u^2+w^2)*cosp)   (v*w*(1-cosp)-u*sinp) ((b*(u^2+w^2)-v*(a*u+c*w))*(1-cosp)+(c*u-a*w)*sinp); ...
            (u*w*(1-cosp)-v*sinp) (v*w*(1-cosp)+u*sinp)  (w^2+(u^2+v^2)*cosp)  ((c*(u^2+v^2)-w*(a*u+b*v))*(1-cosp)+(a*v-b*u)*sinp); ...
            0                    0                      0                                1                               ];
        intermediate = T * [x;y;z;r(ki)];
        intermediates(:,ki) = intermediate(1:3);
    end

    % Draw the curved line
    % Can be improved of course with hggroup etc...
    X = intermediates(1,:);
    Y = intermediates(2,:);
    Z = intermediates(3,:);
    tf = ishold;
    if (~tf), hold on; end
    h = line(X,Y,Z,'Color',arrowcolor,'LineWidth',1.5); hold on;
    %*% replace with arrow3
    % arrow3(X,Y,Z,[arrowcolorname '-' '1.5'])
    quiver3(X(end-1), Y(end-1), Z(end-1), X(end)-X(end-1),...
        Y(end)-Y(end-1), Z(end)-Z(end-1),1,'Color',arrowcolor,'LineWidth',1.5);
end
end

%%*************************************************************************
% rotate by Euler angles in 3D
function [xbody,ybody,zbody,bodyPointsrot] = ...
    EulerRotate(Euler_angle,xbody,ybody,zbody,bodyPoints)

    % convert degrees to radians first
    Euler_angle = Euler_angle*pi/180;

   % --------------create rotation matrix rotm & -------------------------
    % ------------------ rotate the body axes -----------------------------
    % row matrix of intrinsic (point convention) Euler rotation angles in
    % order that agrees with the ZYX default order: [zAngle yAngle xAngle]
    % = [psi theta phi] eul2rotm( ) comes from the Robotics Toolbox which
    % treats rotations as active vector rotations ('point' in the doc).
    % MATLAB: "For a given axis and angle of rotation, point rotation and
    % frame rotation define equivalent angular displacement but in opposite
    % directions."
    rotm = eul2rotm(Euler_angle);
    xbody = rotm*xbody'; xbody = xbody';
    ybody = rotm*ybody'; ybody = ybody';
    zbody = rotm*zbody'; zbody = zbody';
    % ------------------ rotate the mesh ----------------------------------
    bodyPointsrot = rotm*bodyPoints';      
    bodyPointsrot = bodyPointsrot';        % transpose for correct format
end
%%*************************************************************************
function plot_3D_rotation_figure(bodymesh,Xspatial_plot,...
    Yspatial_plot,Zspatial_plot,title_text,Xaxis_label_spacing_factor,...
    Yaxis_label_spacing_factor,Zaxis_label_spacing_factor,facealpha)

    % axis labels
    Xtext = 'X'; Ytext = 'Y'; Ztext = 'Z';
    
    % plot body mesh
    colormap gray; brighten(.8);        % display mesh in grayscale in lighter colors
    trimesh(bodymesh,'FaceAlpha',facealpha,'FaceColor','interp',...
        'LineStyle','none');
    hold on;
    
    title(title_text,'FontSize',12,'FontWeight','Normal'); % unbold title text
    
    % draw spatial frame XYZ axes
    
    quiver3(0,0,0,Xspatial_plot(1),0.9*Xspatial_plot(2),0.9*Xspatial_plot(3),'k','LineWidth',1.5);
    quiver3(0,0,0,Yspatial_plot(1),Yspatial_plot(2),Yspatial_plot(3),'k','LineWidth',1.5);
    quiver3(0,0,0,Zspatial_plot(1),Zspatial_plot(2),Zspatial_plot(3),'k','LineWidth',1.5);
    
    % label XYZ axes
    % plot text labels for each axis
    text_offset = 0.05*norm(Xspatial_plot);
    
    text(4*text_offset+Xspatial_plot(1),Xspatial_plot(2),Xspatial_plot(3),Xtext,'FontSize',13);
    text(Yspatial_plot(1),text_offset+Yspatial_plot(2),Yspatial_plot(3),Ytext,'FontSize',13);
    text(Zspatial_plot(1),Zspatial_plot(2),text_offset+Zspatial_plot(3),Ztext,'FontSize',13);
    
    % hide axes
    h = gca; 
    h.XAxis.Visible = 'off';h.YAxis.Visible = 'off';h.ZAxis.Visible = 'off';
    
    l = light('Position',[-0.1 0.6 0.8],'Style','infinite');
    lighting gouraud; material shiny;
    axis equal; grid off;

end
%%*************************************************************************
function draw_body_axes(xbody,ybody,zbody,rotorder)
% draw body axes with colors, labels matched to the order of rotations
% (1st, 2nd, 3rd)
    htmlgray = [128 128 128]/255;
    linecolors = lines(7);
    BlueLines = linecolors(1,:); RedLines = linecolors(7,:);
    GreenLines = linecolors(5,:);

    switch rotorder
        case 0          % no rotations -- reference orientation
            axisfactor = 0.90;                  % rescales axis lengths
            axiscolor = htmlgray; 
            xtext = 'x'; ytext = 'y'; ztext = 'z';
        case 1          % after 1st rotation
            axisfactor = 0.90;                  % rescales axis lengths
            axiscolor = BlueLines;  
            xtext = ['x' char(39)];ytext = ['y' char(39)];
            ztext = ['z' char(39)];
        case 2          % after 2nd rotation
            axisfactor = 0.90;                  % rescales axis lengths
            axiscolor = RedLines;  
            xtext = ['x' char(39) char(39)];ytext = ['y' char(39) char(39)];
            ztext = ['z' char(39) char(39)];
        case 3          % after 3rd rotation
            axisfactor = 0.90;                  % rescales axis lengths
            axiscolor = GreenLines;  
            xtext = ['x' char(39) char(39) char(39)];
            ytext = ['y' char(39) char(39) char(39)];
            ztext = ['z' char(39) char(39) char(39)];
    end

    % rescale axis lengths
    xbody = axisfactor*xbody; ybody = axisfactor*ybody; zbody = axisfactor*zbody;

    quiver3(0,0,0,xbody(1),xbody(2),xbody(3),'LineWidth',1.5,'Color',axiscolor);
    quiver3(0,0,0,ybody(1),ybody(2),ybody(3),'LineWidth',1.5,'Color',axiscolor);
    quiver3(0,0,0,zbody(1),zbody(2),zbody(3),'LineWidth',1.5,'Color',axiscolor);
    
    % plot text labels for each axis
    text_offset = 0.35*norm(xbody);

    %*% remove child axis labels
    xtext = '';ytext = '';ztext = '';
        
    text(text_offset+xbody(1),1.3*text_offset+xbody(2),1.3*text_offset+xbody(3),xtext,'FontSize',13,'Color',axiscolor);
    text(text_offset+ybody(1),ybody(2),text_offset+ybody(3),ytext,'FontSize',13,'Color',axiscolor);
    text(text_offset+zbody(1),text_offset+zbody(2),zbody(3),ztext,'FontSize',13,'Color',axiscolor);

end
%%*************************************************************************
function hn=arrow3(p1,p2,s,w,h,ip,alpha,beta)
    % ARROW3 (R13)
    %   ARROW3(P1,P2) draws lines from P1 to P2 with directional arrowheads.
    %   P1 and P2 are either nx2 or nx3 matrices.  Each row of P1 is an
    %   initial point, and each row of P2 is a terminal point.
    %
    %   ARROW3(P1,P2,S,W,H,IP,ALPHA,BETA) can be used to specify properties
    %   of the line, initial point marker, and arrowhead.  S is a character
    %   string made with one element from any or all of the following 3
    %   columns:
    %
    %     Color Switches      LineStyle            LineWidth
    %     ------------------  -------------------  --------------------
    %     k  blacK (default)  -  solid (default)   0.5 points (default)
    %     y  Yellow           :  dotted            0   no lines
    %     m  Magenta          -. dashdot           /   LineWidthOrder
    %     c  Cyan             -- dashed
    %     r  Red              *  LineStyleOrder            _______ __  
    %     g  Green                                       ^        |    
    %     b  Blue                                       / \       |    
    %     w  White                        Arrowhead    /   \   Height  
    %     a  Asparagus                                /     \     |    
    %     d  Dark gray                               /       \    |    
    %     e  Evergreen                              /___   ___\ __|__  
    %     f  Firebrick                             |    | |    |       
    %     h  Hot pink                              |-- Width --|       
    %     i  Indigo                                |    | |    |       
    %     j  Jade                                       | |            
    %     l  Light gray                                 | |            
    %     n  Nutbrown                                   | |            
    %     p  Pear                                       | |            
    %     q  kumQuat                      Line       -->| |<--LineWidth
    %     s  Sky blue                                   | |            
    %     t  Tawny                                      | |            
    %     u  bUrgundy                                   | |            
    %     v  Violet                                     | |            
    %     z  aZure                                      | |            
    %     x  random                       Initial      /   \           
    %     o  colorOrder                   Point    -->(     )<--IP     
    %     |  magnitude                    Marker       \_ _/           
    %
    %     -------------------------------------------------------------
    %                          Color Equivalencies
    %     -------------------------------------------------------------
    %     ColorOrder     Arrow3         |     Simulink       Arrow3
    %     ----------     ----------     |     ----------     ----------
    %     Color1         Blue           |     LightBlue      aZure
    %     Color2         Evergreen      |     DarkGreen      Asparagus
    %     Color3         Red            |     Orange         kumQuat
    %     Color4         Sky blue       |     Gray           Light gray
    %     Color5         Violet         |
    %     Color6         Pear           |
    %     Color7         Dark gray      |
    %     -------------------------------------------------------------
    %
    %   The components of S may be specified in any order.  Invalid
    %   characters in S will be ignored and replaced by default settings.
    %
    %   Prefixing the color code with '_' produces a darker shade, e.g.
    %   '_t' is dark tawny; prefixing the color code with '^' produces a
    %   lighter shade, e.g. '^q' is light kumquat.  The relative brightness
    %   of light and dark color shades is controlled by the scalar parameter
    %   BETA.  Color code prefixes do not affect black (k), white (w), or
    %   the special color switches (xo|).
    %
    %   ColorOrder may be achieved in two fashions:  The user may either
    %   set the ColorOrder property (using RGB triples) or define the
    %   global variable ColorOrder (using a string of valid color codes).
    %   If the color switch is specified with 'o', and the global variable
    %   ColorOrder is a string of color codes (color switches less 'xo|',
    %   optionally prefixed with '_' or '^'), then the ColorOrder property
    %   will be set to the sequence of colors indicated by the ColorOrder
    %   variable.  The color sequence 'bersvpd' matches the default
    %   ColorOrder property.  If the color switch is specified with 'o', and
    %   the global variable ColorOrder is empty or invalid, then the current
    %   ColorOrder property will be used.  Note that the ColorOrder variable
    %   takes precedence over the ColorOrder property.
    %
    %   The magnitude color switch is used to visualize vector magnitudes
    %   in conjunction with a colorbar.  If the color switch is specified
    %   with '|', colors are linearly interpolated from the current ColorMap
    %   according to the length of the associated line.  This option sets
    %   CLim to [MinM,MaxM], where MinM and MaxM are the minimum and maximum
    %   magnitudes, respectively.
    %
    %   The current LineStyleOrder property will be used if LineStyle is
    %   specified with '*'.  MATLAB cycles through the line styles defined
    %   by the LineStyleOrder property only after using all colors defined
    %   by the ColorOrder property.  If however, the global variable
    %   LineWidthOrder is defined, and LineWidth is specified with '/',
    %   then each line will be drawn with sequential color, linestyle, and
    %   linewidth.
    %
    %   W (default = 1) is a vector of arrowhead widths; use W = 0 for no
    %   arrowheads.  H (default = 3W) is a vector of arrowhead heights.  If
    %   vector IP is neither empty nor negative, initial point markers will
    %   be plotted with diameter IP; for default diameter W, use IP = 0.
    %   The units of W, H and IP are 1/72 of the PlotBox diagonal.
    %
    %   ALPHA (default = 1) is a vector of FaceAlpha values ranging between
    %   0 (clear) and 1 (opaque).  FaceAlpha is a surface (arrowhead and
    %   initial point marker) property and does not affect lines.  FaceAlpha
    %   is not supported for 2D rendering.
    %
    %   BETA (default = 0.4) is a scalar that controls the relative
    %   brightness of light and dark color shades, ranging between 0 (no
    %   contrast) and 1 (maximum contrast).
    %
    %   ARROW3(P1,P2,S,W,H,'cone',...) will plot cones with bases centered
    %   on P1 in the direction given by P2.  In this instance, P2 is
    %   interpreted as a direction vector instead of a terminal point.
    %   Neither initial point markers nor lines are plotted with the 'cone'
    %   option.
    %
    %   HN = ARROW3(P1,P2,...) returns a vector of handles to line and
    %   surface objects created by ARROW3.
    %
    %   ARROW3 COLORS will plot a table of named colors with default
    %   brightness.  ARROW3('colors',BETA) will plot a table of named
    %   colors with brightness BETA.
    %
    %   ARROW3 attempts to preserve the appearance of existing axes.  In
    %   particular, ARROW3 will not change XYZLim, View, or CameraViewAngle.
    %   ARROW3 does not, however, support stretch-to-fill scaling.  AXIS
    %   NORMAL will restore the current axis box to full size and remove any
    %   restrictions on the scaling of units, but will likely result in
    %   distorted arrowheads and initial point markers.  See
    %   (arrow3_messes_up_my_plots.html).
    %
    %   If a particular aspect ratio or variable limit is required, use
    %   DASPECT, PBASPECT, AXIS, or XYZLIM commands before calling ARROW3.
    %   Changing limits or aspect ratios after calling ARROW3 may alter the
    %   appearance of arrowheads and initial point markers.  ARROW3 sets
    %   XYZCLimMode to manual for all plots, sets DataAspectRatioMode to
    %   manual for linear plots, and sets PlotBoxAspectRatioMode to manual
    %   for log plots and 3D plots.  CameraViewAngleMode is also set to
    %   manual for 3D plots.
    %
    %   ARROW3 UPDATE will restore the appearance of arrowheads and
    %   initial point markers that have become corrupted by changes to
    %   limits or aspect ratios.  ARROW3('update',SF) will redraw initial
    %   point markers and arrowheads with scale factor SF.  If SF has one
    %   element, SF scales W, H and IP.  If SF has two elements, SF(1)
    %   scales W and IP, and SF(2) scales H.  If SF has three elements,
    %   SF(1) scales W, SF(2) scales H, and SF(3) scales IP.  All sizes are
    %   relative to the current PlotBox diagonal.
    %
    %   ARROW3 UPDATE COLORS will update the magnitude coloring of
    %   arrowheads, initial point markers, and lines to conform to the
    %   current ColorMap.
    %
    %   HN = ARROW3('update',...) returns a vector of handles to updated
    %   objects.
    %
    %   EXAMPLES:
    %
    %     % 2D vectors
    %     arrow3([0 0],[1 3])
    %     arrow3([0 0],[1 2],'-.e')
    %     arrow3([0 0],[10 10],'--x2',1)
    %     arrow3(zeros(10,2),50*rand(10,2),'x',1,3)
    %     arrow3(zeros(10,2),[10*rand(10,1),500*rand(10,1)],'u')
    %     arrow3(10*rand(10,2),50*rand(10,2),'x',1,[],1)
    %
    %     % 3D vectors
    %     arrow3([0 0 0],[1 1 1])
    %     arrow3(zeros(20,3),50*rand(20,3),'--x1.5',2)
    %     arrow3(zeros(100,3),50*rand(100,3),'x',1,3)
    %     arrow3(zeros(10,3),[10*rand(10,1),500*rand(10,1),50*rand(10,1)],'a')
    %     arrow3(10*rand(10,3),50*rand(10,3),'x',[],[],0)
    %
    %     % 3D animation
    %     t=(0:pi/40:8*pi)'; u=cos(t); v=sin(t);
    %     plot3(20*t,u,v)
    %     axis([0,600,-1.5,1.5,-1.5,1.5])
    %     grid on, view(35,25)
    %     hold on
    %     pbaspect([1.8,1.4,1])
    %     arrow3(zeros(3),diag([500,1.5,1.5]),'l',0.7,[],0)
    %     p=[20*t,u,v]; inc=4:1:length(t);
    %     p2=p(inc,:); p1=p(inc-1,:);
    %     hn=arrow3(p1(1,:),p2(1,:),'0_b',0.7);
    %     for i=2:1:length(p1)
    %       delete(hn)
    %       hn=arrow3(p1(i,:),p2(i,:),'0_b',0.7);
    %       pause(0.01)
    %     end
    %     hold off
    %
    %     % Cone plot
    %     t=(pi/8:pi/8:2*pi)'; p1=[cos(t) sin(t) t]; p2=repmat([0 0 1],16,1);
    %     arrow3(p1,p2,'x',2,4,'cone'), hold on
    %     plot3(p1(:,1),p1(:,2),p1(:,3)), hold off
    %     pause % change cone size
    %     arrow3('update',[1,2])
    %
    %     % Just for fun
    %     arrow3(zeros(100,3),50*rand(100,3),'x',8,4,[],0.95)
    %     light('position',[-10 -10 -10],'style','local')
    %     light('position',[60,60,60]), lighting gouraud
    %
    %     % ColorOrder variable, color code prefixes, and Beta
    %     global ColorOrder, ColorOrder='^ui^e_hq^v';
    %     theta=[0:pi/22:pi/2]';
    %     arrow3(zeros(12,2),[cos(theta),sin(theta)],'1.5o',1.5,[],[],[],0.5)
    %
    %     % ColorOrder property, LineStyleOrder, and LineWidthOrder
    %     global ColorOrder, ColorOrder=[];
    %     set(gca,'ColorOrder',[1,0,0;0,0,1;0.25,0.75,0.25;0,0,0])
    %     set(gca,'LineStyleOrder',{'-','--','-.',':'})
    %     global LineWidthOrder, LineWidthOrder=[1,2,4,8];
    %     w=[1,2,3,4]; h=[4,6,4,2];
    %     arrow3(zeros(4,2),[10*rand(4,1),500*rand(4,1)],'o*/',w,h,0)
    %
    %     % Magnitude coloring
    %     colormap spring
    %     arrow3(20*randn(20,3),50*randn(20,3),'|',[],[],0)
    %     set(gca,'color',0.7*[1,1,1])
    %     set(gcf,'color',0.5*[1,1,1]), grid on, colorbar
    %     pause % change the ColorMap and update colors
    %     colormap hot
    %     arrow3('update','colors')
    %
    %     % LogLog plot
    %     set(gca,'xscale','log','yscale','log');
    %     axis([1e2,1e8,1e-2,1e-1]); hold on
    %     p1=repmat([1e3,2e-2],15,1);
    %     q1=[1e7,1e6,1e5,1e4,1e3,1e7,1e7,1e7,1e7,1e7,1e7,1e6,1e5,1e4,1e3];
    %     q2=1e-2*[9,9,9,9,9,7,5,4,3,2,1,1,1,1,1]; p2=[q1',q2'];
    %     global ColorOrder, ColorOrder=[];
    %     set(gca,'ColorOrder',rand(15,3))
    %     arrow3(p1,p2,'o'), grid on, hold off
    %
    %     % SemiLogX plot
    %     set(gca,'xscale','log','yscale','linear');
    %     axis([1e2,1e8,1e-2,1e-1]); hold on
    %     p1=repmat([1e3,0.05],15,1);
    %     q1=[1e7,1e6,1e5,1e4,1e3,1e7,1e7,1e7,1e7,1e7,1e7,1e6,1e5,1e4,1e3];
    %     q2=1e-2*[9,9,9,9,9,7,5,4,3,2,1,1,1,1,1]; p2=[q1',q2'];
    %     arrow3(p1,p2,'x'), grid on, hold off
    %
    %     % SemiLogY plot
    %     set(gca,'xscale','linear','yscale','log');
    %     axis([2,8,1e-2,1e-1]); hold on
    %     p1=repmat([3,2e-2],17,1);
    %     q1=[7,6,5,4,3,7,7,7,7,7,7,7,7,6,5,4,3];
    %     q2=1e-2*[9,9,9,9,9,8,7,6,5,4,3,2,1,1,1,1,1]; p2=[q1',q2'];
    %     set(gca,'LineStyleOrder',{'-','--','-.',':'})
    %     arrow3(p1,p2,'*',1,[],0), grid on, hold off
    %
    %     % Color tables
    %     arrow3('colors')           % default color table
    %     arrow3('colors',0.3)       % low contrast color table
    %     arrow3('colors',0.5)       % high contrast color table
    %
    %     % Update initial point markers and arrowheads
    %     % relative to the current PlotBox diagonal
    %     arrow3('update')           % redraw same size
    %     arrow3('update',2)         % redraw double size
    %     arrow3('update',0.5)       % redraw half size
    %     arrow3('update',[0.5,2,1]) % redraw W half size,
    %                                %        H double size, and
    %                                %        IP same size
    %
    %     See also (arrow3_examples.html), (arrow3_messes_up_my_plots.html).
    
    %   Copyright(c)2002-2020 Version 5.18.1
    %     Tom Davis
    %     Jeff Chang
    
    %   Revision History:
    %
    %     08/08/20 - Address ColorOrder, LineStyleOrder property changes
    %                in R2019b. (TD)
    %     06/19/20 - Remove deprecated zbuffer renderer. (TD)
    %     11/02/19 - Address ColorOrder property changes in R2019b. (TD)
    %     08/28/18 - Use explicit EdgeColor. (TD)
    %     01/15/13 - Use AppData instead of UserData. (TD)
    %     07/27/11 - Added animation example. (TD)
    %     05/13/09 - Corrected spelling errors (TD)
    %     03/16/08 - Updated contact information (TD)
    %     10/23/07 - Corrected zero magnitude exclusion (TD)
    %     09/08/07 - Added cone plot option; removed adaptive grid
    %                spacing; corrected scale factor; removed "nearly"
    %                tight limits (TD)
    %     07/24/07 - Ignore zero-magnitude input (TD)
    %     07/08/07 - Modified named colors to match named Simulink
    %                colors; added light and dark shades for basic
    %                colors (ymcrgb) (TD)
    %     07/01/07 - Modified named colors to match default ColorOrder
    %                colors (TD)
    %     06/24/07 - Error checking for empty P1, P2 (TD)
    %     06/17/07 - Trim colors,W,H,IP,ALPHA to LENGTH(P1) (TD)
    %     05/27/07 - Magnitude coloring and documentation revision (TD)
    %     03/10/07 - Improved code metrics (TD)
    %     02/21/07 - Preserve existing axis appearance;
    %                use relative sizes for W, H, and IP;
    %                removed version checking; minor bug fixes (TD) 
    %     01/09/04 - Replaced calls to LINSPACE, INTERP1, and
    %                COLORMAP (TD)
    %     12/17/03 - Semilog examples, CAXIS support, magnitude
    %                coloring, and color updating; use CData instead
    %                of FaceColor; minor bug fixes (TD)
    %     07/17/03 - Changed 2D rendering from OpenGL to ZBuffer;
    %                defined HN for COLORS and UPDATE options (TD)
    %     02/27/03 - Replaced calls to RANDPERM, VIEW, REPMAT, SPHERE,
    %                and CYLINDER; added ZBuffer for log plots, RESET
    %                for CLA and CLF, and ABS for W and H (TD)
    %     02/01/03 - Added UPDATE scale factor and MATLAB version
    %                checking, replaced call to CROSS (TD)
    %     12/26/02 - Added UserData and UPDATE option (TD)
    %     11/16/02 - Added more named colors, color code prefix,
    %                global ColorOrder, ALPHA , and BETA (TD)
    %     10/12/02 - Added global LineWidthOrder,
    %                vectorized W, H and IP (TD)
    %     10/05/02 - Changed CLF to CLA for subplot support,
    %                added ColorOrder and LineStyleOrder support (TD)
    %     04/27/02 - Minor log plot revisions (TD)
    %     03/26/02 - Added log plot support (TD)
    %     03/24/02 - Adaptive grid spacing control to trade off
    %                appearance vs. speed based on size of matrix (JC)
    %     03/16/02 - Added "axis tight" for improved appearance (JC)
    %     03/12/02 - Added initial point marker (TD)
    %     03/03/02 - Added aspect ratio support (TD)
    %     03/02/02 - Enhanced program's user friendliness (JC)
    %                (lump Color, LineStyle, and LineWidth together)
    %     03/01/02 - Replaced call to ROTATE (TD)
    %     02/28/02 - Modified line plotting,
    %                added linewidth and linestyle (TD)
    %     02/27/02 - Minor enhancements on 3D appearance (JC)
    %     02/26/02 - Minor enhancements for speed (TD&JC)
    %     02/26/02 - Optimize PLOT3 and SURF for speed (TD)
    %     02/25/02 - Return handler, error handling, color effect,
    %                generalize for 2D/3D vectors (JC)
    %     02/24/02 - Optimize PLOT3 and SURF for speed (TD)
    %     02/23/02 - First release (JC&TD)
    
    %-------------------------------------------------------------------------
    % Error Checking
    global LineWidthOrder ColorOrder
    if nargin<8 || isempty(beta), beta=0.4; end
    beta=abs(beta(1)); if nargout, hn=[]; end
    if strcmpi(p1,'colors')                            % plot color table
      if nargin>1, beta=abs(p2(1)); end
      LocalColorTable(1,beta); return
    end
    fig=gcf; ax=gca;
    if strcmpi(p1,'update'), ad=getappdata(ax,'Arrow3');    % update
      LocalLogCheck(ax);
      if size(ad,2)<13, error('Invalid AppData'), end
      setappdata(ax,'Arrow3',[]); sf=[1,1,1]; flag=0;
      if nargin>1
        if strcmpi(p2,'colors'), flag=1;               % update colors
        elseif ~isempty(p2)                            % update surfaces
          sf=p2(1)*sf; n=length(p2(:));
          if n>1, sf(2)=p2(2); if n>2, sf(3)=p2(3); end, end
        end
      end
      H=LocalUpdate(fig,ax,ad,sf,flag); if nargout, hn=H; end, return
    end
    InputError=['Invalid input, type HELP ',upper(mfilename),...
      ' for usage examples'];
    if nargin<2, error(InputError), end
    [r1,c1]=size(p1); [r2,c2]=size(p2);
    if c1<2 || c1>3 || r1*r2==0, error(InputError), end
    if r1~=r2, error('P1 and P2 must have same number of rows'), end
    if c1~=c2, error('P1 and P2 must have same number of columns'), end
    p=sum(abs(p2-p1),2)~=0; cone=0;
    if nargin>5 && ~isempty(ip) && strcmpi(ip,'cone')  % cone plot
      cone=1; p=sum(p2,2)~=0;
      if ~any(p), error('P2 cannot equal 0'), end
      set(ax,'tag','Arrow3ConePlot');
    elseif ~any(p), error('P1 cannot equal P2')
    end
    if ~all(p)
      warning('Arrow3:ZeroMagnitude','Zero magnitude ignored')
      p1=p1(p,:); p2=p2(p,:); [r1,c1]=size(p1);
    end
    n=r1; Zeros=zeros(n,1);
    if c1==2, p1=[p1,Zeros]; p2=[p2,Zeros];
    elseif ~any([p1(:,3);p2(:,3)]), c1=2; end
    L=get(ax,'LineStyleOrder'); C=get(ax,'ColorOrder');
    ST=get(ax,'DefaultSurfaceTag'); LT=get(ax,'DefaultLineTag');
    EC=get(ax,'DefaultSurfaceEdgeColor');
    if strcmp(get(ax,'nextplot'),'add') && strcmp(get(fig,'nextplot'),'add')
      Xr=get(ax,'xlim'); Yr=get(ax,'ylim'); Zr=get(ax,'zlim');
      [xs,ys,xys]=LocalLogCheck(ax); restore=1;
      if xys, mode='auto';
        if any([p1(:,3);p2(:,3)]), error('3D log plot not supported'), end
        if (xs && ~all([p1(:,1);p2(:,1)]>0)) || ...
           (ys && ~all([p1(:,2);p2(:,2)]>0))
           error('Nonpositive log data not supported')
        end
      else, mode='manual';
        if strcmp(get(ax,'WarpToFill'),'on')
          warning('Arrow3:WarpToFill',['Stretch-to-fill scaling not ',...
            'supported;\nuse DASPECT or PBASPECT before calling ARROW3.']);
        end
      end
      set(ax,'XLimMode',mode,'YLimMode',mode,'ZLimMode',mode,...
        'CLimMode','manual');
    else, restore=0; cla reset; xys=0; set(fig,'nextplot','add');
      set(ax,'LineStyleOrder',L);
      if c1==2, azel=[0,90]; else, azel=[-37.5,30]; end
      setappdata(ax,'Arrow3',[]);
      set(ax,'nextplot','add','View',azel);
    end
    
    %-------------------------------------------------------------------------
    % Style Control
    [vc,cn]=LocalColorTable(0); prefix='';
    if nargin<3, [c,ls,lw]=LocalValidateCLSW;% default color, linestyle/width
    else 
      [c,ls,lw]=LocalValidateCLSW(s);
      if length(c)>1, if sum('_^'==c(1)), prefix=c(1); end, c=c(2); end
      if c=='x'                              % random named color (less white)
        [~,i]=sort(rand(1,23)); c=cn(i,:);
      elseif c=='o'                                    % ColorOrder
        if ~isempty(ColorOrder)
          [c,failed]=LocalColorMap(lower(ColorOrder),vc,cn,beta);
          if failed, ColorOrderWarning=['Invalid ColorOrder ',...
            'variable, current ColorOrder property will be used'];
            warning('Arrow3:ColorOrder',ColorOrderWarning)
          else, C=c;
          end
        end, c=C;
      elseif c=='|', map=get(fig,'colormap');          % magnitude coloring
        M=(p1-p2); M=sqrt(sum(M.*M,2)); minM=min(M); maxM=max(M);
        if maxM-minM<1, minM=0; end
        set(ax,'clim',[minM,maxM]); c=LocalInterp(minM,maxM,map,M);
      elseif ~sum(vc==c), c='k'; ColorWarning=['Invalid color switch, ',...
        'default color (black) will be used'];
        warning('Arrow3:Color',ColorWarning)
      end
    end
    if length(c)==1                                    % single color
      c=LocalColorMap([prefix,c],vc,cn,beta);
    end
    set(ax,'ColorOrder',c); c=LocalRepmat(c,[ceil(n/size(c,1)),1]);
    if ls~='*'                                         % LineStyleOrder
      set(ax,'LineStyleOrder',ls);
    end
    ls=get(ax,'LineStyleOrder');
    ls=LocalRepmat(cellstr(ls),[ceil(n/size(ls,1)),1]);
    if lw=='/'                                         % LineWidthOrder
      if ~isempty(LineWidthOrder)
        lw=LineWidthOrder;
      else, lw=0.5; LineWidthOrderWarning=['Undefined LineWidthOrder, ',...
        'default width (0.5) will be used'];
        warning('Arrow3:LineWidthOrder',LineWidthOrderWarning)
      end
    end
    lw=LocalRepmat(lw,[ceil(n/length(lw)),1]);
    if nargin<4 || isempty(w), w=1; end                % width
    w=LocalRepmat(abs(w(:)),[ceil(n/length(w)),1]);
    if nargin<5 || isempty(h), h=3*w; end              % height
    h=LocalRepmat(abs(h(:)),[ceil(n/length(h)),1]);
    if nargin>5 && ~isempty(ip) && ~cone               % ip
      ip=LocalRepmat(ip(:),[ceil(n/length(ip)),1]);
      i=find(ip==0); ip(i)=w(i);
    else, ip=-ones(n,1);
    end
    if nargin<7 || isempty(alpha), alpha=1; end
    a=LocalRepmat(alpha(:),[ceil(n/length(alpha)),1]); % FaceAlpha
    
    %-------------------------------------------------------------------------
    % Log Plot
    if xys
      units=get(ax,'units'); set(ax,'units','points');
      pos=get(ax,'position'); set(ax,'units',units);
      if strcmp(get(ax,'PlotBoxAspectRatioMode'),'auto')
        set(ax,'PlotBoxAspectRatio',[pos(3),pos(4),1]);
      end
      par=get(ax,'PlotBoxAspectRatio');
      set(ax,'DataAspectRatio',[par(2),par(1),par(3)]);
      % map coordinates onto unit square
      q=[p1;p2]; xr=Xr; yr=Yr;
      if xs, xr=log10(xr); q(:,1)=log10(q(:,1)); end
      if ys, yr=log10(yr); q(:,2)=log10(q(:,2)); end
      q=q-LocalRepmat([xr(1),yr(1),0],[2*n,1]);
      dx=xr(2)-xr(1); dy=yr(2)-yr(1);
      q=q*diag([1/dx,1/dy,1]);
      q1=q(1:n,:); q2=q(n+1:end,:);
    else, xs=0; ys=0; dx=0; dy=0; xr=0; yr=0;
    end
    
    %-------------------------------------------------------------------------
    % Line
    set(ax,'DefaultLineTag','arrow3');
    if ~cone
      H1=Zeros;
      for i=1:n
        if lw(i)
          H1(i)=plot3([p1(i,1),p2(i,1)],[p1(i,2),p2(i,2)],...
            [p1(i,3),p2(i,3)],ls{i},'Color',c(i,:),'LineWidth',lw(i));
        end
      end
    else                                               % cone plot
      P=zeros(3*n,3); i=1:n;
      P(3*i-2,:)=p1(i,:); P(3*i-1,:)=p1(i,:); P(3*i,1)=NaN;
      H1=plot3(P(:,1),P(:,2),P(:,3));
    end
    
    %-------------------------------------------------------------------------
    % Scale
    if ~restore, axis tight, end
    ar=get(ax,'DataAspectRatio'); ar=sqrt(3)*ar/norm(ar);
    set(ax,'DataAspectRatioMode','manual');
    if xys, sf=1;
    else, xr=get(ax,'xlim'); yr=get(ax,'ylim'); zr=get(ax,'zlim');
      sf=norm(diff([xr;yr;zr],1,2)./ar')/72;
    end
    
    %-------------------------------------------------------------------------
    % AppData
    c=c(1:n,:); w=w(1:n); h=h(1:n); ip=ip(1:n); a=a(1:n);
    setappdata(ax,'Arrow3',[getappdata(ax,'Arrow3');p1,p2,c,w,h,ip,a]);
    
    %-------------------------------------------------------------------------
    % Arrowhead
    whip=sf*[w,h,ip];
    if xys, whip=whip*sqrt(2)/72; p1=q1; p2=q2; end
    w=whip(:,1); h=whip(:,2); ip=whip(:,3);
    if cone                                            % cone plot
      delete(H1), H1=[];
      p2=p2./LocalRepmat(sqrt(sum(p2.*p2,2)),[1,3]);
      p2=p1+p2.*LocalRepmat(ar,[n,1]).*LocalRepmat(h,[1,3]);
    end
    W=(p1-p2)./LocalRepmat(ar,[n,1]);
    W=W./LocalRepmat(sqrt(sum(W.*W,2)),[1,3]);         % new z direction
    U=[-W(:,2),W(:,1),Zeros];
    N=sqrt(sum(U.*U,2)); i=find(N<eps); j=length(i);
    U(i,:)=LocalRepmat([1,0,0],[j,1]); N(i)=ones(j,1);
    U=U./LocalRepmat(N,[1,3]);                         % new x direction
    V=[W(:,2).*U(:,3)-W(:,3).*U(:,2),...               % new y direction
      W(:,3).*U(:,1)-W(:,1).*U(:,3),...
      W(:,1).*U(:,2)-W(:,2).*U(:,1)];
    m=20;                               % surface grid spacing
    set(ax,'DefaultSurfaceTag','arrow3','DefaultSurfaceEdgeColor','none');
    r=[0;1]; theta=(0:m)/m*2*pi; Ones=ones(1,m+1);
    x=r*cos(theta); y=r*sin(theta); z=r*Ones;
    G=surface(x/2,y/2,z); dar=diag(ar);
    set(G,'EdgeColor','none');
    X=get(G,'XData'); Y=get(G,'YData'); Z=get(G,'ZData');
    H2=Zeros; [j,k]=size(X);
    for i=1:n                           % translate, rotate, and scale
      H2(i)=copyobj(G,ax);
      xyz=[w(i)*X(:),w(i)*Y(:),h(i)*Z(:)]*[U(i,:);V(i,:);W(i,:)]*dar;
      x=reshape(xyz(:,1),j,k)+p2(i,1);
      y=reshape(xyz(:,2),j,k)+p2(i,2);
      z=reshape(xyz(:,3),j,k)+p2(i,3);
      LocalSetSurface(xys,xs,ys,dx,dy,xr,yr,...
        x,y,z,a(i),c(i,:),H2(i),2,m+1);
    end
    delete(G);
    
    %-------------------------------------------------------------------------
    % Initial Point Marker
    if any(ip>0)
      theta=(-m:2:m)/m*pi; phi=(-m:2:m)'/m*pi/2; cosphi=cos(phi);
      x=cosphi*cos(theta); y=cosphi*sin(theta); z=sin(phi)*Ones;
      G=surface(x*ar(1)/2,y*ar(2)/2,z*ar(3)/2);
      set(G,'EdgeColor','none');
      X=get(G,'XData'); Y=get(G,'YData'); Z=get(G,'ZData');
      H3=zeros(n,1);
      for i=1:n                                        % translate
        if ip(i)>0
          H3(i)=copyobj(G,ax);
          x=p1(i,1)+X*ip(i); y=p1(i,2)+Y*ip(i); z=p1(i,3)+Z*ip(i);
          LocalSetSurface(xys,xs,ys,dx,dy,xr,yr,...
            x,y,z,a(i),c(i,:),H3(i),m+1,m+1);
        end
      end, delete(G);
    else, H3=[];
    end
    
    %-------------------------------------------------------------------------
    % Finish
    if restore, xr=Xr; yr=Yr; zr=Zr;
      if xys, set(ax,'DataAspectRatioMode','auto'); end
    else
      axis tight
      xr=get(ax,'xlim'); yr=get(ax,'ylim'); zr=get(ax,'zlim');
      set(ax,'nextplot','replace');
    end
    azel=get(ax,'view');
    if abs(azel(2))==90, c1=2; else, c1=3; end
    set(fig,'Renderer','OpenGL');
    set(ax,'LineStyleOrder',L,'ColorOrder',C,'DefaultLineTag',LT,...
      'DefaultSurfaceTag',ST,'DefaultSurfaceEdgeColor',EC,...
      'xlim',xr,'ylim',yr,'zlim',zr,'clim',get(ax,'CLim'));
    if c1==3, set(ax,'CameraViewAngle',get(ax,'CameraViewAngle'),...
       'PlotBoxAspectRatio',get(ax,'PlotBoxAspectRatio'));
    end
    if nargout, hn=[H1(:);H2(:);H3(:)]; end
end
%-------------------------------------------------------------------------
% Local Functions
%-------------------------------------------------------------------------
% Update
function H=LocalUpdate(fig,ax,ad,sf,flag)
    global ColorOrder
    p1=ad(:,1:3); p2=ad(:,4:6); c=ad(:,7:9); a=ad(:,13);
    w=sf(1)*ad(:,10); h=sf(2)*ad(:,11); ip=sf(3)*ad(:,12);
    H=get(ax,'children'); tag=get(H,'tag'); type=get(H,'type');
    delete(H(strcmp(tag,'arrow3') & strcmp(type,'surface')));
    set(fig,'nextplot','add'); set(ax,'nextplot','add'); H1=[];
    if flag, map=get(fig,'colormap');                  % update colors
      M=(p1-p2); M=sqrt(sum(M.*M,2)); minM=min(M); maxM=max(M);
      H1=H(strcmp(tag,'arrow3') & strcmp(type,'line'));
      MagnitudeWarning=['Cannot perform magnitude coloring on lines ',...
        'that\nwere drawn with a single color, linestyle, and linewidth'];
      if length(H1)>1
        for i=1:length(H1)                             % update line colors
          x=get(H1(i),'xdata'); y=get(H1(i),'ydata'); z=get(H1(i),'zdata');
          if length(x)>2                               % multiple lines
            warning('Arrow3:Magnitude',MagnitudeWarning), continue
          end
          m=sqrt((x(1)-x(2))^2+(y(1)-y(2))^2+(z(1)-z(2))^2);
          c=LocalInterp(minM,maxM,map,m); set(H1(i),'color',c);
        end
      elseif length(H1)==1
        warning('Arrow3:Magnitude',MagnitudeWarning)
      end
      c=LocalInterp(minM,maxM,map,M);
    end
    set(ax,'ColorOrder',c);                            % update surfaces
    ColorOrder=[];
    if strcmp(get(ax,'tag'),'Arrow3ConePlot')
          H=arrow3(p1,p2,'o' ,w,h,'cone',a);           % update cones
    else, H=arrow3(p1,p2,'o0',w,h,    ip,a);
    end,  H=[H1(:);H(:)];
    set(ax,'nextplot','replace');
end
%-------------------------------------------------------------------------
% SetSurface
function LocalSetSurface(xys,xs,ys,dx,dy,xr,yr,x,y,z,a,c,H,n,m)
    if xys
      x=x*dx+xr(1); y=y*dy+yr(1);
      if xs, x=10.^x; end
      if ys, y=10.^y; end
    end
    cd=zeros(n,m,3); cd(:,:,1)=c(1); cd(:,:,2)=c(2); cd(:,:,3)=c(3);
    set(H,'XData',x,'YData',y,'ZData',z,'CData',cd,'FaceAlpha',a);
end
%-------------------------------------------------------------------------
% ColorTable
function [vc,cn]=LocalColorTable(n,beta)
    vc='kymcrgbadefhijlnpqstuvzw';                     % valid color codes
    %                k               y               m               c
    cn=[0.00,0.00,0.00; 1.00,1.00,0.00; 1.00,0.00,1.00; 0.00,1.00,1.00;
    %                r               g               b               a
        0.635,0.078,0.184; 0.466,0.674,0.188; 0,0.447,0.741; 0.42,0.59,0.24;
    %                d               e               f               h
        0.25,0.25,0.25; 0.00,0.50,0.00; 0.70,0.13,0.13; 1.00,0.41,0.71;
    %                i               j               l               n
        0.29,0.00,0.51; 0.00,0.66,0.42; 0.50,0.50,0.50; 0.50,0.20,0.00;
    %                p               q               s               t
        0.75,0.75,0.00; 1.00,0.50,0.00; 0.00,0.75,0.75; 0.80,0.34,0.00;
    %                u               v               z               w
        0.50,0.00,0.13; 0.75,0.00,0.75; 0.38,0.74,0.99; 1.00,1.00,1.00];
    
    % Named Simulink Colors (zaql)
    % LightBlue = 0.38  0.74  0.99 = aZure
    % DarkGreen = 0.42  0.59  0.24 = Asparagus
    % Orange    = 1.00  0.50  0.00 = kumQuat
    % Gray      = 0.50  0.50  0.50 = Light gray
    %
    % Default ColorOrder Property Colors (bersvpd)
    % Color1    = 0.00  0.00  1.00 = Blue
    % Color2    = 0.00  0.50  0.00 = Evergreen
    % Color3    = 1.00  0.00  0.00 = Red
    % Color4    = 0.00  0.75  0.75 = Sky blue
    % Color5    = 0.75  0.00  0.75 = Violet
    % Color6    = 0.75  0.75  0.00 = Pear
    % Color7    = 0.25  0.25  0.25 = Dark gray
    
    if n, clf reset                                    % plot color table
      name={'blacK','Yellow','Magenta','Cyan',...
        'Red','Green','Blue','Asparagus',...
        'Dark gray','Evergreen','Firebrick','Hot pink',...
        'Indigo','Jade','Light gray','Nutbrown',...
        'Pear','kumQuat','Sky blue','Tawny',...
        'bUrgundy','Violet','aZure','White'};
      c=['yptn';'gjae';'czsb';'hmvi';'qrfu';'wldk'];
      set(gcf,'DefaultAxesXTick',[],'DefaultAxesYTick',[],...
        'DefaultAxesXTickLabel',[],'DefaultAxesYTickLabel',[],...
        'DefaultAxesXLim',[0,0.75],'DefaultAxesYLim',[0,0.75],...
        'DefaultRectangleEdgeColor','none');
      for i=1:24, subplot(4,6,i); box on
        j=find(vc==c(i)); title(name{j});
        dark=LocalBrighten(cn(j,:),-beta);
        light=LocalBrighten(cn(j,:),beta);
        rectangle('Position',[0,0.00,0.75,0.25],'FaceColor',dark);
        rectangle('Position',[0,0.25,0.75,0.25],'FaceColor',cn(j,:));
        rectangle('Position',[0,0.50,0.75,0.25],'FaceColor',light);
        rectangle('Position',[0,0.00,0.75,0.75],'EdgeColor','k');
        if rem(i,6)==1
          set(gca,'YTickLabel',{'dark','normal','light'},...
            'YTick',[0.125,0.375,0.625]);
          if i==19
            text(0,-0.25,['{\bf\itARROW3}  Named Color Table  ',...
                '( \beta = ',num2str(beta),' )']);
          end
        end
      end
    end
end
%-------------------------------------------------------------------------
% ColorMap
function [C,failed]=LocalColorMap(c,vc,cn,beta)
    n=length(c); failed=0; C=zeros(n,3); i=1; j=1;
    while 1
      if ~sum([vc,'_^']==c(i)), failed=1; break, end
      if sum('_^'==c(i))
        if i+1>n, failed=1; break, end
        if ~sum(vc==c(i+1)), failed=1; break, end
        cc=cn(vc==c(i+1),:); gamma=beta;
        if c(i)=='_', gamma=-beta; end
        C(j,:)=LocalBrighten(cc,gamma); i=i+2;
      else, C(j,:)=cn(vc==c(i),:); i=i+1;
      end
      if i>n, break, end, j=j+1;
    end
    if n>j, C(j+1:n,:)=[]; end
end
%-------------------------------------------------------------------------
% Brighten
function C=LocalBrighten(c,beta)
    if sum([c==0,c==1])==3 && sum(c==0)<3 && sum(c==1)<3
      if beta<0
        C=(1+beta)*c;
      else
        C=c;  C(C==0)=beta;
      end
    else
      C=c.^((1-min(1-sqrt(eps),abs(beta)))^sign(beta));
    end
end
%-------------------------------------------------------------------------
% Repmat
function B=LocalRepmat(A,siz)
    if length(A)==1, B(prod(siz))=A; B(:)=A; B=reshape(B,siz);
    else, [m,n]=size(A); mind=(1:m)'; nind=(1:n)';
      mind=mind(:,ones(1,siz(1))); nind=nind(:,ones(1,siz(2)));
      B=A(mind,nind);
    end
end
%-------------------------------------------------------------------------
% Interp
function v=LocalInterp(xmin,xmax,y,u)
    [m,n]=size(y); h=(xmax-xmin)/(m-1); p=length(u); v=zeros(p,n);
    k=min(max(1+floor((u-xmin)/h),1),m-1); s=(u-xmin)/h-k+1;
    for j=1:n, v(:,j)=y(k,j)+s.*(y(k+1,j)-y(k,j)); end
    v(v<0)=0; v(v>1)=1;
end
%-------------------------------------------------------------------------
% Check for supported log scales
function [xs,ys,xys]=LocalLogCheck(ax)
    xs=strcmp(get(ax,'xscale'),'log');
    ys=strcmp(get(ax,'yscale'),'log');
    zs=strcmp(get(ax,'zscale'),'log');
    if zs, error('Z log scale not supported'), end
    xys=xs+ys;
    if xys, azel=get(ax,'view');
      if abs(azel(2))~=90, error('3D log plot not supported'), end
    end
end
%-------------------------------------------------------------------------
% Generate valid value for color, linestyle and linewidth
function [c,ls,lw]=LocalValidateCLSW(s)
    if nargin<1 || isempty(s), c='k'; ls='-'; lw=0.5;
    else
        % identify linestyle
        if contains(s,'--'), ls='--'; s=strrep(s,'--','');
        elseif contains(s,'-.'), ls='-.'; s=strrep(s,'-.','');
        elseif contains(s,'-'), ls='-'; s=strrep(s,'-','');
        elseif contains(s,':'), ls=':'; s=strrep(s,':','');
        elseif contains(s,'*'), ls='*'; s=strrep(s,'*','');
        else, ls='-';
        end
    
        % identify linewidth
        tmp=double(s);
        tmp=find(tmp>45 & tmp<58);
        if ~isempty(tmp)
            if any(s(tmp)=='/'), lw='/'; else, lw=str2double(s(tmp)); end
            s(tmp)='';
        else, lw=0.5;
        end
    
        % identify color
        if ~isempty(s), s=lower(s);
            if length(s)>1, c=s(1:2);
            else, c=s(1); end
        else, c='k';
        end
    end
end
```
