## Supplementary material for "Using pose estimation and 3D rendered models to study leg-mediated self-righting by lanternflies": Dataset S1: FigS2GHIRightingMethodStatistics.m

```
clear all; close all;

show_swarm = 0;

[num,txt,raw] = xlsread('SLF self-righting video summary statistics',5);
SRtime = [num(:,1) num(:,4) num(:,7)]/1000;
attemptime = [num(:,2) num(:,5) num(:,8)]/1000;
numattempts = [num(:,3) num(:,6) num(:,9)];

boxlabels = {'diagonal rotating','lifted rotating','pitching'};

%% plot all data
figure('Color','white','Position',[100,50,400,750]);
t = tiledlayout(3,1);

%% create swarmchart
if show_swarm
    % plot self-righting times
    nexttile(1);
    swarmchart(1:3,SRtime,'k', 'jitter','on', 'jitterAmount',0.35,...
        'MarkerFaceAlpha',0.20,'MarkerEdgeAlpha',0.20); hold on;
    ylabel('time to self-right (s)','Fontsize',13);
    xticks([1 2 3]);
    set(gca,'Xticklabel',boxlabels)

    % plot time/attempt
    nexttile(2);
    swarmchart(1:3,attemptime,'k', 'jitter','on', 'jitterAmount',0.35,...
        'MarkerFaceAlpha',0.20,'MarkerEdgeAlpha',0.20); hold on;
    ylabel('time/attempt (s)','Fontsize',13);
    xticks([1 2 3]);
    set(gca,'Xticklabel',boxlabels)

    % plot number of attempts
    nexttile(3);
    swarmchart(1:3,numattempts,'k', 'jitter','on', 'jitterAmount',0.35,...
        'MarkerFaceAlpha',0.20,'MarkerEdgeAlpha',0.20); hold on;
    ylabel('number of attempts','Fontsize',13);
    xticks([1 2 3]);
    set(gca,'Xticklabel',boxlabels)

end
%% create box-and-whisker plots

% plot self-righting times
nexttile(1);
SRbox = boxplot(SRtime,'Notch','on','Labels',{'' '' ''});
ylabel('time to self-right (s)','Fontsize',13);
set(gca,'Xticklabel',[]) %to just get rid of the numbers but leave the ticks.
ylim([0 18]);
ax = gca; % current axes
ax.FontSize = 11;

nexttile(2);
attemptimebox =boxplot(attemptime,'Notch','on','Labels',{'' '' ''});
set(gca,'Xticklabel',[]) %to just get rid of the numbers but leave the ticks.
ylabel('time/attempt (s)','Fontsize',13);
ylim([0 1.8]);
ax = gca; % current axes
ax.FontSize = 11;

nexttile(3);
numattemptbox =boxplot(numattempts,'Notch','on','Labels',boxlabels);
set(gca,'XTickLabel',boxlabels,'FontSize',10);
% xtickangle(45);         % rotate for readability
ylabel('number of attempts','Fontsize',12);
ylim([0 38]);
ax = gca; % current axes
ax.FontSize = 11;

t.TileSpacing = 'none';
% t.Padding = 'compact';

hold off;

saveas(gcf,'FigS2GHI.png');
%% statistics to compare different methods for righting
figure;
% do statistics for each measure

display('time to self-right statistics');
[p_SRtime,tbl_SRtime,stats_SRtime] = kruskalwallis(SRtime,[],'off');

c = multcompare(stats_SRtime);
tbl = array2table(c,"VariableNames", ...
    ["Group A","Group B","Lower Limit","A-B","Upper Limit","P-value"])

display('time/attempt statistics');
[p_attemptime,tbl_attemptime,stats_attemptime] = kruskalwallis(attemptime,[],'off');
c = multcompare(stats_attemptime);
tbl = array2table(c,"VariableNames", ...
    ["Group A","Group B","Lower Limit","A-B","Upper Limit","P-value"])

display('number of attempts statistics');
[p_numattempts,tbl_numattempts,stats_numattempts] = kruskalwallis(numattempts,[],'off');
c = multcompare(stats_numattempts);
tbl = array2table(c,"VariableNames", ...
    ["Group A","Group B","Lower Limit","A-B","Upper Limit","P-value"])

%% print out summary statistics

ntrials = isnan(SRtime);ntrials = sum(~ntrials,1);

fprintf('mean time to self-right = %f %f %f s\n',mean(SRtime,1,'omitnan'));

fprintf('SD time to self-right = %f %f s\n',std(SRtime(:,1:2),1,'omitnan'));

fprintf('SEM time to self-right = %f %f s\n',std(SRtime(:,1:2),1,'omitnan')./sqrt(ntrials(1:2)));

fprintf('mean time/attempt = %f %f %f s\n',mean(attemptime,1,'omitnan'));

fprintf('SD time/attempt = %f %f s\n',std(attemptime(:,1:2),1,'omitnan'));
fprintf('SEM time/attempt = %f %f s\n',std(attemptime(:,1:2),1,'omitnan')./sqrt(ntrials(1:2)));

fprintf('mean number attempts = %f %f %f s\n',mean(numattempts,1,'omitnan'));

fprintf('SD number attempts = %f %f s\n',std(numattempts(:,1:2),1,'omitnan'));
fprintf('SEM number attempts = %f %f s\n',std(numattempts(:,1:2),1,'omitnan')./sqrt(ntrials(1:2)));
```
