## Supplementary material for "Using pose estimation and 3D rendered models to study leg-mediated self-righting by lanternflies": Dataset S1: Fit3DpointsToCircle_FigS5.m

```
%% compute local circular arc fit to an ordered set of points arranged
% in an approximate arc

clear; close all;

smoothwindow = 25;                 % 25 ms smoothing time

circlefitwindow = 20;

%% load the .mat file with analysis data
[file, pathstr1] = uigetfile('*_anal.mat','Select .mat file with analysis data');
matdatafilename = [pathstr1 file];
load(matdatafilename);

idx1 = startidx; idx2 = idxmax;

prompt = {'first frame to fit' 'last frame to fit'};
dlgtitle = 'user defined inputs';
fieldsize = [1 45];
definput = {num2str(idx1),num2str(idx2)};
answer = inputdlg(prompt,dlgtitle,fieldsize,definput);
[idx1,ansflag]     = str2num(answer{1});
[idx2,ansflag]     = str2num(answer{2});
if idx2 > nframe; idx2 = nframe; end;

X = COM_whole_insect(:,1);  % all values are in mm
Y = COM_whole_insect(:,2);
Z = COM_whole_insect(:,3);

xfit = COM_whole_insect(idx1:idx2,1);  % all values are in mm
yfit = COM_whole_insect(idx1:idx2,2);
zfit = COM_whole_insect(idx1:idx2,3);

%------------------------------------
% fit the orbit trajectory to the arc of a circle & return its XYZ center,
% radius, and its normal vector with its orgin at the center of the circle
%------------------------------------
% first fit xyz values to a plane
% use singular value decomposition to find the plane in best agreement with
% the points provided
% based on Matt J response, Mathworks Answers:
% https://www.mathworks.com/matlabcentral/answers/125251-3d-plane-with-more-than-3-points

center = [mean(xfit) mean(yfit) mean(zfit)];

% subtract off the centroid from all points before SVD
P=[xfit-center(1),yfit-center(2),zfit-center(3)];
meanPlanePts=mean(P);
P=bsxfun(@minus,P,meanPlanePts);
[U,S,V]=svd(P);

% find the normal to the plane:
planenormal=V(:,end);                      % normal to the plane

% Ground plane equation is ax + by + cz + d = 0
a = planenormal(1);
b = planenormal(2);
c = planenormal(3);
d = - dot(planenormal,meanPlanePts);

% now make it a unit vector
planenormal = planenormal/norm(planenormal);

%% rotate the points to be fit so they are in the XY plane
%% compute the rotation angle and axis to rotate data into XY plane
% rotate planenormal = z = [0;0;1]
rotaxis = cross(-planenormal,[0;0;1]); rotaxis= rotaxis'; % format for rotate_3D
rotangle = acos(dot(-planenormal,[0;0;1]));

% make up the array of values to rotate using rotate_3D
Rvec = [xfit';yfit';zfit']; % Rvec should have 3 rows
Rvec = rotate_3D(Rvec,'any',rotangle,rotaxis','radian');
Xrot = Rvec(1,:);Yrot =Rvec(2,:);Zrot =Rvec(3,:);

%% perform circle fit in the XY plane

XY = [Xrot' Yrot'];

% fit the xyz values to arc of a circle
Par = CircleFitByPratt(XY);
% Par = [a b R] is the fitting circle with center (a,b) and radius R
circlecenter = Par(1:2);
% the circle's radius is the distance from the rotation axis to the orbit
circleradius = Par(3);

% option to show data & fit in the xy plane
figure;
plot(XY(:,1),XY(:,2),'k.'); hold on;  % plot data

% compute fit points
theta = atan2(XY(:,2)-circlecenter(2),XY(:,1)-circlecenter(1));
y=circleradius*sin(theta) + circlecenter(2);

axis equal;
normalvector = planenormal*circleradius;
plot(circlecenter(1),circlecenter(2),'o');
plot(XY(:,1),y,'ro')
viscircles(circlecenter(:,1:2),circleradius);      % plot the fit in the XY plane
title('fit circle to data in xy plane');


%% compute the fit residuals
% first get the difference between the Z values and the plane
Zdiff = Zrot -mean(Zrot);
% next, get difference between the y data and the circle fit
Ydiff = y - XY(:,2);
% add in quadrature
fit_resid = sqrt(Zdiff.*Zdiff + Ydiff.*Ydiff);
mean_fit_resid = mean(fit_resid(:));

%% rotate the center point to the original orientation
Rvec = [circlecenter(1);circlecenter(2);mean(Zrot)]; % Rvec should have 3 rows
% rotate in the opposite direction to restore original orientation
Rvec = rotate_3D(Rvec,'any',-rotangle,rotaxis','radian');
circlecenter(1) = Rvec(1);circlecenter(2) =Rvec(2);circlecenter(3) =Rvec(3);

% get the direction of the normal using the right hand rule
vec_start = [xfit(1),yfit(1),zfit(1)]-circlecenter;
vec_end = [xfit(end),yfit(end),zfit(end)]-circlecenter;
norm_direction = cross(vec_start,vec_end);
if dot(norm_direction,planenormal) < 0
    planenormal = -planenormal;
end
%-----------------------------------
figure;
plot3(X,Y,Z,'k.'); hold on;
plot3(X(1),Y(1),Z(1),'g*','MarkerSize',10)
plot3(X(end),Y(end),Z(end),'r+','MarkerSize',10)
plot3(xfit,yfit,zfit,'.','Color',[0.6 0.6 0.6]);

% draw the fitted circle %*% replace with just the fitted arc
drawCircle(circleradius,circlecenter,planenormal,'b',1);

normalvector = planenormal*circleradius ;

% quiver3(mean(X),mean(Y),mean(Z),...
%     normalvector(1),normalvector(2),normalvector(3),'b');

axis equal; 
xlim([1.1*min(X),1.1*max(X)]); ylim([1.1*min(Y),1.1*max(Y)]);
zlim([0 1.1*max(Z)]);
grid on;

xlabel('X (mm)');ylabel('Y (mm)');zlabel('Z (mm)');

% get polar angle from Z axis of plane normal, convert to elevation angle
% (angle of plane normal above XY plane.
elevation_angle = 180/pi*acos(planenormal(3));
if elevation_angle < 90
    elevation_angle = 90 - elevation_angle;
else
    elevation_angle = elevation_angle - 90;
end

legend({'tracks','start','stop','fitted points' 'fit to circle'}); legend boxoff;

normal_label = sprintf('Z_{max} = %2.1f mm, elevation = %2.1f deg',Z(idxmax),elevation_angle);

% radius predicted by the inverted pendulum template model (Fig. 3F)
% Zmax / d = sin(elevation)
d_template = Z(idxmax)/sin(elevation_angle*pi/180);

radius_text = sprintf('radius = %2.1f mm',circleradius);

title({radius_text ; normal_label})

view(90,0);

hold off;


COMtrajfig = fullfile(pathstr1,'COMtrajectory.jpg');
saveas(gcf,COMtrajfig)

%% print statistics
fprintf('mean, max fit residual = %1.3f, %2.3f mm\n',mean_fit_resid,max(fit_resid(:)));
fprintf('max COM Z = %1.1f mm\n',Z(idxmax));
fprintf('fit circle radius = %3.1f mm\n',circleradius);

fprintf('elevation = %3.1f deg\n',elevation_angle);

fprintf('fitted frames %d to %d \n',idx1,idx2);

%% ************************************************************************
function Par = CircleFitByPratt(XY)

%--------------------------------------------------------------------------
%     Copyright (c) 2009, Nikolai Chernov
%     Circle fit by Pratt
%      V. Pratt, "Direct least-squares fitting of algebraic surfaces",
%      Computer Graphics, Vol. 21, pages 145-152 (1987)
%
%     Input:  XY(n,2) is the array of coordinates of n points x(i)=XY(i,1), y(i)=XY(i,2)
%
%     Output: Par = [a b R] is the fitting circle:
%                           center (a,b) and radius R
%
%     Note: this fit does not use built-in matrix functions (except "mean"),
%           so it can be easily programmed in any programming language
%
%--------------------------------------------------------------------------

n = size(XY,1);      % number of data points

centroid = mean(XY);   % the centroid of the data set

%     computing moments (note: all moments will be normed, i.e. divided by n)

Mxx=0; Myy=0; Mxy=0; Mxz=0; Myz=0; Mzz=0;

for i=1:n
    Xi = XY(i,1) - centroid(1);  %  centering data
    Yi = XY(i,2) - centroid(2);  %  centering data
    Zi = Xi*Xi + Yi*Yi;
    Mxy = Mxy + Xi*Yi;
    Mxx = Mxx + Xi*Xi;
    Myy = Myy + Yi*Yi;
    Mxz = Mxz + Xi*Zi;
    Myz = Myz + Yi*Zi;
    Mzz = Mzz + Zi*Zi;
end

Mxx = Mxx/n;
Myy = Myy/n;
Mxy = Mxy/n;
Mxz = Mxz/n;
Myz = Myz/n;
Mzz = Mzz/n;

%    computing the coefficients of the characteristic polynomial

Mz = Mxx + Myy;
Cov_xy = Mxx*Myy - Mxy*Mxy;
Mxz2 = Mxz*Mxz;
Myz2 = Myz*Myz;

A2 = 4*Cov_xy - 3*Mz*Mz - Mzz;
A1 = Mzz*Mz + 4*Cov_xy*Mz - Mxz2 - Myz2 - Mz*Mz*Mz;
A0 = Mxz2*Myy + Myz2*Mxx - Mzz*Cov_xy - 2*Mxz*Myz*Mxy + Mz*Mz*Cov_xy;
A22 = A2 + A2;

epsilon=1e-12;
ynew=1e+20;
IterMax=20;
xnew = 0;

%    Newton's method starting at x=0

for iter=1:IterMax
    yold = ynew;
    ynew = A0 + xnew*(A1 + xnew*(A2 + 4.*xnew*xnew));
    if (abs(ynew)>abs(yold))
        disp('Newton-Pratt goes wrong direction: |ynew| > |yold|');
        xnew = 0;
        break;
    end
    Dy = A1 + xnew*(A22 + 16*xnew*xnew);
    xold = xnew;
    xnew = xold - ynew/Dy;
    if (abs((xnew-xold)/xnew) < epsilon), break, end
    if (iter >= IterMax)
        disp('Newton-Pratt will not converge');
        xnew = 0;
    end
    if (xnew<0.)
        fprintf(1,'Newton-Pratt negative root:  x=%f\n',xnew);
        xnew = 0;
    end
end

%    computing the circle parameters

DET = xnew*xnew - xnew*Mz + Cov_xy;
Center = [Mxz*(Myy-xnew)-Myz*Mxy , Myz*(Mxx-xnew)-Mxz*Mxy]/DET/2;

Par = [Center+centroid , sqrt(Center*Center'+Mz+2*xnew)];

end    %    CircleFitByPratt
%% ************************************************************************
function drawCircle(rad,pos,n,color,linewidth)
%https://demonstrations.wolfram.com/ParametricEquationOfACircleIn3D/
%draws a 3D circle at position pos with radius rad, normal to the
%circle n, and color color.
phi = atan2(n(2),n(1)); %azimuth angle, in [-pi, pi]
theta = atan2(sqrt(n(1)^2 + n(2)^2) ,n(3));% zenith angle, in [0,pi]
t = 0:pi/32:2*pi;
x = pos(1)- rad*( cos(t)*sin(phi) + sin(t)*cos(theta)*cos(phi) );
y = pos(2)+ rad*( cos(t)*cos(phi) - sin(t)*cos(theta)*sin(phi) );
z = pos(3)+ rad*sin(t)*sin(theta);
plot3(x,y,z,color,'LineWidth',linewidth)
end
%% ************************************************************************
function a = anglebetweenthreepoints(A,B,C)
AB = A-B; CB = C-B;
a = atan2(norm(cross(AB,CB)), dot(AB,CB));
end
%% ************************************************************************
%% ************************************************************************
```
