## Supplementary material for "Using pose estimation and 3D rendered models to study leg-mediated self-righting by lanternflies": Dataset S1: PElandscape.m

```
%% plot PE energy landscape figure

close all; clear;

animate_mesh = 0;       % show mesh at each stage of rotations

animation_flag = 0;     % create an animation of the surface plot rotating

plot_PE_colormap = 1;

rotation_using_euler_angles = 1;

%% load the .mat file with analysis data
[file, pathstr1] = uigetfile('*_anal.mat','Select .mat file with analysis data');
matdatafilename = [pathstr1 file];
load(matdatafilename);


%% plot 3d model PE using pitch and roll trajectory from tracked data & analysis
%--------------------------------------------------------------------------


for i = 1:nframe
    % get Euler angles to rotate initial array by
    Euler_angle = quat2eul(quat_array(i,:));
    yaw_array(i)   = Euler_angle(1); 
    pitch_array(i) = Euler_angle(2); 
    roll_array(i)  = Euler_angle(3);

    % get the angle by which the cranial-caudal (body x-axis) is elevated from
    % the XY spatial plane
  
    % compute rotation matrix
    if rotation_using_euler_angles
        rotm = eul2rotm(Euler_angle);
    else  % use quaternions
        rotm = quat2rotm(quat_array(i,:));
    end

    bodyPointsrot = rotm*bodyPoints';      % perform 3D rotation on mesh
    bodyPointsrot = bodyPointsrot';        % restore correct format

    % make an updated mesh
    body_rot = triangulation(bodyConnect,bodyPointsrot);

    % find lowest point along z
    minz = min(bodyPointsrot(:,3));
    % subtract off to get COM at correct height
    % this assumes the body rests on the ground
    % factor of 1e-3 convert distances in mm to meters
    PE_meas(i) = mass_body_only*g*1e-3*minz;

    if animate_mesh
        colormap bone;
        trimesh(body_rot,'FaceAlpha',bodyalpha,'FaceColor','interp',...
            'LineStyle','none');
        xlim([-5 6]);ylim([-5 6]);zlim([-5 6]);
        az = -37.5; el = 5;
        view(az,el); xlabel('X');ylabel('Y');zlabel('Z');
        % create diffuse lighting source at camera position
        camlight('headlight');
        lighting gouraud; material shiny;
        pause(0.01);
    end

    % convert to the convention used in Li et al., 2019 in Fig. 2 PE
    % landscape figure -- get "pitch" (really elevation from XY plane) in
    % radians
    [~,elev,~] = cart2sph(cranial_caudal_axis(i,1),...
        cranial_caudal_axis(i,2),cranial_caudal_axis(i,3));

    if dorsal_ventral_axis(:,3) > 0 % overturned, so elev = 0 to 90 deg
        pitch_array(i) = elev;
    else                            % flipped, do elev = 90 to 180 deg
        pitch_array(i) = pi - elev;
    end
end

PE_meas = abs(PE_meas);

% plot actual trajectory as a solid line
figure('Color','white','Position',[10,100,500,400]);

% resolve large jumps in extrinsic angles & convert to degrees
% the bank and heading angles have range [-pi,pi]; elevation [-pi/2,pi/2]
% by default, unwrap looks for jumps > pi and adds +/- 2pi until the jump
% is < pi; OK to use as is for bank and heading, but need to multipy
% elevation (pitch) by 2

%convert to degrees for plotting & rotating
roll_array = unwrap(roll_array)*180/pi;
pitch_array = pitch_array*180/pi;

% change to Li et al., 2019 conventions for pitch for plotting
pitch_array = 180-pitch_array;

if mean(roll_array) < 0
    roll_array = -roll_array;
end

plot3(pitch_array,roll_array,PE_meas,'k-','LineWidth',1); hold on;
plot3(pitch_array(1),roll_array(1),PE_meas(1),'v','markersize',10,'MarkerEdgeColor','k',...
    'MarkerFaceColor','w');
plot3(pitch_array(end),roll_array(end),PE_meas(end),'^','markersize',10,'MarkerEdgeColor','k',...
    'MarkerFaceColor','w');

%% create surface plot of PE vs a grid of pitch + roll values for the 3D model chosen
% create values for plotting the 3D model PE landscape plot

% save for creating PE values on mesh of pitch and roll values
minpitch = min(0,min(pitch_array));  maxpitch = max(180,max(pitch_array));

minroll = min(-15,min(roll_array));
maxroll = max(195,max(roll_array));

deltapitch = 5; deltaroll = 3;

minroll = floor(minroll/deltaroll)*deltaroll;
minpitch = floor(minpitch/deltapitch)*deltapitch;
maxroll = ceil(maxroll/deltaroll)*deltaroll;
maxpitch = ceil(maxpitch/deltapitch)*deltapitch;

% mesh of pitch and roll values for surface plotting
[pitch,roll] = meshgrid(minpitch:deltapitch:maxpitch,...
    minroll:deltaroll:maxroll);

% first plot the entire PE landscape surface
for j = 1:size(pitch,2)  % number columns == pitch changes
    for i = 1:size(pitch,1) % rows -- roll changes
        % get Euler angles to rotate initial array by
        psi = 0;                % zero yaw (no effect on PE), first rotation
        theta = pitch(i,j);
        phi = roll(i,j);
        Euler_angle = [psi theta phi];
        % convert degrees to radians
        Euler_angle = Euler_angle*pi/180;

        % compute rotation matrix for this set of Euler angles
        rotm = eul2rotm(Euler_angle); 

        bodyPointsrot = rotm*bodyPoints';      % perform 3D rotation on mesh
        bodyPointsrot = bodyPointsrot';        % restore correct format

        % make an updated mesh
        body_rot = triangulation(bodyConnect,bodyPointsrot);

        % find lowest point along z
        minz = min(bodyPointsrot(:,3));
        % subtract off to get COM at correct height
        % this assumes the body rests on the ground
        % factor of 1e-3 convert distances in mm to meters
        PE_mesh(i,j) = mass_body_only*g*1e-3*minz;
    end
end  % end of rotation loop

PE_mesh = abs(PE_mesh);

h1 = surf(pitch,roll,PE_mesh,'FaceAlpha',0.5,'EdgeColor',[0.5 0.5 0.5]);
colormap gray; brighten(0.25);

xlabel('pitch (deg)');ylabel('roll (deg)');zlabel('PE_{body} (J)');
xlim([minpitch maxpitch]); 
% xticks([0 90 180]);
ylim([minroll  maxroll]); yticks([0 90 180]);
zlim([0 1.1*max(PE_mesh(:))]);

legend({'tracked trajectory' 'start' 'end' '' '' ''},...
    'Location','northoutside','Orientation','horizontal');
legend boxoff;

grid on;

if animation_flag           % make an animated version of the surface plot
    %% create videowriter
    videofilename = fullfile(pathstr,[filename(1:23),'_PELand.mp4']);
    rot3DvideoObj=VideoWriter(videofilename,'MPEG-4');
    rot3DvideoObj.FrameRate = 5;
    open(rot3DvideoObj);

    % initialize 3D plot rotation video
    nrotangles = 100;
    az0 = -37.5; azend = 160; el0 = 30; elend = 30;
    az = linspace(az0,azend,nrotangles);
    el = linspace(el0,elend,nrotangles);
    for j = 1:nrotangles
        view(az(j),el(j));
        % animation frame grab (use gcf not just gca)
        animframe = getframe(gcf);
        writeVideo(rot3DvideoObj,animframe);
    end
    close(rot3DvideoObj)
end

%% generate a heatmap viewed from overhead with contour lines
figure('Color','white','Position',[10,100,500,400]);

[M,cp] =contourf(pitch,roll,PE_mesh);  hold on;
colormap parula; brighten(0.3);

% plot the actual pitch-roll trajectory
if plot_PE_colormap         % use the total PE as a color map for the trajectory markers
    scatter(pitch_array,roll_array,15,PE,'filled',"LineWidth",0.1);
    plot(pitch_array,roll_array,'k-','LineWidth',0.25);
else
    plot(pitch_array,roll_array,'k-','LineWidth',1);
end

plot(pitch_array(1),roll_array(1),'v','markersize',8,'MarkerEdgeColor','k',...
    'MarkerFaceColor','w');
plot(pitch_array(end),roll_array(end),'^','markersize',8,'MarkerEdgeColor','k',...
    'MarkerFaceColor','w');

axis equal;
c = colorbar('southoutside');

c.Label.String = 'PE (J)';

xlabel('pitch (deg)');ylabel('roll (deg)');
xlim([minpitch maxpitch]); 
% xticks([0 90 180]);
ylim([minroll  maxroll]); yticks([0 90 180]);

grid on;

legend({'' '' 'tracked trajectory' 'start' 'end' '' '' ''},...
    'Location','eastoutside','Orientation','vertical');
legend boxoff;

%% save figure as image file
FigPElandfilename = fullfile(pathstr1,[filename(1:23),'_PELand.png']);
saveas(gcf,FigPElandfilename);
```
