## Supplementary material for "Using pose estimation and 3D rendered models to study leg-mediated self-righting by lanternflies": Dataset S1: SLF_terrestrial_self_righting_Fig5BSAFTableS4.m

```
%**************************************************************************
close all;  % cleans up all previous windows
clear all;

color_black = [0 0 0];
colors = lines(7);              % make up six colors using the Matlab plot 
%                                 line color defaults
color_red = colors(7,:);        % red
newcolors(1,:) = color_black;   % third instars
newcolors(2,:) = color_red;     % fourth instars
newcolors(3,:) = color_red;
newcolors(4,:) = color_red;

line_styles = {'-','-.','-','-.'};

%**************************************************************************
% selpath = uigetdir; cd(selpath); % change to directory of interest
%--------------------------------------------------------------------------
%% plot success percentage for self-righting
% get data from sheet 2
[num,txt,raw] = xlsread('SLF self-righting video summary statistics',2);
percent_success = num(1:4,2)./num(1:4,1);              % column 1 = % success
percent_success(:,2) = 1-percent_success;  % column 2 = % failure
percent_success = 100*percent_success;

substrate = txt(2:5,1);
instar = txt(2:5,2);

%--------------------------------------------------------------------------
% Bar graph of success percentage for self-righting for Fig. 4B
figure('Color','white','Position',[10 10 700 700]);
subplot(1,2,1);
success_plot = barh(percent_success,'stacked');
hold on;

% set colors for the bars
barcolors(1,:) = [113,143,80]/255;    % green -- works with color variants
barcolors(2,:) = [241,120,83]/255;    % orange-red -- works with color variants
colororder(barcolors);

yticklabels(substrate);
xlim([0 120] );
xticks([0 50 100]);

xlabel('Percent','Fontsize',14);
ylabel('Substrate','Fontsize',14)
legend('Succeeded','Failed','');
legend 'boxoff'; legend('Location','northoutside','Orientation','vertical');
legend('Fontsize',12);

xtips1 = success_plot(2).YEndPoints + 1.3;
ytips1 = success_plot(2).XEndPoints;
for i = 1:4
    bartip_label{i} = instar{i};
end
text(xtips1,ytips1,bartip_label,'VerticalAlignment','middle','Fontsize',11)

set(gca,'FontSize',11);

hold off;
hsp1 = get(gca, 'Position');            % Get 'Position' for (2,1,1)
% save plot as a file
saveas(gcf,'Fig4B.png');

%**************************************************************************
% plot summary statistics for self-righting time/attempts, number attempts
% get data from sheet 1
[num,txt,raw] = xlsread('SLF self-righting video summary statistics',1);
%--------------------------------------------------------------------------
% parse substrate type
substrate = txt(2:end,1);
% parse datasets using substrate type and instar (num(:,1)
for i = 1:numel(substrate)
    if num(i,1) == 0 
        exp_condition(i) = 1;
    else
        if strcmp(substrate{i},'posterboard')
            exp_condition(i) = 2;
        elseif strcmp(substrate{i},'glass') 
            exp_condition(i) = 3;
        else
            exp_condition(i) = 4;
        end
    end
end

% get indices for each substrate/instar condition
idx_3rdinstar = find(exp_condition == 1);
idx_4thinstar_posterboard = find(exp_condition == 2);
idx_4thinstar_glass = find(exp_condition == 3);
idx_4thinstar_sandpaper = find(exp_condition == 4);

% make up labels
instar_substrate{1} = '3rd instar, posterboard';
instar_substrate{2} = '4th instar, posterboard';
instar_substrate{3} = '4th instar, glass';
instar_substrate{4} = '4th instar, sandpaper';
%-----------------------
% parse numerical data
trial_num = num(:,2);

mean_attempt_time = num(:,8)/1000;median_attempt_time = num(:,12)/1000;
CI_attempt_time = 1.96*num(:,10)/1000;

% because these data are ordinal, we should use medians and ignore the CI
% based on standard errors
median_num_attempts = num(:,11);

mean_search_time = num(:,4)/1000;median_search_time = num(:,6)/1000;
CI_search_time = 1.96*num(:,5)/1000;

%**************************************************************************
% create plots of how these depend on trial number for Fig. 5CDE
figure('Color','white','Position',[100,50,450,900]);
t = tiledlayout(3,1);
legend_labels = {'3rd instar, posterboard','linear fit',...
     '4th instar, posterboard','4th instar, glass','4th instar, sandpaper'};
%--------------------------
% % search time
nexttile;
errorbar(trial_num(idx_3rdinstar) - 0.1,mean_search_time(idx_3rdinstar),...
    CI_search_time(idx_3rdinstar),'o-','MarkerSize',9,...
    'MarkerEdgeColor',newcolors(1,:),'MarkerFaceColor',newcolors(1,:)); hold on;
errorbar(trial_num(idx_4thinstar_posterboard) - 0.05,mean_search_time(idx_4thinstar_posterboard) - 0.05,...
    CI_search_time(idx_4thinstar_posterboard) - 0.05,'--s','MarkerSize',9,...
    'MarkerEdgeColor',newcolors(2,:),'MarkerFaceColor',newcolors(2,:)); hold on;
errorbar(trial_num(idx_4thinstar_glass) + 0.05,mean_search_time(idx_4thinstar_glass),...
    CI_search_time(idx_4thinstar_glass),'-.d','MarkerSize',9,...
    'MarkerEdgeColor',newcolors(3,:),'MarkerFaceColor',newcolors(3,:)); hold on;
errorbar(trial_num(idx_4thinstar_sandpaper) + 0.1,mean_search_time(idx_4thinstar_sandpaper),...
    CI_search_time(idx_4thinstar_sandpaper),':^','MarkerSize',9,...
    'MarkerEdgeColor',newcolors(4,:),'MarkerFaceColor',newcolors(4,:)); hold on;

xticks(''); xlim([0.5,5.5]);ylim([0 4.5]);

ylabel('search time (s)','Fontsize',14);

set(gca,'FontSize',11);

hold off;

%-----------------------
% time/attempt

nexttile;
colororder(newcolors);

errorbar(trial_num(idx_3rdinstar) - 0.1,mean_attempt_time(idx_3rdinstar),...
    CI_attempt_time(idx_3rdinstar),'-o','MarkerSize',9,...
    'MarkerEdgeColor',newcolors(1,:),'MarkerFaceColor',newcolors(1,:)); hold on;
errorbar(trial_num(idx_4thinstar_posterboard) - 0.05,mean_attempt_time(idx_4thinstar_posterboard) - 0.05,...
    CI_attempt_time(idx_4thinstar_posterboard) - 0.05,'--s','MarkerSize',9,...
    'MarkerEdgeColor',newcolors(2,:),'MarkerFaceColor',newcolors(2,:)); hold on;
errorbar(trial_num(idx_4thinstar_glass) + 0.05,mean_attempt_time(idx_4thinstar_glass),...
    CI_attempt_time(idx_4thinstar_glass),'-.d','MarkerSize',9,...
    'MarkerEdgeColor',newcolors(3,:),'MarkerFaceColor',newcolors(3,:)); hold on;
errorbar(trial_num(idx_4thinstar_sandpaper) + 0.1,mean_attempt_time(idx_4thinstar_sandpaper),...
    CI_attempt_time(idx_4thinstar_sandpaper),':^','MarkerSize',9,...
    'MarkerEdgeColor',newcolors(4,:),'MarkerFaceColor',newcolors(4,:)); hold on;

xticks(''); xlim([0.5,5.5]);ylim([0 0.6]);

ylabel('time/attempt (s)','Fontsize',14);

set(gca,'FontSize',11);

hold off;

%% add linear regression vs. trial number

%-----------------------
% number attempts

nexttile;
colororder(newcolors);

plot(trial_num(idx_3rdinstar) - 0.1,median_num_attempts(idx_3rdinstar),...
    '-o','MarkerSize',9,...
    'MarkerEdgeColor',newcolors(1,:),'MarkerFaceColor',newcolors(1,:)); hold on;
plot(trial_num(idx_4thinstar_posterboard) - 0.05,median_num_attempts(idx_4thinstar_posterboard) - 0.05,...
    '--s','MarkerSize',9,...
    'MarkerEdgeColor',newcolors(2,:),'MarkerFaceColor',newcolors(2,:)); hold on;
plot(trial_num(idx_4thinstar_glass) + 0.05,median_num_attempts(idx_4thinstar_glass),...
    '-.d','MarkerSize',9,...
    'MarkerEdgeColor',newcolors(3,:),'MarkerFaceColor',newcolors(3,:)); hold on;
plot(trial_num(idx_4thinstar_sandpaper) + 0.1,median_num_attempts(idx_4thinstar_sandpaper),...
    ':^','MarkerSize',9,...
    'MarkerEdgeColor',newcolors(4,:),'MarkerFaceColor',newcolors(4,:)); hold on;
xticks([1 2 3 4 5]); xlim([0.5,5.5]);

ylabel('number of attempts','Fontsize',14);
ylim([0 max(median_num_attempts)+1]);

set(gca,'FontSize',11);

t.TileSpacing = 'tight';
t.Padding = 'tight';
xlabel(t,'trial number','Fontsize',14);

% global legend tiled layout plot
% https://www.mathworks.com/matlabcentral/answers/542096-adding-a-global-legend-to-a-tiledlayout#answer_502549
lg  = legend(legend_labels);legend 'boxoff';
lg.FontSize = 12; set(lg,'Box','off');
lg.Layout.Tile = 'North'; % <-- place legend north of tiles

hold off;

saveas(gcf,'FigS2ABC.png');

%% perform statistical analysis for Table S2

% save all linear regression results & statistics in Table S2
% outarray_numattempts = [header; outarray];

%**************************************************************************
% plot the data for number attempts, time to self-right, time/attempts as
% box-and-whisker plots for Fig. 5F

% On each box, the central mark indicates the median, and the bottom and
% top edges of the box indicate the 25th and 75th percentiles,
% respectively. The whiskers extend to the most extreme data points not
% considered outliers, and the outliers are plotted individually using the
% '+' marker symbol.

[num,txt,raw] = xlsread('SLF self-righting video summary statistics',3);
SRtime = [num(:,1) num(:,4) num(:,7) num(:,10)];
attemptime = [num(:,2) num(:,5) num(:,8) num(:,11)];
numattempts = [num(:,3) num(:,6) num(:,9) num(:,12)];

boxlabels = {'3rd, posterboard','4th, posterboard','4th, glass','4th, sandpaper'};

figure('Color','white','Position',[100,50,300,700]);
t = tiledlayout(3,1);

% plot self-righting times
nexttile;
SRbox = boxplot(SRtime/1000,'Notch','on','Labels',{'' '' '' ''},'symbol', '');
ylabel('time to self-right (s)','Fontsize',13);
set(gca,'Xticklabel',[]) %to just get rid of the numbers but leave the ticks.
ylim([0 2.8]);
ax = gca; % current axes
ax.FontSize = 11;

nexttile;
attemptimebox =boxplot(attemptime/1000,'Notch','on','Labels',boxlabels,'symbol', '');
set(gca,'Xticklabel',[]) %to just get rid of the numbers but leave the ticks.
ylabel('time/attempt (s)','Fontsize',13);
ylim([0 0.9]);
ax = gca; % current axes
ax.FontSize = 11;

nexttile;
numattemptbox =boxplot(numattempts,'Notch','on','Labels',boxlabels,'symbol', '');
set(gca,'XTickLabel',boxlabels,'FontSize',10);
xtickangle(45);         % rotate for readability
ylabel('number of attempts','Fontsize',12);
ylim([0 13]);
ax = gca; % current axes
ax.FontSize = 11;

t.TileSpacing = 'none';
t.Padding = 'compact';

hold off;

saveas(gcf,'FigS2DEF.png');

%% statistics for Table S4
figure;
% do statistics for each measure

display('time to self-right statistics');
[p_SRtime,tbl_SRtime,stats_SRtime] = kruskalwallis(SRtime,[],'off');

c = multcompare(stats_SRtime);
tbl = array2table(c,"VariableNames", ...
    ["Group A","Group B","Lower Limit","A-B","Upper Limit","P-value"])

display('attempt/time statistics');
[p_attemptime,tbl_attemptime,stats_attemptime] = kruskalwallis(attemptime,[],'off');
c = multcompare(stats_attemptime);
tbl = array2table(c,"VariableNames", ...
    ["Group A","Group B","Lower Limit","A-B","Upper Limit","P-value"])

display('number of attempts statistics');
[p_numattempts,tbl_numattempts,stats_numattempts] = kruskalwallis(numattempts,[],'off');
c = multcompare(stats_numattempts);
tbl = array2table(c,"VariableNames", ...
    ["Group A","Group B","Lower Limit","A-B","Upper Limit","P-value"])
%**************************************************************************

%% ************************************************************************
function [output_data] = lmfit_and_plot(x,y,SE,label_text,...
    marker_type,marker_size,marker_color,line_type)

display(strcat('Fit ',label_text));

if max(SE) > 0
    errorbar(x,y,SE, marker_type,...
        'MarkerSize',marker_size,'Color',marker_color,'LineWidth',1.5);
    hold on;
    weight = 1./(SE.*SE);
    mdl = fitlm(x,y,'Weights',weight);
else
    plot(x,y,marker_type,'MarkerSize',marker_size,...
        'Color',marker_color,'LineWidth',1.5);
    hold on;
    mdl = fitlm(x,y);
end

% get fit statistics

y_CI = coefCI(mdl);

dof = mdl.DFE;
Rsquared = mdl.Rsquared.Ordinary;

meanfitcoeff=mdl.Coefficients.Estimate;  % the fit coefficients
fitparamSE = mdl.Coefficients.SE;
tStat = mdl.Coefficients.tStat;
fitparamP = mdl.Coefficients.pValue;

Pval=mdl.ModelFitVsNullModel.Pvalue;% P value
Fstat = mdl.ModelFitVsNullModel.Fstat;

% plot fitted line if fitparamP(2) < 0.05
if Pval < 0.05
    yfit = meanfitcoeff(1) + meanfitcoeff(2)*x;
    plot(x,yfit,line_type,'Color',marker_color,'LineWidth',1);
end

fprintf('R^2 = %f, Fstat = %f, P = %f\n',Rsquared,Fstat,Pval);
fprintf('intercept = %f , CI = %f %f\n',meanfitcoeff(1),y_CI(1,1),...
    y_CI(1,2));
fprintf('slope = %f , CI = %f %f\n',meanfitcoeff(2),y_CI(2,1),...
    y_CI(2,2));

fprintf('\n *************************************************\n');

% create output array
output_data = {label_text, meanfitcoeff(1), fitparamSE(1),...
    y_CI(1,1),y_CI(1,2),tStat(1),fitparamP(1),meanfitcoeff(2),fitparamSE(2),...
    y_CI(2,1),y_CI(2,2),tStat(2),fitparamP(2),Rsquared,Fstat,Pval,dof};

end
```
