## Supplementary material for "Using pose estimation and 3D rendered models to study leg-mediated self-righting by lanternflies": Dataset S1: SLF_terrestrial_self_righting_TableS2S3.m

```
%**************************************************************************
close all;  % cleans up all previous windows
clear;

line_style = {'-','-.','-','-.'};
marker_style = {'o' '*' 's' '^' 'd' '>'};
newcolors = lines(4);
header = {'source' 'int' 'SE' 'CI' 'CI' 'tstat' 'p'...
    'slope' 'SE' 'CI' 'CI'  'tstat' 'p' 'R2' 'Fstat' 'Pval' 'dof'};
header_chisqu = {'Group A' 'Group B' 'h' 'p-value' 'chi-squared' 'dof'};

% successful vs failed self-righting for Supplementary Tables
[num,txt,raw] = xlsread('SLF self-righting video summary statistics',4);

trial_num = num(:,1);
condition = txt(2:end,2);
condition_list = unique(condition);

% pairs of conditions to test using chi-square
GroupAB = nchoosek(1:numel(condition_list),uint16(2));
[numABpairs,~] = size(GroupAB);

succeeded = num(:,3); failed = num(:,4);
percent_success = succeeded/(succeeded+failed);

% Table S2:  The linear regression statistics 
% for each condition, see if it's correlated with trials number using linear regression
outarray = [];
for i = 1:numel(condition_list)
    idx = find(contains(condition,condition_list{i}));
    x = trial_num(idx); y = percent_success(idx); SE = 0;
    [output_data] = lmfit_and_plot(x,y,SE,condition_list(i),...
        marker_style{i},10,newcolors(i,:),line_style{i});
    outarray = [outarray;output_data];
end
xlabel('trial number');ylabel('self-righting percent success');
legend(condition_list);
hold off;
% save all linear regression results & statistics
outarray_percent_success = [header; outarray];

% because there is no dependence on trial number, 
% now organize by condition, pooling all trials 
percent_success_by_condition = [];
for i = 1:numel(condition_list)
    idx = find(contains(condition,condition_list{i}));
    succeeded_condition(i) = sum(succeeded(idx));
    failed_condition(i) = sum(failed(idx));
end

total_condition = succeeded_condition + failed_condition;

for i = 1:numABpairs
    [h_success(i),p_success(i),chi(i),df(i)] = ...
        prop_test(succeeded_condition(GroupAB(i,:)),total_condition(GroupAB(i,:)), true);
    outarray_success_chisq(i,:) = ...
        {condition_list{GroupAB(i,1)},condition_list{GroupAB(i,2)},...
        h_success(i),p_success(i),chi(i),df(i)};
end

% Data for Table S3
outarray_success_chisq = [header_chisqu; outarray_success_chisq];


% end of main function
%% ************************************************************************
function [output_data] = lmfit_and_plot(x,y,SE,label_text,...
    marker_type,marker_size,marker_color,line_type)

display(strcat('Fit ',label_text));

if max(SE) > 0
    errorbar(x,y,SE, marker_type,...
        'MarkerSize',marker_size,'Color',marker_color,'LineWidth',1.5);
    hold on;
    weight = 1./(SE.*SE);
    mdl = fitlm(x,y,'Weights',weight);
else
    plot(x,y,marker_type,'MarkerSize',marker_size,...
        'Color',marker_color,'LineWidth',1.5);
    hold on;
    mdl = fitlm(x,y);
end

% get fit statistics

y_CI = coefCI(mdl);

dof = mdl.DFE;
Rsquared = mdl.Rsquared.Ordinary;

meanfitcoeff=mdl.Coefficients.Estimate;  % the fit coefficients
fitparamSE = mdl.Coefficients.SE;
tStat = mdl.Coefficients.tStat;
fitparamP = mdl.Coefficients.pValue;

Pval=mdl.ModelFitVsNullModel.Pvalue;% P value
Fstat = mdl.ModelFitVsNullModel.Fstat;

% plot fitted line if fitparamP(2) < 0.05
if Pval < 0.05
    yfit = meanfitcoeff(1) + meanfitcoeff(2)*x;
    plot(x,yfit,line_type,'Color',marker_color,'LineWidth',1);
end

fprintf('R^2 = %f, Fstat = %f, P = %f\n',Rsquared,Fstat,Pval);
fprintf('intercept = %f , CI = %f %f\n',meanfitcoeff(1),y_CI(1,1),...
    y_CI(1,2));
fprintf('slope = %f , CI = %f %f\n',meanfitcoeff(2),y_CI(2,1),...
    y_CI(2,2));

fprintf('\n *************************************************\n');

% create output array
output_data = {label_text, meanfitcoeff(1), fitparamSE(1),...
    y_CI(1,1),y_CI(1,2),tStat(1),fitparamP(1),meanfitcoeff(2),fitparamSE(2),...
    y_CI(2,1),y_CI(2,2),tStat(2),fitparamP(2),Rsquared,Fstat,Pval,dof};

end

%% ************************************************************************
% laurie (2023). Compare two proportions (Chi-square) 
% (https://www.mathworks.com/matlabcentral/fileexchange/45966-compare-two-proportions-chi-square)
%  MATLAB Central File Exchange. Retrieved January 27, 2023. 

function [h,p, chi2stat,df] = prop_test(X , N, correct)

% [h,p, chi2stat,df] = prop_test(X , N, correct)
% 
% A simple Chi-square test to compare two proportions
% It is a 2 sided test with alpha=0.05
%
% Input:
% X = vector with number of success for each sample (e.g. [20 22])
% N = vector of total counts for each sample (e.g. [48 29])
% correct = true/false : Yates continuity correction for small samples?
%
% Output:
% h = hypothesis (H1/H0)
% p = p value
% chi2stat= Chi-square value
% df = degrees of freedom (always equal to 1: 2 samples)
%
% Needs chi2cdf from the Statistics toolbox
% Inspired by prop.test() in "R" but much more basic
%
% Example: [h,p,chi]=prop_test([20 22],[48 29], true)
% The above example tests if 20/48 differs from 22/29 using Yate's correction

if (length(X)~= 2)||(length(X)~=length(N))
    disp('Error: bad vector length')
elseif (X(1)>N(1))|| (X(2)>N(2))
    disp('Error: bad counts (X>N)')
else  
    df=1; % 2 samples
    
    % Observed data
    n1 = X(1);
    n2 = X(2);
    N1 = N(1);
    N2 = N(2);
    
    % Pooled estimate of proportion
    p0 = (n1+n2) / (N1+N2);
    
    % Expected counts under H0 (null hypothesis)
    n10 = N1 * p0;
    n20 = N2 * p0;
    observed = [n1 N1-n1 n2 N2-n2];
    expected = [n10 N1-n10 n20 N2-n20];
    
    if correct == false
        % Standard Chi-square test
        chi2stat = sum((observed-expected).^2 ./ expected);
        p = 1 - chi2cdf(chi2stat,1);
    else
        % Yates continuity correction        
        chi2stat = sum((abs(observed - expected) - 0.5).^2 ./ expected);
        p = 1 - chi2cdf(chi2stat,1);
    end
    
    h=0;
    if p<0.05
        h=1;
    end
end
end
```
