## Supplementary material for "Using pose estimation and 3D rendered models to study leg-mediated self-righting by lanternflies": Dataset S1: RigidBodyParams.m

```
function [RBP,TR]=RigidBodyParams(TR)
% Compute exact rigid body parameters (listed in the OUTPUT below) of a 
% solid object represented by a CLOSED triangular surface mesh. For 
% details, see the accompanying pdf titled "Rigid-body Parameters Using 
% Divergence Theorem".
%
% INPUT:
%   - TR    : triangular surface mesh specified in one of the following
%             formats:
%               a. 'TriRep' object (may be obsolete depending on your 
%                  version of Matlab)
%               b. 'triangulation' object
%               c. Structure with the exact same fields as the one returned 
%                   by the 'isosurface' function; that is  
%                       TR.faces    : M-by-3 list of faces
%                       TR.vertices : N-by-3 list of vertex co-ordinates
%               d. Cell such TR={Tri X} where Tri is a M-by-3 list of faces
%                  and X is a N-by-3 list of vertex co-ordinates
%
% OUTPUT:
%   - RBP   : structure containing all relevant rigid body parameters
%               RBP.volume          : total volume enclosed by the mesh.
%                                     Total mass of the object can be 
%                                     computed by multiplying total volume 
%                                     with density. Note that negative
%                                     volume indicates that mesh vertices
%                                     are pointing inwards.
%               RBP.centroid        : 1-by-3 vector specifying the
%                                     co-ordinates of the object's centroid. 
%               RBP.inertia_tensor  : 3-by-3 symmetric matrix representing 
%                                     the object's inertia tensor.
%               RBP.PAI             : 3-by-3 array containing principal 
%                                     axes of inertia, along columns.
%               RBP.eigs            : 1-by-3 vector containing magnitudes 
%                                     of the principal axes of inertia, so
%                                     that RBP.eigs(i) corresponds to
%                                     RBP.PAI(:,i).
%               RBP.moments         : 1-by-10 vector of moments used to 
%                                     compute the volume, centroid, and
%                                     inertia parameters of the input
%                                     object. Suppose that M=RBP.moments,
%                                     then M=[m000 m100 m010 m001 m110 ...
%                                     m101 m011 m200 m020 m002]. 
%           
% AUTHOR: Anton Semechko
%


% Check input format and retrieve face and vertex lists
TR=CheckFormat(TR);
Tri=TR.faces;
X=TR.vertices;

% Area weighted face normals (magnitude = 2 x triangle area)
X1=X(Tri(:,1),:);
X2=X(Tri(:,2),:);
X3=X(Tri(:,3),:);
FN=cross(X2-X1,X3-X1,2);

% Zeroth order moment (same as volume enclosed by the mesh) ---------------
C=(X1+X2+X3)/3;
m000=sum(sum(FN.*C,2))/6;
if m000<eps
    warning('Mesh has negative volume, indicating that face normal orientations are reversed (i.e., pointing into the surface).')
end

% First order moments (together they specify the centroid of the region 
% enclosed by the mesh) ---------------------------------------------------
x1=X1(:,1); y1=X1(:,2); z1=X1(:,3);
x2=X2(:,1); y2=X2(:,2); z2=X2(:,3);
x3=X3(:,1); y3=X3(:,2); z3=X3(:,3);

x_2=((x1+x2).*(x2+x3) + x1.^2 + x3.^2)/12;
y_2=((y1+y2).*(y2+y3) + y1.^2 + y3.^2)/12;
z_2=((z1+z2).*(z2+z3) + z1.^2 + z3.^2)/12;

xy=((x1+x2+x3).*(y1+y2+y3) + x1.*y1 + x2.*y2 + x3.*y3)/24;
xz=((x1+x2+x3).*(z1+z2+z3) + x1.*z1 + x2.*z2 + x3.*z3)/24;
yz=((y1+y2+y3).*(z1+z2+z3) + y1.*z1 + y2.*z2 + y3.*z3)/24;

m100=sum(sum(FN.*[x_2 2*xy 2*xz],2))/6;
m010=sum(sum(FN.*[2*xy y_2 2*yz],2))/6;
m001=sum(sum(FN.*[2*xz 2*yz z_2],2))/6;

% Second order moments (used to determine elements of the inertia tensor)
% -------------------------------------------------------------------------
x_3=((x1+x2+x3).*(x1.^2+x2.^2+x3.^2) + x1.*x2.*x3)/20; 
y_3=((y1+y2+y3).*(y1.^2+y2.^2+y3.^2) + y1.*y2.*y3)/20; 
z_3=((z1+z2+z3).*(z1.^2+z2.^2+z3.^2) + z1.*z2.*z3)/20; 

x_2y=((3*y1+y2+y3).*x1.^2 + (y1+3*y2+y3).*x2.^2 + (y1+y2+3*y3).*x3.^2 + (2*y1+2*y2+y3).*x1.*x2 + (2*y1+y2+2*y3).*x1.*x3 + (y1+2*y2+2*y3).*x2.*x3)/60;
x_2z=((3*z1+z2+z3).*x1.^2 + (z1+3*z2+z3).*x2.^2 + (z1+z2+3*z3).*x3.^2 + (2*z1+2*z2+z3).*x1.*x2 + (2*z1+z2+2*z3).*x1.*x3 + (z1+2*z2+2*z3).*x2.*x3)/60;

y_2x=((3*x1+x2+x3).*y1.^2 + (x1+3*x2+x3).*y2.^2 + (x1+x2+3*x3).*y3.^2 + (2*x1+2*x2+x3).*y1.*y2 + (2*x1+x2+2*x3).*y1.*y3 + (x1+2*x2+2*x3).*y2.*y3)/60;
y_2z=((3*z1+z2+z3).*y1.^2 + (z1+3*z2+z3).*y2.^2 + (z1+z2+3*z3).*y3.^2 + (2*z1+2*z2+z3).*y1.*y2 + (2*z1+z2+2*z3).*y1.*y3 + (z1+2*z2+2*z3).*y2.*y3)/60;

z_2y=((3*y1+y2+y3).*z1.^2 + (y1+3*y2+y3).*z2.^2 + (y1+y2+3*y3).*z3.^2 + (2*y1+2*y2+y3).*z1.*z2 + (2*y1+y2+2*y3).*z1.*z3 + (y1+2*y2+2*y3).*z2.*z3)/60;
z_2x=((3*x1+x2+x3).*z1.^2 + (x1+3*x2+x3).*z2.^2 + (x1+x2+3*x3).*z3.^2 + (2*x1+2*x2+x3).*z1.*z2 + (2*x1+x2+2*x3).*z1.*z3 + (x1+2*x2+2*x3).*z2.*z3)/60;

xyz=((x1+x2+x3).*(y1+y2+y3).*(z1+z2+z3) - (y2.*z3+y3.*z2-4*y1.*z1).*x1/2 -(y1.*z3+y3.*z1-4*y2.*z2).*x2/2 - (y1.*z2+y2.*z1-4*y3.*z3).*x3/2)/60;

m110=sum(sum(FN.*[x_2y y_2x 2*xyz],2))/6;
m101=sum(sum(FN.*[x_2z 2*xyz z_2x],2))/6;
m011=sum(sum(FN.*[2*xyz y_2z z_2y],2))/6;

m200=sum(sum(FN.*[x_3 3*x_2y 3*x_2z],2))/9;
m020=sum(sum(FN.*[3*y_2x y_3 3*y_2z],2))/9;
m002=sum(sum(FN.*[3*z_2x 3*z_2y z_3],2))/9;

% Inertia tensor ----------------------------------------------------------
Ixx=m020+m002-(m010^2+m001^2)/m000;
Iyy=m200+m002-(m100^2+m001^2)/m000;
Izz=m200+m020-(m100^2+m010^2)/m000;
Ixy=m110-m100*m010/m000;
Ixz=m101-m100*m001/m000;
Iyz=m011-m010*m001/m000;

I=[Ixx -Ixy -Ixz; -Ixy Iyy -Iyz; -Ixz -Iyz Izz];

% Output ------------------------------------------------------------------
[V,D]=svd(I);

RBP.volume=m000;
RBP.centroid=[m100 m010 m001]/m000;
RBP.inertia_tensor=I;
RBP.PAI=V;
RBP.eigs=diag(D)';
RBP.moments=[m000 m100 m010 m001 m110 m101 m011 m200 m020 m002];


function TR=CheckFormat(TR)
% Verify that input format is correct and retrieve face and vertex lists

flag=false;
switch class(TR)
    case 'TriRep'
        X=TR.X;
        Tri=TR.Triangulation;
    case 'triangulation'
        X=TR.Points;
        Tri=TR.ConnectivityList;
    case 'struct'
        if isfield(TR,'faces') && isfield(TR,'vertices')
            X=TR.vertices;
            Tri=TR.faces;
        else
            error('Invalid input format. Structure is missing required fields.')
        end
        flag=true;
    case 'cell'
        if numel(TR)==2
            X=TR{2};
            Tri=TR{1};
        else
            error('Invalid mesh format')
        end 
        flag=true;
    otherwise
        error('Unrecognized mesh format')
end

% Verify that the face and vertex lists are correct
if flag
    if ~ismatrix(Tri) || isempty(Tri) || size(Tri,2)~=3
        error('Face list must be a M-by-3 array')
    end
    if ~ismatrix(X) || isempty(X) || size(X,2)~=3
        error('Vertex list must be a N-by-3 array')
    end
    if ~isequal(Tri,round(Tri)) || min(Tri(:))<1
        error('Specified face list is invalid')
    end
end

if max(Tri(:))~=size(X,1) || numel(unique(Tri(:)))~=size(X,1)
    warning('Some vertices are not referenced by the triangulation')
end

if ~all(isfinite(X(:)))
    error('Vertex list contains one or more undefined entries')
end
    
% Verify that the mesh is closed and manifold
if flag
    try
        TR=triangulation(Tri,X);
    catch
        TR=TriRep(Tri,X);  %#ok<*DTRIREP>
    end
end

EA=edgeAttachments(TR,edges(TR));
Nef=cellfun(@length,EA);
if any(Nef~=2)
    error('Input mesh either contains boundaries or is non-manifold')
end

TR=struct();
TR.faces=Tri;
TR.vertices=X;
```
