## Supplementary material for "Using pose estimation and 3D rendered models to study leg-mediated self-righting by lanternflies": Dataset S1: rotate_3D.m

```
function [R, Rm] = rotate_3D(V, mode, theta, u, angle_unit)
%% rotate_3D : function to compute the rotation of a vector or an array of vectors in 2D or 3D space.
%
% Author && support : nicolas.douillet (at) free.fr, 2017-2020.
%
%
% Syntax
% R = rotate_3D(V, mode, theta);
% R = rotate_3D(V, mode, theta, u);
% R = rotate_3D(V, mode, theta, u, angle_unit);
% [R,Rm] = rotate_3D(V, mode, theta, u, angle_unit);
%
%
% Description
% R = rotate_3D(V, mode, theta) computes the vector R, which results
% from the rotation of V vector around one of the the basis vectors, which
% is choosen in the mode : 'x', 'y', or 'z'.
%
% R = rotate_3D(V, mode, theta, u) computes the vector R, which results
% from the rotation of V vector around u vector and of theta angle in radian.
%
% R = rotate_3D(V, mode, theta, u, angle_unit) uses angle_unit for theta
% unit (radian or degree).
%
% [R,Rm] = rotate_3D(V, mode, theta, u, angle_unit) also returns the
% rotation matrix.
%
% Important NB : in 2D -(xOy) plan- mandatory rotation axis is 'z'. It will
% be set as so by default if input is different. Also in 2D, in case u is missing it
% is automatically set to the origin [0,0]' by default.
%
%
% Input arguments
%
%       [ -Vx- ]
% - V = [ -Vy- ], real (array of) vector(s) double, the vector(s) to rotate. Size(V) = [3,vector_nb]. 
%       [ -Vz- ]
%
% - mode : character string in the set {'x', 'X', 'y', 'Y', 'z', 'Z', 'any', 'ANY'}. 
%
% - theta : real scalar double, the rotation angle in radian.
%
%         [ux]
% - u : = [uy], real column vector double, the rotation axis.
%         [uz]
%
% - angle_unit : character string in the set {'radian','degree','RADIAN,'DEGREE'}. Case insensitive.
%
%
% Output arguments
%
%       [ -Rx- ]
% - R = [ -Ry- ], real (array of) vector(s) double, the resulting rotated vector(s). Size(R) = [3,vector_nb].
%       [ -Rz- ]
%
% - Rm : real matrix double, the resulting rotation matrix, size(Rm) = [3,3].
%
%
% 3D Example #1
%
% 2*pi/3 rotation of i vector around [1 1 1]' is basis vector j.
% V = [1 0 0]';
% mode = 'any';
% u = ones(3,1);
% theta = 2*pi/3;
% R = rotate_3D(V, mode, theta, u)
%
%
% 3D Example #2
%
% 2*pi/3 rotation of basis vector i around [0 0 1]' / z axis. u argument is optional in this case.
% mode = 'z';
% R = rotate_3D(V, mode, theta) % equivalent to rotate_3D([1 0]', mode, theta) here
%
%
% 3D Example #3
%
% Array example
% 2*pi/3 rotation of (i,j,k) basis vectors around [1 1 1]' is its permutation (j,k,i).
% V = eye(3);
% u = ones(3,1);
% mode = 'any';
% R = rotate_3D(V, mode, theta, u)
%
%
% 2D Example #1
%
% 3*pi/2 rotation of basis vector j around basis vector z is basis vector i.
% V = [0 1]';
% mode = 'z';
% theta = 3*pi/2;
% R = rotate_3D(V, mode, theta)
%
%
% 2D Example #2
%
% pi/6 rotation of unitary vector [0.5 sqrt(3)/2]' around the origin is basis vector j.
% (default)
% V = [0.5 sqrt(3)/2]';
% mode = 'z';
% theta = pi/6;
% R = rotate_3D(V, mode, theta)
%
%
% 2D Example #3
%
% 45° rotation of basis vector i around -0.5*[sqrt(2) sqrt(2)]' is basis vector j.
% V = [1 0]';
% u = -0.5*[sqrt(2) sqrt(2)]';
% mode = 'x'; % 2D mode automatically set back to 'z' in case of mistake.
% theta = 45; % angle unit in degree
% R = rotate_3D(V, mode, theta, u, 'degree')


%% Input parsing
Ndim = size(V,1);

assert(nargin > 2, 'Not enough input arguments.');
assert(nargin < 6, 'Too many input arguments.');
assert(Ndim > 1 && Ndim < 4, 'Input argument V must have between one and three rows : 1 < size(V,1) <= 3.');
assert(strcmpi(mode,'x') || strcmpi(mode,'y') || strcmpi(mode,'z') || strcmpi(mode,'any'),...
       'Bad mode argument : mode must be a string in the set {''x'',''X'',''y'',''Y'',''z'',''Z'',''any'',''ANY''}.');

if nargin < 5
    
    angle_unit = 'radian';
    
    if nargin < 4
        
        if Ndim == 2
            
            u = [0,0]';
            
        elseif Ndim == 3
            
            switch mode
                
                case {'x', 'X'}
                    
                    u = [1 0 0]';
                    
                case {'y', 'Y'}
                    
                    u = [0 1 0]';
                    
                case {'z', 'Z'}
                    
                    u = [0 0 1]';
                    
            end
            
        end
        
    else
        
        assert(Ndim < 3 || ~strcmpi(mode,'any') || norm(u) > 0,'3D rotation axis u must not equal null vector.');
        
    end
    
else
    
    assert(strcmpi(angle_unit,'radian') || strcmpi(angle_unit,'degree'),'angle_unit value must be either ''radian'' or ''degree''.');
    
    if strcmpi(angle_unit,'degree')
        
        theta = pi * theta / 180;
        
    end
    
end

% assert(size(u,1) == Ndim,'u (rotation around vector) and V (vector to rotate) must be vectors of the same dimension. Here size(V,1) = %s but size(u,1) = %s',num2str(Ndim),num2str(size(u,1)));
% assert(Ndim < 3 || norm(u) ~= 0 || ~strcmpi(mode,'any'), '3D rotation axis u must not equal null vector.');
% assert(imag(theta) == 0, 'Rotation angle theta must be real number in radian unit.');


%% Body

% Rotation matrix construction and resulting rotated vector computation
switch Ndim
    
    case 2 % rotate around a point (2D vector) in (xOy) plan -> mandatory rotation axis is 'z' 
        Rm = [cos(theta) -sin(theta);
              sin(theta)  cos(theta)];
        
        W = V - u;
        R = Rm * W;
        R = R + u;
                        
    case 3
        
        switch mode
            
            case {'x', 'X'} % X axis rotation matrix ; u = i = [1 0 0]'
                Rm = [1          0           0;
                      0 cos(theta) -sin(theta);
                      0 sin(theta)  cos(theta)];
            case {'y', 'Y'} % Y axis rotation matrix ; u = j = [0 1 0]'
                Rm = [cos(theta)   0  -sin(theta);
                      0            1           0;
                      sin(theta)  0  cos(theta)];
            case {'z', 'Z'} % Z axis rotation matrix ; u = k = [0 0 1]'
                Rm = [cos(theta) -sin(theta) 0;
                      sin(theta)  cos(theta) 0;
                      0           0          1];
            case {'any', 'ANY'} % Any u axis rotation matrix
                
                u = u/norm(u);
                
                Rm = [u(1,1)^2+cos(theta)*(1-u(1,1)^2) (1-cos(theta))*u(1,1)*u(2,1)-u(3,1)*sin(theta) (1-cos(theta))*u(1,1)*u(3,1)+u(2,1)*sin(theta);
                      (1-cos(theta))*u(1,1)*u(2,1)+u(3,1)*sin(theta) u(2,1)^2+cos(theta)*(1-u(2,1)^2) (1-cos(theta))*u(2,1)*u(3,1)-u(1,1)*sin(theta);
                      (1-cos(theta))*u(1,1)*u(3,1)-u(2,1)*sin(theta) (1-cos(theta))*u(2,1)*u(3,1)+u(1,1)*sin(theta) u(3,1)^2+cos(theta)*(1-u(3,1)^2)];
            otherwise
                
                error('Bad mode argument : mode must be a string in the set {''x'',''X'',''y'',''Y'',''z'',''Z'',''any'',''ANY''}.');
                
        end
        
        R = Rm * V;       
        
end


end % rotate_3D
```
