## Supplementary material for "Using pose estimation and 3D rendered models to study leg-mediated self-righting by lanternflies": Dataset S1: SLFrightingposeanalyzer.m

```
%% reads 3D xyz coordinates for body landmarks tracked using DLTdv & DLC,
% plots and analyzes them for mechanical properties

% this version uses separately tracked ground points

%%%%%%%%%%%%%%%%%%%%%%%%%%%%%%%%%%%%%%%%%%%%%%%%%%%%%%%%%%%%%%%%%%%%%%%%%%%
%% initialization
clear; close all;

%-------------------------------------

nboot = 1000;                                   % number of samples for 
                                                % bootstrapped confidence 
                                                % intervals using bootci
% plotting, fitting, and animation
show_landmarks = 0;                             % show tracked landmarks on 
                                                % animation
animpausetime = 0;                              % pause between showing 
                                                % animI_rotation images
save_animation = 1;                             % saves simulation as video

bodyalpha = 0.75;                               % body mesh transparency:
                                                % 0 = transparent
                                                % 1 = opaque 
az = -37.5; el = 5;                             % animation viewing angles  

% flags used to show extra plots for checking results
plot_raw_coords_3D = 1;                         % plot all data in 3D
plot_energy_landscape = 0;                      % plot PE vs body orientation angles 
plotbodyz = 0;

%-------------------------------------
% tracking resolution from pixel/mm and reconstruction error
% tolerance for how far off ground the feet used to define the ground plane
% were -- values of z <= Ztol are assumed to be on the ground plane
Ztol = 0.35;                 % value in mm

% define the ground plane along z at the bottom  of all z 
% landmark values below the generic ground plane for that day to correct
% for small tilt & offsets
update_ground_plane = 1;
% the percentile to use in defining the ground plane (defined to be above
% e.g., the bottom pcutoff % of all points in the dataset)
pcutoff = 10;  
% factor to multiply Ztol by to determine whether or not points are on the
% ground plane when performing fit to plane -- only used if pcfitplane is
% used to determine the ground plane;  for now we are using singular value
% decomposition instead
gnd_plane_factor = 2; 
% factor to multiply Ztol to define upper-limit of points above ground
% plane for ground contact points
support_plane_factor = 2;
%-------------------------------------
animlegendtext = {'ground plane' '\omega_{body spin}',...
    '','','','','','','','','',...
    'support centroid','COM','COM projection','vertical','pivot point',...
    'cranial' 'caudal' 'scutellum L' 'scutellum R'...
    'dorsal' 'ventral' 'tarsi & tibia-femur joints' '','','','','',...
    'cranial-caudal axis'  '' 'lateral axis'  '' 'dorsal-ventral axis'};
animlegendtext_nomarkers = {'ground plane' '\omega_{body spin}',...
    '','', '','','','','','','',...
    'support centroid','COM','COM projection','vertical','pivot point'};
linecolors = lines(7); % red/cyan/green/purple/gold/orange/blue
bluelines = linecolors(1,:); redlines = linecolors(7,:);
greenlines = linecolors(5,:); purplelines = linecolors(4,:);
orangelines = linecolors(2,:); pastelred = [250, 160, 160]/256;
pastelblue = [167, 199, 231]/256;
pastelorange = [250, 200, 152]/256;

landmarkcolors = [linecolors(7,:);linecolors(4,:);...
    linecolors(5,:);linecolors(3,:);linecolors(1:end,:)];

markercolors = lines(10);
htmlgray = 1.2*[128,128,128]/256;   % gray for boxes, etc
darkgray = [128,128,128]/256;
legcolor = [80 80 80]/256;          % color of legs on animation

% markertypes = {'o','s','^','h','d','v','p','>','<','x','*'}; % all markers
markertypes = {'s','v','<','>','o','o','p','h','d','x','+','o','o'};
% legs 1-6 are: right foreleg, right midleg, right hindleg, left hindleg, left midleg,left foreleg
leglegend1 = {'rightfore','rightmid','righthind','lefthind','leftmid',...
    'leftfore'};
leglegend2 = {'rightfore','','rightmid','','righthind','',...
    'lefthind','','leftmid','','leftfore',''};
leglegend3 = {'RF' 'RM' 'RH' 'LH' 'LM' 'LF'};                                                
%-------------------------------------
% fitting & video parameters

% video parameters
framerate = 1000;                               % video frames/sec
dt = 1/framerate;                               % timestep/video frame (s)

smoothwindow = round(25e-3/dt);                 % 25 ms smoothing time
% used for smoothing tracked coordinates and computing slopes

max_NaN_gap = ceil(smoothwindow/3);             % maximum number of missing 
                                                % NaN datapoints interpolated
%-------------------------------------
% acceleration gravity m s^-2
g = 9.80;                                       

%%%%%%%%%%%%%%%%%%%%%%%%%%%%%%%%%%%%%%%%%%%%%%%%%%%%%%%%%%%%%%%%%%%%%%%%%%%
%% Import xyz coordinates for legs and COM data

[file, pathstr] = uigetfile('*.csv','Select DLT tracks from .csv file');
coordsfile = [pathstr file];
% get filename parts for creating output file names
[pathstr,filename,file_ext] = fileparts(coordsfile);

if save_animation
    videofilename = fullfile(pathstr,[filename(1:23),'.mp4']);
    writerObj = VideoWriter(videofilename,'MPEG-4');
    open(writerObj);
end

[num,txt,raw] = xlsread(coordsfile);

nframe = size(num,1);
time = 1/framerate * (1:nframe);

num = 1000*num;                     % convert num (meter) into mm:
nlandmarks = round(size(num,2)/3);  % number tracked body landmarks
ncoords = size(num,2);              % number entries in num

%%%%%%%%%%%%%%%%%%%%%%%%%%%%%%%%%%%%%%%%%%%%%%%%%%%%%%%%%%%%%%%%%%%%%%%%%%%
%% get user defined inputs
prompt = {'framerate(fps)' 'smoothing window(ms)' 'tracking error(mm)' ...
    'view azimuth(deg)','view elevation(deg)',...
    'Body mesh 1=transparent; 0=opaque' 'Show landmarks/body axes(0=No,1=Yes)' ...
    'first frame to plot' 'last frame to plot' 'instar (3 or 4)'};
dlgtitle = 'user defined inputs';
fieldsize = [1 45];
definput = {num2str(framerate),num2str(smoothwindow),num2str(Ztol),...
    num2str(az),num2str(el),num2str(bodyalpha),num2str(show_landmarks),...
    '1',num2str(nframe),'4','0'};
answer = inputdlg(prompt,dlgtitle,fieldsize,definput);
[framerate,ansflag]         = str2num(answer{1});
[smoothingtime,ansflag]     = str2num(answer{2});
[Ztol,ansflag]              = str2num(answer{3});
[az,ansflag]                = str2num(answer{4});
[el,ansflag]                = str2num(answer{5});
[bodyalpha,ansflag]         = str2num(answer{6});
[show_landmarks,ansflag]    = str2num(answer{7});
[firstframe,ansflag]        = str2num(answer{8});
[lastframe,ansflag]         = str2num(answer{9});
[instar,ansflag]            = str2num(answer{10});

dt = 1/framerate;                               % timestep/video frame (s)
smoothwindow = round(smoothingtime*1e-3/dt);    % # frames for smoothing

% account for new range of frames--only analyze and plot selected frames
time = time(firstframe:lastframe) - time(firstframe);
num = num(firstframe:lastframe,:);
nframe = numel(time); firstframe = 1; lastframe = nframe;

Ztol_gnd_plane = Ztol*gnd_plane_factor;
Ztol_support_plane = Ztol*support_plane_factor;

%%%%%%%%%%%%%%%%%%%%%%%%%%%%%%%%%%%%%%%%%%%%%%%%%%%%%%%%%%%%%%%%%%%%%%%%%%%
%-------------------------------------
% initialize spotted lanternfly nymph standard 3D model mass & body 
% dimensions

mass_whole_insect = 28.4e-6;        % mass [kg]

% spotted lanternfly nymph body dimensions from morphometrics
body_length_3Dmodel = 8.9;  % mean body snout-caudal length
if instar == 4
    body_width_3Dmodel = 5.8;
elseif instar == 3
    body_width_3Dmodel = 4.0;
else
    body_width_3Dmodel = 6.0;
end
body_thickness_3Dmodel  = 2.3;    % measured on 3D model--from photos &
% micrographs, body thickness/length ratio ranges from 0.23 to 0.26
% SLF nymph leg dimensions in terms of 
% fraction of body snout-caudal length; fore,mid,hind
coxa_femur_length = [0.70,0.68,0.42]*body_length_3Dmodel;  
% distance from knee to tarsus tip:
tibia_tarsus_length = [0.65,0.63,0.85]*body_length_3Dmodel;
tarsus_length = [0.15 0.15 0.15]*body_length_3Dmodel; % tarsus-only length

% make each array have 6 elements consistent with the leg numbering (Left
% fore-hind, right hind-fore)
coxa_femur_length = [coxa_femur_length fliplr(coxa_femur_length)];
tibia_tarsus_length = [tibia_tarsus_length fliplr(tibia_tarsus_length)];
tarsus_length = [tarsus_length fliplr(tarsus_length)];

% compute body & leg segments mass using their fractions of total body mass
% fore,mid,hind order -- all masses are in [kg]
mass_body_only = 0.70*mass_whole_insect;
femur_coxa_mass = [2.7,2.6,1.6]*0.01*mass_whole_insect;
femur_coxa_mass = [femur_coxa_mass fliplr(femur_coxa_mass)];
tibia_tarsus_mass = [2.5,2.4,3.2]*0.01*mass_whole_insect;
tibia_tarsus_mass = [tibia_tarsus_mass fliplr(tibia_tarsus_mass)];

% compute I_femur and I_tibia about their COM in units of kg m^2
% *1e-6 converts length^2 from mm^2 to m^2
I_femur = (1/12)*femur_coxa_mass.*coxa_femur_length.*coxa_femur_length*1e-6;
I_tibia = (1/12)*tibia_tarsus_mass.*tibia_tarsus_length.*tibia_tarsus_length*1e-6;

%% get body mesh in units of mm
% get the mesh body-only 3D model from file and its properties
% and the points on the body where the legs insert (base of coxa)
[bodymesh,coxacoord,cranialcoord,caudalcoord]= getbodymesh(instar);
bodyConnect = bodymesh.ConnectivityList; % doesn't change
bodyPoints = bodymesh.Points;            % initial orientation

% standard I_rot in the order roll, pitch, yaw required for calculations;
% corresponds to the actual body orientation used

I_rot = [4.64e-11	0	0;
            0	1.25e-10	0;
            0	0	1.50e-10];

% Iyaw, Ipitch, Iroll for standard 3D model
Iroll = I_rot(1,1); Ipitch = I_rot(2,2); Iyaw = I_rot(3,3); %[kg m^2]

% reference orientation of the body frame for rotations (mesh starts
% out with its body axes oriented along XYZ)
referencepts = [1 0 0; 0 1 0; 0 0 1];

%% import file with ground contact points
% [groundptsfile, grndpathstr] = uigetfile('*.csv','Select ground points .csv file');
% groundptsfile = [grndpathstr groundptsfile];
groundptsfile = fullfile(pathstr,'groundplane_xyzpts.csv');

[groundptcoords,~,~]= xlsread(groundptsfile);
groundptcoords = 1000*groundptcoords;            % convert meter into mm:

X_ground = groundptcoords(:,1:3:end);
Y_ground = groundptcoords(:,2:3:end);
Z_ground = groundptcoords(:,3:3:end);

% get the ground plane & correct num for rotation & the offset in z
[num,Z_ground] = get_ground_plane(num,X_ground,Y_ground,Z_ground,nlandmarks,Ztol_gnd_plane);

% After this step, the xy plane should be approximately horizontal 
% and the body +z-axis should point vertically upward, and z should extend
% from Z = 0 upward

%% preprocess body landmark coordinates 
% > fill in very short stretches of missing data & replace outliers
% > smooth using a low-pass filter window to take out only the
% high frequency noise (>= 200 Hz noise for leg points, >= 40 Hz for body 
% points

for k = 1:27                % smooth body points over entire window
    num(:,k) = filloutliers(num(:,k),'pchip',"movmedian",smoothwindow);
    num(:,k) = smooth(num(:,k),smoothwindow);   
    num(:,k) = fillmissing(num(:,k),'pchip','MaxGap',max_NaN_gap);
end

for k = 28:63               % smooth faster leg points over 1/5 window
    num(:,k) = filloutliers(num(:,k),'pchip',"movmedian",smoothwindow);
    num(:,k) = smooth(num(:,k),ceil(smoothwindow/5));   
    num(:,k) = fillmissing(num(:,k),'pchip','MaxGap',max_NaN_gap);
end

%%
% use lowest z-values for each body landmark to estimate the 
% exact ground plane in this video;  rotate & correct for residual z-offset
% in this specific video relative to the larger ground plane on that day
if update_ground_plane
    % select landmarks to use z-values for (3 = cranial, 6 = caudal, etc.)
    Zindx = [30:3:45,51:3:60];     % z indices for all tarsi, mid & hind knees (sometimes on ground)

    leglabels = {'RF tarsus','RM tarsus','RH tarsus','LF tarsus',...
    'LM tarsus','LF tarsus','RM knee','RH knee','LH knee','LM knee'};
    % initialize set of new approximate groundplane points
    X_ground = []; Y_ground = []; Z_ground = [];
    % save only the landmarks with Z < Ztol
    for k = 1:numel(Zindx)
        idxlowZ = find(num(:,Zindx(k))<0);  % if less than zero, correct
        X_ground = [X_ground;num(idxlowZ,Zindx(k)-1)];
        Y_ground = [Y_ground;num(idxlowZ,Zindx(k)-2)];
        Z_ground = [Z_ground;num(idxlowZ,Zindx(k))];
    end

    % as above, assemble into Xground, Yground, Zground & repeat process above
    % get the ground plane & correct num for rotation & offset in z
    [num,Z_ground] = get_ground_plane(num,X_ground,Y_ground,Z_ground,nlandmarks,Ztol_gnd_plane);
    
    % plot corrected landmarks to see if constant z values are level
    figure; 
    for k = 1:numel(Zindx)
        subplot(ceil(numel(Zindx))/2,2,k)
        plot(time,num(:,Zindx(k)),'-'); hold on; grid on;
        ylabel(leglabels{k});xlabel('time (s)');
    end
    %% subtract off estimate of corrected ground plane z based on last few 
    % frames where the tarsis are on the ground
    % estimate z offset
    tarsi_righted = num((end-5):end,30:3:45); 
    Zmin = median(tarsi_righted(tarsi_righted < Ztol));
    % subtract it off
    num(:,Zindx) = num(:,Zindx) - Zmin;
    % plot landmarks for comparison with uncorrected values
    for k = 1:numel(Zindx)
        subplot(ceil(numel(Zindx))/2,2,k)
        plot(time,num(:,Zindx(k)),'--');
    end
end

%% subtract off an approximate x,y center of mass point (estimated by the 
% mean of the cranial and caudal points) to center body in the animation
num(:,1:3:end) = num(:,1:3:end)  - mean((num(:,1)+num(:,4))/2,'omitnan');
num(:,2:3:end) = num(:,2:3:end)  - mean((num(:,2)+num(:,5))/2,'omitnan');

%% Match preprocessed tracked coordinates to body landmarks
% cranial = pt 1
% caudal = pt 2
% left scuttellum = pt 3
% right scutellum = pt 4
% pt 5,6 = spots that we didn't track in the end
% dorsal = pt 7
% ventral = pt 8
% body pivot point on ground = pt 9
% tarsi = pts 10-15
% knees = pts 16-21 (tibia-femur joint)

cranial = num(:,1:3);  caudal = num(:,4:6); 

scutLeft = num(:,7:9); scutRight = num(:,10:12);

% ignore spots -- weren't tracked (13:15 & 16:18 = NaN) 
num(:,13:15)=nan(size(num(:,13:15)));num(:,16:18)=nan(size(num(:,16:18)));

dorsal = num(:,19:21); ventral = num(:,22:24);

pivot_pt = num(:,25:27);        % tracked body pivot point on ground

%% compute the body axes (roll-pitch-yaw axes using MATLAB conventions)
%--------------------------
% 1. get the cranial-caudal (roll) axis <-- we trust this one unless
% it's missing altogether
cranial_caudal_axis = (cranial-caudal)./vecnorm(cranial - caudal,2,2);            % X (roll)

%--------------------------
% 2. compute the lateral (pitch) axis
lateral_axis = (scutLeft - scutRight)./vecnorm(scutRight - scutLeft,2,2);     % Y (pitch)    

% ensure lateral is perpendicular to cranial-caudal axis
lateral_axis = lateral_axis - dot(lateral_axis,cranial_caudal_axis,2).*cranial_caudal_axis;

%--------------------------
% 3. compute the dorsal-ventral axis (yaw) from other axes

% for NaN dorsal and/or ventral points, get dv axis from other body axes
dorsal_ventral_axis = cross(cranial_caudal_axis,lateral_axis);  % Z (yaw)

% compute the dorsal-ventral axis from tracked coords
dorsal_ventral_axis_tracked = (ventral - dorsal)./vecnorm(ventral - dorsal,2,2);  % Z (yaw)

%--------------------------
% use body axes to define the body-fixed reference frame using standard
% orientations:
Xbody = cranial_caudal_axis; Ybody = lateral_axis;
Zbody = dorsal_ventral_axis;

validbodyaxes = ~isnan(Xbody.*Ybody.*Zbody);  % all entries that aren't NaN
if sum(sum(~validbodyaxes)) ~= 0
    fprintf('Number missing values of computed body axes = %d\n',sum(sum(~validbodyaxes)));
end
%% find time when dorsal-ventral axis flips (i.e., is approximately horizontal)
% the dorsal-ventral axis's Z component, Z_dv = dorsal_ventral_axis(:,3)
% The dorsal-ventral axis points upward (Z_dv>0) when overturned, and downward
% (<0) when upright; it "flips" when changing sign (~0)
[~,idxdvz0] = min(abs(dorsal_ventral_axis(:,3)));

%% find factor for rescaling the tracked body coordinates so they have 
%  approx. same body length as the 3D model mesh

% initial tracked body length in each frame
% cranial_caudal_axis is a unit vector
body_length = abs(dot((cranial-caudal),cranial_caudal_axis,2));

rescale_body_length = body_length_3Dmodel/median(body_length(:),'omitnan');

% rescaled body length in each frame matches 3D model's on average
body_length = body_length*rescale_body_length;  

%% rescale all coordinates by *rescale_body_length so all dimensions are
% standardized to agree with those of the 3D model

num = num*rescale_body_length;

cranial = cranial*rescale_body_length;  caudal = caudal*rescale_body_length; 
scutLeft = scutLeft*rescale_body_length; scutRight = scutRight*rescale_body_length;
dorsal = dorsal*rescale_body_length; ventral = ventral*rescale_body_length;

%% save the tarsus and knee coordinates
tarsuscoords = num(:,[28:45]);
kneecoords = num(:,[46:63]);

%% rescale the body mesh so its width and thickness agree with those of the
% tracked points

body_width = vecnorm((scutLeft - scutRight),2,2);
% use trimmed mean to avoid outliers & very large midtracked points
% independent of percentage used over 10-25%
rescale_body_width = trimmean(body_width(:),15)/body_width_3Dmodel;

% rescales the 3D model mesh dimensions along y and z in body frame 
% for modeling and animations
if rescale_body_width < 1
    fprintf('rescaling based on body width\n')
    coxacoord(:,2)=coxacoord(:,2)*rescale_body_width;
    bodyPoints(:,2)=bodyPoints(:,2)*rescale_body_width;
    bodyPoints(:,3)=bodyPoints(:,3)*rescale_body_width;
    coxacoord(:,3)=coxacoord(:,3)*rescale_body_width;
end

body_thickness = vecnorm((dorsal-ventral),2,2);
% use trimmed mean to avoid outliers & very large midtracked points
% independent of percentage used over 10-25%
rescale_body_thickness = trimmean(body_thickness(:),25)/body_thickness_3Dmodel;
if rescale_body_thickness < rescale_body_width
    fprintf('also rescaling body z values based on body thickness\n')
    bodyPoints(:,3)=bodyPoints(:,3)*rescale_body_thickness/rescale_body_width;
    coxacoord(:,3)=coxacoord(:,3)*rescale_body_thickness/rescale_body_width;
else 
    rescale_body_thickness = rescale_body_width;
end

%% Compute the body COM: rotate the cranial and caudal points on the 3D 
%  model and use their current positions to get shift in COM
COM_body = nan(nframe,3);

for i = 1:nframe 
    %---------------------
    % compute body mesh with orientation defined by array with 
    % columns = current orientations of body-fixed axes

    targetpts = [Xbody(i,:)',Ybody(i,:)',Zbody(i,:)'];

    temp = isnan(targetpts(:));                 % cannot do if any NaNs

    if sum(temp) == 0                           % if no NaNs
        % get rotation matrix rotm to perform the
        % current rotation from the reference orientation to the current
        % orientation
        rotm = get_rot_matrix(referencepts,targetpts);
        % rotate cranial and caudal points to agree with current body axis 
        % orientation and 
        cranial_rot = rotm*cranialcoord';       % rotate cranial coords
        caudal_rot  = rotm*caudalcoord';         % rotate caudal coords

        % shift COM from origin to current value using a weighted average 
        % shift of the cranial and caudal points from their position on the
        % 3D model
        COM_body(i,:) = 0.75*(cranial(i,:) - cranial_rot') + ...
            0.25*(caudal(i,:)- caudal_rot');
    end
end

% get distance from COM to ground plane for potential energy measurements
COM_height = COM_body(:,3);

%% find the maximum values of all body coordinates except the untracked
% spots and the hard-to-track-pivot point; these are used for plot axis
% ranges
allbodyX = num(:,1:3:end); allbodyY = num(:,2:3:end); allbodyZ = num(:,3:3:end);
allbodyX(:,[5,6,9]) = NaN ;allbodyY(:,[5,6,9]) = NaN ; allbodyZ(:,[5,6,9]) = NaN;
maxX = max(allbodyX,[],'omitnan');minX = min(allbodyX,[],'omitnan');
maxY = max(allbodyY,[],'omitnan');minY = min(allbodyY,[],'omitnan');
maxZ = max(allbodyZ,[],'omitnan');
minZ = min(allbodyZ,[],'omitnan');
minZ = -1;                          % truncate minimum Z plotted at -1 mm
maxX = max(maxX,[],'omitnan'); minX = min(minX,[],'omitnan');
maxY = max(maxY,[],'omitnan'); minY = min(minY,[],'omitnan');
% make the XY area square:
minX = min(minY,minX);minY = minX; maxX = max(maxY,maxX);maxY = maxX;

maxZ = max(maxZ,[],'omitnan'); minZ = min(minZ,[],'omitnan');

%%%%%%%%%%%%%%%%%%%%%%%%%%%%%%%%%%%%%%%%%%%%%%%%%%%%%%%%%%%%%%%%%%%%%%%%%%%
%% create animation while computing rotational properties, energies, etc.
figure('Position',[50 10 900 900]);

% initialize data arrays
stability_margin = nan(nframe,1);ideal_stability_margin = nan(nframe,1); 
bodymeshminz = nan(nframe,1);
femur_COM = nan(nframe,6,3);       % femur COM
tibia_COM = nan(nframe,6,3);       % tibia COM
quat_array = nan(nframe,4);
I_spin_body = nan(1,nframe);
I_spin_each_leg = nan(nframe,3); 
PE_legs = zeros(nframe,1);
KE_spin_body = nan(nframe,1);
I_spin_legs = zeros(nframe,1);
angular_speed = nan(1,nframe);
rotation_axis = nan(nframe,3);
rotation_axis_body_frame = nan(nframe,3);
angle_rot_axis_bodyX = nan(1,nframe);
leg_COM_mass = nan(nframe,6,3);
w_orbital = nan(nframe,1);
pivot_pt_on_ground = nan(nframe,2);
orbitalvector = nan(nframe,3);

ncontactpts = zeros(nframe,1);          % number ground contact points
nbodycontactpts = zeros(nframe,1);      % " " " body-only
ntarsicontactpts = zeros(nframe,1);     % " " " tarsi-only
nkneecontactpts = zeros(nframe,1);      % " " " knees-only
nlegcontactpts = zeros(nframe,1);       % " " " legs-only
mean_legs_contactXY = zeros(nframe,2);  % mean contact point on ground for all legs

% perform calculations & create animation
for i = 1:nframe                        % loop over all frames in video
                                        % showing 3D coords of tracked
                                        % points + body axes as arrows

    title(['time = ',num2str(time(i),'%.3f'),' s'],'FontSize',12);
    %---------------------
    % plot the ground plane as a transparent gray rectangular patch
    XGP = [minX maxX maxX minX]; 
    YGP = [minY minY maxY maxY];
    ZGP = zeros(size(XGP));
    patch(XGP,YGP,ZGP,[192/256,192/256,192/256],'FaceAlpha',0.5,...
        'EdgeColor',[192/256,192/256,192/256]); hold on;
    %---------------------
    % animation axis limits
    axis([minX maxX minY maxY minZ maxZ ]);
    axis equal;   
    %---------------------
    % compute the spin angular velocity & show as a vector on the animation
    if i > 1
        sourcepts = [Xbody(i-1,:)',Ybody(i-1,:)',Zbody(i-1,:)'];
        targetpts = [Xbody(i,:)',Ybody(i,:)',Zbody(i,:)'];
        temp = [sourcepts(:),targetpts(:)]; temp = isnan(temp(:));
        if sum(temp) == 0                           % if no NaNs

            % compute the angular velocity vector in the spatial frame
            w = get_ang_vel(sourcepts,targetpts,dt); % w in Hz = rev/s
            angular_speed(i) = vecnorm(w,2,2);
            angular_velocity(i,:) = w;

            rotation_axis(i,:) = w./angular_speed(i); % spatial frame

            % this version gives orientation relative to the body axes
            % (i.e., convert to the body frame coordinates x,y,z)
            rotation_axis_body_frame(i,:) = ...
                [dot(rotation_axis(i,:),Xbody(i,:)),...
                 dot(rotation_axis(i,:),Ybody(i,:)),...
                 dot(rotation_axis(i,:),Zbody(i,:)),];
            % double-check for normalization
            rotation_axis_body_frame(i,:) = rotation_axis_body_frame(i,:)/...
                vecnorm(rotation_axis_body_frame(i,:),2,2);

            I_spin_body(i) = rotation_axis_body_frame(i,:)*I_rot*...
                rotation_axis_body_frame(i,:)';

            % kinetic energy of rotation 
            % (factor of (2*pi)^2 converts Hz = rev/s to rad/s)
            KE_spin_body(i) = 0.5*I_spin_body(i)*angular_speed(i)*...
                angular_speed(i)*(2*pi)^2; 

            % plot the angular velocity vector as an arrow
            warrowscale = 0.35*body_length(i);
           
            P1 = [COM_body(i,1),COM_body(i,2),COM_body(i,3)]; 
            P2 = P1 + [warrowscale*rotation_axis(i,1),...
                warrowscale*rotation_axis(i,2),...
                warrowscale*rotation_axis(i,3)];
            arrow3(P1,P2,'q-1');  % plot angular velocity (gold/orange)
        else
            w = NaN; % w in Hz = rev/s
            angular_speed(i) = NaN;
            rotation_axis(i,:) = [NaN NaN NaN];
            I_spin_body(i) = NaN;
        end
    end
    clear temp;

    %----------------------
    % find ground contact points on legs
    % (will test if any body mesh contact point below)

    Xtarsi = []; Ytarsi = [];
    Xtemp = []; Ytemp = [];         % temporary arrays for holding ground 
                                    % support coords because we reorder 
                                    % points before creating the polygon

    % find tarsi & "knees" in contact with the ground (ignore those values for
    % which there are NaN coordinates)
    for j = 1:6
        if tarsuscoords(i,3*j) <=Ztol_support_plane && sum(isnan(tarsuscoords(i,(1:3)+(j-1)*3))) == 0
            % save ground contact points
            Xtemp = [Xtemp tarsuscoords(i,1+(j-1)*3)];
            Ytemp = [Ytemp tarsuscoords(i,2+(j-1)*3)];
            ncontactpts(i) = ncontactpts(i) + 1;
            % also keep track of tarsi only contact points
            Xtarsi = [Xtarsi tarsuscoords(i,1+(j-1)*3)];
            Ytarsi = [Ytarsi tarsuscoords(i,2+(j-1)*3)];

            ntarsicontactpts(i) = ntarsicontactpts(i) + 1;
        end
        if kneecoords(i,3*j) <=Ztol_support_plane && sum(isnan(kneecoords(i,(1:3)+(j-1)*3))) == 0
            % save knee contact points
            nkneecontactpts(i) = nkneecontactpts(i) + 1;
            Xtemp = [Xtemp kneecoords(i,1+(j-1)*3)];
            Ytemp = [Ytemp kneecoords(i,2+(j-1)*3)];
            ncontactpts(i) = ncontactpts(i) + 1;
        end
        if (tarsuscoords(i,3*j) <=Ztol_support_plane && ...
            sum(isnan(tarsuscoords(i,(1:3)+(j-1)*3))) == 0) || ...
                (kneecoords(i,3*j) <=Ztol_support_plane && ...
                sum(isnan(kneecoords(i,(1:3)+(j-1)*3))) == 0)
            nlegcontactpts(i) = nlegcontactpts(i) + 1;
        end
    end

    %---------------------
    % compute & plot body mesh with orientation defined by array with 
    % columns = current orientations of body-fixed axes

    targetpts = [Xbody(i,:)',Ybody(i,:)',Zbody(i,:)'];

    temp = isnan(targetpts(:));                 % cannot plot if any NaNs
    if sum(temp) == 0                           % if no NaNs
        % get rotation matrix rotm to perform the
        % current rotation from the reference orientation to the current
        % orientation
        rotm = get_rot_matrix(referencepts,targetpts);
        % save quaternions
        quat_array(i,:) = rotm2quat(rotm);
        % rotate body mesh to agree with current body axis orientation
        % and shift COM from origin to current value
        bodyPointsrot = rotm*bodyPoints';     % now perform 3D rotation on mesh
        bodyPointsrot = bodyPointsrot'+ COM_body(i,:);  % restore correct format & COM

        % find lowest body point & test to see if it is on the ground
        [bodymeshminz(i),bodymeshminzidx] = min(bodyPointsrot(:,3));
        if bodymeshminz(i) <= Ztol_support_plane  % if on the ground, save as a ground 
                                 % support point & pivot point
            Xtemp = [Xtemp bodyPointsrot(bodymeshminzidx,1)];
            Ytemp = [Ytemp bodyPointsrot(bodymeshminzidx,2)];
            pivot_pt_on_ground(i,1) = bodyPointsrot(bodymeshminzidx,1);
            pivot_pt_on_ground(i,2) = bodyPointsrot(bodymeshminzidx,2);
            ncontactpts(i) = ncontactpts(i) + 1;
            nbodycontactpts(i) = nbodycontactpts(i) + 1;
        end
        % make an updated mesh
        body_rot = triangulation(bodyConnect,bodyPointsrot);

        % rotate the coxa coordinates on the body
        legcoxa_rot = rotm*coxacoord';
        % shift to correct height
        legcoxa_rot = legcoxa_rot' + COM_body(i,:);

        % plot rotated mesh
        colormap bone;   
        trimesh(body_rot,'FaceAlpha',bodyalpha,'FaceColor','interp',...
            'LineStyle','none');
        hold on;
 
        % create diffuse lighting source at camera position
        camlight('headlight');
        lighting gouraud; material shiny;
    end
    clear temp;
    %---------------------
    % plot the legs & compute their contribution to the PE & KE
    for j = 1:6
        if exist('legcoxa_rot','var')  % if the leg poses could be computed
            % create an ankle joint that's only for display -- not used for
            % calculations
            % unit vector that points along femur from coxa to "knee"
            femur = legcoxa_rot(j,:)-kneecoords(i,(1:3)+(j-1)*3); 
            femur = femur/norm(femur);
            % approximate "ankle" = tarsus-tibia joint coordinates
            % estimated from the microscopic morphometric measurements
            anklecoord = tarsuscoords(i,(1:3)+(j-1)*3)+tarsus_length(j)*femur;
            % prevent the ankle from going below the tarsus (it's only for
            % display)
            anklecoord(3) = max(tarsuscoords(i,3+(j-1)*3),anklecoord(3));

            % compute femur and tibia line segments
            % compute unit vector along each leg
            femur_line_segment(i,j,:) = legcoxa_rot(j,:)-kneecoords(i,(1:3)+(j-1)*3);
            % reshape for calculations
            femur_ls = reshape(femur_line_segment(i,j,:),[1,3]);
            femur_ls = femur_ls/norm(femur_ls);     % normalize
            femur_line_segment(i,j,:) = femur_ls;   % save for later

            tibia_line_segment(i,j,:) = kneecoords(i,(1:3)+(j-1)*3)-tarsuscoords(i,(1:3)+(j-1)*3);
            % reshape for calculations
            tibia_ls = reshape(tibia_line_segment(i,j,:),[1,3]);
            tibia_ls = tibia_ls/norm(tibia_ls);     % normalize
            tibia_line_segment(i,j,:) = tibia_ls;   % save for later

            % plot legs
            plot3([legcoxa_rot(j,1),kneecoords(i,1+(j-1)*3),anklecoord(1),tarsuscoords(i,1+(j-1)*3)],...
                [legcoxa_rot(j,2),kneecoords(i,2+(j-1)*3),anklecoord(2),tarsuscoords(i,2+(j-1)*3)],...
                [legcoxa_rot(j,3),kneecoords(i,3+(j-1)*3),anklecoord(3),tarsuscoords(i,3+(j-1)*3)],'-',...
                'LineWidth',2.5,'Color',legcolor); hold on;

            % get the leg COM coordinates for each leg for KE calculation
            for kk = 1:3            % sum over kk = XYZ contributions
                femur_COM(i,j,kk) = (legcoxa_rot(j,kk)+kneecoords(i,kk+(j-1)*3))*0.5;
                tibia_COM(i,j,kk) = (kneecoords(i,kk+(j-1)*3)+tarsuscoords(i,kk+(j-1)*3))*0.5;
                % add contribution to PE for kk == 3 Z only
                if kk == 3
                    % PE for legs is m_leg*g* height above the ground (z); approx. for
                    % now as height halfway from tarsus tip to coxa
                    temp = femur_coxa_mass(j)*g*femur_COM(i,j,3) + ...
                        tibia_tarsus_mass(j)*g*tibia_COM(i,j,3);
                    if ~isnan(temp)
                        PE_legs(i) = PE_legs(i) + temp;
                    end
                    clear temp;
                end
                % save the contribution of each leg to the whole insect COM
                % z-component for later use
                leg_COM_mass(i,j,kk) = femur_coxa_mass(j)*femur_COM(i,j,kk) + ...
                    tibia_tarsus_mass(j)*tibia_COM(i,j,kk);
            end
            % if the rotation axis is defined (not NaN)
            % compute each leg's contribution to the moment of inertia, its
            % angular speed, and its rotational KE
            if sum(isnan(rotation_axis(i,:))) == 0
                % compute angle (rad) between leg segment & rotation_axis
                femur_angle = acos(dot(rotation_axis(i,:),femur_ls));
                tibia_angle = acos(dot(rotation_axis(i,:),tibia_ls));
                % put leg segment COM in correct format
                femurpt = squeeze(femur_COM(i,j,:)); femurpt = femurpt';
                tibiapt = squeeze(tibia_COM(i,j,:)); tibiapt = tibiapt';
                % make a line segment long enough that we can compute
                % distance to it for each leg
                rot_line_coords = rotation_axis(i,:)*15; 

                % compute full spin moment of inertia for each leg using
                % the initial I values about the tibia and femur COM
                % The factors of sin^2 correct for the rotational axis's
                % angle with respect to the rod-like leg sement long
                % directions.
                % We also add a term based on the parallel axis theorem 
                % proportional to the segment mass and square of the 
                % the distance from leg segment COM to line parallel to the
                % rotation axis passing through the body COM
                dist_femur = point_to_line_distance(femurpt,...
                    -rot_line_coords,rot_line_coords);
                dist_tibia = point_to_line_distance(tibiapt,...
                    -rot_line_coords,rot_line_coords);
                % *1e-6 converts dist_femur and tibia from mm^2 to m^2
                I_spin_each_leg(i,j) = I_tibia(j)*sin(tibia_angle)^2 + ...
                    tibia_tarsus_mass(j)*dist_tibia^2*1e-6 + ...
                    I_femur(j)*sin(femur_angle)^2 + ...
                    femur_coxa_mass(j)*dist_femur^2*1e-6;
                I_spin_legs(i) = I_spin_legs(i) + I_spin_each_leg(i,j);
            end
        end
    end
    %---------------------
    % plot gray polygon corresponding to support plane, defined as all
    % points within a threshold distance of the ground plane
    % make partially transparent so we can see points below it
    
    % reorder the ground contact points in CW ofder
    [Xtemp,Ytemp] = poly2cw(Xtemp,Ytemp);
    % create support polygon with these as vertices 
    X_support{i} = Xtemp;
    Y_support{i} = Ytemp;
    Z_support{i} = zeros(size(Xtemp));
    X_tarsi_support{i} = Xtarsi;
    Y_tarsi_support{i} = Ytarsi;

    if numel(X_support{i}) > 2 
        % compute the convex hull of the support polygon points
        supportpts = [X_support{i}' Y_support{i}'];
        cvidx = convhull(supportpts,'Simplify',true);
        % plot support polygon as a gray patch
        patch(supportpts(cvidx,1),...
            supportpts(cvidx,2),...
            zeros(size(supportpts(cvidx,1))),...
            [192/256,192/256,192/256],'FaceAlpha',0.5);
        % make up a polygon for the support pattern
        support_polygon = polyshape(supportpts(cvidx,1),...
            supportpts(cvidx,2));

        % compute & plot the support pattern centroid
        % if it agrees with the COM, the pose is maximally stable
        [x_support_polygon,y_support_polygon] = centroid(support_polygon);
        plot(x_support_polygon,y_support_polygon,'xk','MarkerSize',15,'LineWidth',1.5);

        % compute stability margin = minimum distance from the COM to 
        % the support polygon boundary; ideal stability margin = min dist
        % from the support polygon centroid to its boundaries
        [stability_margin(i),~,~] = ...
            p_poly_dist(COM_body(i,1),COM_body(i,2),supportpts(cvidx,1),...
            supportpts(cvidx,2));
        [ideal_stability_margin(i),~,~] = ...
        p_poly_dist(x_support_polygon,y_support_polygon,supportpts(cvidx,1),...
            supportpts(cvidx,2));

    elseif numel(X_support{i}) == 2    % show 2 ground contact pts as a line
        plot3(X_support{i},Y_support{i},Z_support{i},'-','Color',...
            [192/256,192/256,192/256]);
    end

    %---------------------
    % plot COM on ground plane vs time
    if ~anynan(COM_body(i,:))                        % only plot if not NaN
        % plot COM as open o
        plot3(COM_body(i,1),COM_body(i,2),COM_body(i,3),'ko','MarkerSize',9);
        % plot COM projected onto ground as closed o
        plot3(COM_body(i,1),COM_body(i,2),0,'ko','MarkerSize',9,'MarkerFaceColor','k');
        % plot vertical downward from body COM
        plot3([COM_body(i,1),COM_body(i,1)],[COM_body(i,2),COM_body(i,2)],...
            [0,COM_body(i,3)],'k--');
    end

    %----------------------
    % plot pivot_pt_on_ground if not NaN
    if ~anynan(pivot_pt_on_ground(i,:))          % only plot if not NaN
        plot3(pivot_pt_on_ground(i,1),pivot_pt_on_ground(i,2),0,'k*');
    end
    %----------------------
    if show_landmarks
        %---------------------
        % plot all tracked body landmarks as markers
        plot3(cranial(i,1),cranial(i,2),cranial(i,3),...
            markertypes{1},'MarkerEdgeColor',landmarkcolors(1,:),...
            'LineWidth',1); hold on;
        plot3(caudal(i,1),caudal(i,2),caudal(i,3),...
            markertypes{1},'MarkerEdgeColor',landmarkcolors(1,:),...
            'LineWidth',1);
        plot3(scutLeft(i,1),scutLeft(i,2),scutLeft(i,3),...
            markertypes{2},'MarkerEdgeColor',landmarkcolors(2,:),...
            'LineWidth',1);
        plot3(scutRight(i,1),scutRight(i,2),scutRight(i,3),...
            markertypes{2},'MarkerEdgeColor',landmarkcolors(2,:),...
            'LineWidth',1);
        if dorsal(i,3) >= 0  % only plot dorsum if not covered up and estimated below the surface)
            plot3(dorsal(i,1),dorsal(i,2),dorsal(i,3),...
                markertypes{3},'MarkerEdgeColor',landmarkcolors(3,:),...
            'LineWidth',1); hold on;
        else plot3(NaN,NaN,NaN,markertypes{3},'MarkerEdgeColor',landmarkcolors(3,:));
        end
        if ventral(i,3) >= 0  % only plot ventrum if not covered up and estimated below the surface)
            plot3(ventral(i,1),ventral(i,2),ventral(i,3),...
                markertypes{3},'MarkerEdgeColor',landmarkcolors(3,:),...
            'LineWidth',1);
        else plot3(NaN,NaN,NaN,markertypes{3},'MarkerEdgeColor',landmarkcolors(3,:),...
            'LineWidth',1);
        end
        for j = 1:6
        plot3([kneecoords(i,1+(j-1)*3),tarsuscoords(i,1+(j-1)*3)],...
            [kneecoords(i,2+(j-1)*3),tarsuscoords(i,2+(j-1)*3)],...
            [kneecoords(i,3+(j-1)*3),tarsuscoords(i,3+(j-1)*3)],'+',...
            'LineWidth',2.5,'Color',legcolor); hold on;
        end
        %---------------------
        % plot body axes
        % cranial-caudal axis arrow
        P1 = [COM_body(i,1),COM_body(i,2),COM_body(i,3)];
        P2 = P1 + 0.75*[body_length(i)*Xbody(i,1),...
            body_length(i)*Xbody(i,2),body_length(i)*Xbody(i,3)];
        arrow3(P1,P2,'f-1');  % red
        % lateral axis arrow
        P2 = P1 + 0.75*[body_width(i)*Ybody(i,1),...
            body_width(i)*Ybody(i,2),body_width(i)*Ybody(i,3)];
        arrow3(P1,P2,'b-1');   % blue
        % dorsal-ventral axis arrow
        P2 = P1 + 0.3*[body_length(i)*Zbody(i,1),...
            body_length(i)*Zbody(i,2),body_length(i)*Zbody(i,3)];
        arrow3(P1,P2,'a-1');   % green
    end
    %---------------------
    xlabel('X(mm)');ylabel('Y(mm)'); zlabel('Z(mm)');
    grid off;

    view(az,el);                              

    % create legend for the animation
    if show_landmarks
        legend(animlegendtext,'Location','bestoutside','Orientation',...
            'horizontal','NumColumns',5,'FontSize',10);
    else
        legend(animlegendtext_nomarkers,'Location','bestoutside',...
            'Orientation','horizontal','NumColumns',5,'FontSize',10); 
    end
    legend boxoff; 
    drawnow; pause(animpausetime);

    if save_animation               % save frame for animation video
        % animation frame grab (use gcf not just gca)
        animframe = getframe(gcf);
        writeVideo(writerObj,animframe);
    end

    % save an image of when it's flipped (i.e., when the dorsal-ventral
    % axis z component, Z_dv ~= 0 (changes sign)
    if i == idxdvz0
        % plot the angular velocity vector as an arrow with its origin at
        % the pivot point
        warrowscale = body_length(i);
        P1 = [pivot_pt_on_ground(i,1),pivot_pt_on_ground(i,2),0];
        P2 = P1 + [warrowscale*rotation_axis(i,1),...
            warrowscale*rotation_axis(i,2),...
            warrowscale*rotation_axis(i,3)];
        arrow3(P1,P2,'q-1');  % plot angular velocity (gold/orange)
        flipfilename = fullfile(pathstr,[filename(1:23),'_flip.png']);
        saveas(gcf,flipfilename);
    elseif i == firstframe
        firstfilename = fullfile(pathstr,[filename(1:23),'_overt.png']);
        saveas(gcf,firstfilename);
    elseif i == lastframe
        lastfilename = fullfile(pathstr,[filename(1:23),'_right.png']);
        saveas(gcf,lastfilename);        
    end

    if i ~= nframe; clf; end

end             % end of simulation time loop

% close animation video
if save_animation; close(writerObj); end

%%%%%%%%%%%%%%%%%%%%%%%%%%%%%%%%%%%%%%%%%%%%%%%%%%%%%%%%%%%%%%%%%%%%%%%%%%%
%% compute combined kinematic parameters before plotting & saving

%%--------------------
% compute the periodicity of the tarsi motions along body frame z_body
tarsus_zbody = nan(nframe,6);
for j = 1:6
    tarsus_zbody(:,j) = dot((tarsuscoords(:,(1:3)+(j-1)*3)-COM_body),Zbody,2);
end

%--------------------------------------------------------------------------
% compute whole insect COM
% first get mass*COM coordinates for all legs in each frame
mass_COM_all_legs = zeros(nframe,3);
for j = 1:6
    mass_COM_all_legs = mass_COM_all_legs + squeeze(leg_COM_mass(:,j,:));
end
COM_whole_insect = (mass_body_only*COM_body + mass_COM_all_legs)/...
    mass_whole_insect;
diffCOM = COM_whole_insect - COM_body;

% maximum height of COM
dmax = max(COM_whole_insect(:,3));
%--------------------------------------------------------------------------
% compute moment of inertia for combined body parts, rotational plus
% translationalI_spin_total_legs
I_spin_legs = I_spin_legs';     % in units of kg m^2 already

% remove outliers, then smooth
I_spin_body = filloutliers(I_spin_body,'pchip',"movmedian",smoothwindow);
I_spin_body = smooth(I_spin_body,smoothwindow,'rloess');
I_spin_legs = filloutliers(I_spin_legs,'pchip',"movmedian",smoothwindow);
I_spin_legs = smooth(I_spin_legs,smoothwindow,'rloess');

I_spin_total = I_spin_legs + I_spin_body;

%--------------------------------------------------------------------------
% compute potential energy vs time

PE = mass_body_only*g*COM_height;           % PE due to body only 
PE_body = PE;

% add contributions due to legs
PE(~isnan(PE_legs)) = PE(~isnan(PE_legs))+ PE_legs(~isnan(PE_legs));

PE = PE*1e-3; % convert PE units to [J] (height: 1e-3 mm to m)
PE_body = PE_body*1e-3; % same

% PE error bars [J] for now due to uncertainty in tracking 
% (*1e-3 converts Ztol from mm to m)
PE_errorbar = mass_whole_insect*g*Ztol*1e-3;

[PEmax,idxmax]  = max(PE);                   % maximum PE & index = idxmax 
[PE_bodymax,~]  = max(PE_body);              % maximum body-only PE
[PEmin,~]       = min(PE);                   % minimum PE 
[PE_bodymin,~]  = min(PE_body);              % minimum body-only PE 

% position of COM on ground when PE is at its maximum
COM_whole_insect_PEmax = COM_whole_insect(idxmax,:);
% replace NaNs after righting with the effective pendulum pivot point on
% ground
pivot_pt_nan = isnan(pivot_pt_on_ground(:,1));
pivot_pt_on_ground(pivot_pt_nan,:) = repmat(COM_whole_insect_PEmax(1:2),...
    sum(pivot_pt_nan),1);

%%--------------------------------------------------------------------------
% compute the kinetic energy

%----------------------
% compute COM velocity to use in computing the body COM kinetic energy
polyorder = 2;             % order of local polynomial fit
KE_COM = zeros(nframe,1);
for k = 1:3
    COM_body_vel(:,k) =movingslope(COM_body(:,k),smoothwindow,polyorder,dt);
    KE_COM = KE_COM + 0.5*mass_body_only*COM_body_vel(:,k).*COM_body_vel(:,k);
end
% convert speed^2 from (mm/s)^2 to (m/2)^2 in KE
KE_COM = KE_COM*1e-6; 

%----------------------
% total translational KE for all legs, including all directions
KE_legs = zeros(nframe,1);
KE_each_leg = zeros(nframe,6);
for j = 1:6
    % compute the speed of each leg segment
    femurvel2 = 0; tibiavel2 = 0;
    for kk = 1:3   % compute each kth xyz componet of jth leg's velocity 
                   % sum vx^2 + vy^2 + vz^2
        femurvel = movingslope(femur_COM(:,j,kk),smoothwindow,polyorder,dt);
        femurvel2 = femurvel.*femurvel + femurvel2;
        tibiavel = movingslope(tibia_COM(:,j,kk),smoothwindow,polyorder,dt);
        tibiavel2 = tibiavel.*tibiavel + tibiavel2;
    end
    KE_each_leg(:,j) = 0.5*(femur_coxa_mass(j).*femurvel2 + tibia_tarsus_mass(j).*tibiavel2);
    KE_legs = KE_legs + KE_each_leg(:,j);
end

% convert speed^2 from (mm/s)^2 to (m/2)^2 in KE
KE_legs = KE_legs*1e-6;

%-----------------
% compute total translational KE
KE_trans = KE_COM + KE_legs;      % total translational KE

%----------------------
% compute spin rotational KE for the legs (note this will automatically be
% in units of [J] because we use angular velocity in rad/s and moment of
% inertia in kg m^2
KE_spin_legs = zeros(nframe,1);
KE_spin_per_leg = zeros(nframe,6);
femur_ang_speed = nan(nframe,6); tibia_ang_speed = nan(nframe,6);
% compute unit vector that is normal to each leg -- used in calculating the
% angular momentum of each leg below
for i = 1:nframe
    for j = 1:6
        leg_normal(i,j,1:3) = cross(femur_line_segment(i,j,:),tibia_line_segment(i,j,:));
        leg_normal(i,j,1:3) = leg_normal(i,j,1:3)/vecnorm(leg_normal(i,j,1:3),2,3);
    end
end
% compute the angular speed of the femur and tibia in [rad/s]
for i = 2:nframe
    for j = 1:6
        femur1 = squeeze(femur_line_segment(i-1,j,:));
        femur2 = squeeze(femur_line_segment(i,j,:));
        tibia1 = squeeze(tibia_line_segment(i-1,j,:));
        tibia2 = squeeze(tibia_line_segment(i,j,:));
        % use formula to avoid instability in acos(dot) that gives complex
        % values; computes angle in radians, so angular speed is [rad/s]
        femur_ang_speed(i,j) = atan2(norm(cross(femur1,femur2)),dot(femur1,femur2))/dt;
        tibia_ang_speed(i,j) = atan2(norm(cross(tibia1,tibia2)),dot(tibia1,tibia2))/dt;
    end
end
% set first frame of the speeds, which are not defined, = second frame
femur_ang_speed(1,:) = femur_ang_speed(2,:); 
tibia_ang_speed(1,:) = tibia_ang_speed(2,:);

% compute spin kinetic energies
for j = 1:6
    % compute mean leg angular speeds
    leg_ang_speed(:,j) = 0.5*(femur_ang_speed(:,j)+tibia_ang_speed(:,j));

    KE_spin_per_leg(:,j) = 0.5*I_spin_each_leg(:,j).*leg_ang_speed(:,j).*leg_ang_speed(:,j);
    KE_spin_legs = KE_spin_legs + KE_spin_per_leg(:,j);   
end
KE_spin_legs = smooth(KE_spin_legs,smoothwindow,'rloess');

% set first frame of the KE, which is not defined, = second frame
KE_spin_per_leg(1,:) = KE_spin_per_leg(2,:);
for j = 1:6
    KE_spin_per_leg(:,j) = smooth(KE_spin_per_leg(:,j),smoothwindow,'rloess');
end

%---------------------
% reminder: all KE_spin are already in units of kg m^2

KE_spin_total = KE_spin_body + KE_spin_legs;

%--------------------
% compute total KE
KE = KE_trans + KE_spin_total;    % total KE--trans+spin, body + legs

%%--------------------------------------------------------------------------
% get KE of orbital motion of the effective pendulum

% angular velocity = speed/radius, where we approximate radius as dmax, the
% maximum height of the orbital
for k = 1:3
    whole_insect_COM_vel(:,k) =movingslope(COM_whole_insect(:,k),smoothwindow,polyorder,dt);
end
% unit vector along local orbital velocity
whole_insect_COM_vel_norm = whole_insect_COM_vel./vecnorm(whole_insect_COM_vel,2,2);

% the COM and dmax are both in units of mm here
w_orbital = vecnorm(whole_insect_COM_vel,2,2)/dmax; % units of rad/s

dmax = dmax*1e-3;     % rescale from mm to m for KE calculation

% use parallel axis theorem to get KE for both spin and orbital motions
I_pendulum = (I_spin_total + mass_whole_insect*dmax^2);
% ratio of elements in I_pendulum
I_rot_ratio = I_spin_total./(mass_whole_insect*dmax^2);

% total pendulum KE in [J]
KE_pendulum = 0.5*I_pendulum.*w_orbital.*w_orbital;

%% --------------------------------------------------------------------------
% find active overturning times from dorsal-ventral axis
% orientation relative to z-axis:
fig_gettimes = figure;
plot(time,dorsal_ventral_axis(:,3),'k-',...
    time,zeros(size(time)),'k--');
title({"Click add points,del to remove","shift-, right-, or double-click","adds a final point & exits"})
[time_active,~] = getpts;
% get indices of the start and end times for righting
[~,startidx] = min(abs(time-time_active(1)));
[~,endidx] = min(abs(time-time_active(2)));
% close the figure (these times are shown in the first panel figure below)
close(fig_gettimes);

%% compute Delta PE to overcome barrier/right at time t (max PE at index 
% idxmax -- only plot values during righting
Delta_PE = PEmax-PE;

% orbital KE for the overturning sequence for plotting convenience
KE_pendulum_righting = KE_pendulum;

% compute the righting ratio, RN_dynamic
RN_dynamic = KE_pendulum./Delta_PE;
idxRN100 = find(RN_dynamic>0.99,1,'first'); % find when it first reaches 100%
idxRN75 = find(RN_dynamic>0.749,1,'first'); % find when it first reaches 75%
idxRN50 = find(RN_dynamic>0.49,1,'first'); % find when it first reaches 50%
idxRN25 = find(RN_dynamic>0.249,1,'first'); % find when it first reaches 25%

%% compute summary stats for moment of inertia
% compute the moment of inertia of the body before and during righting
% using the same time interval as righting
I_body_before = mean(I_spin_body(firstframe:startidx),'omitnan');
I_body_before_SD = std(I_spin_body(firstframe:startidx),'omitnan');
I_body_active_mean = mean(I_spin_body(startidx:idxmax),'omitnan');
% compute 95% CI using bootstrapping
bootdata = I_spin_body(startidx:idxmax);
I_body_active_CI = bootci(nboot,{@(v)mean(v,'omitnan'),bootdata});
I_body_active_CI = 0.5*(I_body_active_CI(2)-I_body_active_CI(1));

% compute the median body axis of rotation in the body frame during righting
temp = rotation_axis_body_frame(startidx:idxmax,:);
temp = rmoutliers(temp);  % remove outliers so they don't bias the mean
rotation_axis_body_frame_during = mean(temp,'omitnan'); clear temp;
% normalize
rotation_axis_body_frame_during = rotation_axis_body_frame_during/...
    norm(rotation_axis_body_frame_during);
[rot_azi,rot_elev,~] = cart2sph(rotation_axis_body_frame_during(1),...
    rotation_axis_body_frame_during(2),rotation_axis_body_frame_during(3));

I_total_before = mean(I_spin_total(firstframe:startidx));
I_total_during = mean(I_spin_total(startidx:idxmax));

% compute mean pendulum moment of inertia during active overturning
I_pendulum_active_mean = mean(I_pendulum(startidx:idxmax),'omitnan');
% compute 95% CI using bootstrapping
bootdata = I_pendulum(startidx:idxmax);
I_pendulum_active_CI = bootci(nboot,{@(v)mean(v,'omitnan'),bootdata});
I_pendulum_active_CI = 0.5*(I_pendulum_active_CI(2)-I_pendulum_active_CI(1));


% compute and display the fraction of I_rot due to the legs -- entire
% sequence
fraction_I_spin_legs = I_spin_legs./I_spin_total;
fraction_I_spin_legs_all_mean = mean(fraction_I_spin_legs(:),'omitnan');
% compute 95% CI using bootstrapping
bootdata = fraction_I_spin_legs;
fraction_I_spin_legs_all_CI = bootci(nboot,{@(v)mean(v,'omitnan'),bootdata});
fraction_I_spin_legs_all_CI = 0.5*(fraction_I_spin_legs_all_CI(2)-fraction_I_spin_legs_all_CI(1));

%% compute torque & reaction force
% compute the pendulum's effective orbital angular acceleration
% w_orbital has units of [rad/s] already
angular_accel_orbital = movingslope(w_orbital,smoothwindow,polyorder,dt);

% convert to net torque in Newton*meters
torque_orbital_net = angular_accel_orbital.*I_pendulum;

% torque_orbital_net = torque_propel_net + torque_gravity
% correct for torque due to gravity
% torque_gravity = (distance from COM on ground to COM at max PE)*MG
% correct for torque due to gravity
% factor of 1e-3* converts mm to m

torque_gravity = mass_whole_insect*g*...
    1e-3*vecnorm(COM_whole_insect_PEmax(1:2)-COM_whole_insect(:,1:2),2,2);

% explanation of signs used to compute torque due to leg
% propeling = pushing or pulling forces:
% at times before the tipping point, negative net torque decreases
% the angular velocity;  when falling, it increases it
% at times before the tipping point, the torque due to gravity slows
% down orbital speed (negative torque); after it's positive

torque_propel_net = +torque_orbital_net(1:idxmax)...
    + torque_gravity(1:idxmax);
torque_propel_net = [torque_propel_net; torque_orbital_net(idxmax+1:end)...
    - torque_gravity(idxmax+1:end)];

% estimate & plot the required propel net force converted to microN
% for comparison with locomotion propelling forces measured in the
% literature for insects

% to figure out the signs here, we need to know whether the angular
% acceleration corresponds to an increasing or decreasing upward or
% downward angular velocity

% estimated moment arm, r_perp, for net propelling force = max distance
% between the tarsal ground contact points and the pivot point

% first compute the maximum distance on the ground from the mean leg ground 
% support point to the COM at maximum PE; factor of 1e-3 converts mm to m
r_perp = zeros(nframe,1);
% find each tarsus in contact with the ground (ignore those values for
% which there are NaN coordinates)

% get direction in XY plane that's perpendicular to whole insect com
% velocity in-plane component
% this ensures dot(whole_insect_COM_vel_perp,whole_insect_COM_vel_norm) = 0
whole_insect_COM_vel_perp(:,1) =  whole_insect_COM_vel_norm(:,2);
whole_insect_COM_vel_perp(:,2) = -whole_insect_COM_vel_norm(:,1);

for i = 1:nframe
    for j = 1:6
        if tarsuscoords(i,3*j) <=Ztol_support_plane && sum(isnan(tarsuscoords(i,(1:3)+(j-1)*3))) == 0
            % compute vector from the tarsus ground contact point to the 
            % pivot point
            r_leg_ground = tarsuscoords(i,(1:2)+(j-1)*3) - ...
                pivot_pt_on_ground(i,:);               
            % add absolute value of the component perpendicular to the 
            % orbital velocity to r_perp & convert from mm to m
            r_perp(i) = r_perp(i) + 1e-3*...
                abs(dot(r_leg_ground,whole_insect_COM_vel_perp(i,1:2)));
        end
    end
end
r_perp(r_perp < 1e-3*3*Ztol) = NaN;    % ignore points close to zero
r_perp = fillmissing(r_perp,'pchip');
r_perp = filloutliers(r_perp,'pchip',"movmedian",smoothwindow);
r_perp = smooth(r_perp,smoothwindow,'rloess');

r_perp_leg = r_perp./ntarsicontactpts;
r_perp_leg_active_mean = mean(r_perp_leg(firstframe:idxmax),'omitnan');
% compute 95% CI using bootstrapping
bootdata = r_perp_leg(firstframe:idxmax);
r_perp_leg_active_CI = bootci(nboot,{@(v)mean(v,'omitnan'),bootdata});
r_perp_leg_active_CI = 0.5*(r_perp_leg_active_CI(2)-r_perp_leg_active_CI(1));

% factor of 1e6 converts force units from Newton to microNewton
force_propel_net = 1e6*torque_propel_net./r_perp;

% divide by body weight
force_propel_net = force_propel_net/(g*mass_whole_insect*1e6);

%%%%%%%%%%%%%%%%%%%%%%%%%%%%%%%%%%%%%%%%%%%%%%%%%%%%%%%%%%%%%%%%%%%%%%%%%%%
%%--------------------------------------------------------------------------
% plot dorsal ventral z component (for orientation), PE and KE vs time
figure('Color','white','Position',[10,10,500,1000]);
tiledlayout(5,1,"TileSpacing",'none','Padding','compact');
%--------------------------
% plot dorsal ventral z component (shows when/whether'EdgeColor','white' it is righted or not)
nexttile;

plot(time,dorsal_ventral_axis(:,3),'k-'); hold on;
yticks([-1 0 1]); grid on;

set(gca,'XTick',[]); % xlabel('time (s)');
ylabel({'Z_{dv}'});
xlim([time(firstframe) time(lastframe)]); ylim([-1.2 1.2]);

% plot vertical gray line when dv axis becomes horizontal
plot([time(idxdvz0),time(idxdvz0)],[-1.2 1.2],'-','Color',htmlgray);

% plot vertical dark gray line when PE is at its maximum
plot([time(idxmax),time(idxmax)],[-1.2 1.2],'-','Color',0.5*htmlgray,'LineWidth',0.5);

xregion(time_active(1),time_active(2),'FaceColor',htmlgray,'EdgeColor','white');

%-----------------
% plot PE and pendulum KE
nexttile;
 
plot(time,1e6*PE,'k-'); hold on;

% fill region between PE and its max value with light gray
coord_up = [time',1e6*PEmax*ones(size(PE))];
coord_low = [time',1e6*PE];
coord_combine = [coord_up;flipud(coord_low)];
fill(coord_combine(:,1),coord_combine(:,2),1.2*htmlgray,'FaceAlpha',0.5);

plot(time,1e6*PE_body,'k--'); % plot body-only PE

plot(time,1e6*KE_pendulum,'-','Color',bluelines);

set(gca,'XTick',[]); %xlabel('time (s)'); 
xlim([time(firstframe) time(lastframe)]);
ylabel('Energy (10^{-6} J)'); 
ylim([0 1.2]);

% plot vertical gray line when dv axis becomes horizontal
Yminmax = ylim;
plot([time(idxdvz0),time(idxdvz0)],Yminmax,'-','Color',htmlgray);

% plot vertical dark gray line when PE is at its maximum
plot([time(idxmax),time(idxmax)],Yminmax,'-','Color',0.5*htmlgray,'LineWidth',0.5);

xregion(time_active(1),time_active(2),'FaceColor',htmlgray,'EdgeColor','white');

% plot vertical arrow to show how Delta PE is defined
idxarrow = firstframe + round((startidx-firstframe)/2);
plot(time(idxarrow),1e6*PEmax,'k^','MarkerSize',4,'MarkerFaceColor','k');
plot(time(idxarrow)*[1,1],[1e6*PE(idxarrow),...
    1e6*PEmax],'k:','LineWidth',1.5);
text(time(idxarrow),1e6*0.5*(PEmax +PE(idxarrow)),' \DeltaPE(t)');

legend({'PE_{total}','','PE_{body}','KE_{avail}','','','',''},...
    'Location','eastoutside',...
    'Orientation','vertical','NumColumns',1,'FontSize',10); legend boxoff;
%------------------
% plot available KE from z-motion and Delta_PE required to right 
nexttile;

% plot KE and Delta_PE during overturning
yyaxis left; hold off;

% only show KE and Delta_PE for the values when they are comparable
plot(time,1e6*Delta_PE,'k:','LineWidth',1.5); hold on;
plot(time,1e6*KE_pendulum,'-','Color',bluelines);

xlim([time(firstframe) time(lastframe)]);
ylim([0,1e6*3*max(KE_pendulum)]);
set(gca,'XTick',[]); %xlabel('time (s)','FontSize',11);
ylabel('Energy (10^{-6} J)');

% make both axes black
ax = gca; ax.YAxis(1).Color = 'k'; ax.YAxis(2).Color = 'k';

% plot vertical gray line when dv axis becomes horizontal
Yminmax = ylim;
plot([time(idxdvz0),time(idxdvz0)],Yminmax,'-','Color',htmlgray);

% plot vertical dark gray line when PE is at its maximum
plot([time(idxmax),time(idxmax)],Yminmax,'-','Color',0.5*htmlgray,'LineWidth',0.5);

% shade righting region in gray
xregion(time_active(1),time_active(2),'FaceColor',htmlgray,'EdgeColor','white');

% plot the righting ratio, RN_dynamic as %
yyaxis right; hold off;
plot(time,100*RN_dynamic,'-.','Color',0.65*greenlines,'LineWidth',1);hold on;

ylabel('RN (%)'); 
xlim([time(firstframe) time(lastframe)]);
ylim([0 110]);

legend({'\Delta PE','KE_{avail}','','','','RN'},...
    'Location','eastoutside',...
    'Orientation','vertical','FontSize',10); legend boxoff;
%-------------------
% plot angular speed vs time
nexttile;

% compute the body's roll and pitch component of angular velocity [Hz=rev/s]
% w_spin in the manuscript
w_spin = sqrt(vecnorm(angular_velocity.*Xbody,2,2).^2 + ...
    vecnorm(angular_velocity.*Ybody,2,2).^2);
w_spin = filloutliers(w_spin,"pchip","movmedian",smoothwindow);
w_spin = smooth(w_spin,smoothwindow,'rloess');

% compute the ratio of the body spin roll & pitch to orbital angular speeds
% use bootstrapping for the mean & 95% CI
% active overturning

% first use bootstrapping to get nboot samples of the mean for each, then
% take their ratio & use to get mean & CI of the ratio
bootdata = w_spin(startidx:idxmax);
w_spin_meanarray = bootstrp(nboot,@mean,bootdata);
bootdata = w_orbital(startidx:idxmax)/(2*pi); % get in same units as w_spin
w_orbital_meanarray = bootstrp(nboot,@mean,bootdata);
w_spin_to_w_orbital = w_spin_meanarray./w_orbital_meanarray;
wspin_to_worbital_active_mean = mean(w_spin_to_w_orbital,'omitnan');
bootdata = w_spin_to_w_orbital;
wspin_to_worbital_active_CI = bootci(nboot,{@(v)mean(v,'omitnan'),bootdata});
wspin_to_worbital_active_CI = 0.5*(wspin_to_worbital_active_CI(2)-wspin_to_worbital_active_CI(1));

% before overturning -- same procedure
bootdata = w_spin(firstframe:startidx);
w_spin_meanarray = bootstrp(nboot,@mean,bootdata);
bootdata = w_orbital(firstframe:startidx)/(2*pi); % get in same units as w_spin
w_orbital_meanarray = bootstrp(nboot,@mean,bootdata);
w_spin_to_w_orbital = w_spin_meanarray./w_orbital_meanarray;
wspin_to_worbital_before_mean = mean(w_spin_to_w_orbital,'omitnan');
bootdata = w_spin_to_w_orbital;
wspin_to_worbital_before_CI = bootci(nboot,{@(v)mean(v,'omitnan'),bootdata});
wspin_to_worbital_before_CI = 0.5*(wspin_to_worbital_before_CI(2)-wspin_to_worbital_before_CI(1));

% plot them finally!

plot(time,w_orbital/(2*pi),'-','Color',bluelines); hold on;
plot(time,w_spin,'--','Color',redlines);

% xlabel('time (s)'); 
set(gca,'XTick',[]);
xlim([time(firstframe) time(lastframe)]);
ylabel('angular speed (Hz)');
ylim([0,1.15*max(max(w_orbital/(2*pi),[],'omitnan'),...
    max(w_spin,[],'omitnan'))])

% plot vertical gray line when dv axis becomes horizontal
Yminmax = ylim;
plot([time(idxdvz0),time(idxdvz0)],[Yminmax(1),Yminmax(2)],'-','Color',htmlgray);

% plot vertical dark gray line when PE is at its maximum
plot([time(idxmax),time(idxmax)],Yminmax,'-','Color',0.5*htmlgray,'LineWidth',0.5);

% shade righting region in gray
xregion(time_active(1),time_active(2),'FaceColor',htmlgray,'EdgeColor','white');

legend({'\omega_{orbital}' '\omega_{spin}' '' '' ''},...
    'Location','eastoutside',...
    'Orientation','vertical','FontSize',10);
legend boxoff;

%-------------------------------------------------------------------------
% plot the legs' tarsus z body frame positions vs time
nexttile;

% the tarsi are in this order:  
% 'right foreleg','right midleg','right hindleg' = 1 2 3
% 'left hindleg','left midleg''left foreleg' = 4 5 6
% plot in this order:  1 2 3 6 5 4
plotorder = [1 2 3 6 5 4];
% same linestyle each side, same color each pair of contralateral legs
% tarsuscolors = lines(6);
tarsuscolors = [0 0 0;0 0 0;0 0 0;darkgray;darkgray;darkgray];
% tarsuslinestyles = {'-' '--' '-.' '-.' '--' '-'};
tarsuslinestyles = {'-' '--' '-.' '-.' '--' '-'};

for j = 1:6
    % tarsus positions
    k = plotorder(j);
    plot(time,tarsus_zbody(:,k),tarsuslinestyles{k},'Color',tarsuscolors(k,:)); hold on;
end
maxtarsz = max(tarsus_zbody(firstframe:lastframe,:),[],'omitnan');
maxtarsz = max(tarsus_zbody(:))+0.5;
mintarsz = min(tarsus_zbody(firstframe:lastframe,:),[],'omitnan'); 
mintarsz = min(mintarsz(:));mintarsz = min([ -1 mintarsz])-0.5;
ylabel({'z_{tarsus} (mm)' '(body frame)'}); ylim([min(-1,mintarsz) maxtarsz]);

xlim([time(firstframe) time(lastframe)]); 
xlabel('time (s)'); % set(gca,'XTick',[]);

% plot vertical gray line when dv axis becomes horizontal
Yminmax = ylim;
plot([time(idxdvz0),time(idxdvz0)],[Yminmax(1),Yminmax(2)],'-','Color',htmlgray);

% plot vertical dark gray line when PE is at its maximum
plot([time(idxmax),time(idxmax)],Yminmax,'-','Color',0.5*htmlgray,'LineWidth',0.5);

% shade righting region in gray
xregion(time_active(1),time_active(2),'FaceColor',htmlgray,'EdgeColor','white');

legend({'right foreleg','right midleg','right hindleg',...
        'left foreleg','left midleg','left hindleg',...
    '' '' '' ''},'Location','eastoutside',...
    'Orientation','vertical','NumColumns',1,'FontSize',10); legend boxoff

%--------------------
% save as figure file
panel1filename = fullfile(pathstr,[filename(1:23),'_5P1.png']);
saveas(gcf,panel1filename);

%%%%%%%%%%%%%%%%%%%%%%%%%%%%%%%%%%%%%%%%%%%%%%%%%%%%%%%%%%%%%%%%%%%%%%%%%%%
%% Supplemental figures / supporting figures

%% Plots of moment of inertia
figure('Color','white','Position',[10,10,500,500]);
tiledlayout(2,1,"TileSpacing",'tight','Padding','loose');

%-----------------------
% plot all components of moment of inertia vs time
nexttile;

%----------
plot(time,1e10*I_pendulum,'-','Color',bluelines,'LineWidth',1.0);hold on;

% plot means
plot(time(firstframe:startidx),1e10*mean(I_pendulum(firstframe:startidx),'omitnan')*...
    ones(size(time(firstframe:startidx))),'-','Color',pastelblue,'LineWidth',1.0);
    hold on;
plot(time(startidx:idxmax),1e10*mean(I_pendulum(startidx:idxmax),'omitnan')*...
    ones(size(time(startidx:idxmax))),...
    '-','Color',pastelblue,'LineWidth',1.0);hold on;
%----------
plot(time,1e10*I_spin_total,'-',...
    'Color',orangelines,'LineWidth',1.0);

% plot means
plot(time(firstframe:startidx),1e10*mean(I_spin_total(firstframe:startidx),'omitnan')*...
    ones(size(time(firstframe:startidx))),...
    '-','Color',pastelorange,'LineWidth',1.0);hold on;
plot(time(startidx:idxmax),1e10*mean(I_spin_total(startidx:idxmax),'omitnan')*...
    ones(size(time(startidx:idxmax))),...
    '-','Color',pastelorange,'LineWidth',1.0);hold on;
%----------
plot(time,1e10*I_spin_body,'-','Color',redlines,'LineWidth',1.0);
% plot means
plot(time(firstframe:startidx),1e10*mean(I_spin_body(firstframe:startidx),'omitnan')*...
    ones(size(time(firstframe:startidx))),...
    '-','Color',pastelred,'LineWidth',1.0);hold on;
plot(time(startidx:idxmax),1e10*mean(I_spin_body(startidx:idxmax),'omitnan')*...
    ones(size(time(startidx:idxmax))),...
    '-','Color',pastelred,'LineWidth',1.0);hold on;

xlabel('time (s)');
ylabel({'moment of' 'inertia (10^{-10} kg m^2)'});
xlim([time(firstframe) time(lastframe)]);
ylim([0 9])

% plot vertical gray line when dv axis becomes horizontal
Yminmax = ylim;
plot([time(idxdvz0),time(idxdvz0)],[Yminmax(1),Yminmax(2)],'-','Color',htmlgray);

% plot vertical dark gray line when PE is at its maximum
plot([time(idxmax),time(idxmax)],Yminmax,'-','Color',0.5*htmlgray,'LineWidth',0.5);

% shade righting region in gray
xregion(time_active(1),time_active(2),'FaceColor',htmlgray,'EdgeColor','white');

% draw a box around the region replotted in the next figure
rectangle('Position',[time_active(1),0,...
    (time_active(2)-time_active(1)),1.1*1e10*Iyaw],...
    'LineStyle','--');

% draw pale lines at means for each region

legend({'pendulum','mean','' 'total spin','mean','' 'total body','mean','', ...
    '' '' ''},'Location','eastoutside',...
    'Orientation','vertical','NumColumns',1,'FontSize',10); legend boxoff;
%-----------------------
% plot body moment of inertia vs time during righting
nexttile;

plot(time,1e10*I_spin_body,'-','Color',redlines,'LineWidth',1.0); hold on;
plot(time,1e10*Iroll*ones(size(time)),"--",...
    time,1e10*Ipitch*ones(size(time)),"-.",...
    time,1e10*Iyaw*ones(size(time)),":",'Color',redlines,'LineWidth',1.5);
plot(time(startidx:idxmax),1e10*ones(size(time(startidx:idxmax)))*mean(I_spin_body(startidx:idxmax),'omitnan'),":",...
    'Color',pastelred,'LineWidth',1.5);

xlim([time_active(1),time_active(2)]);
ylim([0,1.1*1e10*Iyaw]);

% plot vertical gray line when dv axis becomes horizontal
Yminmax = ylim;
plot([time(idxdvz0),time(idxdvz0)],[Yminmax(1),Yminmax(2)],'-','Color',htmlgray);

% plot vertical dark gray line when PE is at its maximum
plot([time(idxmax),time(idxmax)],Yminmax,'-','Color',0.5*htmlgray,'LineWidth',0.5);

% shade righting region in gray
xregion(time_active(1),time_active(2),'FaceColor',htmlgray,'EdgeColor','white');

xlabel('time (s)');ylabel({'body spin moment' ' of inertia (10^{-10} kg m^2)'});
legend({'total','roll' 'pitch' 'yaw' 'mean' ...
    '' '' ''},'Location','eastoutside',...
    'Orientation','vertical','NumColumns',1,'FontSize',10); legend boxoff;
%--------------------
% save as figure file
FigS6filename = fullfile(pathstr,[filename(1:23),'_S6.png']);
saveas(gcf,FigS6filename);

%%-------------------------------------------------------------------------
% plot the tarsi zbody, ground contact pts, & force vs time
figure('Color','white','Position',[525,50,500,800]);

tiledlayout(3,1,"TileSpacing",'none',"Padding","compact");

%------------------------
% plot number of contact points vs time
nexttile;

plot(time,(ncontactpts-nbodycontactpts),'k-'); 
hold on;
plot(time,nbodycontactpts,'k-.');

xlim([time(firstframe) time(lastframe)]); 
set(gca,'XTick',[]); %xlabel('time (s)'); 
ylim([0 max(9,1.2*max(ncontactpts(:)))]);
ylabel({'number of ground';'contact points'});
% plot vertical gray line when dv axis becomes horizontal; gray box during
% overturning
Yminmax = ylim;
plot([time(idxdvz0),time(idxdvz0)],[Yminmax(1),Yminmax(2)],'-','Color',htmlgray);

% plot vertical dark gray line when PE is at its maximum
plot([time(idxmax),time(idxmax)],Yminmax,'-','Color',0.5*htmlgray,'LineWidth',0.5);

% shade righting region in gray
xregion(time_active(1),time_active(2),'FaceColor',htmlgray,'EdgeColor','white');

legend({'tarsi + tibia-femur joints' 'body' '' '' ''},...
    'Location','northwest','Orientation','horizontal','FontSize',10); legend boxoff;
% ------------------------------------------------------------------------
% plot stability margin vs time
nexttile;

% compute percent stability margin
SM = 100*stability_margin./ideal_stability_margin;
SM = filloutliers(SM,'linear');
SM = min(100,SM);             % clip at 100% (digitizing/rounding errors)

plot(time,SM,'k-'); hold on;
plot(time,zeros(size(time)),'k-','LineWidth',0.35);  % SM = 0

xlim([time(firstframe) time(lastframe)]); 
ylim([min(0,1.1*min(SM)),120]);

set(gca,'XTick',[]); %xlabel('time (s)');  
ylabel('stability margin (%)');

% plot vertical gray line when dv axis becomes horizontal
Yminmax = ylim;
plot([time(idxdvz0),time(idxdvz0)],[Yminmax(1),Yminmax(2)],'-','Color',htmlgray);

% plot vertical dark gray line when PE is at its maximum
plot([time(idxmax),time(idxmax)],Yminmax,'-','Color',0.5*htmlgray,'LineWidth',0.5);

% shade righting region in gray
xregion(time_active(1),time_active(2),'FaceColor',htmlgray,'EdgeColor','white');
%-----------------------

%------------------------
% plot mean reaction force/leg vs time
nexttile;
yyaxis left;
plot(time,force_propel_net,'k-'); hold on;
ylabel({'mean force/leg' '(body weight)'});

force_active_mean = mean(force_propel_net(firstframe:idxmax),'omitnan');
% compute 95% CI using bootstrapping
bootdata = force_propel_net(firstframe:idxmax);
force_active_CI = bootci(nboot,{@(v)mean(v,'omitnan'),bootdata});
force_active_CI = 0.5*(force_active_CI(2)-force_active_CI(1));
% compute 95 percentile
force_95 = prctile(force_propel_net(firstframe:lastframe),95);

plot(time(firstframe:idxmax),ones(size(time(firstframe:idxmax)))*force_active_mean,'k:','LineWidth',1.5);

force_max = max(force_propel_net(firstframe:lastframe),[],"omitnan");
force_min = min(force_propel_net,[],"omitnan");
ylim([min(0,1.05*force_min) 1.05*force_max])
xlim([time(firstframe) time(lastframe)]); xlabel('time (s)','FontSize',11);

% plot of reaction torque based on angular acceleration vs time
yyaxis right;

torque_active_mean = mean(torque_propel_net(firstframe:idxmax),'omitnan');
% compute 95% CI using bootstrapping
bootdata = torque_propel_net(firstframe:idxmax);
torque_active_CI = bootci(nboot,{@(v)mean(v,'omitnan'),bootdata});
torque_active_CI = 0.5*(torque_active_CI(2)-torque_active_CI(1));

% compute 95 percentile
torque_95 = prctile(torque_propel_net(firstframe:lastframe),95);

plot(time,torque_propel_net,'--','Color',greenlines,'LineWidth',0.75); hold on;
plot(time(firstframe:idxmax),ones(size(time(firstframe:idxmax)))*torque_active_mean,':','LineWidth',1.5,'Color',greenlines);
ylabel('total torque (Nm)');

torque_max = max(torque_propel_net,[],"omitnan");
torque_min = min(torque_propel_net,[],"omitnan");
ylim([min(0,1.05*torque_min) 1.05*torque_max]);

xlim([time(firstframe) time(lastframe)]);

% make both axes black
ax = gca; ax.YAxis(1).Color = 'k'; ax.YAxis(2).Color = 'k';

% 
%%align zero for left and right
yyaxis right; ylimr = get(gca,'Ylim');ratio = ylimr(1)/ylimr(2);
yyaxis left; yliml = get(gca,'Ylim');
if yliml(2)*ratio<yliml(1)
    set(gca,'Ylim',[yliml(2)*ratio yliml(2)])
else
    set(gca,'Ylim',[yliml(1) yliml(1)/ratio])
end
% plot thin line at 0
plot(time,zeros(size(time)),'k-','LineWidth',0.25);
% plot vertical gray line when dv axis becomes horizontal
Yminmax = ylim;
plot([time(idxdvz0),time(idxdvz0)],Yminmax,'-','Color',htmlgray);
% plot vertical dark gray line when PE is at its maximum
plot([time(idxmax),time(idxmax)],Yminmax,'-','Color',0.5*htmlgray,'LineWidth',0.5);
% shade righting region in gray
xregion(time_active(1),time_active(2),'FaceColor',htmlgray,'EdgeColor','white');


legend({'force/leg' 'mean' '' '' ''  'torque' 'mean' }); legend boxoff;

%--------------------
% save as figure file
FigS5filename = fullfile(pathstr,[filename(1:23),'_S5.png']);
saveas(gcf,FigS5filename);

%%%%%%%%%%%%%%%%%%%%%%%%%%%%%%%%%%%%%%%%%%%%%%%%%%%%%%%%%%%%%%%%%%%%%%%%%%%
%% compute & save summary statistics
outarray = []; 

fprintf('video analyzed = %s, frames %d to %d analyzed\n',file,firstframe,lastframe);

outarray = {'active overturning (s)' abs(time_active(1)-time_active(2))};

% PE
outarray = [outarray;{'PEmax(J)',PEmax}];
outarray = [outarray;{'PEmin(J)',PEmin}];
outarray = [outarray;{'PE body-only max(J)',PE_bodymax}];
outarray = [outarray;{'PE body-only min(J)',PE_bodymin}];
outarray = [outarray;{'PE error bar (J)',PE_errorbar}];
outarray = [outarray;{'PE body error bar (J)',PE_errorbar*mass_body_only/mass_whole_insect}];

% Moments of inertia -- mean, SD during active overturning
outarray = [outarray;{'I_pendulum, active mean (kg m^2)' I_pendulum_active_mean}];
outarray = [outarray;{'I_pendulum, active CI (kg m^2)' I_pendulum_active_CI}];
outarray = [outarray;{'I_body, active mean (kg m^2)' I_body_active_mean}];
outarray = [outarray;{'I_body, active CI (kg m^2)' I_body_active_CI}];
% compute error bar in body spin moment of inertia using the orientation of
% the rotation axis and the error bars in each component of I_body
I_body_err = dot(rotation_axis_body_frame_during,[0.11*Iroll,0.01*Ipitch,0.03*Iyaw]);
outarray = [outarray;{'I_body, righting err (kg m^2)' I_body_err}];

% fraction of I_{total spin} due to legs
outarray = [outarray;{'%% Ispin legs mean (all)',100*fraction_I_spin_legs_all_mean}];
outarray = [outarray;{'%% Ispin legs CI (all)',100*fraction_I_spin_legs_all_CI}];

% mean body rotational axis orientation during righting:
% [x,y,z] coords in body frame =
outarray = [outarray;{'mean rot axis',sprintf('[%1.2f,%1.2f,%1.2f]',rotation_axis_body_frame_during(1),...
    rotation_axis_body_frame_during(2),rotation_axis_body_frame_during(3))}];
% angle between cranial-caudal axis and rotation axis during righting
mean_spin_angle = acos(cos(rot_azi)*cos(rot_elev))*180/pi;
% we only care about the angle with the axis, not whether it's with the +
% or - direction
mean_spin_angle = min(mean_spin_angle,abs(180-abs(mean_spin_angle)));
outarray = [outarray;{'mean spin angle(deg)',acos(cos(rot_azi)*cos(rot_elev))*180/pi }];

outarray = [outarray;{'wspin/worbital before mean',100*wspin_to_worbital_before_mean}];
outarray = [outarray;{'wspin/worbital before mean CI',100*wspin_to_worbital_before_CI}];
outarray = [outarray;{'wspin/worbital active mean',100*wspin_to_worbital_active_mean}];
outarray = [outarray;{'wspin/worbital active CI',100*wspin_to_worbital_active_CI}];

% force, torque, moment arm means from start until flipping point
outarray = [outarray;{'95 percentile force/leg(BW)',force_95}];
outarray = [outarray;{'force/leg active mean (BW)',force_active_mean}];
outarray = [outarray;{'force/leg active CI (BW)',force_active_CI}];

outarray = [outarray;{'95 percentile  torque(Nm)',torque_95}];
outarray = [outarray;{'torque active mean (Nm)',torque_active_mean}];
outarray = [outarray;{'torque/leg active CI (Nm)',torque_active_CI}];

outarray = [outarray;{'moment arm active mean (mm)',1000*r_perp_leg_active_mean}];
outarray = [outarray;{'moment arm active CI (mm)',1000*r_perp_leg_active_CI}];

%SM (mean, SD) (start to dv z horizontal)
outarray = [outarray;{'SM all to apex mean(%)',mean(SM(firstframe:idxmax),'omitnan')}];
% compute 95% CI using bootstrapping
bootdata = SM(firstframe:idxmax);
SM_CI = bootci(nboot,{@(v)mean(v,'omitnan'),bootdata});
SM_CI = 0.5*(SM_CI(2)-SM_CI(1));

outarray = [outarray;{'SM all to apex CI(%)',SM_CI}];

%number tarsi contacts (mean, SD) (start to dv z horizontal)
alllegcontacts = ncontactpts(firstframe:idxmax)-nbodycontactpts(firstframe:idxmax);
alllegcontacts_mean = mean(alllegcontacts,'omitnan');
% compute 95% CI using bootstrapping
bootdata = alllegcontacts;
alllegcontacts_CI = bootci(nboot,{@(v)mean(v,'omitnan'),bootdata});
alllegcontacts_CI = 0.5*(alllegcontacts_CI(2)-alllegcontacts_CI(1));

SDalllegcontacts = std((ncontactpts(firstframe:idxmax)-nbodycontactpts(firstframe:idxmax)),'omitnan');
outarray = [outarray;{'number leg contacts all mean',alllegcontacts_mean}];
outarray = [outarray;{'number leg contacts all CI',alllegcontacts_CI}];
outarray = [outarray;{'ground plane SD(mm)',std(Z_ground)}];
outarray = [outarray;{'body min z < 0 SD(mm)',std(bodymeshminz(bodymeshminz<0))}];

outarray = [outarray;{'time before apex RN=0.25 (ms)',idxmax-idxRN25}];
outarray = [outarray;{'time before apex RN=0.5 (ms)',idxmax-idxRN50}];
outarray = [outarray;{'time before apex RN=0.75 (ms)',idxmax-idxRN75}];
outarray = [outarray;{'time before apex RN=1 (ms)',idxmax-idxRN100}];
outarray = [outarray;{'time before apex Z_{dv}=0 (ms)',idxmax-idxdvz0}];
outarray = [outarray;{'time at apex (ms)',idxmax}];

%%%%%%%%%%%%%%%%%%%%%%%%%%%%%%%%%%%%%%%%%%%%%%%%%%%%%%%%%%%%%%%%%%%%%%%%%%%
%% Save workspace data
matdatafilename = fullfile(pathstr,[filename(1:23),'_anal.mat']);
save(matdatafilename);

% save data as a spreadsheet
spreadsheetfilename = fullfile(pathstr,[filename(1:23),'.xlsx']);
xlswrite(spreadsheetfilename,outarray);

%%**************************************************************************
% function:	p_poly_dist
% Description:	distance from point to polygon whose vertices are specified by the
%              vectors xv and yv
% Input:  
%    x - point's x coordinate
%    y - point's y coordinate
%    xv - vector of polygon vertices x coordinates
%    yv - vector of polygon vertices x coordinates
% Output: 
%    d - distance from point to polygon (defined as a minimal distance from 
%        point to any of polygon's ribs, positive if the point is outside the
%        polygon and negative otherwise)
%    x_poly: x coordinate of the point in the polygon closest to x,y
%    y_poly: y coordinate of the point in the polygon closest to x,y
%
% Routines: p_poly_dist.m
% Revision history:
%    03/31/2008 - return the point of the polygon closest to x,y
%               - added the test for the case where a polygon rib is 
%                 either horizontal or vertical. From Eric Schmitz.
%               - Changes by Alejandro Weinstein
%    7/9/2006  - case when all projections are outside of polygon ribs
%    23/5/2004 - created by Michael Yoshpe 
% https://www.mathworks.com/matlabcentral/fileexchange/19398-distance-from-a-point-to-polygon
%  Alejandro Weinstein (2021). Distance from a point to polygon
%  (https://www.mathworks.com/matlabcentral/fileexchange/19398-distance-from-a-point-to-polygon),
%  MATLAB Central File Exchange. Retrieved October 9, 2021.
%**************************************************************************
function [d,x_poly,y_poly] = p_poly_dist(x, y, xv, yv) 
% If (xv,yv) is not closed, close it.
xv = xv(:);
yv = yv(:);
Nv = length(xv);
if ((xv(1) ~= xv(Nv)) || (yv(1) ~= yv(Nv)))
    xv = [xv ; xv(1)];
    yv = [yv ; yv(1)];
%     Nv = Nv + 1;
end
% linear parameters of segments that connect the vertices
% Ax + By + C = 0
A = -diff(yv);
B =  diff(xv);
C = yv(2:end).*xv(1:end-1) - xv(2:end).*yv(1:end-1);
% find the projection of point (x,y) on each rib
AB = 1./(A.^2 + B.^2);
vv = (A*x+B*y+C);
xp = x - (A.*AB).*vv;
yp = y - (B.*AB).*vv;
% Test for the case where a polygon rib is 
% either horizontal or vertical. From Eric Schmitz
id = find(diff(xv)==0);
xp(id)=xv(id);
clear id
id = find(diff(yv)==0);
yp(id)=yv(id);
% find all cases where projected point is inside the segment
idx_x = (((xp>=xv(1:end-1)) & (xp<=xv(2:end))) | ((xp>=xv(2:end)) & (xp<=xv(1:end-1))));
idx_y = (((yp>=yv(1:end-1)) & (yp<=yv(2:end))) | ((yp>=yv(2:end)) & (yp<=yv(1:end-1))));
idx = idx_x & idx_y;
% distance from point (x,y) to the vertices
dv = sqrt((xv(1:end-1)-x).^2 + (yv(1:end-1)-y).^2);
if(~any(idx)) % all projections are outside of polygon ribs
   [d,I] = min(dv);
   x_poly = xv(I);
   y_poly = yv(I);
else
   % distance from point (x,y) to the projection on ribs
   dp = sqrt((xp(idx)-x).^2 + (yp(idx)-y).^2);
   [min_dv,I1] = min(dv);
   [min_dp,I2] = min(dp);
   [d,I] = min([min_dv min_dp]);
   if I==1, %the closest point is one of the vertices
       x_poly = xv(I1);
       y_poly = yv(I1);
   elseif I==2, %the closest point is one of the projections
       idxs = find(idx);
       x_poly = xp(idxs(I2));
       y_poly = yp(idxs(I2));
   end
end
if ~inpolygon(x, y, xv, yv)    % SM < 0 if the COM is not inside the polygon
   d = -d;
end
end

%**************************************************************************
function [rotm] = get_rot_matrix(sourcepts,targetpts)
    % takes two sets of xyz axes at times t1 and t2,computes the quaternion
    % required to rotate between the two, uses it to get the rotation
    % matrix required to perform the rotation

    %  sourcepts: 3xN matrix whose columns are the coordinates of N source points.
    %  targetpts: 3xN matrix whose columns are the coordinates of N target points.

    % create quaternion to rotate the sourcepts into target pts
    [regParams,Bfit,ErrorStats]=absor(sourcepts,targetpts);
    temp = regParams.q;
    quat = quaternion(temp(1),temp(2),temp(3),temp(4));
      
    % compute the rotation matrix to rotate from sourcpts to target pts
    rotm = quat2rotm(quat);
end
%**************************************************************************
function [w] = get_ang_vel(sourcepts,targetpts,dt)
    % takes two sets of xyz axes at times t1 and t2,computes the quaternion
    % required to rotate between the two, uses it to get the angular
    % velocity during the rotation

    %  sourcepts: 3xN matrix whose columns are the coordinates of N source points.
    %  targetpts: 3xN matrix whose columns are the coordinates of N target points.

    % create quaternion to rotate the sourcepts into target pts
    [regParams,Bfit,ErrorStats]=absor(sourcepts,targetpts);
    temp = regParams.q;
    quat = quaternion(temp(1),temp(2),temp(3),temp(4));
    
    % compute the angular velocity from the quaternion
    w = angvel(quat,dt,'frame');       % w = [rad/s]
    wmag = vecnorm(w,2,2);          % magnitude
    
    w = w * 1/(2*pi);               % w = = [Hz] = [rev/s]
    wmag = vecnorm(w,2,2);          % magnitude
    
end
%% ************************************************************************
function [regParams,Bfit,ErrorStats]=absor(A,B,varargin)
% https://www.mathworks.com/matlabcentral/fileexchange/26186-absolute-orientation-horn-s-method?s_tid=srchtitle
%ABSOR is a tool for finding the rotation -- and optionally also the
%scaling and translation -- that best maps one collection of point coordinates to 
%another in a least squares sense. It is based on Horn's quaternion-based method. The
%function works for both 2D and 3D coordinates, and also gives the option of weighting the 
%coordinates non-uniformly. The code avoids for-loops so as to maximize speed.
%
%DESCRIPTION:
%
%As input data, one has
%
%  A: a 2xN or 3xN matrix whos columns are the coordinates of N source points.
%  B: a 2xN or 3xN matrix whos columns are the coordinates of N target points.
%
%The basic syntax
%
%     [regParams,Bfit,ErrorStats]=absor(A,B) 
%
%solves the unweighted/unscaled registration problem
%
%           min. sum_i ||R*A(:,i) + t - B(:,i)||^2 
%
%for unknown rotation matrix R and unknown translation vector t. 
%
%ABSOR can also solve the more general problem
%
%           min. sum_i w(i)*||s*R*A(:,i) + t - B(:,i)||^2   
%
%where s>=0 is an unknown global scale factor to be estimated along with R and t
%and w is a user-supplied N-vector of  weights. One can include/exclude any 
%combination of s, w, and translation t in the problem formulation. Which 
%parameters participate is controlled using the syntax,
%
%  [regParams,Bfit,ErrorStats]=absor(A,B,'param1',value1,'param2',value2,...)
%
%with parameter/value pair options,
%
%  'doScale' -  Boolean flag. If TRUE, the global scale factor, s, is included.
%               Otherwise, it is assumed that s=1. Default=FALSE.
%
%  'doTrans' -  Boolean flag. If TRUE, the translation, t, is included. Otherwise,
%               zero translation is assumed. Default=TRUE.
%               
%  'weights' - The length N-vector of weights, w. Default, no weighting.
%
%
%OUTPUTS:
%
%
% regParams: structure output with estimated registration parameters,
%
%     regParams.R:   The estimated rotation matrix, R
%     regParams.t:   The estimated translation vector, t
%     regParams.s:   The estimated scale factor.
%     regParams.M:   Homogenous coordinate transform matrix [s*R,t;[0 0 ... 1]].
%
%     For 3D problems, the structure includes
%
%        regParams.q:   A unit quaternion [q0 qx qy qz] corresponding to R and
%                       signed to satisfy max(q)=max(abs(q))>0
%
%     For 2D problems, it includes
%
%        regParams.theta: the counter-clockwise rotation angle about the
%                         2D origin
%
%
%  Bfit: The rotation, translation, and scaling (as applicable) of A that 
%        best matches B.
%
%
% ErrorStats: structure output with error statistics. In particular,
%             defining err(i)=sqrt(w(i))*norm( Bfit(:,i)-B(:,i) ), 
%             it contains
%
%      ErrorStats.errlsq = norm(err) 
%      ErrorStats.errmax = max(err) 
%
%
%
%%%%%%%%%%%%%%%%%%%%%%%%%%%%%%
% Author: Matt Jacobson
% Copyright, Xoran Technologies, Inc.  http://www.xorantech.com


%%Input option processing and set up

 options.doScale = 0;
 options.doTrans = 1;
 options.weights = []; 
    
for ii=1:2:length(varargin)
    param=varargin{ii};
    val=varargin{ii+1};
    if strcmpi(param,'doScale'), 
        options.doScale=val;
    elseif strcmpi(param,'weights')
        options.weights=val;
    elseif strcmpi(param,'doTrans')
        options.doTrans=val;
    else
       error(['Option ''' param ''' not recognized']);
    end
end

 doScale = options.doScale;
 doTrans = options.doTrans;
 weights = options.weights;             

if ~verLessThan('matlab','9.1') %implicit expansion introduced
 matmvec=@(M,v)M-v;
 mattvec=@(M,v)M.*v;
elseif ~verLessThan('matlab','7.4')
 matmvec=@(M,v) bsxfun(@minus,M,v); %matrix-minus-vector
 mattvec=@(M,v) bsxfun(@times,M,v); %matrix-times-vector
else
 matmvec=@matmvecHandle; %matrix-minus-vector
 mattvec=@mattvecHandle; %matrix-times-vector
end

dimension=size(A,1);

if dimension~=size(B,1),
    error 'The number of points to be registered must be the same'
end


%%Centering/weighting of input data

if doTrans
                    if isempty(weights)

                     sumwts=1;   

                     lc=mean(A,2);  rc=mean(B,2);  %Centroids
                     left  = matmvec(A,lc); %Center coordinates at centroids
                     right = matmvec(B,rc); 

                    else

                      sumwts=sum(weights);

                      weights=full(weights)/sumwts;
                         weights=weights(:);
                      sqrtwts=sqrt(weights.');


                      lc=A*weights;   rc=B*weights; %weighted centroids

                      left  = matmvec(A,lc);  
                         left  = mattvec(left,sqrtwts);
                      right = matmvec(B,rc);
                        right = mattvec(right,sqrtwts);  

                    end
                    
else
    
                    if isempty(weights)

                     sumwts=1;   
                     left=A; right=B;

                    else

                      sumwts=sum(weights);

                      weights=full(weights)/sumwts;
                         weights=weights(:);
                      sqrtwts=sqrt(weights.');
 
                       left  = mattvec(A,sqrtwts);
                       right = mattvec(B,sqrtwts);  

                     end
                    
end

M=left*right.';


%%Compute rotation matrix

switch dimension

    case 2
    
        
        Nxx=M(1)+M(4); Nyx=M(3)-M(2);

        N=[Nxx   Nyx;...
           Nyx   -Nxx];

        [V,D]=eig(N);

        [trash,emax]=max(real(  diag(D)  )); emax=emax(1);

        q=V(:,emax); %Gets eigenvector corresponding to maximum eigenvalue
        q=real(q);   %Get rid of imaginary part caused by numerical error


        q=q*sign(q(2)+(q(2)>=0)); %Sign ambiguity
        q=q./norm(q);

        R11=q(1)^2-q(2)^2;
        R21=prod(q)*2;

        R=[R11 -R21;R21 R11]; %map to orthogonal matrix


        
        
    case 3
        
        [Sxx,Syx,Szx,  Sxy,Syy,Szy,   Sxz,Syz,Szz]=dealr(M(:));

        N=[(Sxx+Syy+Szz)  (Syz-Szy)      (Szx-Sxz)      (Sxy-Syx);...
           (Syz-Szy)      (Sxx-Syy-Szz)  (Sxy+Syx)      (Szx+Sxz);...
           (Szx-Sxz)      (Sxy+Syx)     (-Sxx+Syy-Szz)  (Syz+Szy);...
           (Sxy-Syx)      (Szx+Sxz)      (Syz+Szy)      (-Sxx-Syy+Szz)];

        [V,D]=eig(N);

        [trash,emax]=max(real(  diag(D)  )); emax=emax(1);

        q=V(:,emax); %Gets eigenvector corresponding to maximum eigenvalue
        q=real(q);   %Get rid of imaginary part caused by numerical error

        [trash,ii]=max(abs(q)); sgn=sign(q(ii(1)));
         q=q*sgn; %Sign ambiguity

        %map to orthogonal matrix
        
        quat=q(:);
        nrm=norm(quat);
        if ~nrm
         disp 'Quaternion distribution is 0'    
        end

        quat=quat./norm(quat);

        q0=quat(1);
        qx=quat(2); 
        qy=quat(3); 
        qz=quat(4);
        v =quat(2:4);


        Z=[q0 -qz qy;...
           qz q0 -qx;...
          -qy qx  q0 ];

        R=v*v.' + Z^2;

    otherwise
        error 'Points must be either 2D or 3D'

end

%%

if doScale
   
     summ = @(M) sum(M(:));
  
     sss=summ( right.*(R*left))/summ(left.^2);
     if doTrans
      t=rc-R*(lc*sss);
     else
      t=zeros(dimension,1);   
     end
     
else
    
    sss=1;
    if doTrans
     t=rc-R*lc;
    else
      t=zeros(dimension,1);  
    end
 
end


regParams.R=R;
regParams.t=t;
regParams.s=sss;


if dimension==2
    
    regParams.M=[sss*R,t;[0 0 1]];
    regParams.theta=atan2(q(2),q(1))*360/pi;
    
else%dimension=3
    
    regParams.M=[sss*R,t;[0 0 0 1]];
    regParams.q=q/norm(q);
    
end

if nargout>1
    
    Bfit=(sss*R)*A;
    
    if doTrans
     Bfit=matmvec(Bfit,-t);
    end   
end

if nargout>2
    
    l2norm = @(M,dim) sqrt(sum(M.^2,dim));

    err=l2norm(Bfit-B,1);
    
    if ~isempty(weights), err=err.*sqrtwts; end

    ErrorStats.errlsq=norm(err)*sqrt(sumwts); %unnormalize the weights
    ErrorStats.errmax=max(err);
    
   
end
    

end
%%----------------------------
function M=matmvecHandle(M,v)
%Matrix-minus-vector

    for ii=1:size(M,1)
     M(ii,:)=M(ii,:)-v(ii);
    end
end
%%----------------------------
function M=mattvecHandle(M,v)
%Matrix-times-vector

    for ii=1:size(M,1)
     M(ii,:)=M(ii,:).*v;
    end
end 
%%----------------------------
function varargout=dealr(v)
  varargout=num2cell(v);     
end
%%*************************************************************************
function [bodymesh,coxacoord,cranialcoord,caudalcoord] = ...
    getbodymesh(instar)

    % get the mesh body-only 3D model from file and its properties
    if instar == 3
        bodyonly = stlread('SLFBody3rdinstar.stl'); %% Load STL mesh
    elseif instar == 4
        bodyonly = stlread('SLFBody4thinstar.stl'); %% Load STL mesh
    else
        bodyonly = stlread('SLFBodystandard.stl'); %% Load STL mesh
    end
    % get xyz points and connectivity
    bodyPoints = bodyonly.Points;
    bodyConnect = bodyonly.ConnectivityList;
    % the body mesh's COM is at the origin

    rotm = roty(4-180);           % rotate so its cranial-caudal axis = +X
    % the -4 deg is so this is the snout-caudal end axis, which is what we track
    %  L-R lateral axis = +Y & ventral-to-dorsal axis = +Z
    
    bodyPointsrot = rotm*bodyPoints';      % now perform 3D rotation on mesh
    bodyPointsrot = bodyPointsrot';        % restore correct format
    
    % make an updated mesh
    bodymesh = triangulation(bodyConnect,bodyPointsrot);

    % get the COM, moment of inertia tensor and principal axes
    RBP=RigidBodyParams(bodymesh);
 
    % make an updated mesh
    body_rot = triangulation(bodyConnect,bodyPointsrot);
    
    % get the legs insert onto the body at the coxa on the rotated model
    L1 = [2.17,-1.05,0.426];
    L2 = [0.933,-1.58,0.630];
    L3 = [-0.59,-0.755,1.10];
    L4 = [-0.59,0.755,1.10];
    L5 = [0.93,1.58,0.630];
    L6 = [2.17,+1.05,0.426];
    
    coxacoord = [L1;L2;L3;L4;L5;L6];
    cranialcoord = [5.2031,0,0.2184];
    caudalcoord  = [-3.64,0,0.4296]; 
end
%% ************************************************************************
function Dvec = movingslope(vec,supportlength,modelorder,dt)
% movingslope: estimate local slope for a sequence of points, using a sliding window
% usage: Dvec = movingslope(vec)
% usage: Dvec = movingslope(vec,supportlength)
% usage: Dvec = movingslope(vec,supportlength,modelorder)
% usage: Dvec = movingslope(vec,supportlength,modelorder,dt)
%
%
% movingslope uses filter to determine the slope of a curve stored
% as an equally (unit) spaced sequence of points. A patch is applied
% at each end where filter will have problems. A non-unit spacing
% can be supplied.
%
% Note that with a 3 point window and equally spaced data sequence,
% this code should be similar to gradient. However, with wider
% windows this tool will be more robust to noisy data sequences.
%
% arguments: (input)
%  vec - row of column vector, to be differentiated. vec must be of
%        length at least 2
%
%  supportlength - (OPTIONAL) scalar integer - defines the number of
%        points used for the moving window. supportlength may be no
%        more than the length of vec.
%
%        supportlength must be at least 2, but no more than length(vec)
%
%        If supportlength is an odd number, then the sliding window
%        will be central. If it is an even number, then the window
%        will be slid backwards by one element. Thus a 2 point window
%        will result in a backwards differences used, except at the
%        very first point, where a forward difference will be used.
%
%        DEFAULT: supportlength = 3
%
%  modelorder - (OPTIONAL) - scalar - Defines the order of the windowed
%        model used to estimate the slope. When model order is 1, the
%        model is a linear one. If modelorder is less than supportlength-1.
%        then the sliding window will be a regression one. If modelorder
%        is equal to supportlength-1, then the window will result in a
%        sliding Lagrange interpolant.
%
%        modelorder must be at least 1, but not exceeding
%        min(10,supportlength-1)
%
%        DEFAULT: modelorder = 1
%
%  dt - (OPTIONAL) - scalar - spacing for sequences which do not have
%        a unit spacing.
%
%        DEFAULT: dt = 1
%
% arguments: (output)
%  Dvec = vector of derivative estimates, Dvec will be of the same size
%        and shape as is vec.
% 
%
% Example:
%  Estimate the first derivative using a 7 point window with first through
%  fourth order models in the sliding window. Note that the higher order
%  approximations provide better accuracy on this curve with no noise.
%  
%  t = 0:.1:1;
%  vec = exp(t);
%
%  Dvec = movingslope(vec,7,1,.1)
%  Dvec =
%  Columns 1 through 7
%    1.3657  1.3657  1.3657  1.3657  1.5093  1.668  1.8435
%  Columns 8 through 11
%    2.0373  2.0373  2.0373  2.0373
%
%  Dvec = movingslope(vec,7,2,.1)
%  Dvec =
%  Columns 1 through 7
%    0.95747 1.0935  1.2296  1.3657  1.5093  1.668  1.8435
%  Columns 8 through 11
%    2.0373  2.2403  2.4433  2.6463
%
%  Dvec = movingslope(vec,7,3,.1)
%  Dvec =
%  Columns 1 through 7
%    1.0027  1.1049  1.2206  1.3498  1.4918  1.6487  1.8221
%  Columns 8 through 11
%    2.0137  2.2268  2.4602  2.7138
%
%  Dvec = movingslope(vec,7,4,.1)
%  Dvec =
%    Columns 1 through 7
%    0.99988 1.1052  1.2214  1.3498  1.4918  1.6487  1.8221
%  Columns 8 through 11
%    2.0137  2.2255  2.4597  2.7181
%
%
% Example:
%  Estimate the slope of a noisy curve, using a locally quadratic
%  approximation. In this case, use a straight line so that we know
%  the true slope should be 1. Use a wide window, since we have
%  noisy data.
%  
%  t = 0:100;
%  vec = t + randn(size(t));
%  Dvec = movingslope(vec,10,2,1)
%  mean(Dvec)
%  ans = 
%     1.0013
%  std(Dvec)
%  ans =
%     0.10598
%
%  By way of comparison, gradient gives a much noisier estimate
%  of the slope of this curve.
%
%  std(gradient(vec))
%  ans =
%     0.69847
%
%
% Example:
%  As a time test, generate random data vector of length 500000.
%  Compute the slopes using a window of width 10.
%
%  vec = rand(1,500000);
%  tic
%  Dvec = movingslope(vec,10,2);
%  toc
%
%  Elapsed time is 0.626021 seconds.
%
%
% See also: gradient
%
%
% Author: John D'Errico
%
% Release: 1.0
% Release date: 10/19/07
% Cite as:  John D'Errico (2021). Movingslope
%  (https://www.mathworks.com/matlabcentral/fileexchange/16997-movingslope),
%  MATLAB Central File Exchange. Retrieved August 11, 2021.
%**************************************************************************
% how long is vec? is it a vector?
if (nargin==0)
  help movingslope
  return
end
if ~isvector(vec)
  error('vec must be a row or column vector')
end
n = length(vec);                    % n = number of input data points

% -------------------------------------------------------------------------
% supply defaults
if (nargin<4) || isempty(dt)
  dt = 1;
end
if (nargin<3) || isempty(modelorder)
  modelorder = 1;
end
if (nargin<2) || isempty(supportlength)
  supportlength = 3;
end
% check the parameters for problems
if (length(supportlength)~=1) || (supportlength<=1) || (supportlength>n) || (supportlength~=floor(supportlength))
  error('supportlength must be a scalar integer, >= 2, and no more than length(vec)')
end
if (length(modelorder)~=1) || (modelorder<1) || (modelorder>min(10,supportlength-1)) || (modelorder~=floor(modelorder))
  error('modelorder must be a scalar integer, >= 1, and no more than min(10,supportlength-1)')
end
if (length(dt)~=1) || (dt<0)
  error('dt must be a positive scalar numeric variable')
end

% -------------------------------------------------------------------------
% now build the filter coefficients to estimate the slope
if mod(supportlength,2) == 1
  parity = 1; % odd parity
else
  parity = 0;
end
s = (supportlength-parity)/2;       % s = approx. half window size 
t = ((-s+1-parity):s)';             % t = window for filter calculations
coef = getcoef(t,supportlength,modelorder);

% -------------------------------------------------------------------------
% Apply the filter to the entire vector
f = filter(-coef,1,vec);
Dvec = zeros(size(vec));
Dvec(s+(1:(n-supportlength+1))) = f(supportlength:end);
% --------------------------
% patch each end
vec = vec(:);
for i = 1:s
  % patch the first few points (1 to s) so they only involve fits to the
  % the appropriate points
  t = (1:supportlength)' - i;
  coef = getcoef(t,supportlength,modelorder);
  
  Dvec(i) = coef*vec(1:supportlength);
  
  % patch the end points ((n - s + 1) to n) so they only involve fits over
  % the appropriate points
  if i<(s + parity)
    t = (1:supportlength)' - supportlength + i - 1;
    coef = getcoef(t,supportlength,modelorder);
    Dvec(n - i + 1) = coef*vec(n + (0:(supportlength-1)) + 1 - supportlength);
  end
end

% -------------------------------------------------------------------------
% scale by the supplied spacing
Dvec = Dvec/dt;
% all done
end % mainline end
% =========================================================
% subfunction, used to compute the filter coefficients
function coef = getcoef(t,supportlength,modelorder)
% Note: bsxfun would have worked here as well, but some people
% might not yet have that release of matlab.
A = repmat(t,1,modelorder+1).^repmat(0:modelorder,supportlength,1);
pinvA = pinv(A);
% we only need the linear term
coef = pinvA(2,:);
end % nested function end
%% ************************************************************************
% Rik (2023). point to line distance (https://github.com/thrynae/point_to_line_distance/releases/tag/1.3.2), GitHub. Retrieved October 30, 2023. 
function distance=point_to_line_distance(pt, v1, v2)
    %Calculate distance between a point and a line in 2D or 3D.
    % syntax:
    % distance = point_to_line(pt, v1, v2)
    % pt is a nx3 matrix with xyz coordinates for n points
    % v1 and v2 are vertices on the line (each 1x3)
    % d is a nx1 vector with the orthogonal distances
    %
    % 2D inputs are extended to 3D by setting all z-values to 0, all inputs should be either nx2 or nx3
    % (1x2 or 1x3 for v1 and v2).
    %
    % The actual calculation is a slightly edit version of this line (which only works for 3D points):
    % distance=norm(cross(v1-v2,pt-v2))/norm(v1-v2)
    %
    %  _______________________________________________________________________
    % | Compatibility | Windows 10  | Ubuntu 20.04 LTS | MacOS 10.15 Catalina |
    % |---------------|-------------|------------------|----------------------|
    % | ML R2020a     |  works      |  not tested      |  not tested          |
    % | ML R2018a     |  works      |  works           |  not tested          |
    % | ML R2015a     |  works      |  works           |  not tested          |
    % | ML R2011a     |  works      |  works           |  not tested          |
    % | ML 6.5 (R13)  |  works      |  not tested      |  not tested          |
    % | Octave 5.2.0  |  works      |  works           |  not tested          |
    % | Octave 4.4.1  |  works      |  not tested      |  works               |
    % """""""""""""""""""""""""""""""""""""""""""""""""""""""""""""""""""""""""
    %
    % Version: 1.3.2
    % Date:    2020-06-28
    % Author:  H.J. Wisselink
    % Licence: CC by-nc-sa 4.0 ( creativecommons.org/licenses/by-nc-sa/4.0 )
    % Email=  'h_j_wisselink*alumnus_utwente_nl';
    % Real_email = regexprep(Email,{'*','_'},{'@','.'})
    %check inputs
    if nargin~=3
        error('HJW:point_to_line_distance:nargin',...
            'Incorrect number of inputs, expected 3.');
    end
    if ~isnumeric(pt) || ~any(size(pt,2)==[2 3]) || any(size(pt)==0)
        error('HJW:point_to_line_distance:pt_type_size',...
            'First input (pt) is not numeric or has an incorrect shape.')
    end
    if ~isnumeric(v1) || numel(v1)~=size(pt,2)
        error('HJW:point_to_line_distance:v_type_size',...
            ['Second input (v1) is not numeric or has an incorrect size.',char(10),...
            'Expected 1x3 or 1x2, which should match the first input.']) %#ok<CHARTEN>
    end
    if ~isnumeric(v2) || numel(v2)~=size(pt,2)
        error('HJW:point_to_line_distance:v_type_size',...
            ['Third input (v2) is not numeric or has an incorrect size.',char(10),...
            'Expected 1x3 or 1x2, which should match the first input.']) %#ok<CHARTEN>
    end
    %prepare inputs
    v1=v1(:)';%force 1x3 or 1x2
    v2=v2(:)';%force 1x3 or 1x2
    if length(v1)==2,v1(3)=0;  end%extend 1x2 to 1x3 if needed
    if length(v2)==2,v2(3)=0;  end%extend 1x2 to 1x3 if needed
    if size(pt,2)==2,pt(1,3)=0;end%extend nx2 to nx3 if needed
    v1_ = repmat(v1,size(pt,1),1);
    v2_ = repmat(v2,size(pt,1),1);
    %actual calculation
    a = v1_ - v2_;
    b = pt - v2_;
    distance = sqrt(sum(cross(a,b,2).^2,2)) ./ sqrt(sum(a.^2,2));
    %this is equivalent to the following line for a single point
    %distance=norm(cross(v1-v2,pt-v2))/norm(v1-v2)
end

%% ************************************************************************
function [dydt,d2ydt2]...
    = quadratic_piecewise_fitter(t,y,inc_window)
    half_window = round(inc_window/2);
    idx_window = 1:numel(t);
    % similar to methods recommended in: Walker, J. A. (1998). Estimating velocities
    % and accelerations of animal locomotion: a simulation experiment comparing
    % numerical differentiation algorithms. Journal of Experimental Biology,
    % 201(7), 981-995. 
    % Fit model to data & plot fit to piecewise quadratic curve
    for jj = 1:numel(t)
        % compute window over which to compute orientation rate
        min_idx = max(1,(jj-half_window));
        max_idx = min(numel(t),(jj+half_window));
        % generate arrays with just this window of data
        t_win = t(min_idx:max_idx); y_win = y(min_idx:max_idx);
        % fit over this window
        [fitresult, gof] = fit(t_win, y_win, fittype('poly2'));
        % save results
        yfit(jj) = fitresult.p1*t(jj)*t(jj)...
            + fitresult.p2*t(jj) + fitresult.p3;
        dydt(jj) = 2*fitresult.p1*t(jj) + fitresult.p2;
        d2ydt2(jj) = 2*fitresult.p1; 
    end
    % done fitting
end
%% ************************************************************************
function a = anglebetweenthreepoints(A,B,C) 
% angle between line segments AB and BC in radians
    AB = A-B; CB = C-B;
    a = atan2(norm(cross(AB,CB)), dot(AB,CB));
end
%% ************************************************************************
function a = vecangle2pi(v1,v2,n)
% James Tursa at https://www.mathworks.com/matlabcentral/answers/501449-angle-betwen-two-3d-vectors-in-the-range-0-360-degree
% make sure v1 v2 and n are column vectors
% v1 and v2 are vectors, while n is a vector not in the same plane as these
% two
% correct format if it's 3 x 1 instead of 1 x 3
if size(v1,2) == 1; v1 = v1'; end
if size(v2,2) == 1; v2 = v2'; end
if size(n,2)  == 1; n  = n' ; end

x = cross(v1,v2);
c = sign(dot(x,n)) * norm(x);
a = atan2(c,dot(v1,v2));
end

%% ************************************************************************
function [num,Z_ground] = ...
    get_ground_plane(num,X_ground,Y_ground,Z_ground,nlandmarks,Ztol_gnd_plane)

    ground_plane_fit_method = 'SVD';
    % ground_plane_fit_method = 'pcfitplane';

    %----------------- estimate ground plane using
    % use singular value decomposition SVD to find the plane in best agreement with
    % ALL the support polygon points provided
    % based on Matt J response, Mathworks Answers:
    % https://www.mathworks.com/matlabcentral/answers/125251-3d-plane-with-more-than-3-points
    
    % subtract off initial ground plane centroid from body coordinates
    % Because the rotation could cause a change in the
    % centroid, we do this a second time -- see **rotated** below
    num(:,1:3:end) = num(:,1:3:end)  - mean(X_ground(:));
    num(:,2:3:end) = num(:,2:3:end)  - mean(Y_ground(:));
    num(:,3:3:end) = num(:,3:3:end)  - mean(Z_ground(:));
    
    % subtract off centroid from all ground points before SVD
    X_ground = X_ground - mean(X_ground(:));
    Y_ground = Y_ground - mean(Y_ground(:));
    Z_ground = Z_ground - mean(Z_ground(:));
    
    switch ground_plane_fit_method
        case 'SVD'
    
            % create array of points P used for SVD
            P=[X_ground(:),Y_ground(:), Z_ground(:)];
    
            meanPlanePts=mean(P);
            P=bsxfun(@minus,P,meanPlanePts);
            [U,S,V]=svd(P);
    
            %---------------------------------
            % find the groundplanenormal vector, which is perpendicular to the ground
            % plane -- at this point it could point up or down--will correct later
            groundplanenormal=V(:,end);                      % normal to the plane
    
            % Ground plane equation is ax + by + cz + d = 0
            a = groundplanenormal(1);
            b = groundplanenormal(2);
            c = groundplanenormal(3);
            d = - dot(groundplanenormal,meanPlanePts);
    
            % now make it a unit vector
            groundplanenormal = groundplanenormal/norm(groundplanenormal);
   
        case 'pcfitplane'
            % first create point cloud
            ptCloudGndPlane = pointCloud([X_ground Y_ground Z_ground]);
            
            maxDistance = Ztol_gnd_plane;
            % use robust RANSAC method to
            % fit data to the plane equation, ax + by + cz + d = 0.
            [model,inlierIndices,outlierIndices,meanError] = ...
                pcfitplane(ptCloudGndPlane,maxDistance);
            % plot point clouds
            ptcloudin = select(ptCloudGndPlane,inlierIndices);
            ptcloudin.Color = "black";
            ptcloudout = select(ptCloudGndPlane,outlierIndices);
            ptcloudout.Color = "red";
            figure;
            pcshow(ptcloudin);hold on; set(gcf,'color','w');set(gca,'color','w');
            pcshow(ptcloudout);set(gcf,'color','w');set(gca,'color','w');
            % show the fit model
            plot(model);set(gcf,'color','w');set(gca,'color','w');

            % get plane parameters in case we want them
            params = model.Parameters;
            a = params(1); b = params(2); c = params(3); d = params(4);
            % get ground plane normal
            groundplanenormal = model.Normal;           % normal to fit plane
            groundplanenormal = groundplanenormal';     % transpose for consistency with SVD
    end
    
    % make sure it points upward
    if groundplanenormal(3)<0
        groundplanenormal = -groundplanenormal;
    end
    
    % example for how to compute a sample point in the ground plane
    % P0(1:2) = 0; P0(3) = -d/c;
    
    % end of ground plane estimation
    %---------------------------------
    % Compute the rotation angle and axis to rotate by such that the
    % ground plane is horizontal & its normal (groundplanenormal) is vertical
    % We know that +z is up in the actual experiment.  Rotate so the
    % groundplanenormal = +z-axis = [0;0;1] & the ground plane is the xy-plane.
    
    rotangle = acos(dot(groundplanenormal,[0;0;1]));    % rotation angle
    
    % axis by which to rotate is perpendicular to +z and current normal
    rotaxis = cross(groundplanenormal,[0;0;1]);
    rotaxis= rotaxis';                         % transpose for use in rotate_3D
    
    %---------------------------------
    % rotate the ground plane points about rotaxis by rotangle(radians)
    if rotangle ~= 0
        % rotate ground points to correct orientation
        Rvec = [X_ground(:)';Y_ground(:)';Z_ground(:)']; % Rvec should have 3 rows
        Rvec = rotate_3D(Rvec,'any',rotangle,rotaxis','radian');
        X_ground(:) = Rvec(1,:);Y_ground(:) =Rvec(2,:);Z_ground(:) =Rvec(3,:);
    end
    
    %---------------------------------
    % rotate the groundplanenormal vector (should be +z-axis afterward)
    Rvec = rotate_3D(groundplanenormal,'any',rotangle,rotaxis','radian');
    groundplanenormal = Rvec';
    
    %---------------------------------
    % rotate the tracked body and leg coordinates
    % rotate each set of coordinates (requires fussy reformatting)
    for j = 1:nlandmarks        % loop over every body landmark xyz coord
        Rvec = [num(:,1+(j-1)*3)';num(:,2+(j-1)*3)';num(:,3+(j-1)*3)'];
        Rvec = rotate_3D(Rvec,'any',rotangle,rotaxis','radian');
        num(:,1+(j-1)*3) = Rvec(1,:);num(:,2+(j-1)*3)=Rvec(2,:);
        num(:,3+(j-1)*3) = Rvec(3,:);
    end
    %---------------------------------
    % print out the results
    fprintf('subtract off rotated ground point centroid along Z = %f\n',mean(Z_ground(:)));
    fprintf('ground plane normal = %f %f %f\n',groundplanenormal);

    %---------------------------------
    % subtract off **rotated** ground plane centroid from body coordinates
    % (this is not redundant because the rotation could cause a change in the
    % centroid of the ground plane points, which are not symmetrically
    % distributed about)
    % After this step, z = 0 should be the ground plane
    num(:,3:3:end) = num(:,3:3:end)  - mean(Z_ground(:));
    % zero Z_ground after this point
    Z_ground = Z_ground - mean(Z_ground(:));
    %---------------------------------
    % plot the ground support points & vertical arrow
    figure;
    plot3(X_ground,Y_ground,Z_ground,'o'); hold on;

    verticalarrow = groundplanenormal * max([X_ground;Y_ground;Z_ground]);
    quiver3(0,0,0,verticalarrow(1),verticalarrow(2),verticalarrow(3));

    % plot the ground plane as a transparent gray rectangular patch
    minX = min(X_ground); maxX = max(X_ground);
    minY = min(Y_ground); maxY = max(Y_ground);
    XGP = 1.1*[minX maxX maxX minX];
    YGP = 1.1*[minY minY maxY maxY];
    patch(XGP,YGP,[192/256,192/256,192/256],'FaceAlpha',0.5); hold on;

    axis equal; xlabel('X');ylabel('Y');zlabel('Z');
    title('xy plane = horizontal ground support plane; +z = vertical arrow')

end
%% ************************************************************************
```
